# Uncovering transposable element variants and their potential adaptive impact in urban populations of the malaria vector *Anopheles coluzzii*

**DOI:** 10.1101/2020.11.22.393231

**Authors:** Carlos Vargas-Chavez, Neil Michel Longo Pendy, Sandrine E. Nsango, Laura Aguilera, Diego Ayala, Josefa González

## Abstract

**Background:** *Anopheles coluzzii* is one of the primary vectors of human malaria in sub-Saharan Africa. Recently, it has colonized the main cities of Central Africa threatening vector control programs. The adaptation of *An. coluzzii* to urban environments is partly due to an increased tolerance to organic pollution and insecticides. While some of the molecular mechanisms for ecological adaptation, including chromosome rearrangements and introgressions, are known, the role of transposable elements (TEs) in the adaptive processes of this species has not been studied yet. To assess the role of TEs in rapid urban adaptation, the first step is to accurately annotate TE insertions in the genomes of natural populations collected in urban settings.

**Results:** We sequenced using long-reads six *An. coluzzii* genomes from natural breeding sites in two major Central Africa cities. We *de novo* annotated the complete set of TEs in these genomes and in an additional high-quality *An. coluzzii* genome available and identified 64 previously undescribed TE families. TEs were non-randomly distributed throughout the genome with significant differences in the number of insertions of several superfamilies across the studied genomes. We identified seven putatively active families with insertions near genes with functions related to vectorial capacity. Moreover, we identified several TE insertions providing promoter and transcription factor binding sites to insecticide resistance and immune-related genes.

**Conclusions:** The analysis of multiple genomes sequenced using long-read technologies allowed us to generate the most comprehensive TE annotations in this species to date. We identified several TE insertions that could potentially impact both genome architecture and the regulation of functionally relevant genes in *An. coluzzii*. These results provide a basis for future studies of the impact of TEs on the biology of *An. coluzzii*.

## Introduction

The deadly success of the malaria mosquito *Anopheles coluzzii* is rooted in its extraordinary ecological plasticity, inhabiting virtually every habitat in West and Central Africa where it spreads the human malaria parasite (1, 2). Noteworthy, the larvae of *An. coluzzii* exploit more disturbed and anthropogenic sites than its sister species *An. gambiae. An. coluzzii* exhibits a higher tolerance to salinity and organic pollution, and as a consequence, is the predominant species in coastal and urban areas (2-4). However, this mosquito not only has a greater resilience to ion-rich aquatic environments, but it has also become resistant to DDT and pyrethroid insecticides used for vector control (5). Actually, insecticide resistant populations of this malaria mosquito are present across its geographical range, driving *An. coluzzii* evolution across the continent (6, 7). The adaptive flexibility of this mosquito has been also highlighted by its rapid competence to expand its range of peak biting times in order to avoid insecticide treated bed-nets (8). This extraordinary adaptative capacity makes this malaria vector a threat for malaria control. Thus, elucidating the natural genetic variants underlying the ecological and the physiological responses to fluctuating environments in this species is key for its control.

At the molecular level, a variety of genetic mechanisms have been related back to the myriad of adaptation processes present in this mosquito. The most prominent and historically studied examples are chromosomal inversions (9, 10). *An. coluzzii* exhibits a large number of polymorphic chromosomal rearrangements (11, 12). Many of these inversions have been associated to environmental adaptation through environmental clines and/or correlation with specific climatic variables (10, 13), such as the inversion 2La associated with aridity tolerance capacity in adults (14, 15). Other types of rearrangements, such as gene duplications, have been involved in insecticide resistance. For example, the acetylcholinesterase (*Ace-1*) gene has been duplicated, maintaining at least a sensitive and a resistance copy, in order to counteract the fitness cost of the resistant phenotype (16-18). Moreover, a recent genome-wide analysis showed that genes containing copy number variants were enriched for insecticide functions (19). Other examples of gene selection due to anthropogenic activities have been found in genes related with detoxification or immunity, particularly in new colonized urban settings (3, 20-22). These adaptive processes have been repeated across West and Central African populations, reducing the efficacity of vector control measures (6). However, while several of the candidate genes responsible for the adaptive capacity of *An. coluzzii* have been identified, our knowledge of the genetic variants underlying differences in these genes lags behind. In particular, very little is known about natural variation in transposable element (TE) insertions in *An. coluzzii*.

TEs are key players in multiple adaptive processes across species, due to their capacity to generate a wide variety of mutations and to contribute to rapid responses to environmental changes (23, 24). TEs can disperse across the genome regulatory sequences such as promoters, enhancers, insulators, and repressive elements thus affecting nearby gene expression (25). Additionally, they can also act as substrates for ectopic recombination leading to structural mutations such as chromosomal rearrangements (26-28). However, TEs are often ignored when analyzing functional variants in genomes. This is because due to their repetitive nature, TE insertions are difficult to annotate and reads derived from TEs are often discarded in genome-wide analyses (29). Long-read sequencing techniques are needed to get a comprehensive view of TE variation in genomes, as these technique allow the annotation of TE insertions in the genome rather than inferring their position (30, 31).

Although TE insertions have been annotated genome-wide in several anopheline species including *An. coluzzii*, most studies to date have characterized the TE repertoire in a single genome for each species (32-40). To capture the full extent of TE natural variation and the potential consequences of TE insertions, it is necessary to evaluate multiple genomes in order to comprehensively assess diversity within a species (41-43). This becomes especially relevant when attempting to identify recent TE insertions and their effect in the genome structure and genome function, given that they might be restricted to local populations. So far, our knowledge of *An. coluzzii* genome variation due to TE insertions is limited to a few well-characterized families that have been found to vary across genomes (44-48).

In this work, we sequenced, using long-read technologies, and assembled the genomes of *An. coluzzii* larvae collected in six natural breeding sites in two major cities in Central Africa: Douala (Cameroon) and Libreville (Gabon). We performed a *de novo* TE annotation of the six newly assembled genomes, and we also annotated the previously available *An. coluzzii* genome from Yaoundé (Cameroon) (49). We identified 64 new anopheline TE families and showed that the availability of multiple genomes substantially improves the discovery of TE variants. We further analyzed individual TE insertions that could be acting as enhancers and promoters and that are located nearby genes with functions relevant for the vectorial capacity of the mosquitoes.

## Methods

### Sample collection and DNA isolation

We sampled *An. coluzzii* larvae in two cities of Central Africa: Libreville, Gabon, in January 2016 and Douala, Cameroon, in April 2018 (Additional File 1: Table S1). A systematic inspection of potential breeding sites was conducted to determine the presence of *Anopheles* larvae. We manually separated the anopheline from the culicine larvae based on morphological recognition and positioning of their bodies on or under the water surface (Robert, 2017). We collected immature 3rd and 4th stage larvae of *Anopheles* from water bodies using the standard dipping method (Service, 1993). Larvae were stored in 1.5 ml of absolute ethanol. After each daily sampling session, the samples were stored at -20 °C.

All the samples were PCR tested to differentiate *An. coluzzii* larvae from *An. gambiae* larvae before library preparation, using primers SINE200_F (TCGCCTTAGACCTTGCGTTA) and SINE200_R (CGCTTCAAGAATTCGAGATAC) (46). These primers target a single copy *SINE200* transposable element insertion that is fixed in *An. coluzzii* and absent in *An. gambiae*.

For PacBio sequencing, DNA from a single *An. coluzzii* larva from the *LBV11* site was extracted using the MagAttract HMW DNA extraction kit (Qiagen) following manufacturer’s instructions. Briefly, the larva was air-dried and lysed in 240 µl of buffer ATL (proteinase K added) shaking overnight at 56 ºC. Next, the DNA was isolated using the MagAttract magnetic beads and eluted twice in 50 µl of buffer AE. The DNA concentration was measured using a Qubit fluorometer.

For Nanopore sequencing, DNA from six larvae from each of the five breeding sites was extracted either with the QiaAMP UCP DNA kit (Qiagen) or MagAttract HMW DNA extraction kit (Qiagen). We performed individual larvae extractions as our objective was to use the minimum number of larvae possible to avoid the presence of excess polymorphisms that could affect the genome assembly. For the QiaAMP UCP DNA kit, we followed the manufacturer’s instructions. Each larva was air-dried and lysed in 200 µl of buffer AUT (proteinase K added) shaking overnight at 56 ºC, then DNA was isolated using a QIAamp UCP MinElute column and eluted twice in 25 µl of buffer AUE. For the MagAttract HMW DNA extraction kit, we followed manufacturer’s instructions but using lower buffer amounts to increase DNA concentration. Briefly, each larva was lysed in 120 µl of buffer ATL (proteinase K added) shaking overnight at 56 ºC, then DNA was isolated using the MagAttract magnetic beads and eluted twice in 25 µl of buffer AE. The DNA concentration was measured using a Qubit fluorometer. Both elutions of the same sample were mixed before library preparation. For Illumina sequencing, DNA from one larva from each of the six different breeding sites was extracted following the same extraction protocol as for Nanopore sequencing.

### Library preparation and sequencing

Quality control of the DNA sample for PacBio sequencing (Qubit, NanoDrop and Fragment analyzer) was performed at the Center for Genomic Research facility of the University of Liverpool prior to library preparation. The library was prepared by shearing DNA to obtain fragments of approximately 30 kb and sequenced on 2 SMRT cells using Sequel SMRT cell, 3.0 chemistry. Nanopore libraries were constructed using the Native Barcoding Expansion 1-12 (PCR-free) and the Ligation Sequencing Kit following manufacturer’s instructions. A minimum of 400 ng of DNA from each larva was used to start with the library workflow. For each breeding site, six larvae were barcoded, and equal amounts of each barcoded sample were pooled prior to sequencing. The samples from the same breeding site were ran in a single R9.4 flow cell in a 48-hour run, except for sample *DLA112* which was run in two flow cells. The DNA concentration was assessed during the whole procedure to ensure enough DNA was available for sequencing.

The quality control of the samples, library preparation and Illumina sequencing was performed at the Center for Genomic Research facility of the University of Liverpool. Low input libraries were prepared with the NEBNext Ultra II FS DNA library kit (300 bp inserts) on the Mosquito platform, using a 1/10 reduced volume protocol. Paired-end sequencing was performed on the Illumina Novaseq platform using S2 chemistry (2×150 bp).

### Genome Assemblies

The PacBio sequenced genome was assembled using *Canu* version 1.8 (130) with an estimated genome size of 250Mb and parameters: ‘*stopOnLowCoverage*=5, *corMinCoverage*=0, *correctedErrorRate*=0.105, *CorMhapFilterThreshold*=0.0000000002, *corMhapOptions=“--threshold 0*.*80 --num-hashes 512 --num-min-matches 3 --ordered-sketch-size 1000 --ordered-kmer-size 14 --min-olap-length 2000 --repeat-idf-scale 50*” *mhapMemory*=60g, *mhapBlockSize*=500, *ovlMerDistinct*=0.975’. The parameter stopOnLowCoverage was set to 5 to prevent fragmentation given that some of our samples had medium coverage. corMinCoverage was set to 0 to conserve the full length of the reads during the correction stage. correctedErrorRate was set to 0.105 following the recommendations in Canu’s manual for low coverage genome assemblies. All of the remaining parameters were set to reduce disk space and run time following the recommendations for repetitive genomes.Next, we identified and removed allelic variants using *purge_haplotigs* version 1.0.4 (131) with the “*-l 15 -m 100 -h 195*” parameters.

The Nanopore genomes were assembled using *Canu* version 1.8 using the same parameters as previously described, except for *correctedErrorRate* which was set to 0.16, followed by a round of polishing using *racon* version 1.3.3 (132), followed by *nanopolish* version 0.11.1 (133) and *pilon* version 1.23-0 (134) with the fix parameter set on ‘*bases*’. *Pilon* requires high coverage short-read data to perform the polishing and these data came from the aforementioned single larvae sequenced from each of the sites. Finally, *blobtools* version 1.1.1 (135) was used to remove contamination from all six genome assemblies taking into consideration fragment sizes, their taxonomic assignation and the coverage using the Illumina reads.

As a proxy of the completeness, the BUSCO values for the six newly assembled genomes plus the *AcolN1* genome were obtained using BUSCO version 3.0.2 (50) with the *diptera_odb9* set as reference. Finally, the contigs for all seven assemblies were ordered and merged with *RaGOO* v1.1 (136) using the chromosome level *An. gambiae* AgamP4 assembly.

### Gene annotation transfer

The *gff* for the genome annotation for AgamP4 was transferred into the newly assembled genomes using *Liftoff* (137) with default parameters. The annotation was manually inspected using *UGENE* version 35 (138) and whenever needed the annotation was accordingly corrected. 96% of the AgamP4 genes were correctly transferred.

### Construction of the curated TE library and *de novo* TE annotation

We ran the *TEdenovo* pipeline (51) independently on each of the seven genomes with default parameters. The obtained consensus in each genome were further filtered by discarding those generated with (i) only one sequence; (ii) with less than one full-length fragment mapping to the genome; (iii) with less than three full-length copies; and (iv) shorter than 100bp (Additional file 1: Table S2). The remaining consensuses were manually curated to remove redundant sequences and artifacts by manual inspection of coverage plots generated using the *plotCoverage* tool from REPET and visualization of the structural features on the genome browser IGV version 2.4.19 (139).

To ensure that we identified as much of the TE diversity as possible, the *TEfam* (tefam.biochem.vt.edu) database, which contains the TE libraries for several species of mosquitoes, was used to annotate the seven genomes using *RepeatMasker* version open-4.0.9 (Smit et al. 2015). Families with more than three matches longer than 90% in any genome were selected and their hit with the highest identity from each genome was extracted. These sequences were added to the REPET library and all the consensuses were clustered using CD-HIT version 4.8.1 (140) with the *-c* and *-s* parameters set to 0.8. These filters ensured that all TEs with an identity greater than 80% throughout more than 80% of their sequence were grouped in the same family. 85 clusters contained sequences only identified by *TEfam*. The sequences belonging to the same cluster were used to perform a multiple sequence alignment and the consensuses were obtained.

The consensuses were classified using PASTEC (141) with default parameters. Next their bidirectional best-hits were calculated using BLAST (142) against the *TEfam* (tefam.biochem.vt.edu), AnoTExcel (143) and Repbase (144) databases. When more than 80% of a consensus matched to a feature from the databases with an identity higher than 80%, the classification was transferred to the consensus. While not an order *per se*, MITEs were grouped together for subsequent analysis. Additionally, we classified the families based on the conservation of features characteristic of their orders into putative autonomous, putative autonomous lacking terminal inverted repeats (TIRs) or long terminal repeats (LTRs), putative non-autonomous, such as MITEs and TRIMs, and degenerated (Additional file 1: Table S4) (Fonseca et al 2019). These classified consensuses were used to re-annotate the assembled genomes with the *TEannot* pipeline using default parameters and we discarded copies whose length overlapped >80% with satellite annotations (52).

### Transfer of TE annotations to the *AcolN1* reference genome

We transferred the euchromatic TE annotations from the six genomes we sequenced to the *AcolN1* genome. First, we built a gff file composed by the coordinates of two 500 bp long “anchors” adjacent to each TE. We transferred these features considering each pair of anchors as exons from a single gene using the *Liftoff* tool with the -exclude_partial -overlap 1 -s 0.8 parameters (137). We conserved only transfers where both anchors were transferred to the AcolN1 genome. When both anchors were separated by less than 10 bp we considered the TE to be absent. This allowed us to identify most TE insertions, both present and absent in the AcolN1 genome. Next, following the same strategy we transferred these regions from the AcolN1 genome to the other six genomes. To summarize this information, we built a matrix containing the status for all of the TE insertions transferred to the AcolN1 genome in every genome. When the anchors were found more than 10 bp away the TE was considered to be present and this was represented with a 1, when the distance was less than 10 bp the TE was considered to be absent and it was represented with a -1. When any of the anchors was not transferred the TE was considered to be not transferred and represented with a 0.

Overall, we were able to transfer to the AcolN1 genome 71.93% to 75.29% of the TEs present in each of the six genomes leading to a total of 67,548 TEs transferred. We then attempt to transferred back these 67,548 from the AcolN1 reference to the remaining six genomes and we were able to confidently transfer 53,893 (79.78%) of these regions to at least three of the genomes with 32,185 (47.65%) transferred to all six genomes. We checked whether the TEs that failed to be transferred were enriched for nested TEs and we found that this was the case: while 68.58% and 51.18% of the TEs located more and less than 500 bp away from other TEs, respectively, were transferred, only 38.26% of the TEs overlapping other TEs were transferred to the six genomes.

### Identification of newly described families in other species

We analyzed all 10 available fully sequenced species from the Pyretophorus series, which belongs to the Cellia subgenus. We also included an additional five *Anopheles* species, three from each of the other series from the Cellia subgenus and two from the other subgenera with available fully sequenced species. As outgroups we included the genomes of *Cx. quinquefasciatus, Ae. aegypti* and *D. melanogaster. RepeatMasker* version open-4.0.9 (Smit et al. 2015) was run with default parameters using the 64 newly described families as the library on the following genomes: *An. albimanus* (AalbS2), *An. atroparvus* (AatrE3), *An. farauti* (AfarF2), *An. funestus* (AfunF3), *An. stephensi* (AsteS1), *An. epiroticus* (AepiE1), *An. christyi* (AchrA1), *An. merus* (AmerM2), *An. gambiae* (AgamP4), *An. coluzzii* (AcolN1), *An. melas* (AmelC2), *An. arabiensis* (AaraD1), *An. quadriannulatus* (AquaS1), *An. bwambae* (Abwa2) and *An. fontenillei* (ASM881789v1), *Cx. quinquefasciatus* (CulPip1.0), *Ae. aegypti* (AaegL5.0) and *D. melanogaster* (ISO1 release 6).

### Identification of heterochromatin

The coordinates for the pericentric heterochromatin, compact intercalary heterochromatin, and diffuse intercalary heterochromatin in *An. gambiae* AgamP3 were obtained from a previous work (62). The *An. gambiae* AgamP3 genome assembly was mapped against the seven *An. coluzzii* genome assemblies using *progressiveMauve* (145) and the corresponding coordinates on each of the assemblies were retrieved (Additional file 1: Table S1C). To identify families enriched in either euchromatin or heterochromatin a χ^2^ test of independence was performed.

### Transfer of known inversion breakpoints

The coordinates for inversions 2La, 2Rb, 2Rc and 2Rd were obtained from Corbett-Detig et al. (2019) (69) and coordinates for 2Ru were obtained from White et al (2009) (70). 50 kb regions flanking each side of the insertion were obtained and mapped using *minimap2* (146) against the scaffolded genome assemblies to transfer the breakpoints. To validate the breakpoints, we determined if long reads spanned the breakpoints using the genome browser IGV version 2.4.19 (139).

### Detection of putatively active TE families

To identify potentially active TE families, we identified families with more than two identical full-length fragment copies in at least six of the seven annotated genomes. We determined the fraction of identical copies of these families by identifying all their insertions in the genome and calculating the sequence identity of all their bases against the consensus by performing a nucleotide BLAST. Given that long-read have a higher rate of sequencing errors that could affect the age estimation (although we used Illumina reads for polishing), we also used dnaPipeTE (147) to estimate the relative age of the TE families using the raw Illumina reads for the six genomes that we sequenced. We compared the TE landscape obtained using dnaPipeTE with that obtained using the BLAST procedure, using a Kolmogorov-Smirnov test corrected for multiple testing using the Benjamini–Hochberg procedure (Additional file 1: Table S16). Given that we observed few significant differences, we continued using the landscape data obtained using the BLAST procedure. We identified the families where the majority of the bases of their insertions were on the peak of identical sequences in the TE landscape (>50% of the bases with >99% base identity) in more than five of the seven genomes we analyzed. Finally, we assessed the ability to actively transpose of strong candidates by identifying their intact ORFs, LTRs (in the case of LTR retrotransposons) and target site duplication (TSD), and estimated the % identity between the two LTR of each TE copy.

### Classification of TEs by their genomic location

To determine the location of TEs we used the *findOverlaps* function from the *GenomicAlignments* R package (148) using default parameters. Both the TE and the gene annotation were converted to *GenomicRanges* objects ignoring strand information in the case of TEs.

### Insecticide resistance genes

A list with a total of 43 relevant insecticide resistance genes was generated taking several works into consideration (3, 100-102) (Additional file 1: Table S14). To determine the position of the L to M nonsynonymous substitution that we observed in AGAP004707 (*para*) we used the position from the CAM12801.1 reference sequence.

### Immune-related genes

The full list of 414 immune-related genes from *An. gambiae* was downloaded from ImmunoDB (104). We conserved the 281 most reliable genes filtering by the STATUS field and conserving only those with A or B scores (A refers to genes confirmed with high confidence and expert-refined cDNA supplied, and B refers to genes confirmed with high confidence, no refinement required).

### TFBS and promoter identification

The matrices for *dl* (MA0022.1), *cnc::maf-S* (MA0530.1) and *Stat92E* (MA0532.1) were downloaded from JASPAR (http://jaspar.genereg.net/) (149). The sequences for the TEs of interest were obtained using *getSeq* from the *Biostrings* R package. The TFBS in the sequences were identified using the web version of FIMO (150) from the MEME SUITE (151) with default parameters. The ElemeNT online tool was used to identify promoter motifs (152).

## Results

### Six new whole-genome assemblies of *An. coluzzii* from two major cities in Central Africa

To explore the TE diversity in *An. coluzzii*, we used long-read sequencing and performed whole genome assemblies and scaffolding of larvae collected from six natural urban breeding sites: three from Douala, Cameroon, and three from Libreville, Gabon, Central Africa (Figure 1A, Additional File 1: Table S1A; see Material and Methods). Additionally, we performed a reference-guided scaffolding of the available *An. coluzzii* reference genome *AcolN1* using the chromosome level assembly *AgamP4* of *An. gambiae* (49). Although the genomes analyzed varied in sequencing coverage and read length, these differences only had an effect on contig N50 but did not affect other assembly and scaffolding metrics (Additional File 1: Table S1B). Overall the number of scaffolds varied from 5 to 107, with the median being 20, however, the scaffolds’ N50 was similar across the seven genomes (Table 1).

**Figure 1.**
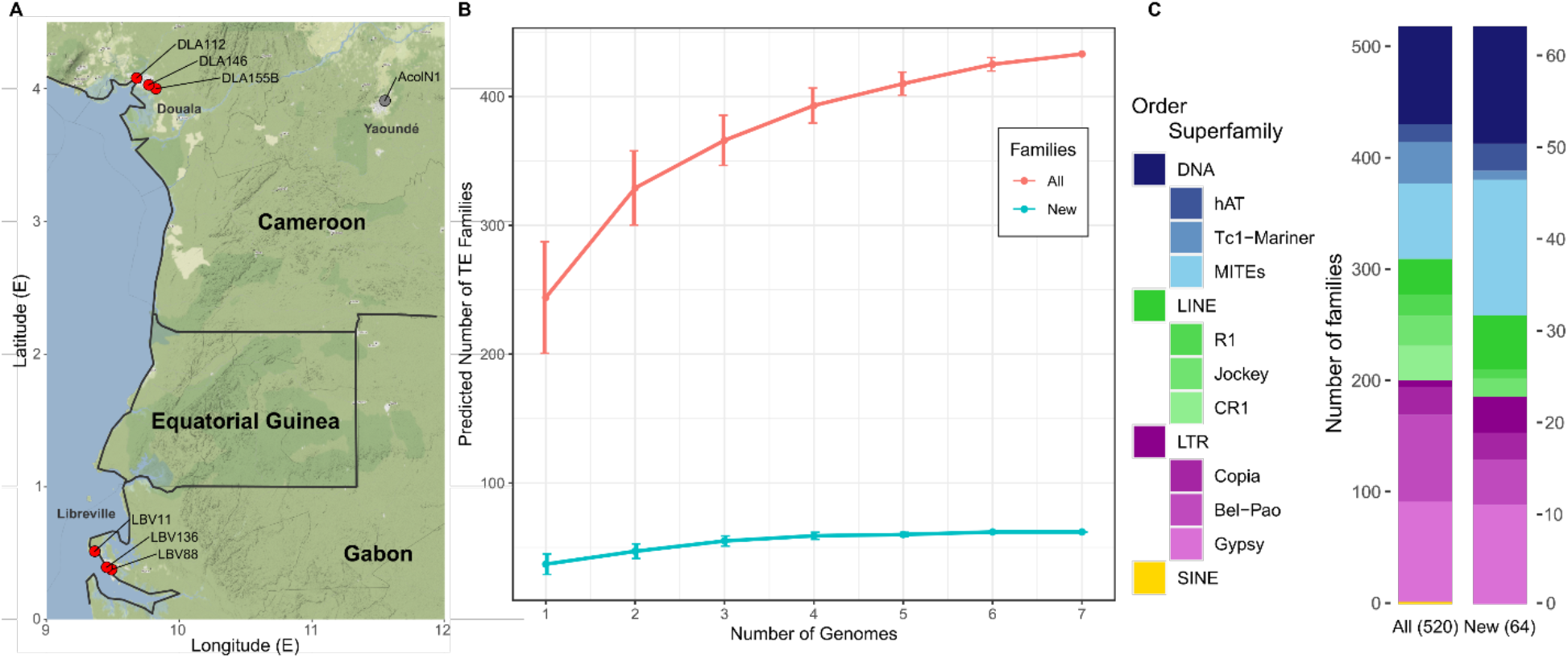
Transposable elements in *An. coluzzii*. A) Geographic location of the six breeding sites analyzed (in red) and of the place of origin of the Ngousso colony (in grey) which was used to generate the *AcolN1* genome. B) Number of TE families identified when using a single genome or when using all possible combinations of more than one genome. The red line shows the total number of TE families and the blue line shows the number of newly described families. Note that 76% of all the TE copies where already identified when analyzing a single genome (Additional File 2). C) Classification of all TE families and newly described families in *An. coluzzii*. The three most abundant superfamilies from each order are shown.

**Table 1.** Genome assemblies and scaffolds’ statistics for the *An. coluzzii* genomes analyzed in this work. Three genomes were collected in Douala (DLA) and three in Libreville (LBV). ^*a*^ *LBV11* was sequenced using PacBio technologies, while the other five genomes were sequenced using Oxford Nanopore Technologies. ^b^ Genome statistics for *AcolN1*, the high quality *de novo* genome assembly reported by Kingan *et al*., (49) are also included.

| Genome | Long reads coverage | Illumina coverage | Assembly size (Mb) | Number of contigs | Number of scaffolds | Scaffolds N50 (kb) | Complete BUSCO genes | Genes transferred | TE families identified |
| --- | --- | --- | --- | --- | --- | --- | --- | --- | --- |
| *DLA112* | 55X | 59X | 252 | 3,917 | 107 | 54,591 | 96.6% | 13,469 | 244 |
| *DLA155B* | 28X | 19X | 236 | 2,081 | 24 | 52,031 | 89.5% | 13,303 | 243 |
| *DLA146* | 28X | 42X | 247 | 2,036 | 14 | 54,960 | 95.1% | 13,314 | 193 |
| *LBV88* | 31X | 41X | 245 | 2,576 | 19 | 54,450 | 94.5% | 13,328 | 280 |
| *LBV136* | 34X | 130X | 236 | 2,911 | 28 | 52,053 | 95.2% | 13,307 | 172 |
| *LBV11<sup>a</sup>* | 89X | 61X | 246 | 2,608 | 20 | 53,712 | 94.2% | 13,393 | 294 |
| *AcoIN1<sup>b</sup>* | ~270X | - | 251 | 205 | 5 | 53,057 | 98.9% | 13,487 | 283 |

We assessed the genomes completeness using BUSCO with the dipteran set of genes (50). We obtained percentages of complete genes ranging from 94.2% to 96.6% except for the *DLA155B* sample which had a lower completeness value (89.5%; Table 1). These completeness values for most (5/6) of the samples were similar to those from the *AcolN1* genome assembly which contained 98.9% complete genes (Table 1; Additional file 1: Table S1).

Overall the analyzed genomes are comparable in terms of scaffold contiguity and completeness (Table 1 and Additional File 1: Table S1).

### 64 new anopheline TE families discovered in *An. coluzzii*

To identify the TE families present in each of the genomes, we used the *TEdenovo* pipeline from the REPET package (51). After several rounds of manual curation, we identified between 172 and 294 TE families for each genome (Table 1; Additional file 1: Table S2). Remarkably, while using a single reference would have only allowed the identification of a median of 244 TE families, clustering the TE libraries from an increasing number of genomes allowed the identification of a total of 435 well supported TE families (Figure 1B; see Material and Methods). Interestingly, 64 of these families (32 DNA, 9 LINEs and 23 LTRs) are described here for the first time. The majority of the new families (43/64) had partial matches to other known TEs, thus allowing us to classify them at the superfamily level (Additional file 1: Table S3). The use of multiple references was especially relevant for identifying these previously undescribed families given that using a single genome would have only allowed to identify a median of 37 (25-48) novel TE families (Figure 1B).

To further characterize these novel families, we estimated the average number of insertions in the seven *An. coluzzii* genomes, and their distribution and abundance in other species from the *Anopheles* genus (Figure 2; Additional file 3: Figure S2; Additional file 1: Table S3). To do this, we first annotated individual TE insertions in the seven *An. coluzzii* genomes using the *TEannot* pipeline from the REPET package (52). To ensure that our annotation was as complete as possible, in addition to the 435 families identified using REPET *de novo* approach, we also added to our library 85 TE families from the *TEfam* database that we found to also be present in the *An. coluzzii* genomes reported in this work (see material and methods). Most probably, these 85 families were either not identified by REPET or were discarded during manual curation (Additional file 1: Table S4; see Material and Methods) (53). The final total of 520 families were classified into 23 superfamilies and then further grouped into four orders (DNA, LINE, LTR and SINE; Figure 1C).

**Figure 2.**
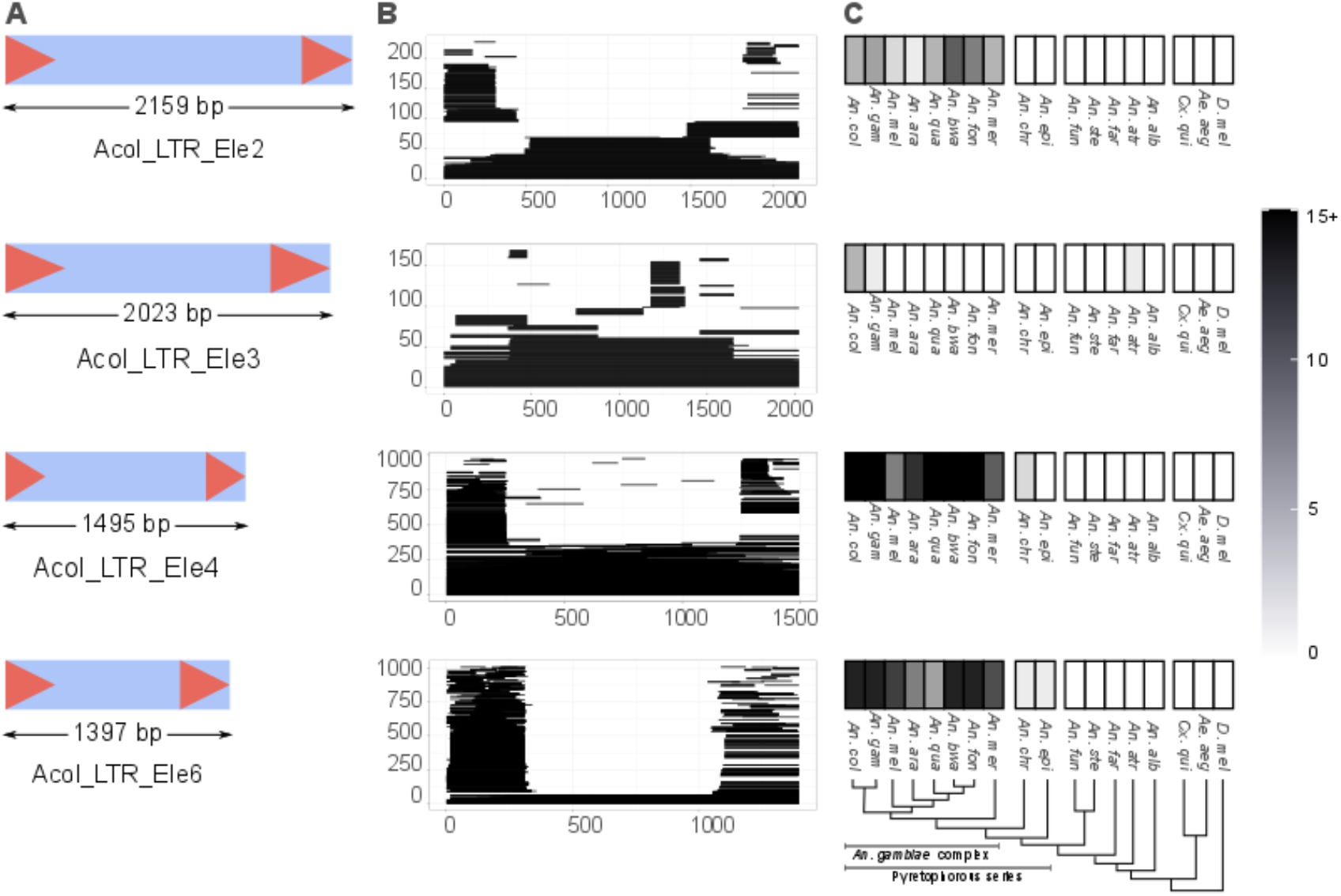
Structure, abundance and phylogenetic distribution of novel TE families. The four newly identified TRIMs families are shown, for the remaining 60 novel families see Additional file 3: Figure S2. A) The structure of each new family is displayed: the light blue box represents the full extension of the TE and the red arrows represent LTRs. B) All insertions for each TE family were identified and are shown as a coverage plot in which each lines represents a copy in the genome. Note that the large number of stacked horizontal lines in the extremes of the plot represent and abundance of solo LTRs. C) Phylogenetic distribution of the TE family insertions in 15 members of the *Anopheles* genus, *Culex quinquefasciatus, Ae. Aegypti* and *D. melanogaster*. The number of insertions with more than 80% identity and spanning at least 80% of the consensus, in each species is shown using a black and white gradient. Species with no insertions are shown in white while species with 15 or more insertions are shown in black.

Copies from all 64 new families were found in all seven *An. coluzzii* genomes, further suggesting that these are *bona fide* families. Although the majority of families contain full-length copies in at least one of the seven genomes analyzed, truncated copies were the most abundant (Figure 2B; Additional file 1: Table S3). We identified a median of 72 insertions (ranging from 16 to 1445) per family and genome (Figure 2B; Additional file 3: Figure S2B). Two out of the four TRIM elements identified (*Acol_LTR_Ele 4* and *Acol_LTR_Ele 6*) are among the most abundant new families, with more than 150 insertions (Figure 2B). TRIM elements are non-autonomous retrotransposons flanked by LTRs and lacking coding capacity (Figure 2A). These elements have not been previously described in anopheline genomes and are still underexplored in insect genomes in general (54-56). However, they might be important players in insect genome evolution: in plants there are some examples of TRIM elements showing the capacity to restructure genomes by acting as target sites for retrotransposon insertions, alter host gene structure, and transduce host genes (57, 58). While we found TRIM elements in *An. coluzzii* genomes to be underrepresented in gene bodies (χ^2^ test *p-value* > 0.05), they were overrepresented in nested insertions (χ^2^ test *p-value* < 0.05).

We also assessed the phylogenetic distribution of the 64 new TE families in 15 species of the *Anopheles* genus, including the eight members of the *An. gambiae* complex, two more distantly related mosquitoes species (*Culex quinquefasciatus, Aedes aegypti*) and *Drosophila melanogaster* (Additional file 1: Table S3) (36, 59, 60). We found that the new families were unevenly distributed among the members of the *Anopheles* genus (Figure 2C and Additional file 3: Figure S2C). Ten families were exclusively found in members of the Pyretophorus series, suggesting that these elements emerged after the split of this series from the Cellia subgenus. Moreover, 13 families were also found in at least one of the other three non-anopheline species (Additional file 3: Figure S2C). The distribution of these 13 families was patchy, with some of them present only in distantly related species while others were present in members of the *Anopheles* genus or in members of the Pyretophorus series. These suggests that some of these families might have been acquired through horizontal transfer events (Additional file 1: Table S3) (40).

### TE superfamily abundance varies across the seven genomes analyzed

The percentage of the genome represented by TEs across the seven genomes varied between 16.94% and 20.21% (Table 2). Note that differences across genomes in assembly and scaffolding statistics did not explained the differences in TE content or superfamily abundance (Additional File 1: Table S1B).

**Table 2.** TE content in the seven genomes analyzed. TE copy number, TE content in megabases and percentage of the genome represented by TEs. Values are given for the whole genome and for the euchromatin and heterochromatin compartments separately.

| Genome | Whole genome |  |  | Euchromatin |  |  | Heterochromatin |  |  |
| --- | --- | --- | --- | --- | --- | --- | --- | --- | --- |
|  | TE copy number | Mb | Genome % | Copy number | Mb | Region % | Copy number | Mb | Region % |
| *DLA112* | 72901 | 48.00 | 19.02 | 49853 | 28.18 | 12.67 | 22930 | 19.74 | 70.34 |
| *DLA155B* | 62999 | 40.08 | 16.94 | 45592 | 25.39 | 11.86 | 17371 | 14.66 | 65.76 |
| *DLA146* | 68658 | 45.42 | 18.40 | 47874 | 27.22 | 12.36 | 20682 | 18.15 | 68.35 |
| *LBV88* | 68593 | 45.81 | 18.70 | 48922 | 28.06 | 12.81 | 19582 | 17.68 | 68.74 |
| *LBV136* | 64343 | 40.79 | 17.26 | 45792 | 24.97 | 11.73 | 18406 | 15.73 | 67.59 |
| *LBV11* | 71803 | 47.59 | 19.58 | 50187 | 28.95 | 13.40 | 21564 | 18.60 | 70.22 |
| *AcoIN1* | 75745 | 50.81 | 20.21 | 48537 | 26.10 | 11.95 | 27205 | 24.70 | 74.77 |

We found a positive correlation between TE content and genome size as has been previously described in *Anopheles* and other species (Pearson’s r = 0.90, significance = 0.007; Additional file 4: Figure S3) (40, 61). As expected due to heterochromatin being a TE rich region and thus challenging to assemble (62), most of the differences in TE content across genomes were found in the heterochromatin compartment (Table 2; χ^2^ test for variance, *p-value* = 3.57e-3).

To assess whether differences in TE content at the family and superfamily level existed among the seven genomes, we focused on the TE copy number in euchromatic regions as we are mostly interested in the potential functional impact of TE insertions. We found significant differences at the order and superfamily levels (χ^2^ test *p-value* = 1.07e-21 and *p-value* = 1.69e-14, respectively). The largest differences were found in the LTR order: LTRs were more abundant in the *DLA112* and *LBV88* genomes and less abundant in *AcolN1* (Figure 3A). At the superfamily level, we found that the largest differences were in the *Gypsy* superfamily, which belongs to the LTR order. We also observed an enrichment of the *RTE* superfamily in *LBV11*, of the *CR1* and *Bel-Pao* superfamilies in *DLA112*, and a depletion of the *CR1* superfamily in *AcolN1* (Figure 3). Therefore, most of the differences in TE content between the evaluated genomes appear to be in retrotransposon families.

**Figure 3.**
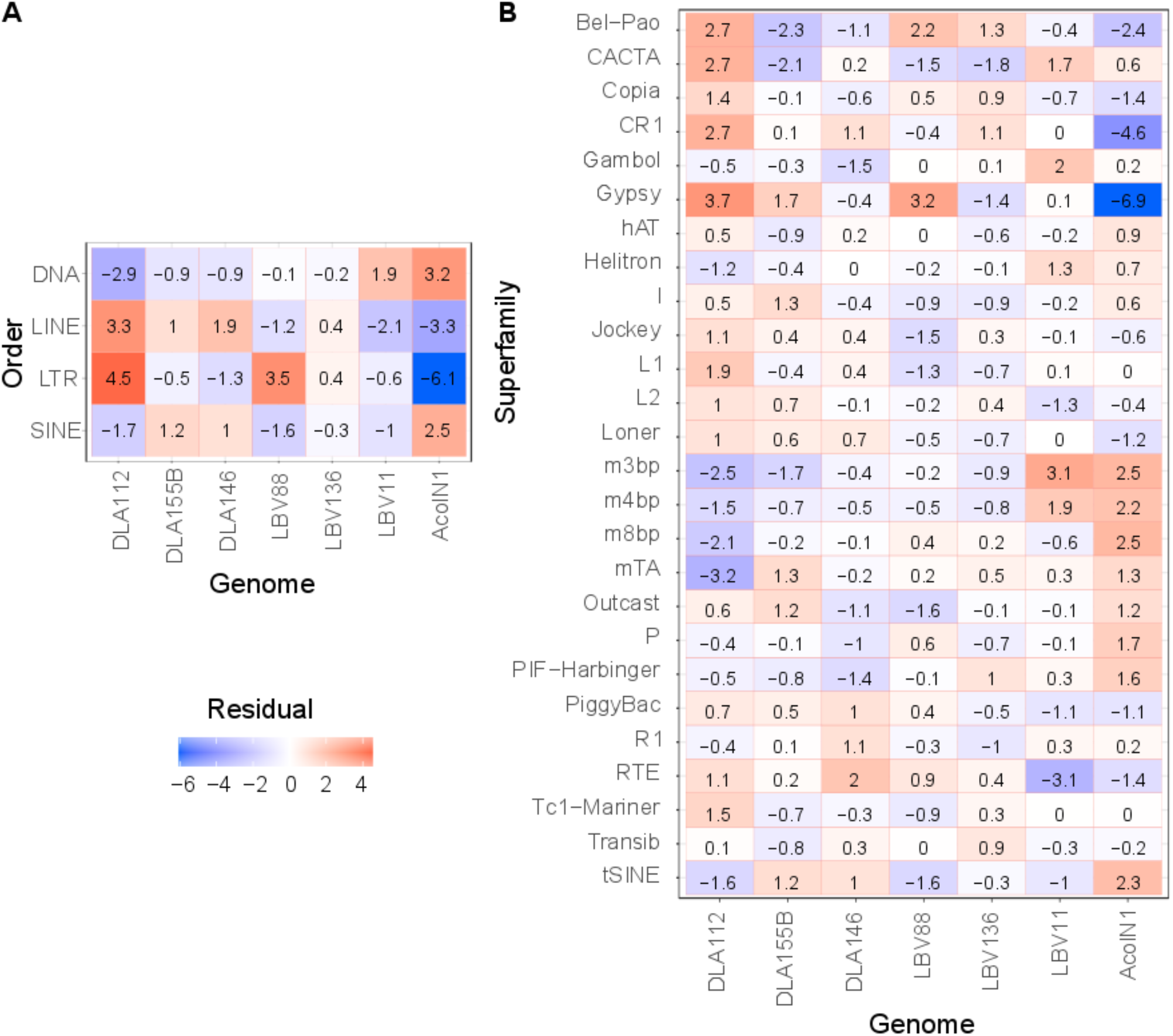
Differences in TE content between the seven *An. coluzzii* genomes. Differences are shown at the (A) order and (B) superfamily levels. χ^2^ tests were performed for the number of insertions and the Person’s residuals are shown. Note that MITEs are divided into the m3bp, m4bp, m8bp and mTA superfamilies.

### TEs are nonrandomly distributed throughout the genome

As expected, we found that the percentage of TEs in euchromatin, 11.73%-13.40%, is much lower than the percentage of TEs in heterochromatin, 65.76%-74.77% (χ^2^ test *p-value* < 0.05; Table 2 and Figure 4A). None of the TE families identified were exclusive to either the euchromatin or heterochromatin. However, 45 families were enriched in the euchromatin (χ^2^ test, *p-value* < 0.01) including 12 out of the 32 *mTA* MITE families (Additional file 1: Table S5). This is in line with what has been previously reported in *Ae. aegypti* (63). We also observed that the TE distribution was uneven between the chromosomes, and as expected, the X chromosome had a larger fraction of its euchromatin spanned by TEs (Figure 4B) (64).

**Figure 4.**
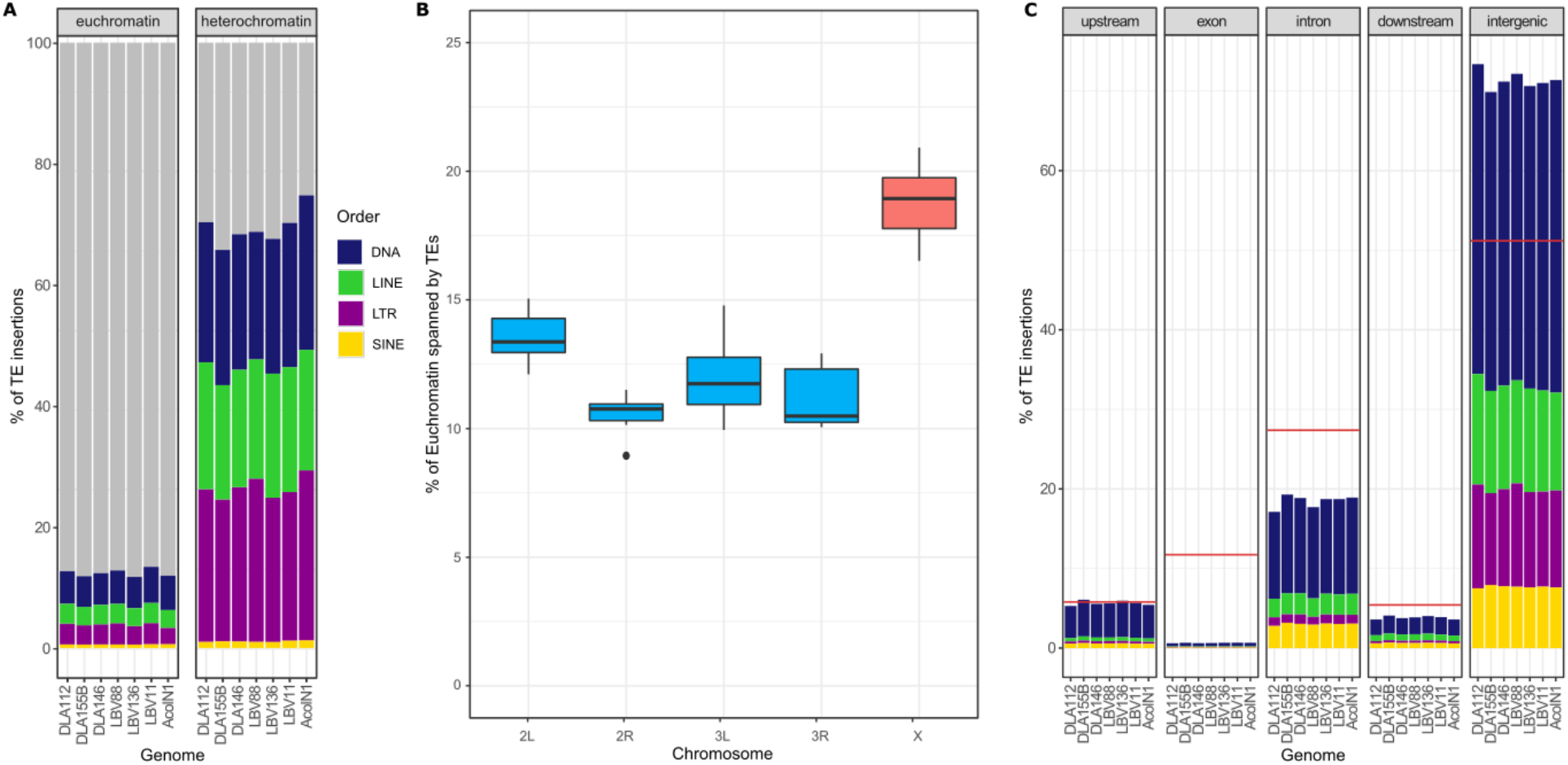
TE insertions distribution throughout the genomes. A) Percentage of euchromatin and heterochromatin occupied by TEs in each of the seven analyzed genomes. Each order is shown in a different color. B) Boxplots of the percentage of the euchromatin of each chromosome covered by TEs. Autosomes are shown in blue and the X chromosome in red. C) Percentage of TE insertions in each genome that fall in a specific genomic region. A red line is used to display the expected percentage that should be covered by TEs taking in consideration the size of the genomic region. Each order is shown in a different color as in A).

Finally, we also determine the distribution of TE insertions regarding genes. We divided the genome in five regions: 1 kb upstream, exon, intron, 1 kb downstream and intergenic (65). More than half of the genes (7239) in *An. coluzzii* had TEs either in their body or 1 kb upstream or downstream. Many of these genes (3888/7239) had insertions in all seven genomes, while 1065 genes have an insertion only in one genome. We found that the number of insertions in intergenic regions was higher than expected by chance while the number of insertions in exons was lower (χ^2^ test *p-value* < 0.001; Figure 4C; Additional file 1: Table S6). The upstream and downstream regions behaved differently: the downstream region had a smaller amount of TEs than expected by chance and the upstream region was neither enriched nor depleted for TE insertions (χ^2^ test *p-value* = 0; Additional file 1: Table S6). This is possibly linked with the chromatin state of these regions given that downstream regions are more commonly in a closed chromatin state (65).

Focusing on the TE orders, we observed that LTR elements were more abundant on intergenic regions while SINEs were more abundant on introns, and DNA elements were more abundant in introns and in the upstream region (χ^2^ test *p-value* < 2.03e-3; Figure 4C; Additional file 1: Table S7A). MITEs, which are non-autonomous DNA elements have been reported to be more abundant in the introns and flanking regions of genes (66). We observed the same behavior for *mTA* and *m3bp* MITEs, which are more abundant in upstream regions and introns, and *m8bp* MITEs which are more abundant in introns (Additional file 1: Table S7B).

Overall, TEs are not randomly distributed in the genome, as they are more abundant in heterochromatic than in euchromatic regions, more abundant in the X chromosome than in autosomes, and more abundant in intergenic regions than in gene bodies or gene flanking regions.

### MITE insertions are present in several inversion breakpoints

TEs have been suggested to be involved in chromosome rearrangements within the *An. gambiae* complex. Indeed, TEs have been found in close proximity to the breakpoints of the 2La in *An. gambiae* and *An. melas*, and to the breakpoints of the 2Rb inversions in *An. gambiae* and *An. coluzzii* (67, 68). We thus explored the TE content in the breakpoints of the 2La and 2Rb, and three other common polymorphic inversions in *An. coluzzii*: 2Rc, 2Rd, and 2Ru (69). We were able to identify the two breakpoints regions in at least one genome for all the insertions except for the inversion *2Ru* for which we could only identify the distal one (Figure 5). The analysis of these breakpoint regions suggested that analyzed genomes have the standard conformation for all five inversions (see Material and Methods; Additional file 1: Table S8). We identified several TEs nearby the proximal and the distal breakpoints of 2La and 2Rb, in agreement with previous studies (Figure 5) (26, 67, 68). For the standard 2La proximal breakpoint, Sharakhov et al. (67) identified several DNA transposons and a SINE insertion. We also identified a cluster of MITE insertions, which are DNA transposons; however, we additionally identified an *Outcast* (LINE) element (Figure 5). Regarding the standard 2La distal breakpoint, we observed two MITEs similar to one of the insertions in the proximal breakpoint, which was in agreement with the findings by Sharakhov et al. (67) (Figure 5). We also observed similar behavior in the 2Rb breakpoints, such as the one described by Lobo et al., (68): tandem repeats flanking the inversion in the standard and inverted forms, and TEs in the internal sequences of both breakpoints (Figure 5). For the 2Rd inversion, we identified MITEs near both breakpoints. Finally, we have also described here for the first time, TE insertions that are present in the distal breakpoint of inversion 2Ru but not near the estimated proximal breakpoint; although in the latter case we were able to identify reads spanning the breakpoints in the seven genomes (70).

**Figure 5.**
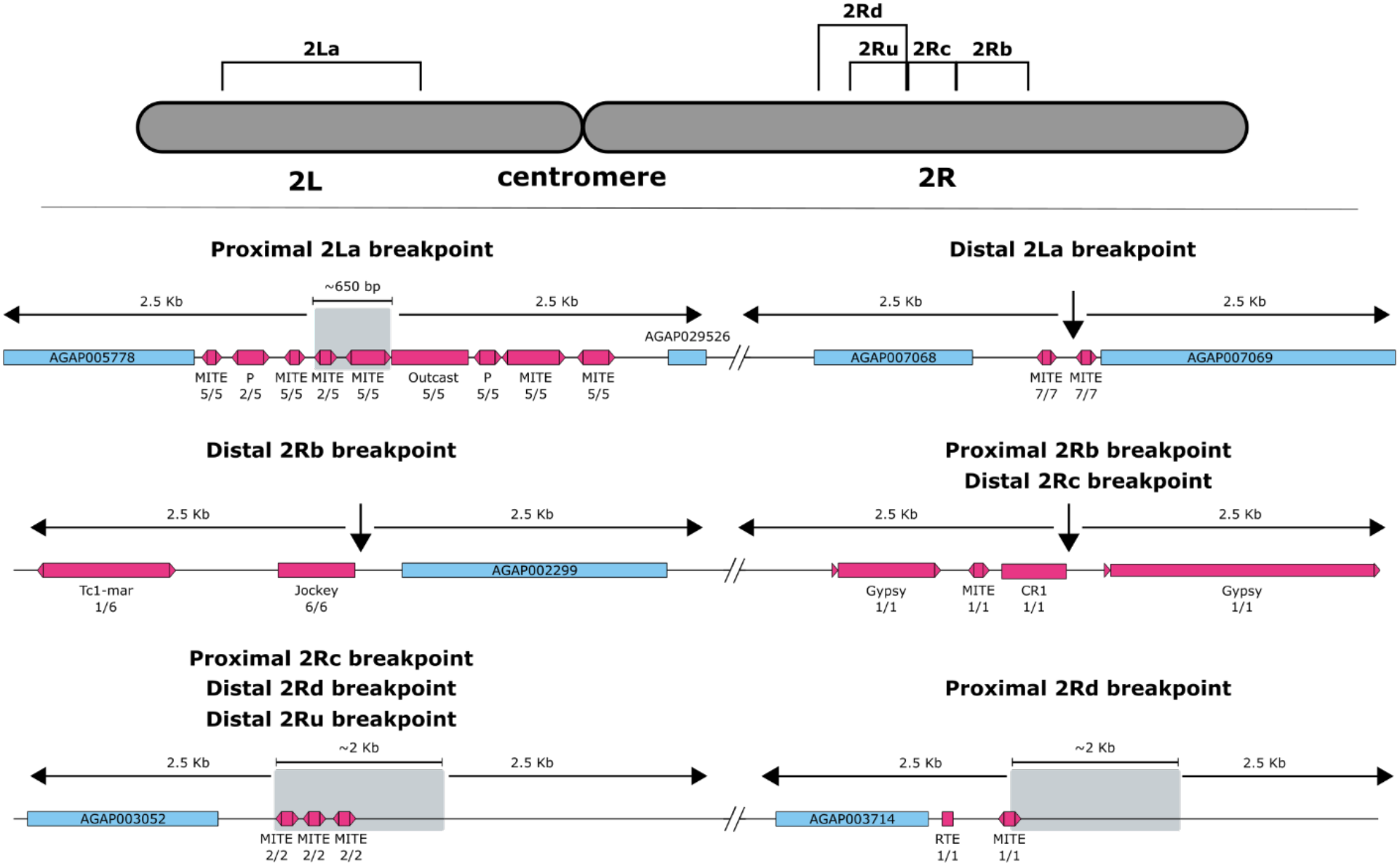
TE insertions near known inversion breakpoints. Diagram of chromosome 2 with the analyzed inversions. For each inversion both breakpoints, proximal (closer to the centromere) and distal (farther from the centromere), plus 2.5 kb to each side are shown. When the position of a breakpoint was not identified at the single base pair level, the interval where the breakpoint is predicted to be is shown in a grey box. Genes are shown as blue boxes while TEs are shown as pink boxes. Below each TE, the family of the TE is shown and below the family name the number of genomes where the insertion was found and the number of genomes where the breakpoint region was identified. Note that breakpoints are shared among some of the inversions.

### Six of the seven potential active families are LTR insertions

To identify potentially active TE families, we first estimated their relative age by analyzing the TE landscapes (71, 72). We observed an “L” shape landscape in all genomes which is indicative of a recent TE burst (Additional file 5: Figure S4) (73). This “L” shape landscape, dominated by retrotransposons, had previously been described for the sister species *An. gambiae* (72, 74), where numerous Gypsy LTR Retrotransposons (up to 75%) might currently be active (75, 76). We further investigated the families in the peak of the landscape and we identified eight families with more than two identical full-length fragments and with more than half of their copies identical to the consensus (Additional file 1: Table S9A). Additionally, we assessed the potential ability of our candidates to actively transpose by identifying their intact open read frames (ORFs), LTRs (in the case of LTR retrotransposons), and target site duplications (TSDs), and determined that seven of these families are potentially fully capable of transposing. For the LTR families, we further evaluated the identity between LTRs of the same copy. Mean identities ranged from 97.38% to 99.44% across the six LTR families, with 17.64% to 40,35% of the copies having identical LTRs (Additional File 1: Table S9B). These results further suggest that these families might be responsible for the recent retrotransposon burst in *An. coluzzii* (Additional file 1: Table S9A). Note that LTR elements accounted for most of the differences in TE content across genomes (Figure 3).

As a first step towards assessing the potential functional consequences of the TE insertions from these seven putatively active families, we focused on insertions that occurred in introns, exons, and 1 kb upstream or downstream of a gene. We identified 66 genes with insertions from these families, with four genes containing up to two insertions in the same gene region (Additional File 6: Figure S5; Additional File 1: Table S10 and Table S11). Six of the genes have functions related to vectorial capacity: insecticide resistance, immunity, and biting ability (Table 3). We checked whether the TE insertions nearby these genes contained binding sites for transcription factors or promoter motifs (Additional file 1: Table S12; Additional file 1: Table S13). We focused on identifying binding sites for three transcription factors that are known to be involved in response to xenobiotics (cap’n’collar: *cnc*) and in immune response and development (dorsal: *dl* and signal transducer and activator of transcription: *STAT*) given the availability of matrix profiles from *D. melanogaster* (77, 78). We identified binding sites for either *dl, STAT* or both in three insertions; interestingly the *Acol_gypsy_Ele18* and the *Acol_copia_Ele8* insertions have more than three binding sites for the same transcription factors, suggesting that they might be functional sites (Table 3) (79). Additionally, the genes that contained these TE insertions also contained binding sites for the same transcription factors, which suggests that these transcription factors already played a prior role in their regulation. We also identified a putative promoter sequence in the *Acol_copia_Ele24* insertion found upstream of the CLIPA1 protease encoded by AGAP011794 which could also potentially lead to changes in the regulation of this gene (Additional file 1: Table S14).

**Table 3.** Seven TE insertions from putatively active families are located nearby genes related with vectorial capacity. TE Freq. specifies the number of genomes where the TE insertion was found and the number of genomes where the gene structure was correctly transferred (genes where some exons were missing were not taken into consideration in this analysis). References in the Phenotype column are as follows: 1 (80), 2 (81), 3 (82), 4 (83), 5 (84), 6 (85), 7 (86), 8 (87). The number in parenthesis in the transcription factor binding site (TFBS) column refers to the number (or range) of TFBS found in the TE. *The insertion size in parenthesis refers to an insertion found in one of the genomes corresponding to a solo-LTR insertion.

| TE family | Insert size (bp) | TE Freq. | Gene | Function | Possible phenotype [Reference] | TFBS |
| --- | --- | --- | --- | --- | --- | --- |
| *Acol_copia_EI e8* | 3230 (200)* | 2/4 | AGAP012466 | cuticular protein RR-2 family 146 | Development, insecticide resistance [2, 3] | *dl* (3) and *STAT* (5) |
| *Acol_copia_EI e24* | 167 | 4/4 |  |  |  | - |
| *Acol_gypsy_EI e65* | 185 | 3/3 | AGAP010620 | Peptidase S1, PA clan | Immunity, digestion [4, 5] | - |
| *Acol_gypsy_EI e18* | 4858 | 1/6 | AGAP029191 | Defective proboscis extension response | "Bendy" proboscis [6] | *dl* (3-7) and *STAT* (1-6) |
| *Acol_copia_EI e24* | 168 | 1/6 | AGAP011794 | CLIPA1 protein | Digestion, immunity or development [7] | - |
| *Acol_gypsy_EI e18* | 235 | 1/7 | AGAP002633 | Gustatory receptor 53 | Vectorial capacity [8] | - |
| *Acol_gypsy_EI e65* | 141 | 1/3 | AGAP028069 | Peptidase S1, PA clan | Immunity, digestion [5, 6] | *dl* (1) |

### TE insertions could influence the regulation of genes involved in insecticide resistance

The usage of pyrethroids, carbamates, and DDT as vector control mechanisms has led to the rapid dispersion of insecticide resistance alleles in natural populations (88-92). Among the best characterized resistance point mutations are L1014F (*kdr-west*), L1014S (*kdr-east*), and N1575Y in the voltage gated sodium channel *para* (also known as *vgsc*), and G119S in the acetylcholinesterase *ace-1* gene (93-95). We first investigated whether the seven genomes analyzed in this work contained these resistance alleles. We found the *kdr-west* mutation in the six genomes from Douala and Libreville but not in *AcolN1* genome (49). None of the other mutations were identified, however a previously undescribed nonsynonymous substitution (L1688M) in the fourth domain of *para* was identified in the aforementioned six genomes. Whether this replacement also increases insecticide resistance is yet to be assessed.

TEs have been hypothesized to play a relevant role specifically in response to insecticides (96-98), and a few individual insertions affecting insecticide tolerance in anopheline mosquitoes have already been described (99). Thus, we searched for TE insertions in the neighborhood of insecticide-related genes that could potentially lead to differences in their regulation. We focused on eight well-known insecticide resistance genes: *Ace1, Cyp6p3, Gstd1-6, para, rdl, Cyp6m2, Cyp6z1* and *Gste2*. We also considered genes that have been shown to be differentially expressed in *An. gambiae* when exposed to insecticides (Additional file 1: Table S14; Additional file 7: Figure S6) (3, 100-102). We found that 21 out of the 43 genes analyzed contained at least one TE insertion. We also observed that *para* had the largest number of TE insertions (48 in average per genome, mainly in its introns) from this set of genes. This is an exception, given that the average number of insertions per gene is 2.95 for members of this set which falls within the expected number of insertions per gene in all the genome (t-test, *p-values* > 0.2).

Only one of the insertions, a solo LTR element of *Acol_Pao_Bel_Ele43* from the *Pao-Bel* superfamily and present in all the genomes analyzed, was located in the 3’ UTR of *GSTE2*. Interestingly, an upstream insertion possibly affecting the expression level of this gene has previously been identified in *An. funestus* (99). To determine if TEs could influence the regulation of insecticide-resistance genes, we focused on polymorphic (present in two or more genomes) and fixed (present in all seven genomes analyzed) insertions located in introns or 1 kb upstream or downstream of the gene. We searched for *cnc* binding sites, and for those insertions located in gene upstream regions we also looked for promoter motifs (Additional file 1: Table S12; Additional file 1: Table S13). We identified 15 insertions in 10 genes containing either *cnc* binding sites or promoter sequences. One insertion located in *CYP4C28* and two insertions in *para* contained binding sites for *cnc*, although the genes did not contain binding sites for this transcription factor. Additionally, we identified 12 insertions containing promoter motifs and located nearby nine genes (Figure 6). In some cases, such as the *Acol_m2bp_Ele10* MITE insertion in *ABCA4* or the *tSINE* insertion in *GSTMS2*, while the same TE insertion was found in six and seven genomes respectively, the promoter motifs were found only in four and one genome respectively (Figure 6; Additional file 1: Table S13). We analyzed the consensus sequence of these two families and we found that while the *Acol_m2bp_Ele10* had the promoter motif, the *tSINE* did not, suggesting that some of the *Acol_m2bp_Ele10* elements lost the promoter motifs while the *tSINE* copies acquired them.

**Figure 6.**
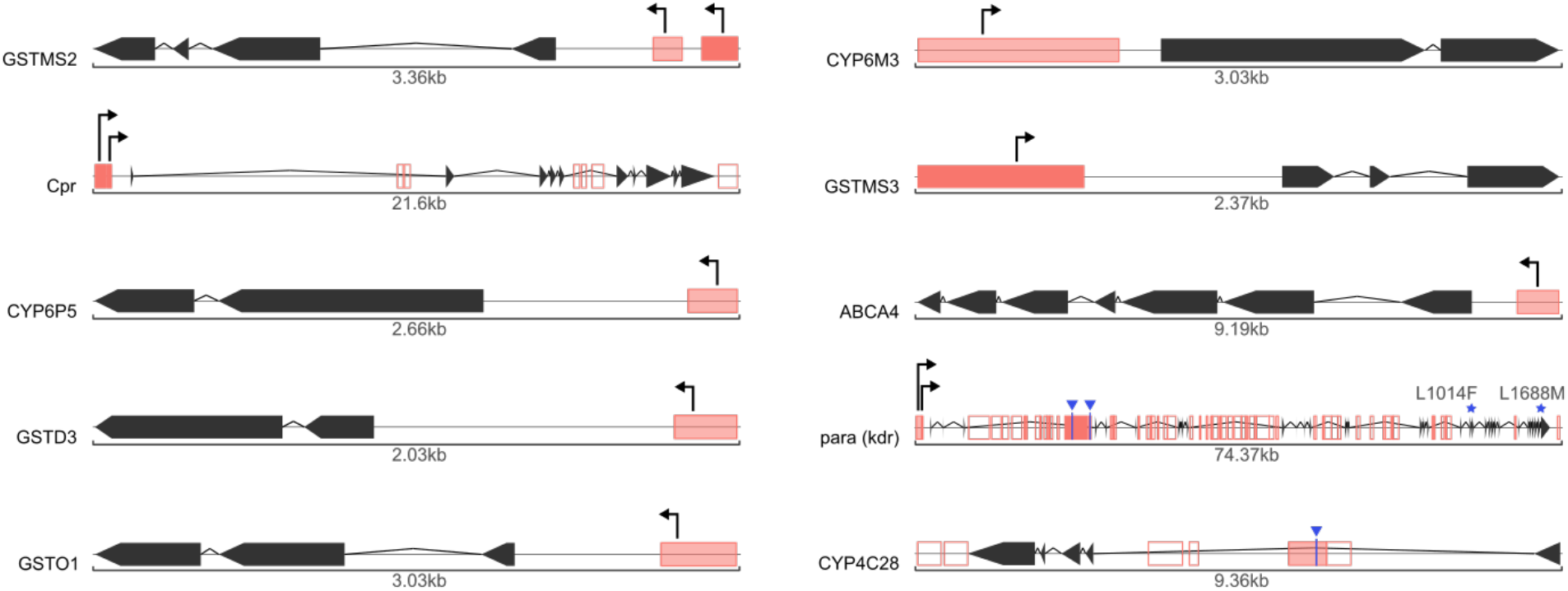
TE insertions in the neighborhood of genes involved in insecticide resistance. The gene structure is shown in black with arrows representing the exons. TE insertions are depicted as red boxes. When containing a TFBS for *cnc* or a promoter they are filled in red, otherwise they are empty. The red color is darker on fixed TEs and lighter on polymorphic TEs. Promoters are shown as arrows while *cnc* binding sites are shown in blue. Resistance alleles are shown for *para* (*kdr*).

### Immune response genes could also be potentially affected by TEs

Mosquitoes breeding in urban and polluted aquatic environments overexpress immune-related genes suggesting that immune response is relevant for urban adaptation (103). To assess the potential role of TEs in immune response, we searched for TE insertions in genes putatively involved in immunity according to ImmunoDB (104) (Additional file 1: Table S15). We identified 466 TE insertions in 148 out of the 281 genes analyzed. The number of insertions in each gene varied greatly going from 58 genes with a single insertion to AGAP000940, a gene coding for a C-type lectin and spanning 107.2 kb, with 48 insertions. The frequency of these insertions was also variable with 186 (41.1%) of the insertions being fixed, 202 (44.6%) polymorphic and 65 (14.3%) unique. We further explored polymorphic and fixed insertions and identified binding sites for *dl* and *STAT* and promoter motifs. We found that 19 TEs contained bindings sites for *dl*, 21 TEs contained binding sites for *STAT* and 12 TEs contained binding sites both for *dl* and *STAT* (Additional file 1: Table S15). Additionally, we identified 81 insertions, in the upstream region of 56 genes, which carried putative promoter sequences.

We identified TE insertions in three different antimicrobial peptides (AMPs). AMPs form the first line of host defense against infection and are a key component of the innate immune system, however none had transcription factor binding sites (TFBS) for *dl* or *STAT*. It is important to keep in mind that there are other TF that participate in the regulation of AMPs and that both *dl* and *STAT* are also involved in other biological processes (105). Interestingly we also identified TEs with TFBS for *dl* in the vicinity of *STAT1* and *STAT2* which might lead to novel regulatory mechanisms of the JAK/STAT signaling pathway. Furthermore, 11 of the 156 genes containing TE insertions are differentially expressed in response to a *Plasmodium* invasion. These genes participate in several pathways of the immune response including the small regulatory RNA pathway, pathogen recognition, the nitric oxide response and ookinete melanization (78, 106-108). Four of the TEs affecting these genes added TFBS and promoter sequences, thus suggesting that these TE insertions can potentially influence the response to this pathogen (109) (Table 4).

**Table 4.**
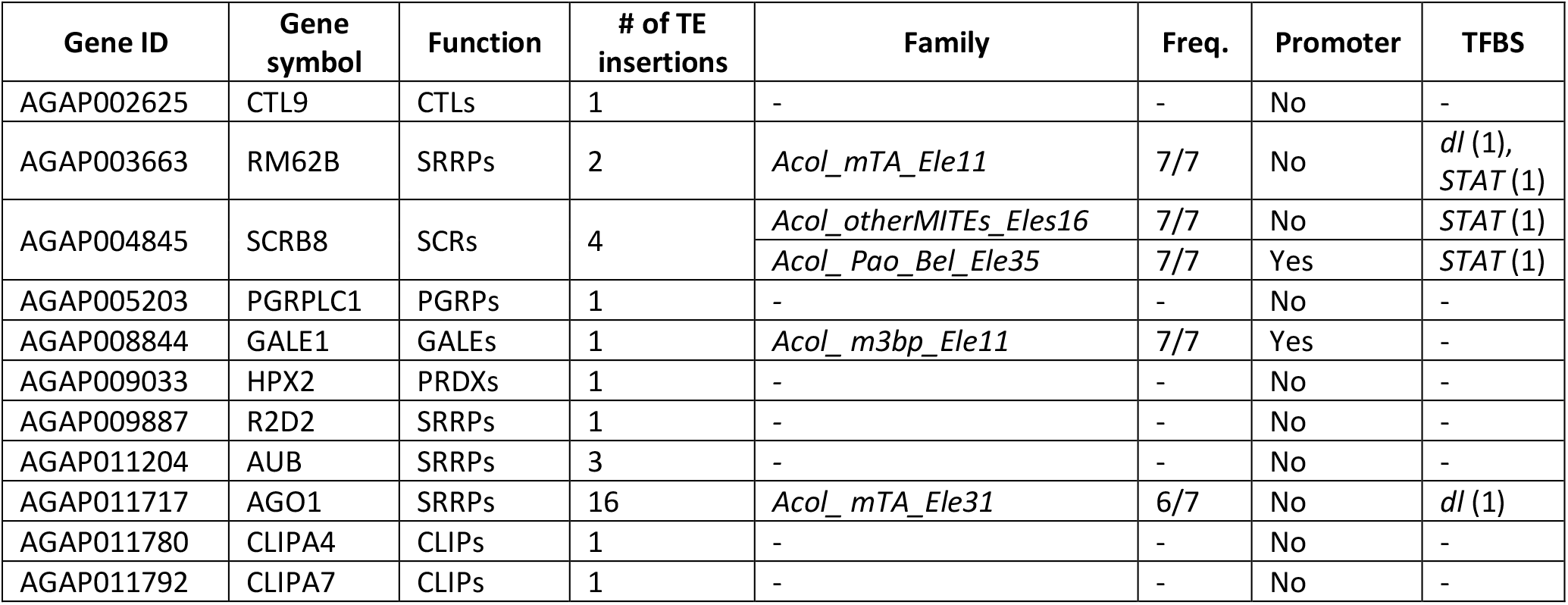
TE insertions in *Plasmodium* responsive genes from the immune system. Family and frequency are only shown for TEs with TFBS or promoter sequences. In the Function column the following abbreviations are used: C-Type Lectins (CTLs), Small Regulatory RNA Pathway Members (SRRPs), Scavenger Receptors (SCRs), Peptidoglycan Recognition Proteins (PGRPs), Galactoside-Binding Lectins (GALEs), Peroxidases (PRDXs), CLIP-Domain Serine Proteases (CLIPs).

## Discussion

In this study, we *de novo* annotated transposable element (TE) insertions in seven genomes of *An. coluzzii*, six of them sequenced here. A comprehensive genome-wide TE annotation was possible because we used long-read technologies to perform the genome sequencing and assembly. Long-reads allow identifying TE insertions with high confidence given that the entire TE insertion sequence can be spanned by a single read (30, 31). While the genome-wide TE repertoire has been studied in other anopheline species, particularly in *An. gambiae*, to our knowledge there are no other studies that have explored TE variation in multiple genomes from a single species (32, 33, 36, 40, 72, 110). We observed that increasing the number of available genomes analyzed allowed us to increase the number of identified TE families from a median of 244 (172-294) to 435 (Figure 1B). Moreover, having the full sequences of seven genomes also allowed us to discover 64 new TE families, including four TRIM families previously undescribed in anopheline genomes. This might be relevant as TRIM elements have been shown to be important players in genome evolution in other species (57, 58). The wide range of families identified across genomes was not directly related to the quality of the genome assembly taking into consideration the more generally used quality parameters such as read length, number of contigs, and contig N50 (111). This suggests that there are possibly other characteristics of each genome that affect the identification of high-quality TE families, such as biases in the location of the TE insertions given that TE families are challenging to identify in regions with low complexity or with numerous nested TEs. Nonetheless, the identification of TE families is dependent on the methodology used to perform TE annotations, therefore different annotation strategies could lead to the discovery of still undescribed families (60).

The availability of several genome assemblies also allowed us to determine that the majority of the intraspecies differences in the TE content were in heterochromatic regions, most likely due to differences in the quality of the genome assembly. Nevertheless, there were also significant differences in the TE content in euchromatic regions, reflecting true intraspecific variability as has been previously observed in several organisms including Drosophila (112, 113), mammals (114, 115), maize (116) and Arabidopsis (117). TE insertions were not randomly distributed throughout the genome and instead were consistently enriched in intergenic regions, most likely due to purifying selection, as suggested in the wild grass *Brachypodium distachyon* (118). In Drosophila, TE enrichment in intergenic regions was also observed in addition to enrichment in the intronic region, which we did not observe in *An. coluzzii* (119). We also analyzed the TE content in the breakpoints of five common polymorphic inversions, three of them analyzed here for the first time. We found TE insertions in all but one of the inversion breakpoints, with MITE elements being the most common TE family, as already described in the 2Rd’ inversion in *An. arabiensis* (26) (Figure 5).

As a first step towards identifying the potential role of TEs in rapid adaptation to novel habitats (120), we focused on insertions from recently active families located near genes that are relevant for the vectorial capacity of *An. coluzzii* (Table 3). Because adaptation can also happen from standing variation, in the case of insecticide resistance genes, which have been shown to be shaped by TE insertions in several organisms, and immune-related genes, we analyzed all insertions independently of their age (Figure 6 and Table 4) (99, 121, 122). While the role of nonsynonymous substitutions and copy number variation in resistance to insecticides commonly used in urban environments has been studied, the potential role of TEs has not yet been comprehensively assessed in *An. coluzzii* or any other anopheline species (19, 102, 123-125). In the genomes we assessed, we identified several insertions that were polymorphic or fixed nearby functionally relevant genes (Table 3, Table 4 and Figure 6). Some of the identified candidate insertions contained binding sites for transcription factors related to the function of the nearby genes and promoter regions. Besides adding regulatory regions, TEs can also affect the regulation of nearby genes by affecting gene splicing and generating long non-coding RNAs among many other molecular mechanisms (25, 126-129). Thus, it is possible that the candidate TE insertions identified, which lack binding sites and promoters, could be affecting nearby genes through other molecular mechanisms. Our results are a first approximation to the potential role of TEs in *An. coluzzii* adaptation to the challenging environment that urban ecosystems entail. Establishing a direct link between the TEs and the traits involved in urban adaptation will require sampling a larger number of individuals and characterizing the phenotypes associated with the insertions.

## Conclusions

The long-read sequencing of seven *An. coluzzii* genomes from urban environments allowed us to capture to a larger extent the diversity of TE families and TE insertions and to assess their potential impact in the genome architecture and genome function in this species. While there was an enrichment of TE insertions in intergenic regions, we found several insertions located in the 1 kb flanking regions or inside genes relevant for the vectorial capacity of this species. Furthermore, we found that some of these TE insertions are adding regulatory regions suggesting that they could influence the regulation of these genes. Further studies are needed to confirm the potential functional effect of these insertions. The genomic resources and the results that we present in this work provide a basis for future studies of the impact of TEs in the biology of *An. coluzzii*. This will allow increasing our knowledge on a species which besides being interesting from an evolutionary perspective, given its high levels of genetic diversity and the strong anthropogenic pressures it faces, is of great importance to human health. A better understanding of the biology of *An. coluzzii* and its ability to rapidly adapt to urban environments will further facilitate the development of novel strategies to combat malaria. Better management strategies can be implemented if we understand and are able to predict changes in the frequency of genetic variants relevant for the vectorial capacity of this species.

## Data accessibility

All the genome sequencing data obtained in this work, as well as the genome assemblies are available in NCBI SRA and NCBI Genbank respectively, under the BioProject accession number PRJNA676011. TE annotations in each of the seven genomes in gff files, transferred annotations across the seven genomes in txt format, and the TE library in fasta format are also available at https://digital.csic.es/handle/10261/224416.

## Acknowledgements

We thank members of the González Lab for comments on the manuscript. We thank the Ecology of Vectorial Systems team at the CIMRF (Franceville, Gabon) for their support in field collections. We thank Jean Pierre Agbor and Serge Donfanck for their commitment in larvae collections in Douala (Cameroon). Version 3 of this preprint has been peer-reviewed and recommended by Peer Community In Genomics (https://doi.org/10.24072/pci.genomics.100006)

## Appendix

File name: Additional file 1

File format: Microsoft Excel Binary File format (xls)

Title of data: Supplementary Tables

Description of data: Supplementary Tables

File name: Additional file 2

File format: Portable document format (pdf)

Title of data: Figure S1. Number of TE copies identified when using the TE libraries from an increasing number of genomes.

Description of data: Number of TE copies identified when using the TE library of a single genome or when using all possible combinations of more than one genome.

File name: Additional file 3

File format: Portable document format (pdf)

Title of data: Figure S2. Novel TE families

Description of data: Newly described families. A) The structure of each new family is displayed: the light blue box represents the full extension of the TE and the red arrows represent LTRs. B) All insertions for each TE family are shown as a coverage plot where each line represents a copy in a genome. C) Phylogenetic distribution of the TE family insertions in 15 members of the Anopheles genus, Culex quinquefasciatus, Ae. Aegypti and D. melanogaster. The number of insertions with more than 80% identity and spanning at least 80% of the consensus, in each species is shown using a black and white gradient. Species with no insertions are shown in white while species with 50 or more insertions are shown in black.

File name: Additional file 4

File format: Portable document format (pdf)

Title of data: Figure S3. Number of TE insertions vs genome size

Description of data: Comparison of the bases spanned by TEs in each genome with their full genome sizes.

File name: Additional file 5

File format: Portable document format (pdf)

Title of data: Figure S4. TE landscapes

Description of data: TE landscapes for the six genomes sequenced in this work generated using dnaPipeTE

File name: Additional file 6

File format: Portable document format (pdf)

Title of data: Figure S5. Genes with TE insertions from active families

Description of data: Diagrams of TE insertions closer than 1 kb to genes showing the gene structure and the TE insertion

File name: Additional file 7

File format: Portable document format (pdf)

Title of data: Figure S6. Genes associated with insecticide resistance with TE insertions

Description of data: Diagrams of genes associated with insecticide resistance showing the gene structure and the TE insertions closer than 1 kb to gene.

## Cite as

Vargas-Chavez, C., Longo Pendy, N. M., Nsango, S., Aguilera, L., Ayala, D., and González, J. (2021). Uncovering transposable element variants and their potential adaptive impact in urban populations of the malaria vector Anopheles coluzzii. bioRxiv, 2020.11.22.393231, ver. 3 peer-reviewed and recommended by Peer community in Genomics.

