## Supplementary figures and images for "Uncovering transposable element variants and their potential adaptive impact in urban populations of the malaria vector *Anopheles coluzzii*"

### Additional File 2

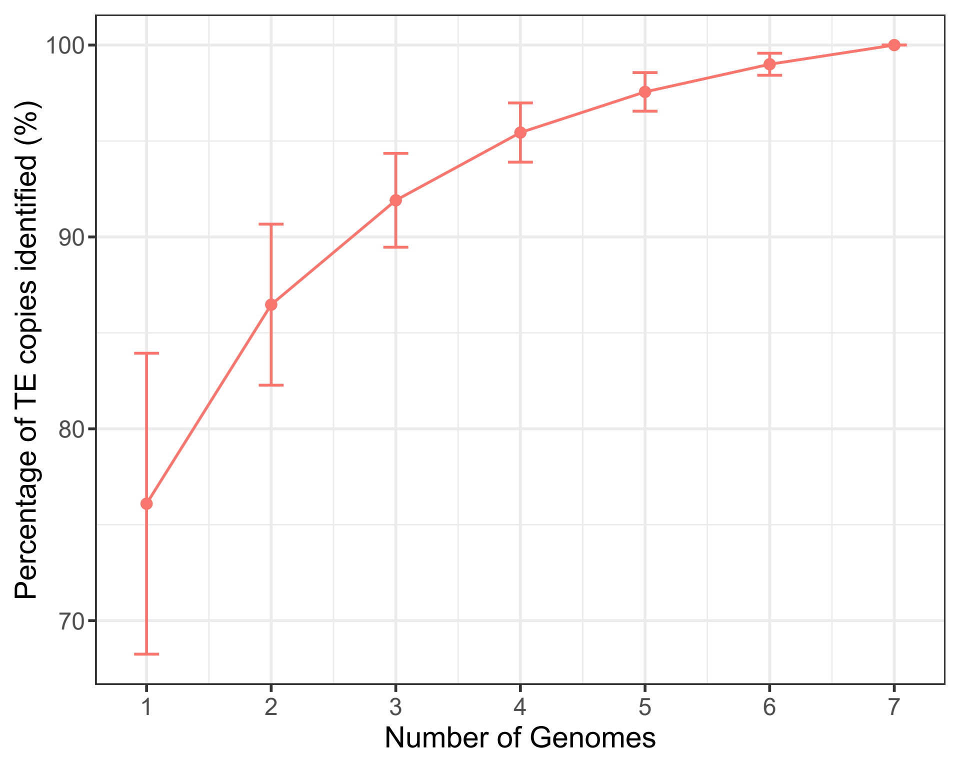

### Additional File 3

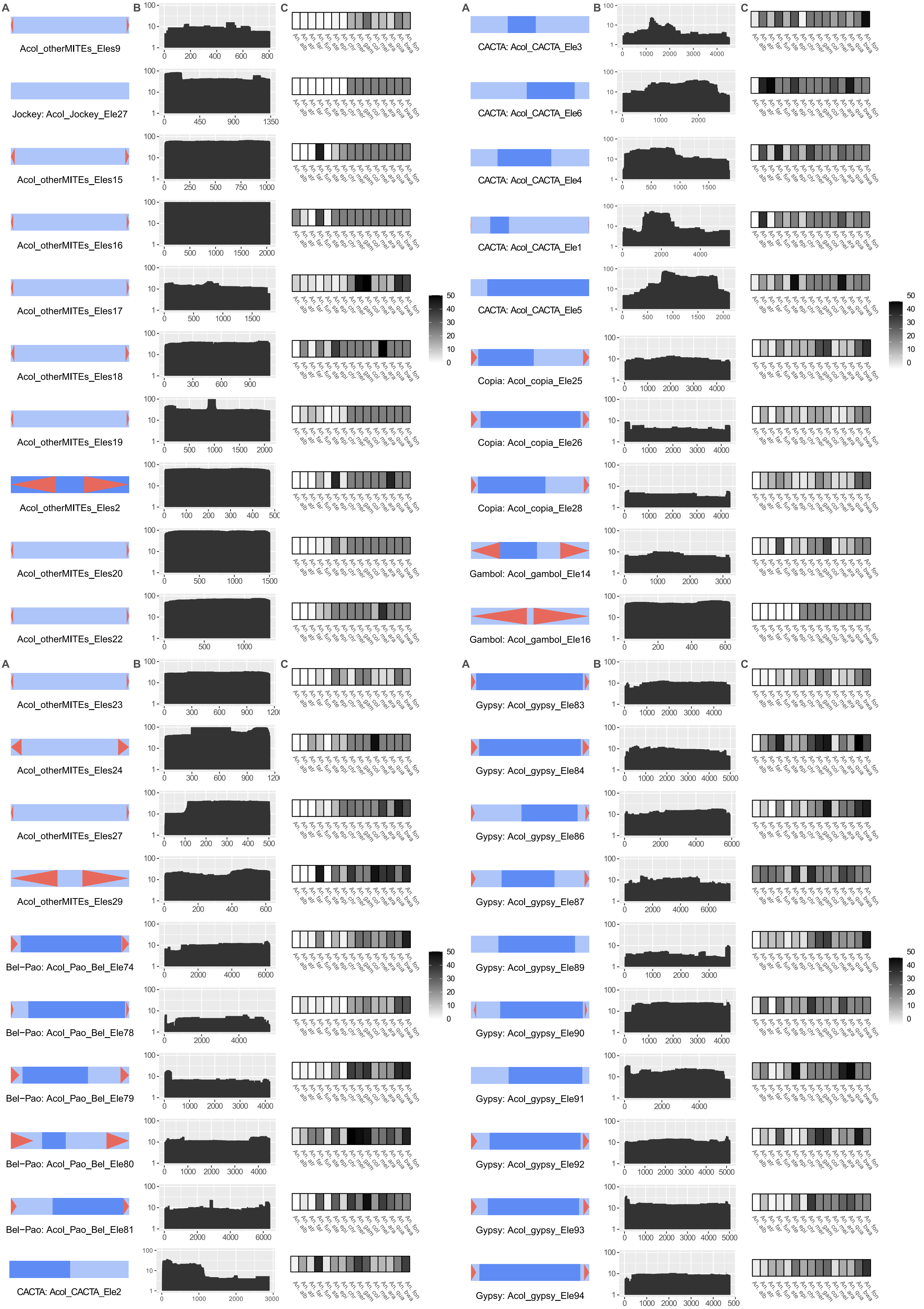

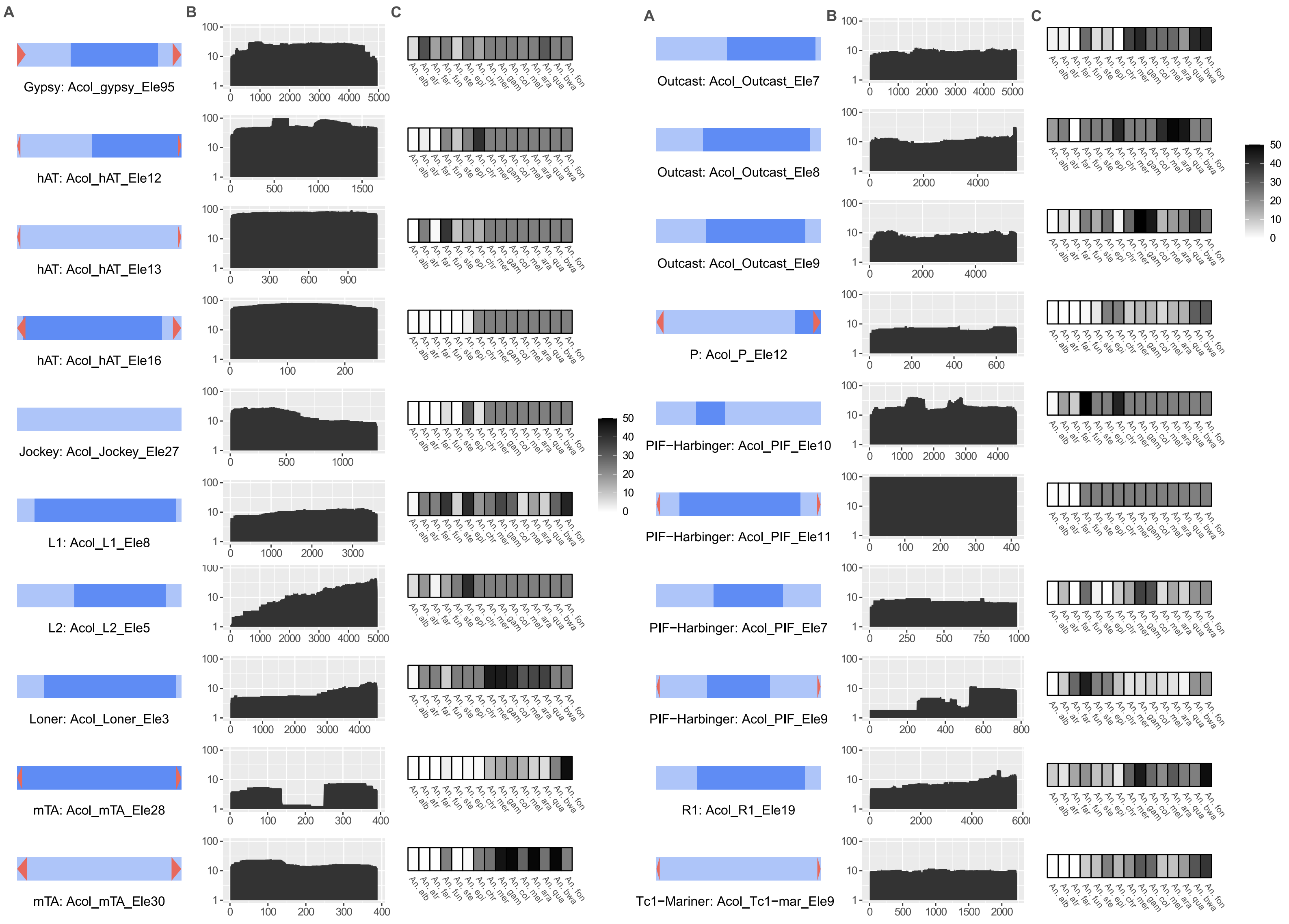

### Additional File 4

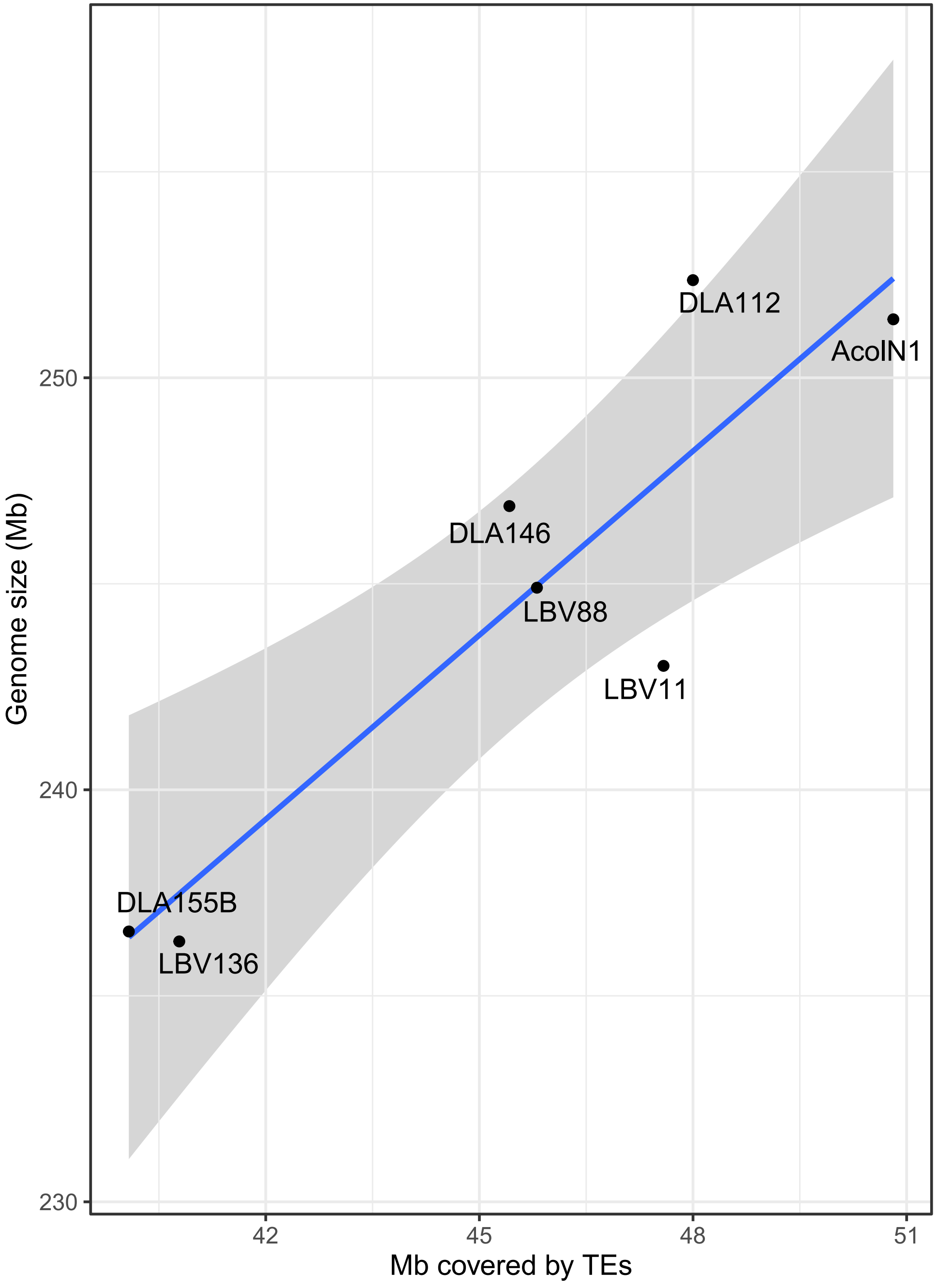

### Additional File 5

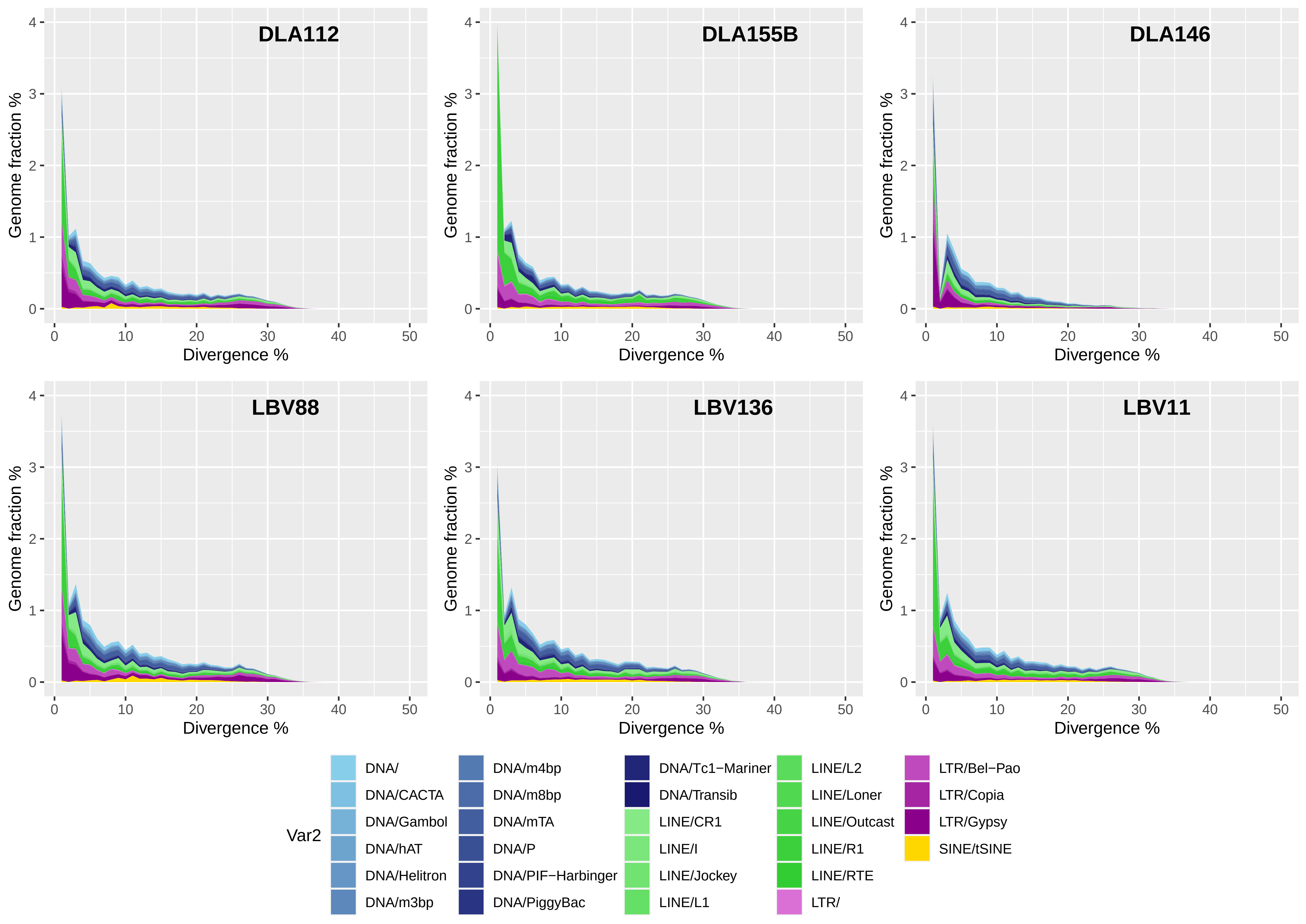
