## Additional File 6 for "Uncovering transposable element variants and their potential adaptive impact in urban populations of the malaria vector *Anopheles coluzzii*"

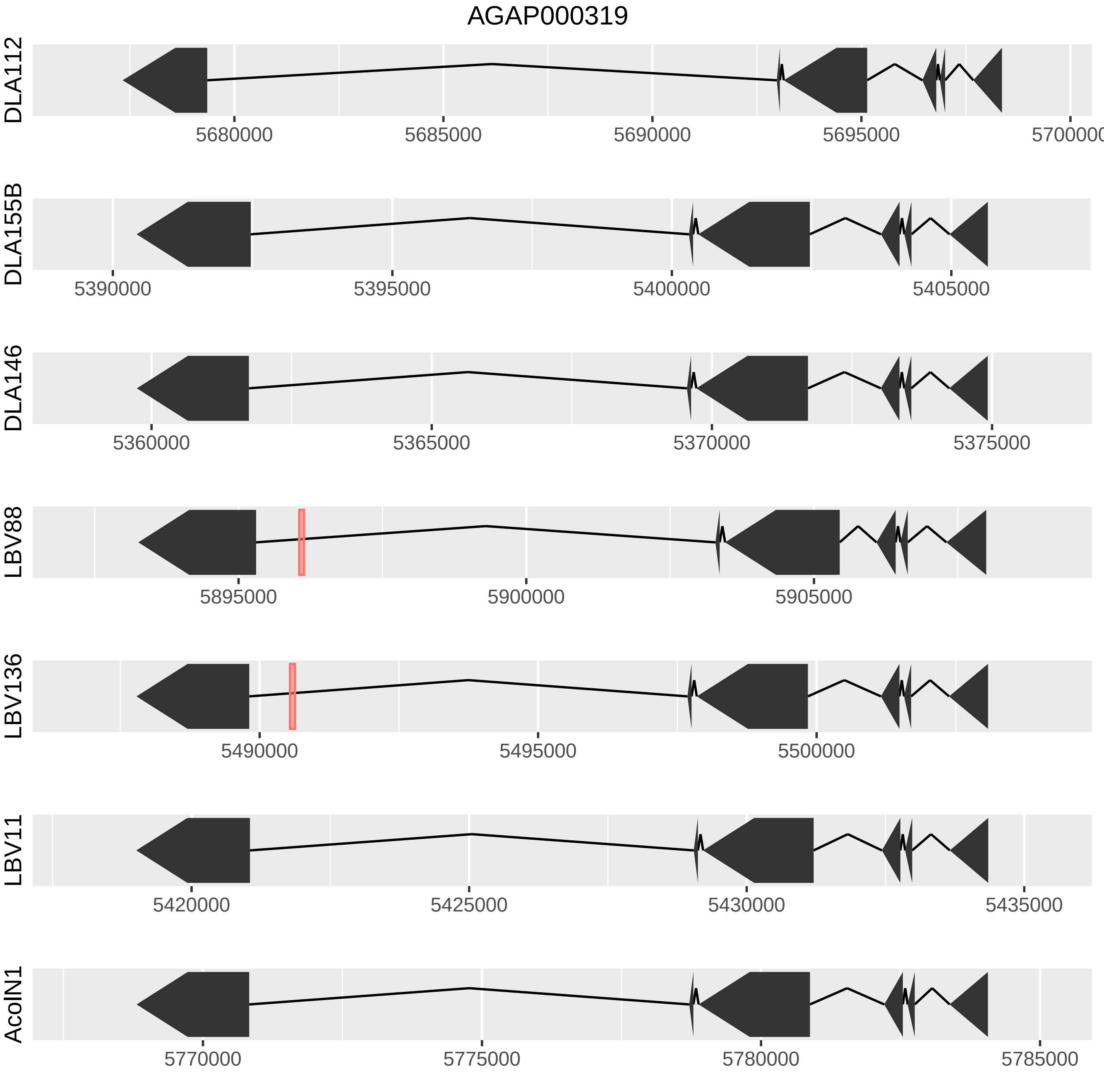

AGAP000319
DLA112
5680000
5685000
5690000
5695000
5700000
DLA155B
5390000
5395000
5400000
5405000
DLA146
5360000
5365000
5370000
5375000
LBV88
5895000
5900000
5905000
LBV136
5490000
5495000
5500000
LBV11
5420000
5425000
5430000
5435000
AcolN1
5770000
5775000
5780000
5785000

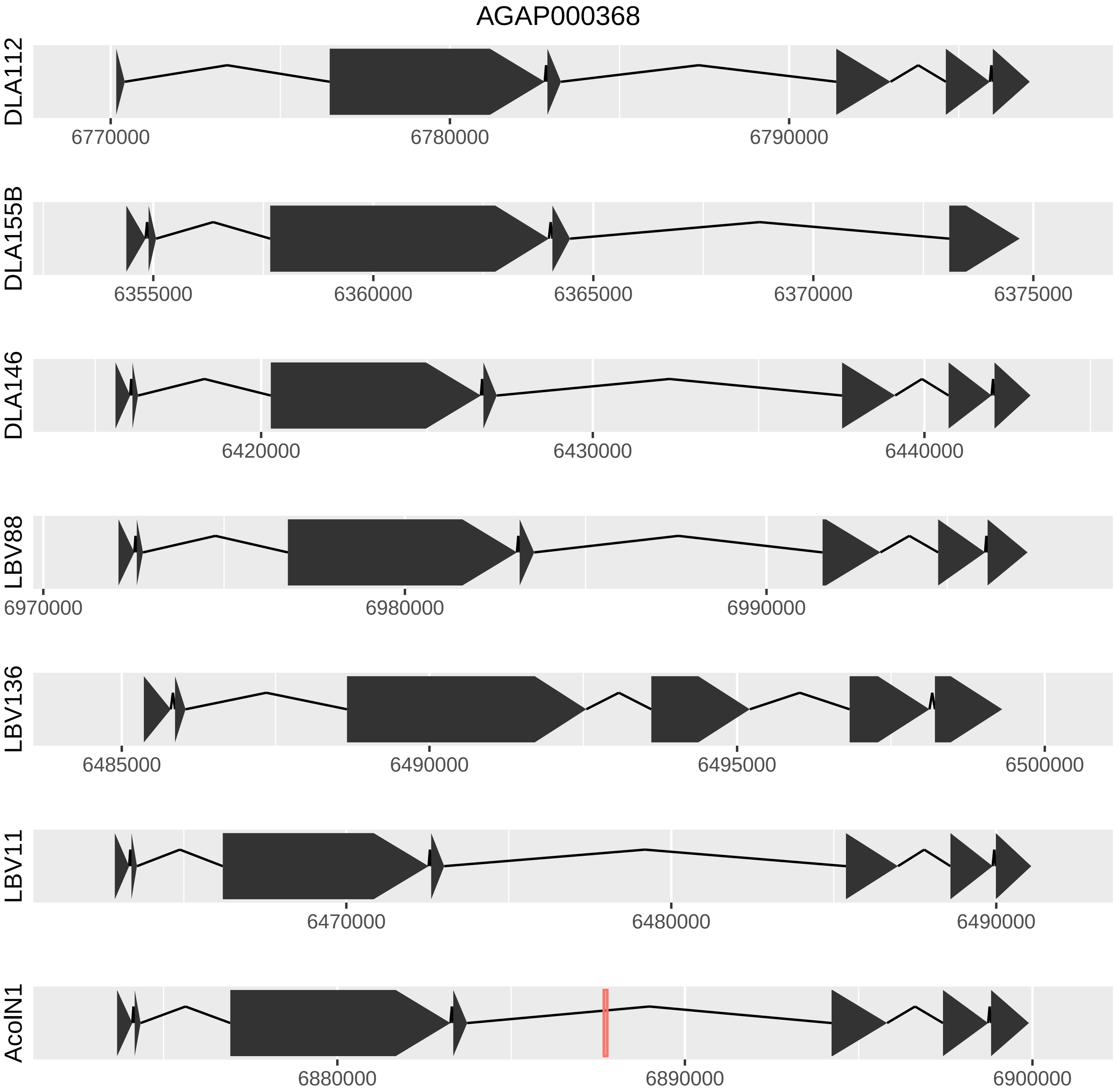
AGAP000368
DLA112
6770000
6780000
6790000
DLA155B
6355000
6360000
6365000
6370000
6375000
DLA146
6420000
6430000
6440000
LBV88
6970000
6980000
6990000
LBV136
6485000
6490000
6495000
6500000
LBV11
6470000
6480000
6490000
AcolN1
6880000
6890000
6900000

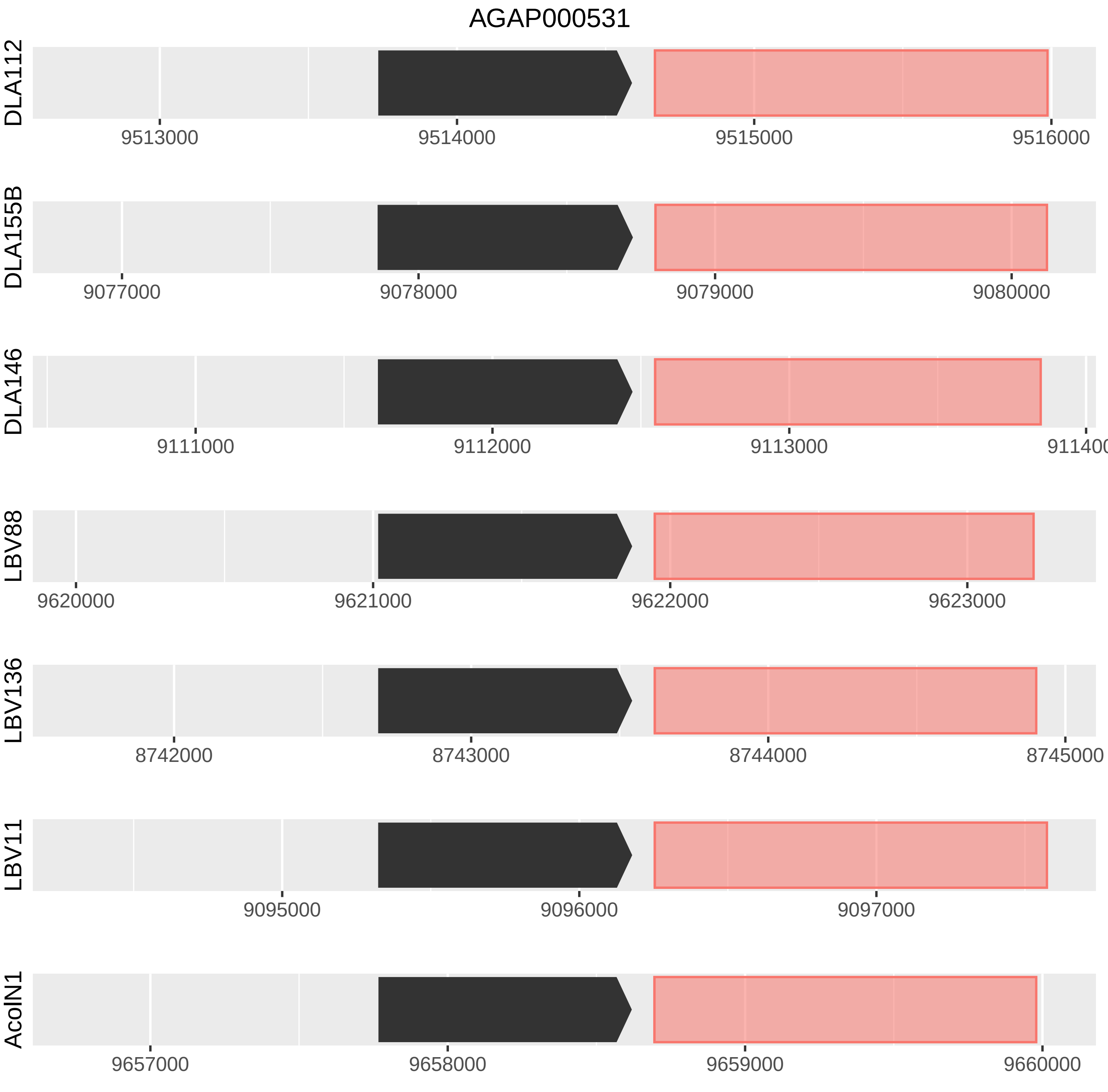
AGAP000531
DLA112
9513000
9514000
9515000
9516000
DLA155B
9077000
9078000
9079000
9080000
DLA146
9111000
9112000
9113000
LBV88
9620000
9621000
9622000
9623000
LBV136
8742000
8743000
8744000
8745000
LBV11
9095000
9096000
9097000
AcolN1
9657000
9658000
9659000
9660000

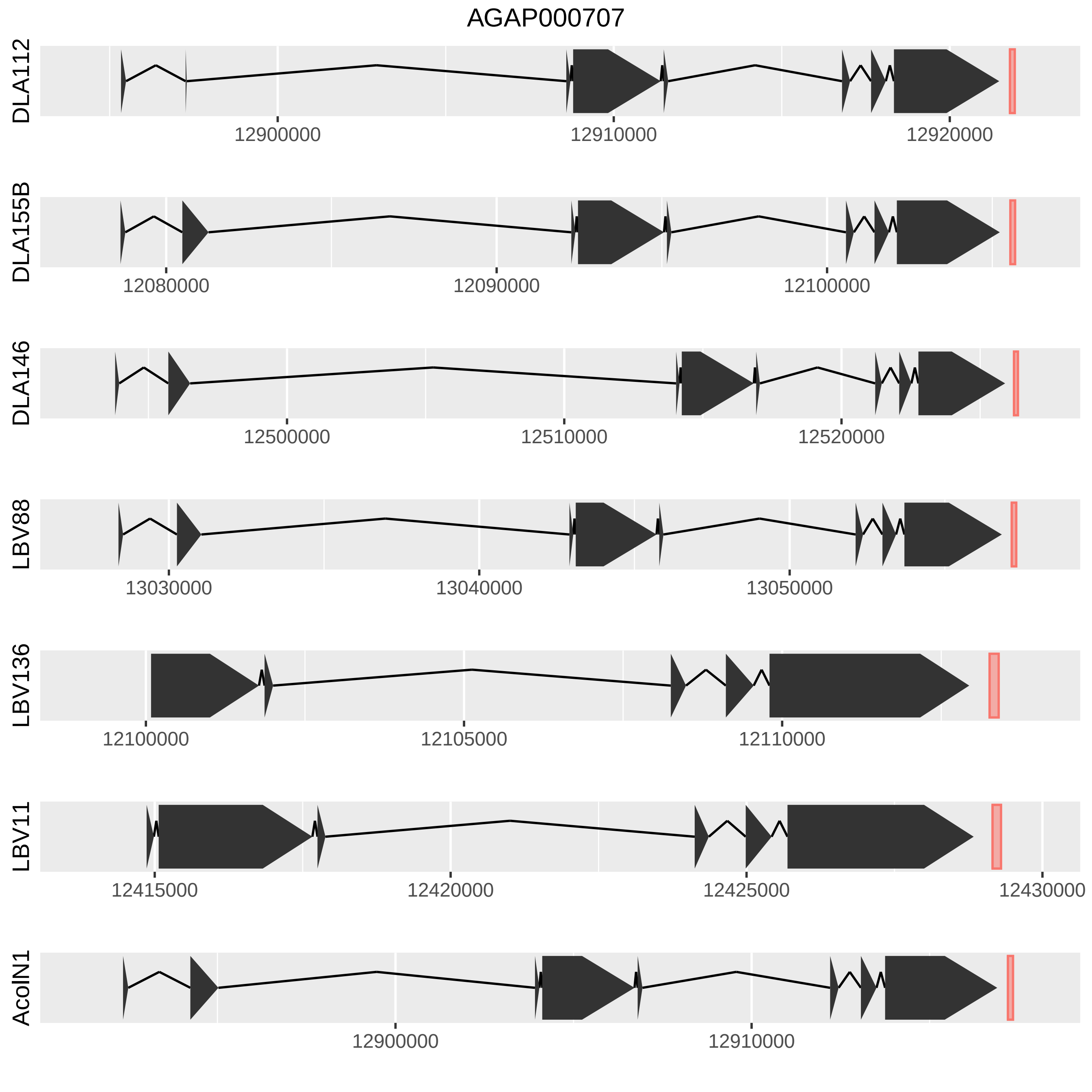
AGAP000707
DLA112
12900000
12910000
12920000
DLA155B
12080000
12090000
12100000
DLA146
12500000
12510000
12520000
LBV88
13030000
13040000
13050000
LBV136
12100000
12105000
12110000
LBV11
12415000
12420000
12425000
12430000
AcoIN1
12900000
12910000

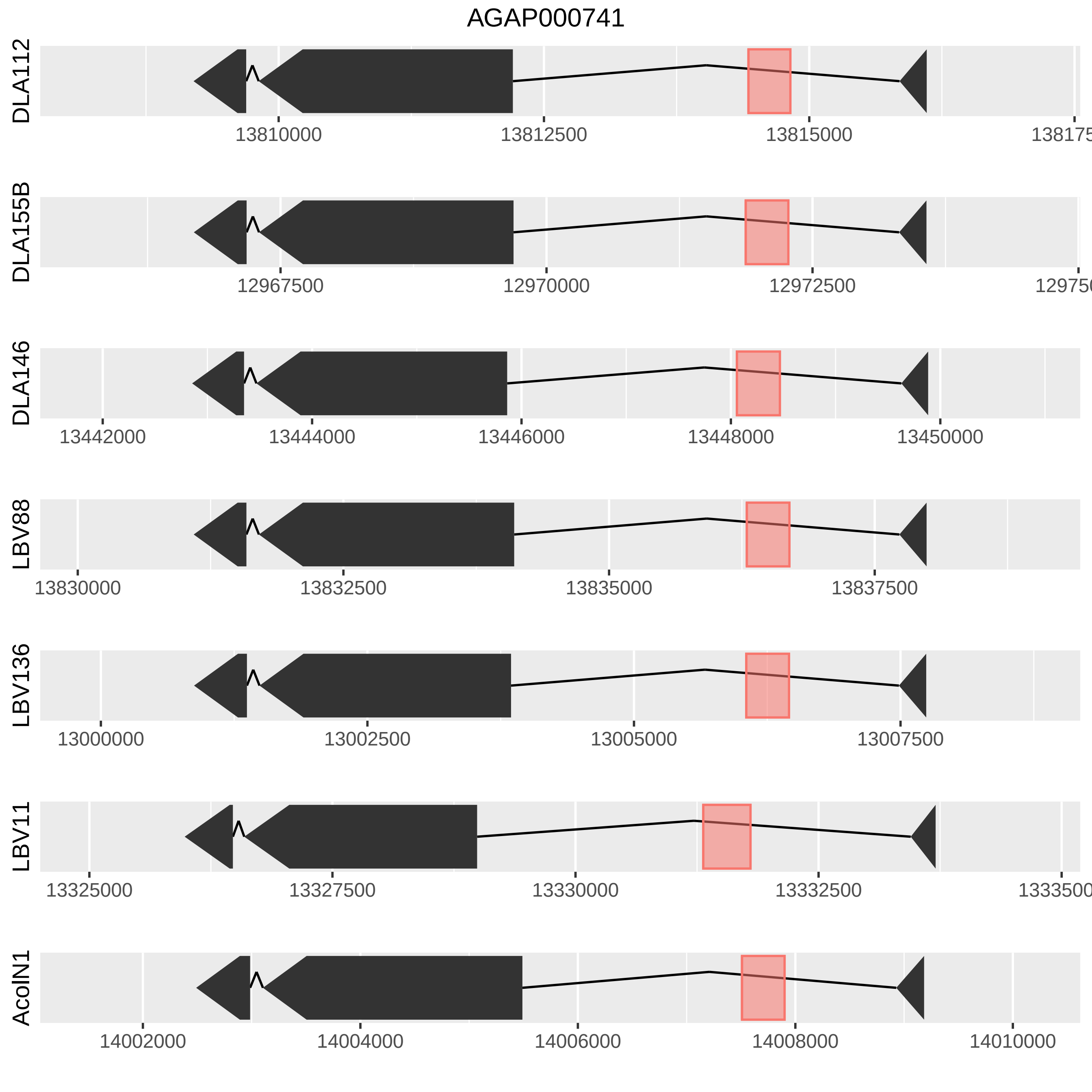
AGAP000741
DLA112
13810000
13812500
13815000
DLA155B
12967500
12970000
12972500
DLA146
13442000
13444000
13446000
13448000
13450000
LBV88
13830000
13832500
13835000
13837500
LBV136
13000000
13002500
13005000
13007500
LBV11
13325000
13327500
13330000
13332500
AcolN1
14002000
14004000
14006000
14008000
14010000

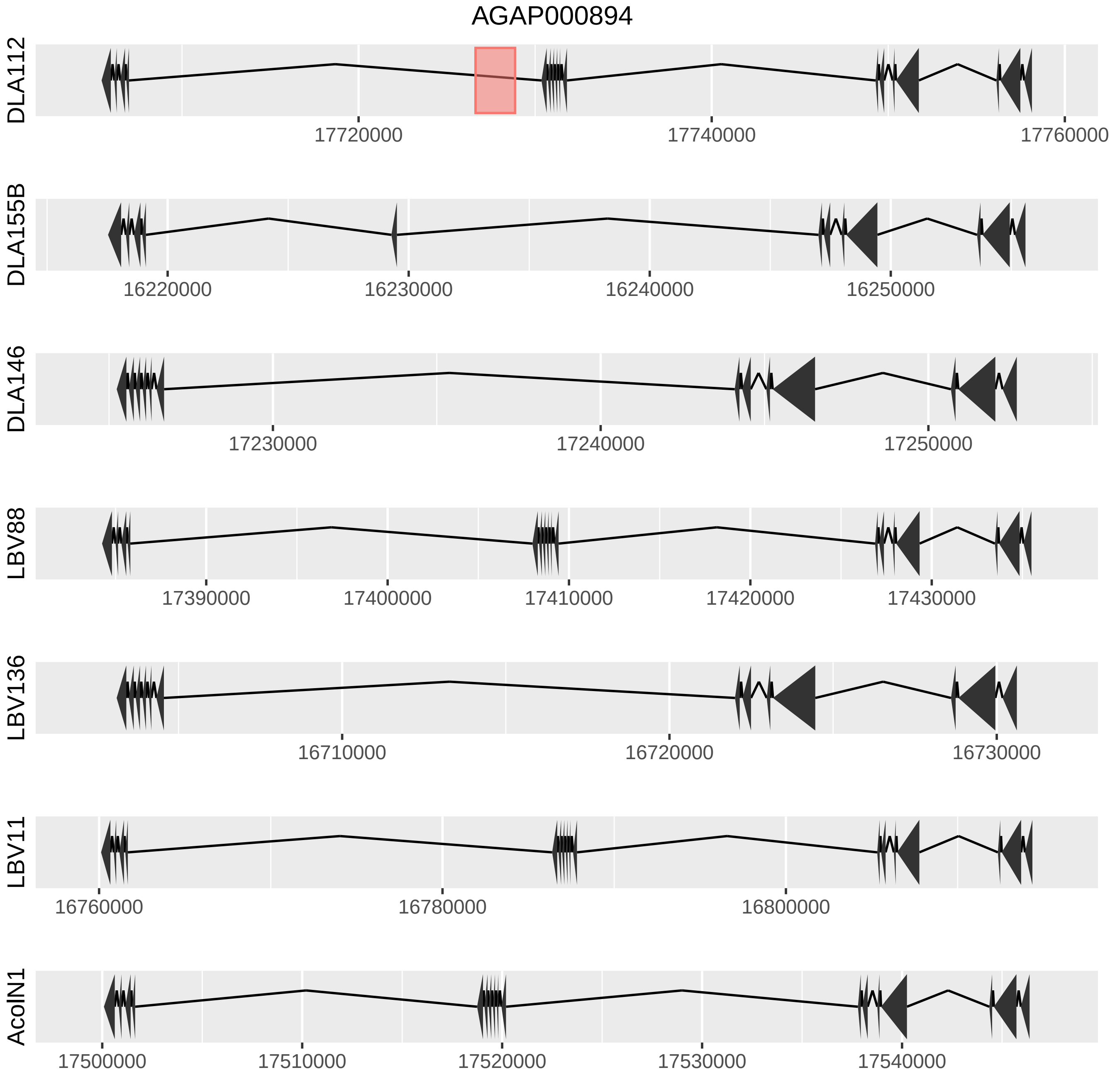
AGAP000894
DLA112
17720000
17740000
17760000
DLA155B
16220000
16230000
16240000
16250000
DLA146
17230000
17240000
17250000
LBV88
17390000
17400000
17410000
17420000
17430000
LBV136
16710000
16720000
16730000
LBV11
16760000
16780000
16800000
AcolN1
17500000
17510000
17520000
17530000
17540000

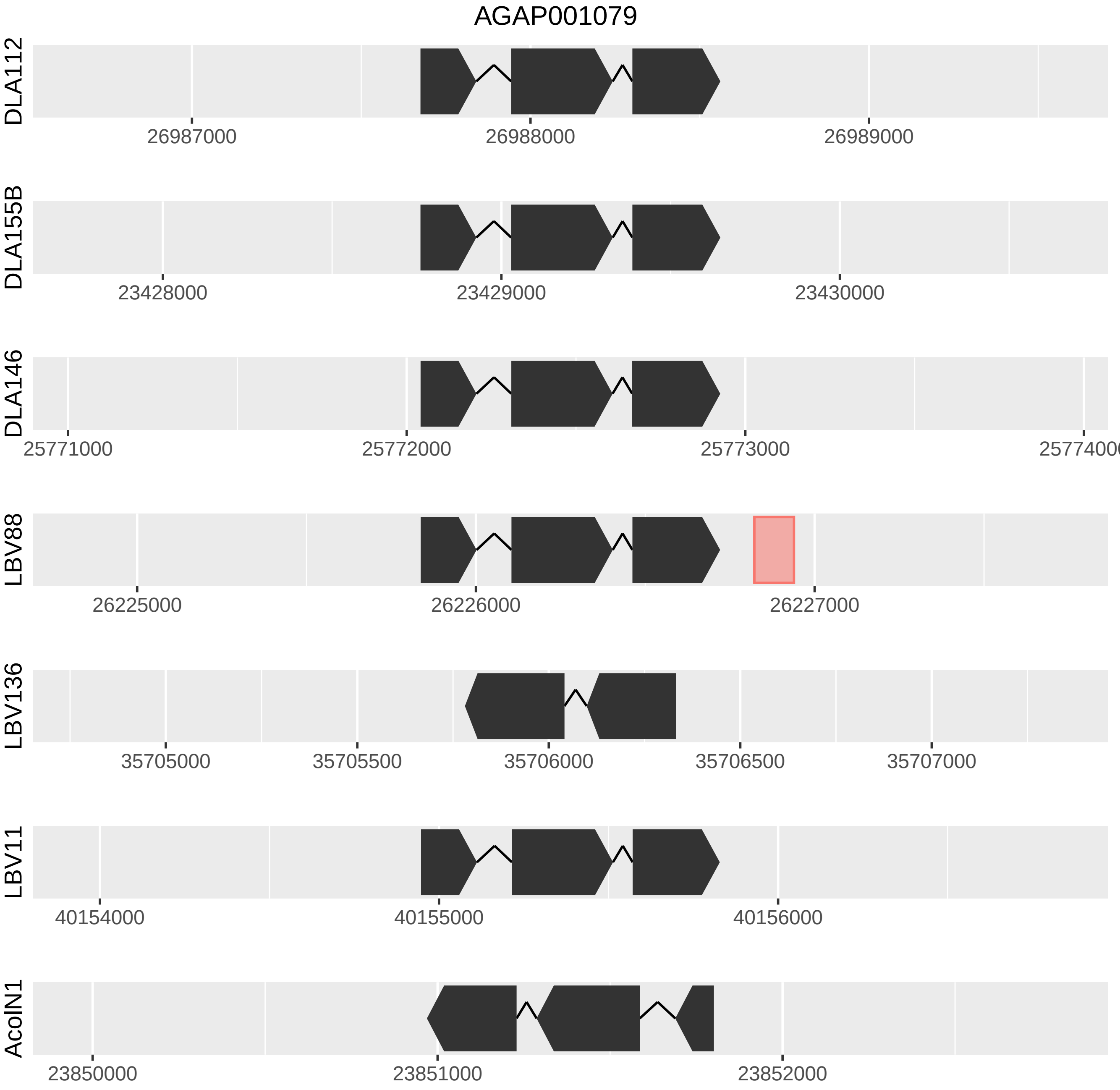
AGAP001079
DLA112
26987000
26988000
26989000
DLA155B
23428000
23429000
23430000
DLA146
25771000
25772000
25773000
LBV88
26225000
26226000
26227000
LBV136
35705000
35705500
35706000
35706500
35707000
LBV11
40154000
40155000
40156000
AcolN1
23850000
23851000
23852000

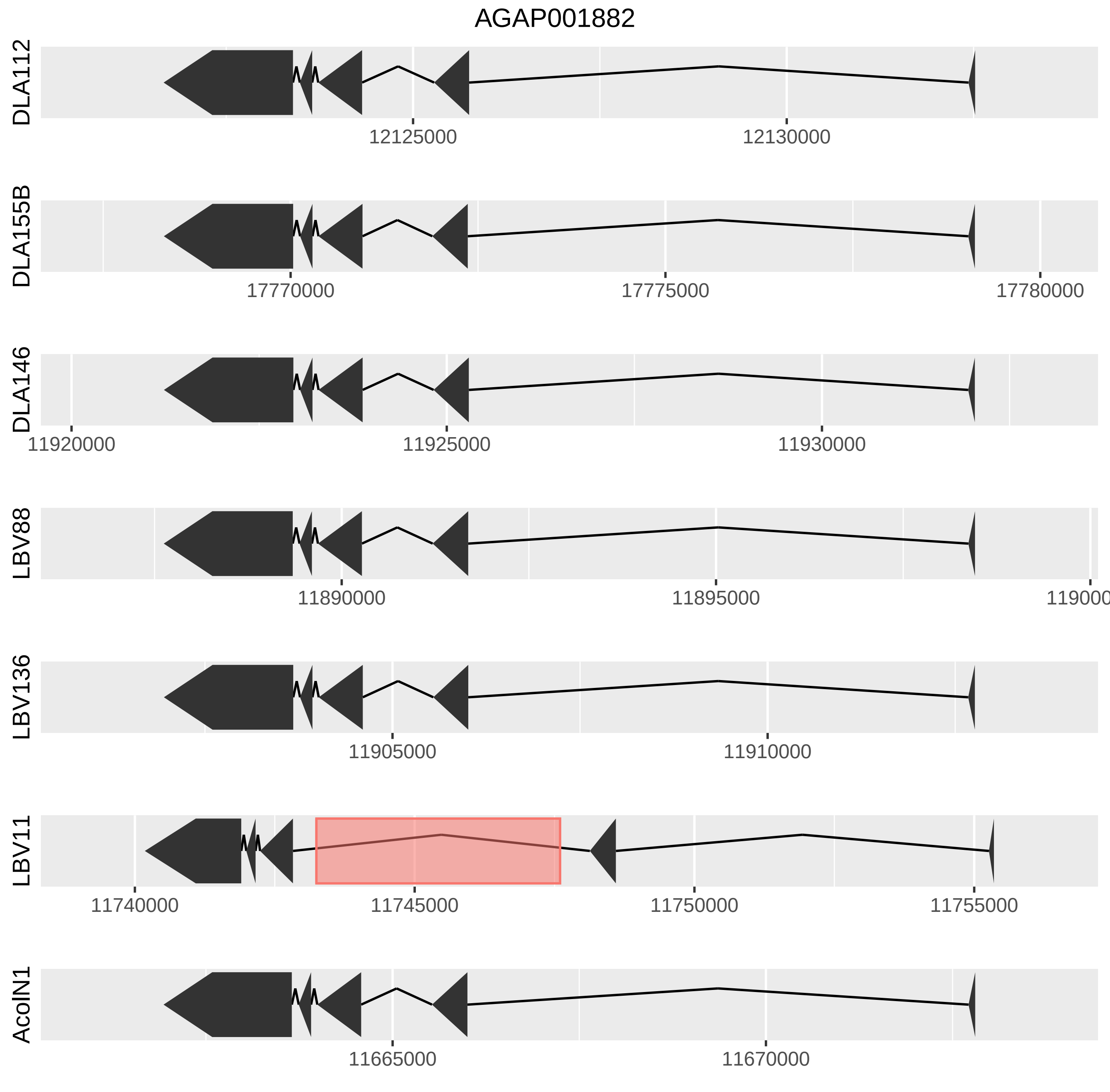
AGAP001882
DLA112
12125000
12130000
DLA155B
17770000
17775000
17780000
DLA146
11920000
11925000
11930000
LBV88
11890000
11895000
LBV136
11905000
11910000
LBV11
11740000
11745000
11750000
11755000
AcolN1
11665000
11670000

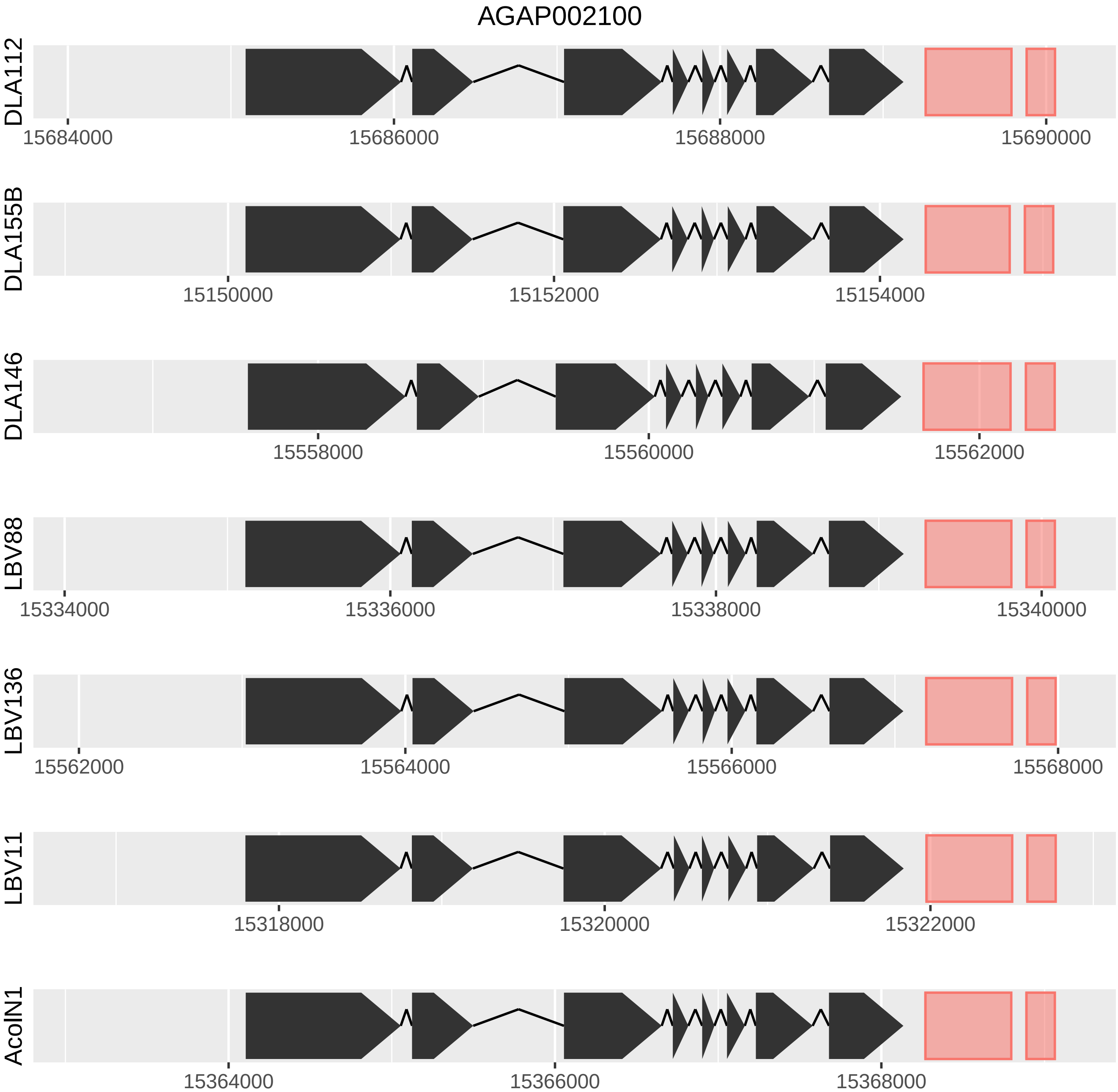
AGAP002100
DLA112
15684000
15686000
15688000
15690000
DLA155B
15150000
15152000
15154000
DLA146
15558000
15560000
15562000
LBV88
15334000
15336000
15338000
15340000
LBV136
15562000
15564000
15566000
15568000
LBV11
15318000
15320000
15322000
AcoIN1
15364000
15366000
15368000

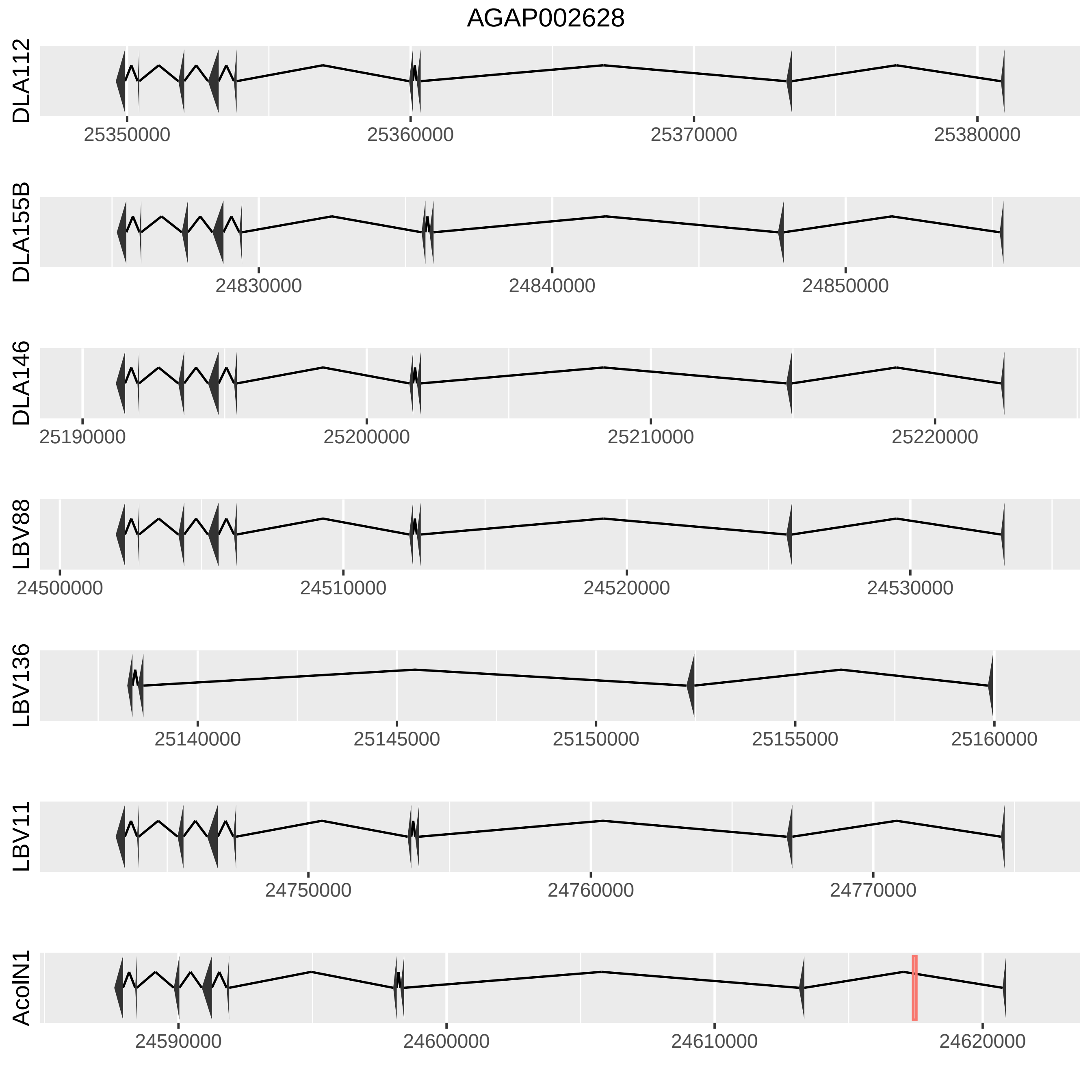

AGAP002628
DLA112
25350000
25360000
25370000
25380000
DLA155B
24830000
24840000
24850000
DLA146
25190000
25200000
25210000
25220000
LBV88
24500000
24510000
24520000
24530000
LBV136
25140000
25145000
25150000
25155000
25160000
LBV11
24750000
24760000
24770000
AcolN1
24590000
24600000
24610000
24620000

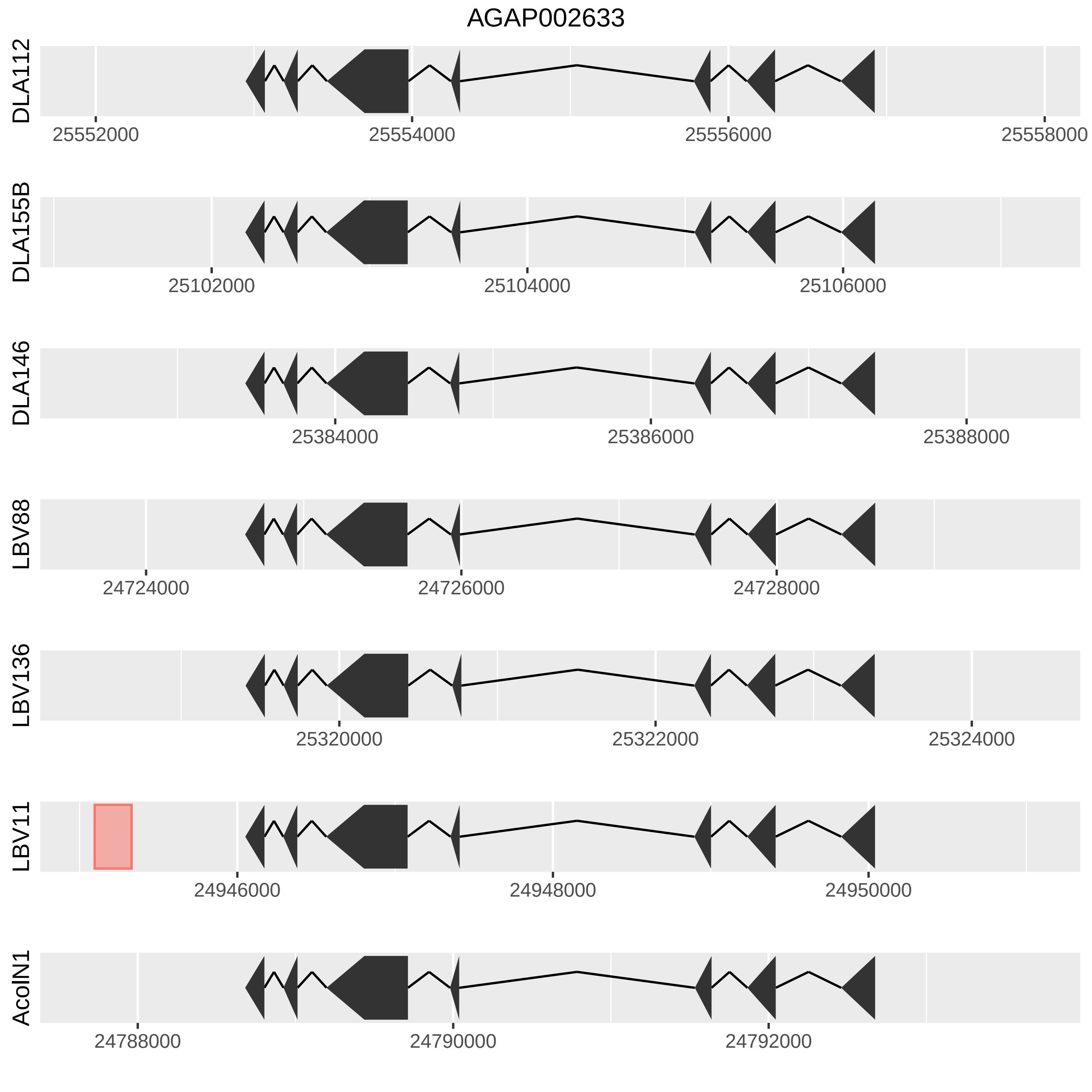

AGAP002633
DLA112
25552000
25554000
25556000
25558000
DLA155B
25102000
25104000
25106000
DLA146
25384000
25386000
25388000
LBV88
24724000
24726000
24728000
LBV136
25320000
25322000
25324000
LBV11
24946000
24948000
24950000
AcolN1
24788000
24790000
24792000

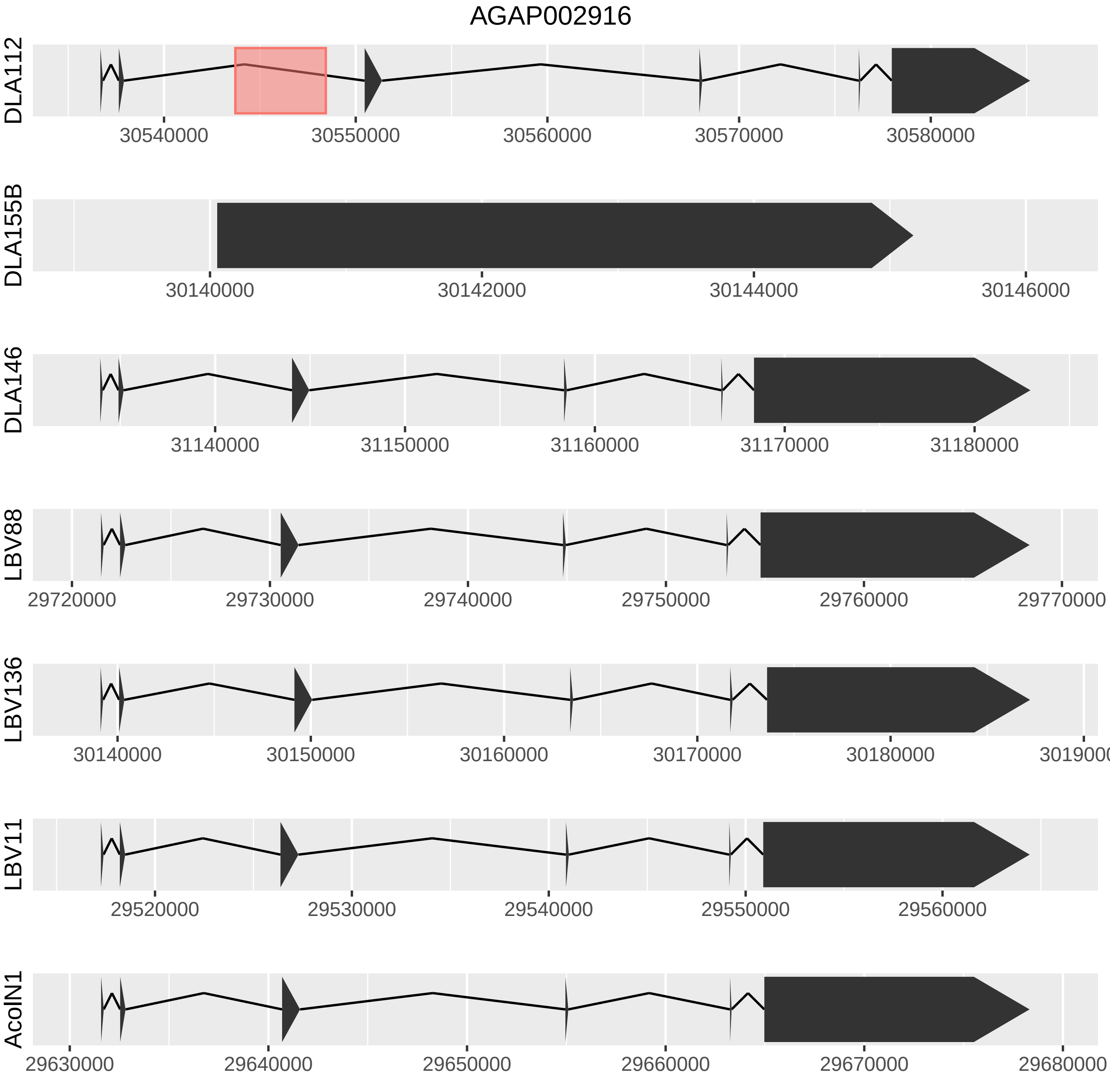

AGAP002916
DLA112
30540000
30550000
30560000
30570000
30580000
DLA155B
30140000
30142000
30144000
30146000
DLA146
31140000
31150000
31160000
31170000
31180000
LBV88
29720000
29730000
29740000
29750000
29760000
29770000
LBV136
30140000
30150000
30160000
30170000
30180000
3019000
LBV11
29520000
29530000
29540000
29550000
29560000
AcolN1
29630000
29640000
29650000
29660000
29670000
29680000

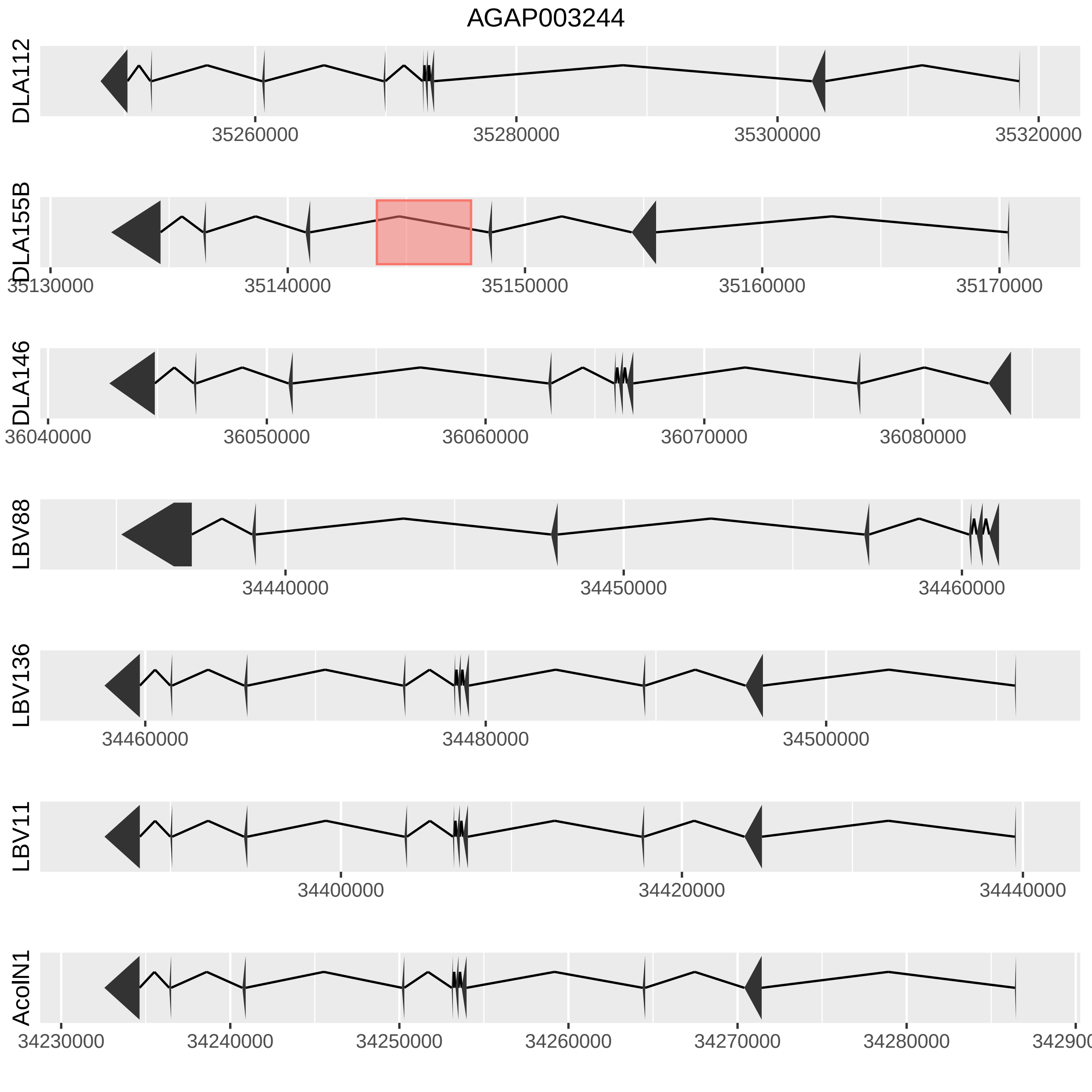
AGAP003244
DLA112
35260000
35280000
35300000
35320000
DLA155B
35130000
35140000
35150000
35160000
35170000
DLA146
36040000
36050000
36060000
36070000
36080000
LBV88
34440000
34450000
34460000
LBV136
34460000
34480000
34500000
LBV11
34400000
34420000
34440000
AcolN1
34230000
34240000
34250000
34260000
34270000
34280000

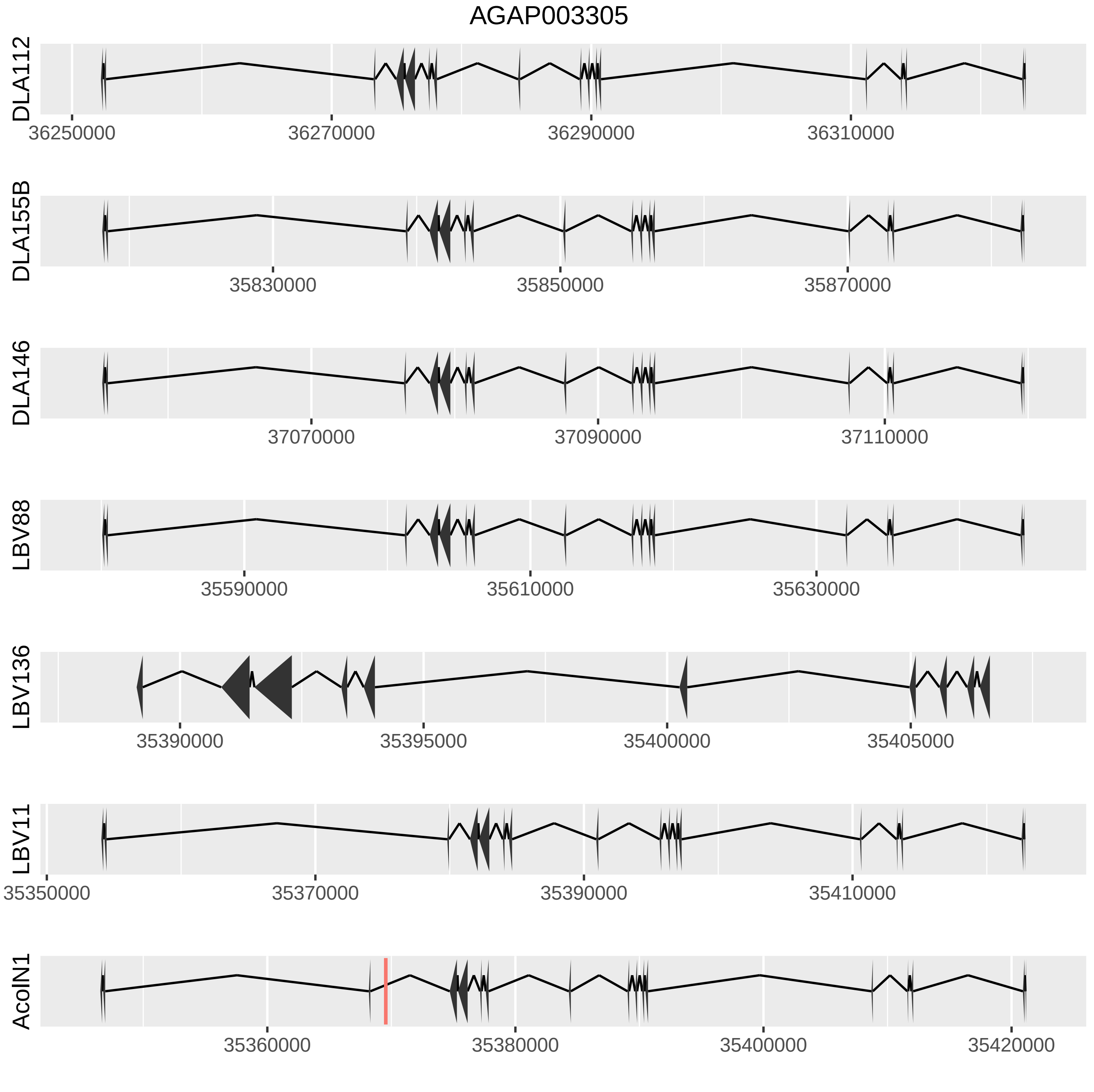

AGAP003305
DLA112
36250000
36270000
36290000
36310000
DLA155B
35830000
35850000
35870000
DLA146
37070000
37090000
37110000
LBV88
35590000
35610000
35630000
LBV136
35390000
35395000
35400000
35405000
LBV11
35350000
35370000
35390000
35410000
AcolN1
35360000
35380000
35400000
35420000

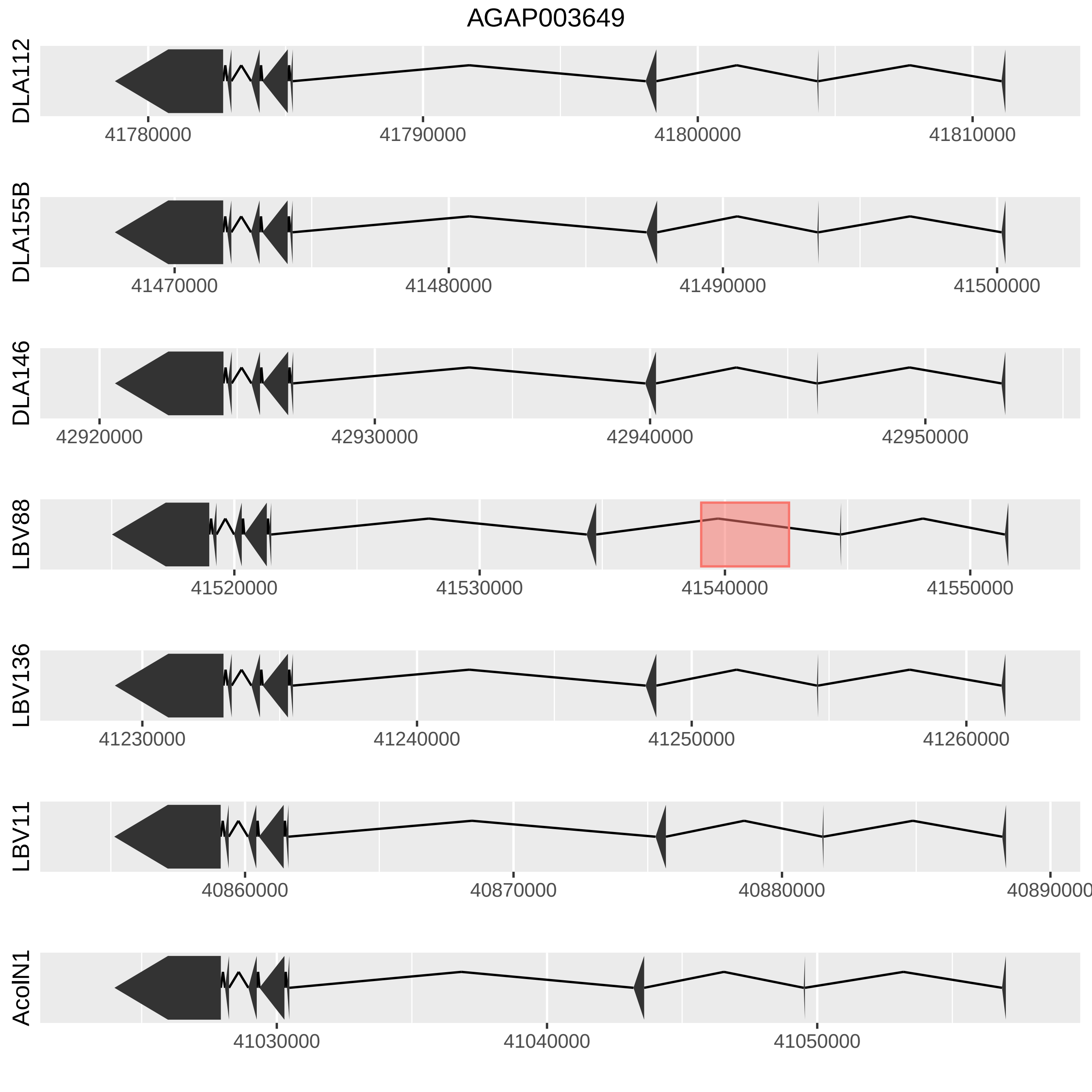

AGAP003649
DLA112
41780000
41790000
41800000
41810000
DLA155B
41470000
41480000
41490000
41500000
DLA146
42920000
42930000
42940000
42950000
LBV88
41520000
41530000
41540000
41550000
LBV136
41230000
41240000
41250000
41260000
LBV11
40860000
40870000
40880000
40890000
AcolN1
41030000
41040000
41050000

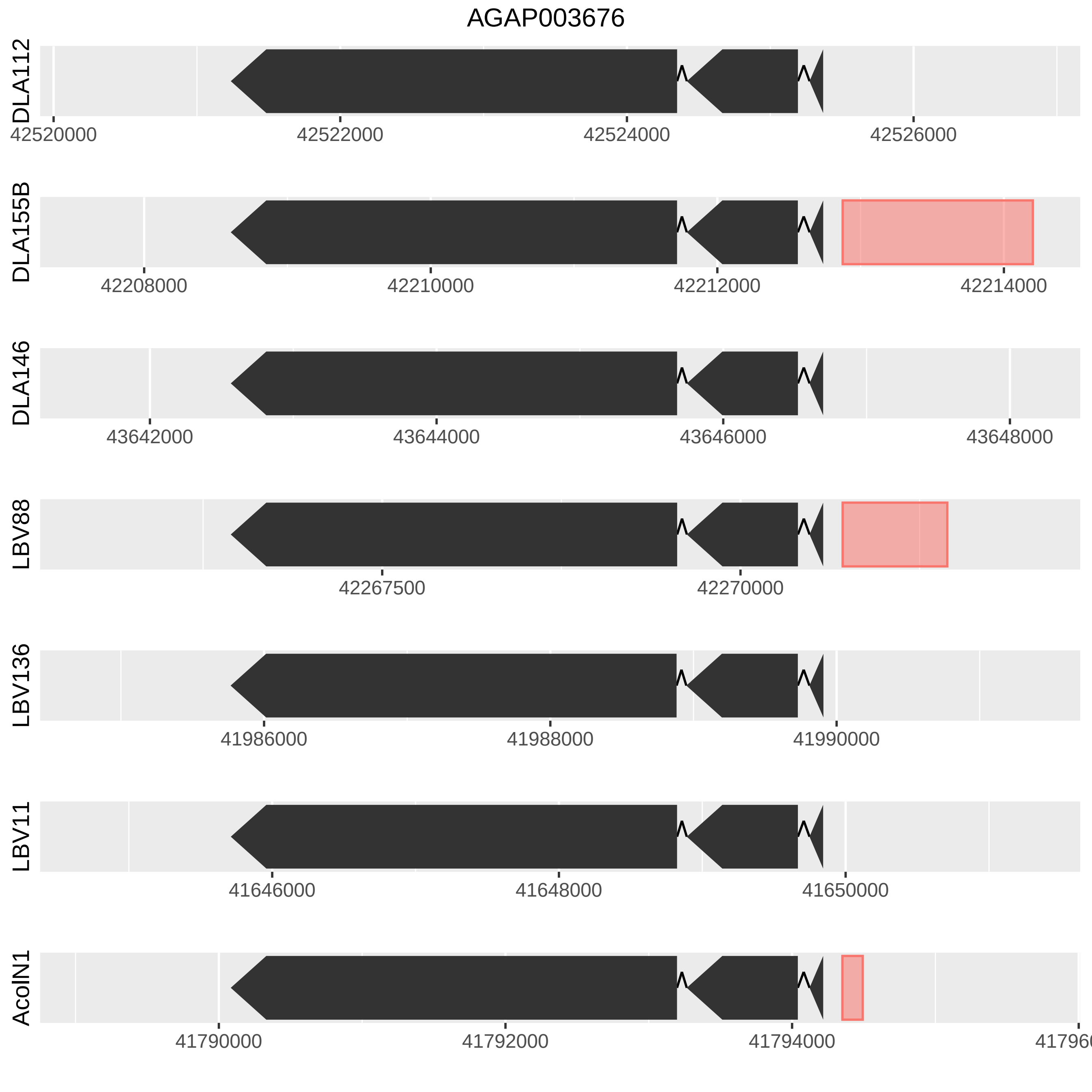
AGAP003676
DLA112
42520000
42522000
42524000
42526000
DLA155B
42208000
42210000
42212000
42214000
DLA146
43642000
43644000
43646000
43648000
LBV88
42267500
42270000
LBV136
41986000
41988000
41990000
LBV11
41646000
41648000
41650000
AcolN1
41790000
41792000
41794000

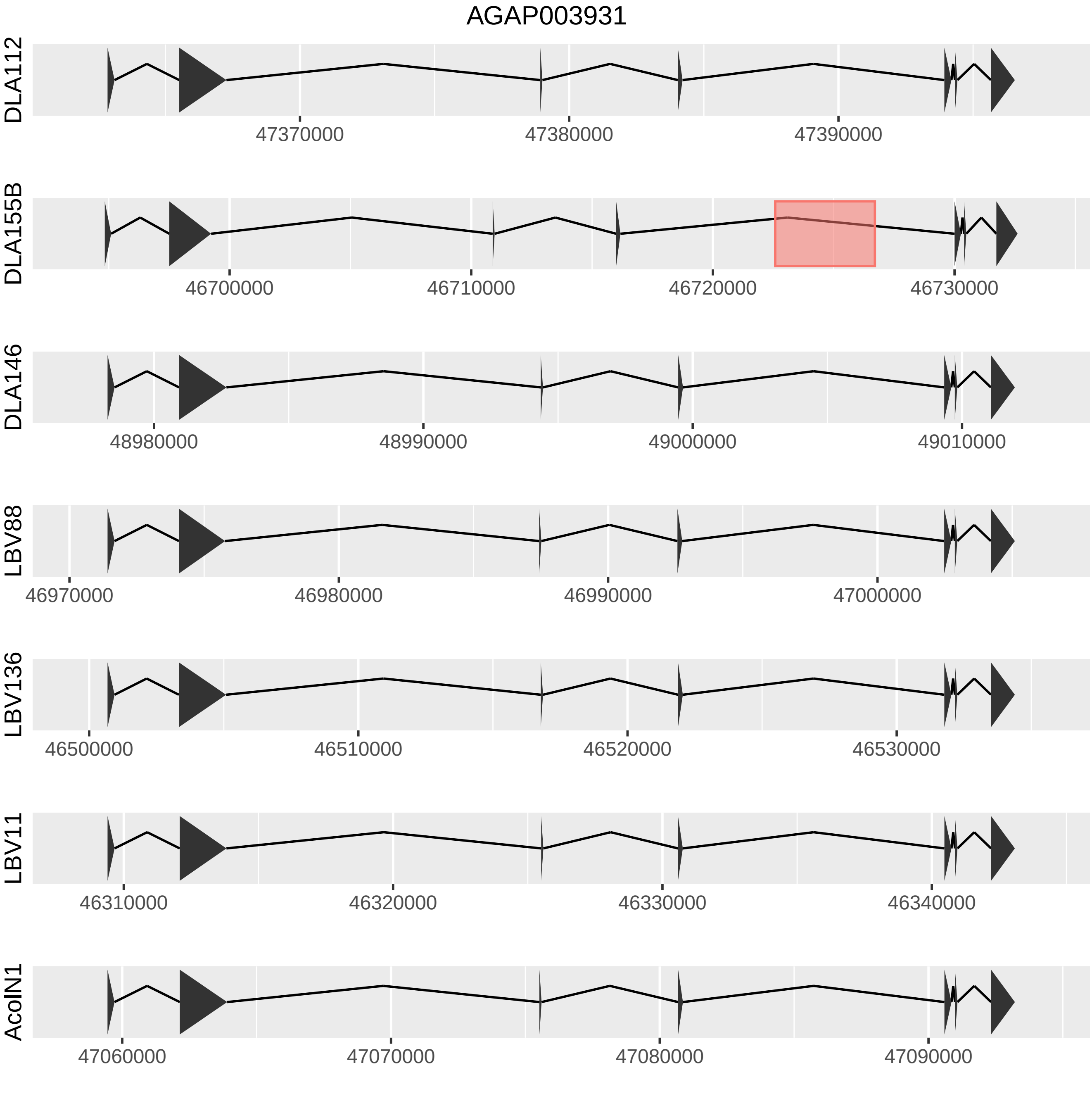
AGAP003931
DLA112
47370000
47380000
47390000
DLA155B
46700000
46710000
46720000
46730000
DLA146
48980000
48990000
49000000
49010000
LBV88
46970000
46980000
46990000
47000000
LBV136
46500000
46510000
46520000
46530000
LBV11
46310000
46320000
46330000
46340000
AcolN1
47060000
47070000
47080000
47090000

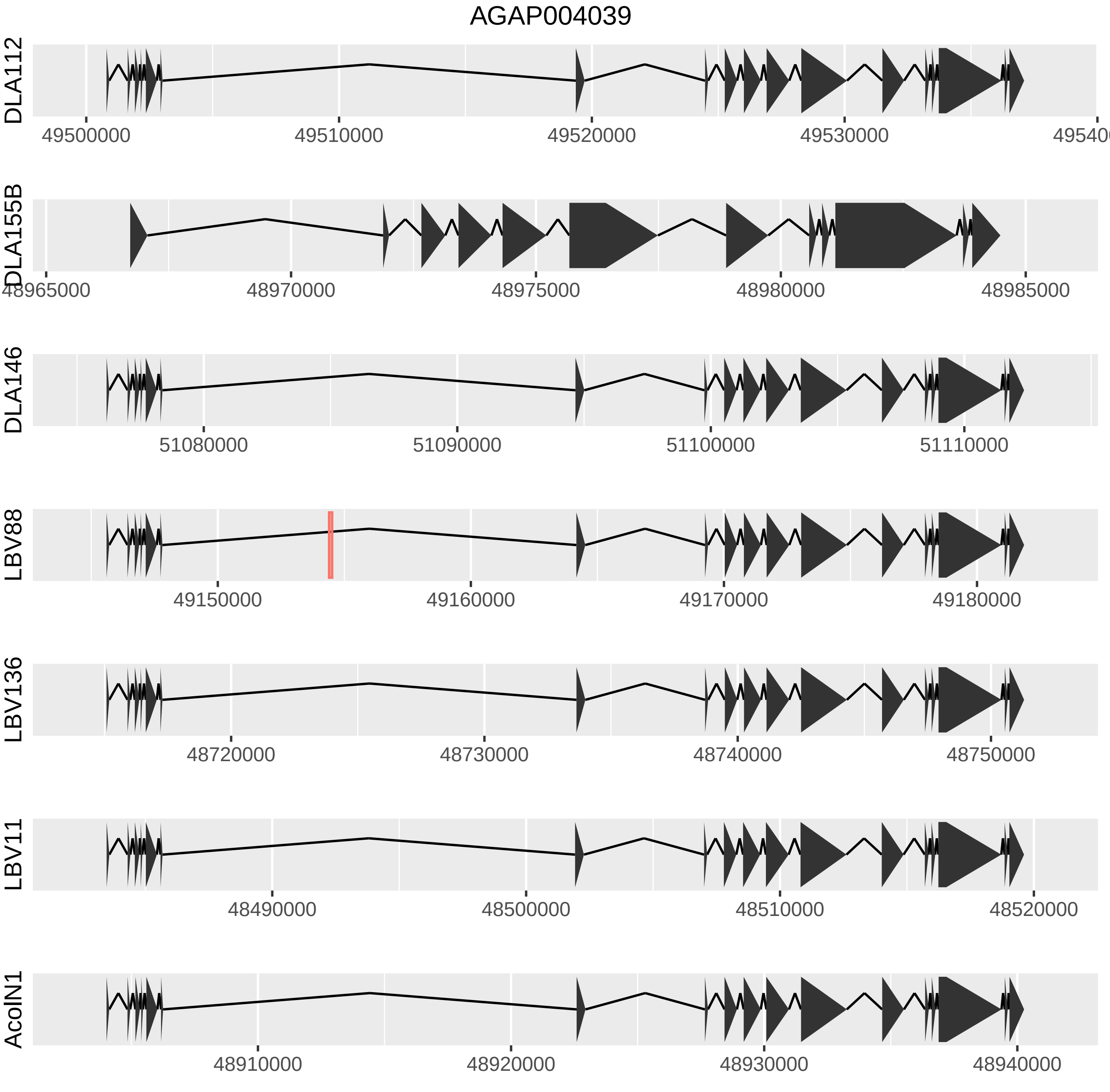

AGAP004039
DLA112
49500000
49510000
49520000
49530000
DLA155B
48965000
48970000
48975000
48980000
48985000
DLA146
51080000
51090000
51100000
51110000
LBV88
49150000
49160000
49170000
49180000
LBV136
48720000
48730000
48740000
48750000
LBV11
48490000
48500000
48510000
48520000
AcolN1
48910000
48920000
48930000
48940000

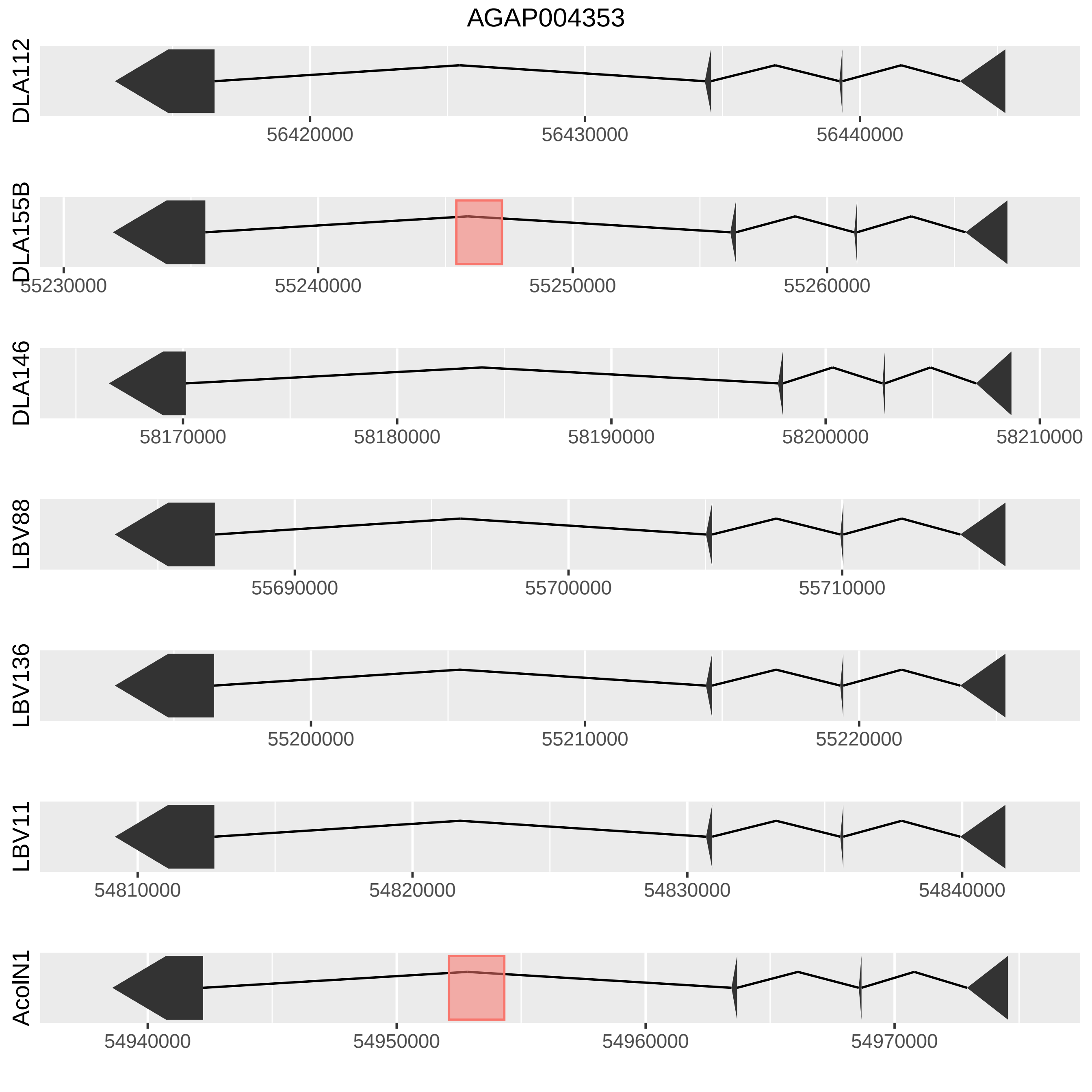

AGAP004353
DLA112
56420000
56430000
56440000
DLA155B
55230000
55240000
55250000
55260000
DLA146
58170000
58180000
58190000
58200000
58210000
LBV88
55690000
55700000
55710000
LBV136
55200000
55210000
55220000
LBV11
54810000
54820000
54830000
54840000
AcolN1
54940000
54950000
54960000
54970000

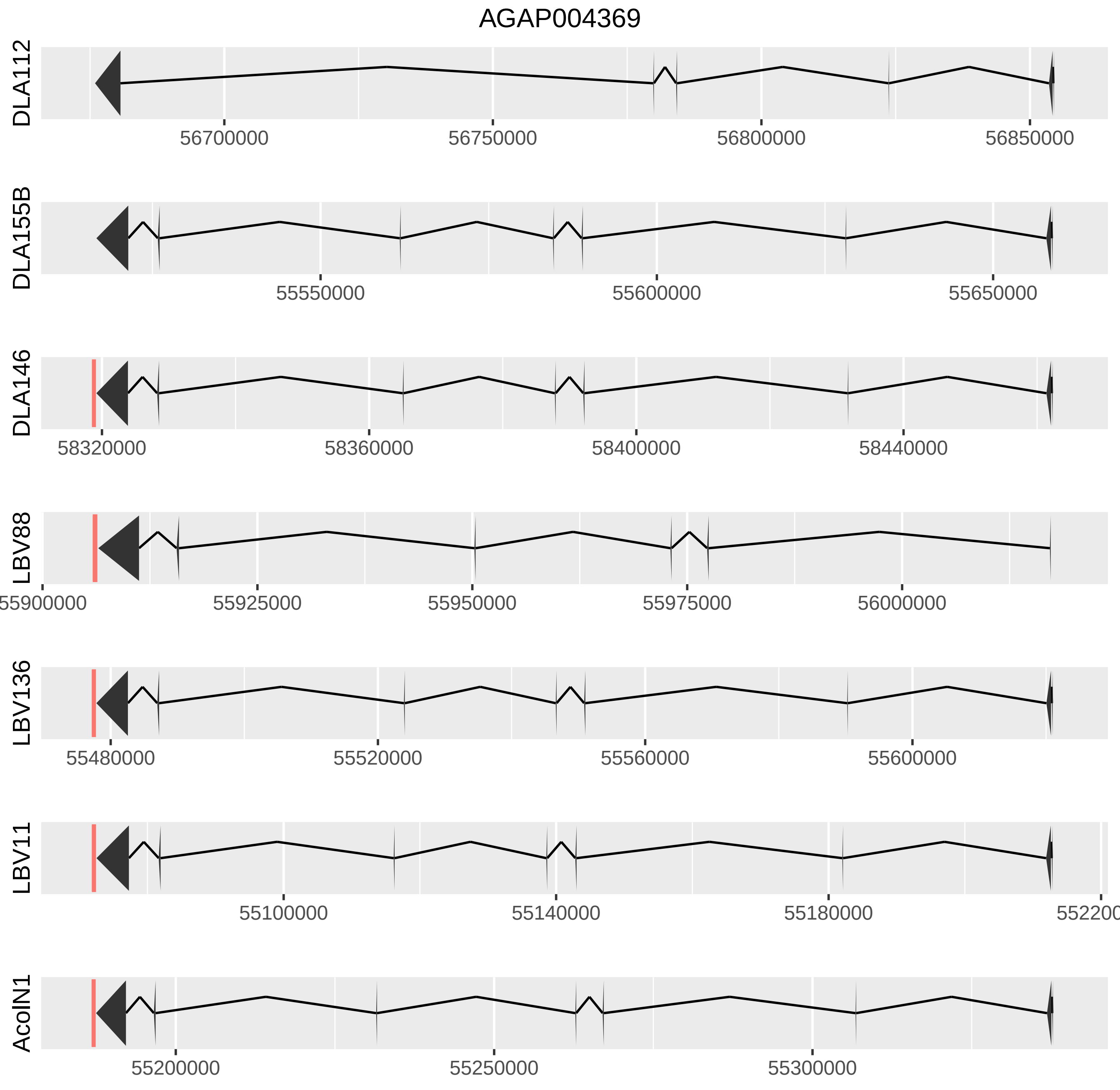

AGAP004369
DLA112
56700000
56750000
56800000
56850000
DLA155B
55550000
55600000
55650000
DLA146
58320000
58360000
58400000
58440000
LBV88
55900000
55925000
55950000
55975000
56000000
LBV136
55480000
55520000
55560000
55600000
LBV11
55100000
55140000
55180000
AcoIN1
55200000
55250000
55300000

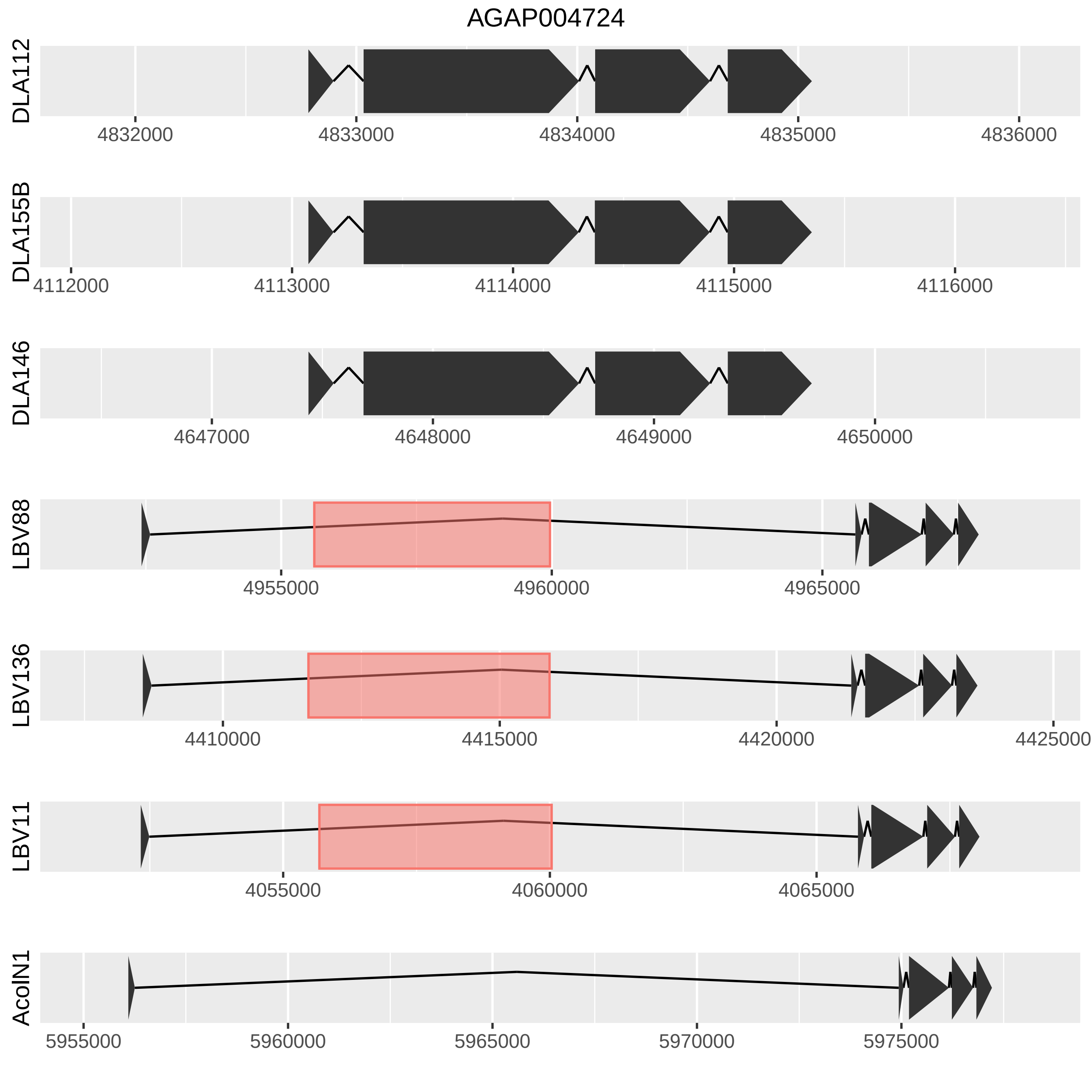
AGAP004724
DLA112
4832000
4833000
4834000
4835000
4836000
DLA155B
4112000
4113000
4114000
4115000
4116000
DLA146
4647000
4648000
4649000
4650000
LBV88
4955000
4960000
4965000
LBV136
4410000
4415000
4420000
4425000
LBV11
4055000
4060000
4065000
AcolN1
5955000
5960000
5965000
5970000
5975000

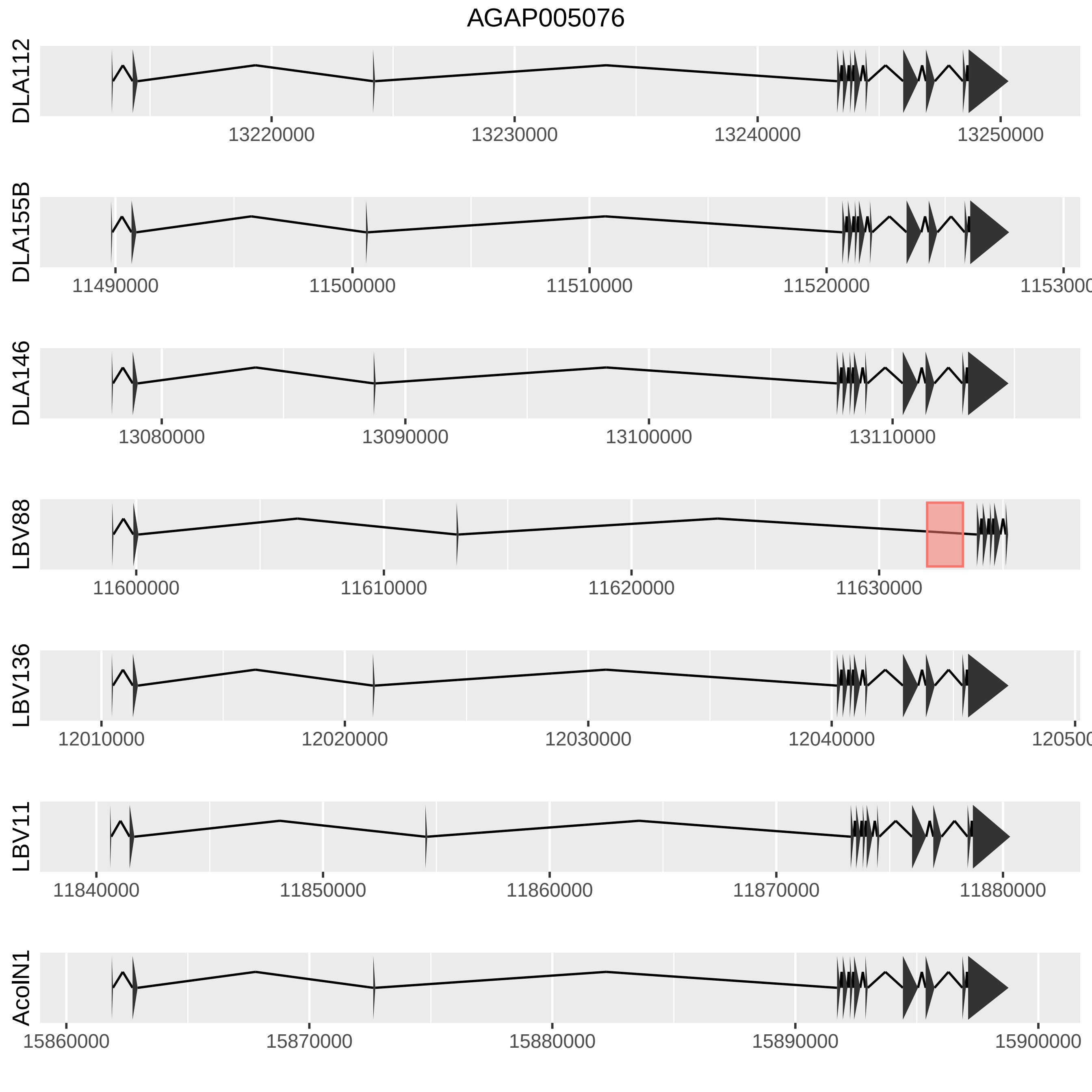

AGAP005076
DLA112
13220000
13230000
13240000
13250000
DLA155B
11490000
11500000
11510000
11520000
DLA146
13080000
13090000
13100000
13110000
LBV88
11600000
11610000
11620000
11630000
LBV136
12010000
12020000
12030000
12040000
LBV11
11840000
11850000
11860000
11870000
11880000
AcolN1
15860000
15870000
15880000
15890000
15900000

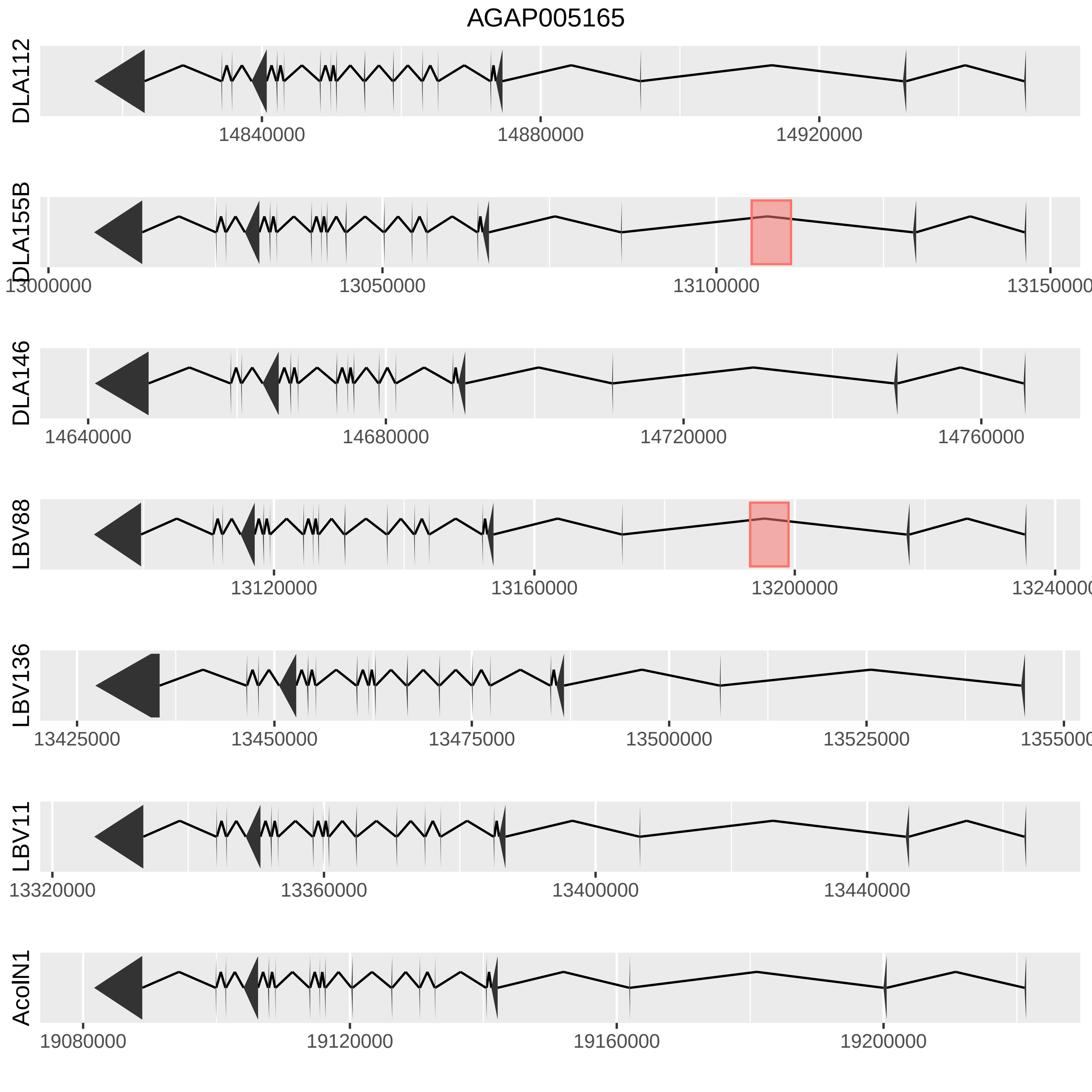

AGAP005165
DLA112
14840000
14880000
14920000
DLA155B
13000000
13050000
13100000
13150000
DLA146
14640000
14680000
14720000
14760000
LBV88
13120000
13160000
13200000
13240000
LBV136
13425000
13450000
13475000
13500000
13525000
13550000
LBV11
13320000
13360000
13400000
13440000
AcolN1
19080000
19120000
19160000
19200000

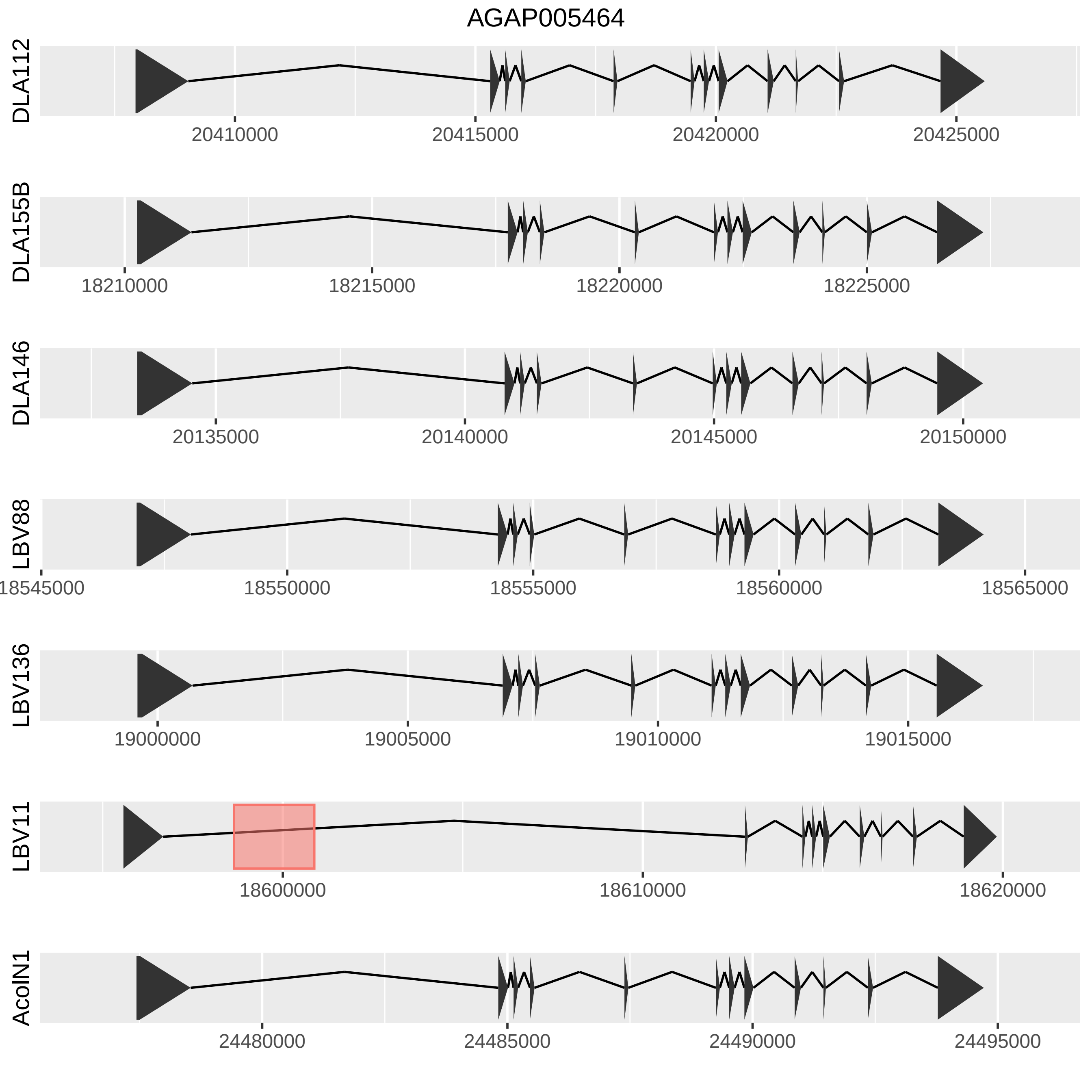

AGAP005464
DLA112
20410000
20415000
20420000
20425000
DLA155B
18210000
18215000
18220000
18225000
DLA146
20135000
20140000
20145000
20150000
LBV88
18545000
18550000
18555000
18560000
18565000
LBV136
19000000
19005000
19010000
19015000
LBV11
18600000
18610000
18620000
AcoIN1
24480000
24485000
24490000
24495000

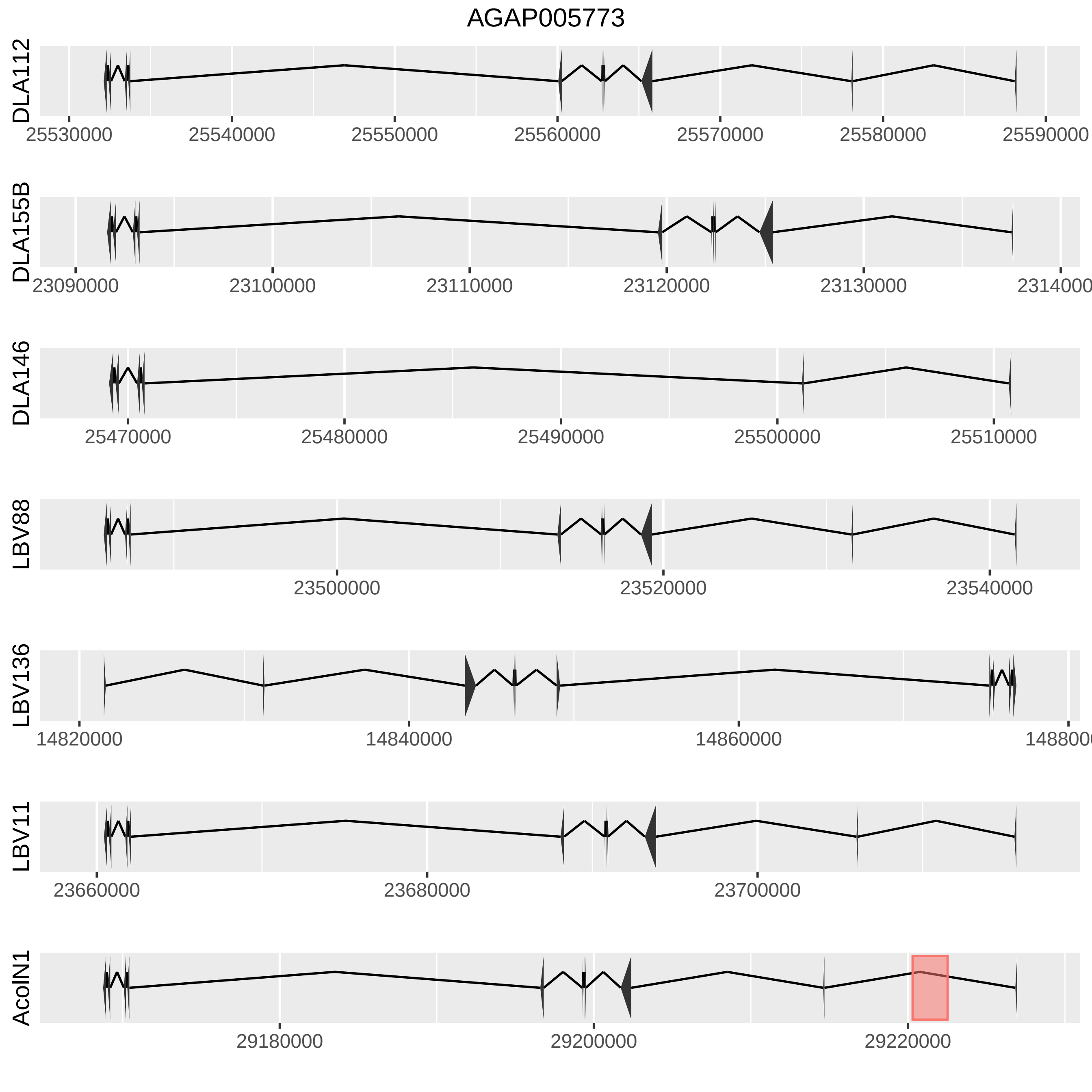
AGAP005773
DLA112
25530000
25540000
25550000
25560000
25570000
25580000
25590000
DLA155B
23090000
23100000
23110000
23120000
23130000
2314000
DLA146
25470000
25480000
25490000
25500000
25510000
LBV88
23500000
23520000
23540000
LBV136
14820000
14840000
14860000
148800
LBV11
23660000
23680000
23700000
AcolN1
29180000
29200000
29220000

AGAP005806

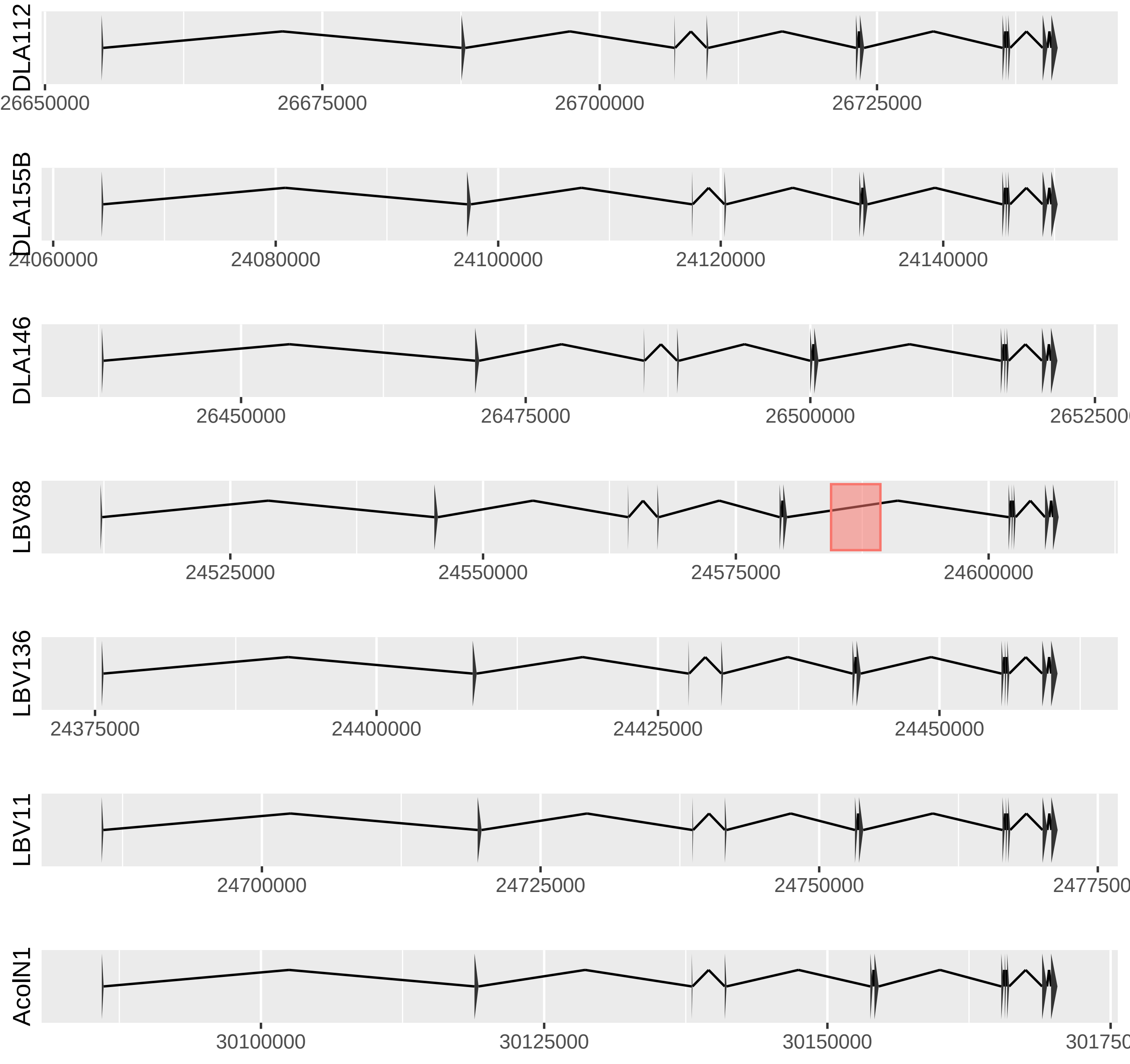

DLA112
26650000
26675000
26700000
26725000
DLA155B
24060000
24080000
24100000
24120000
24140000
DLA146
26450000
26475000
26500000
LBV88
24525000
24550000
24575000
24600000
LBV136
24375000
24400000
24425000
24450000
LBV11
24700000
24725000
24750000
AcolN1
30100000
30125000
30150000

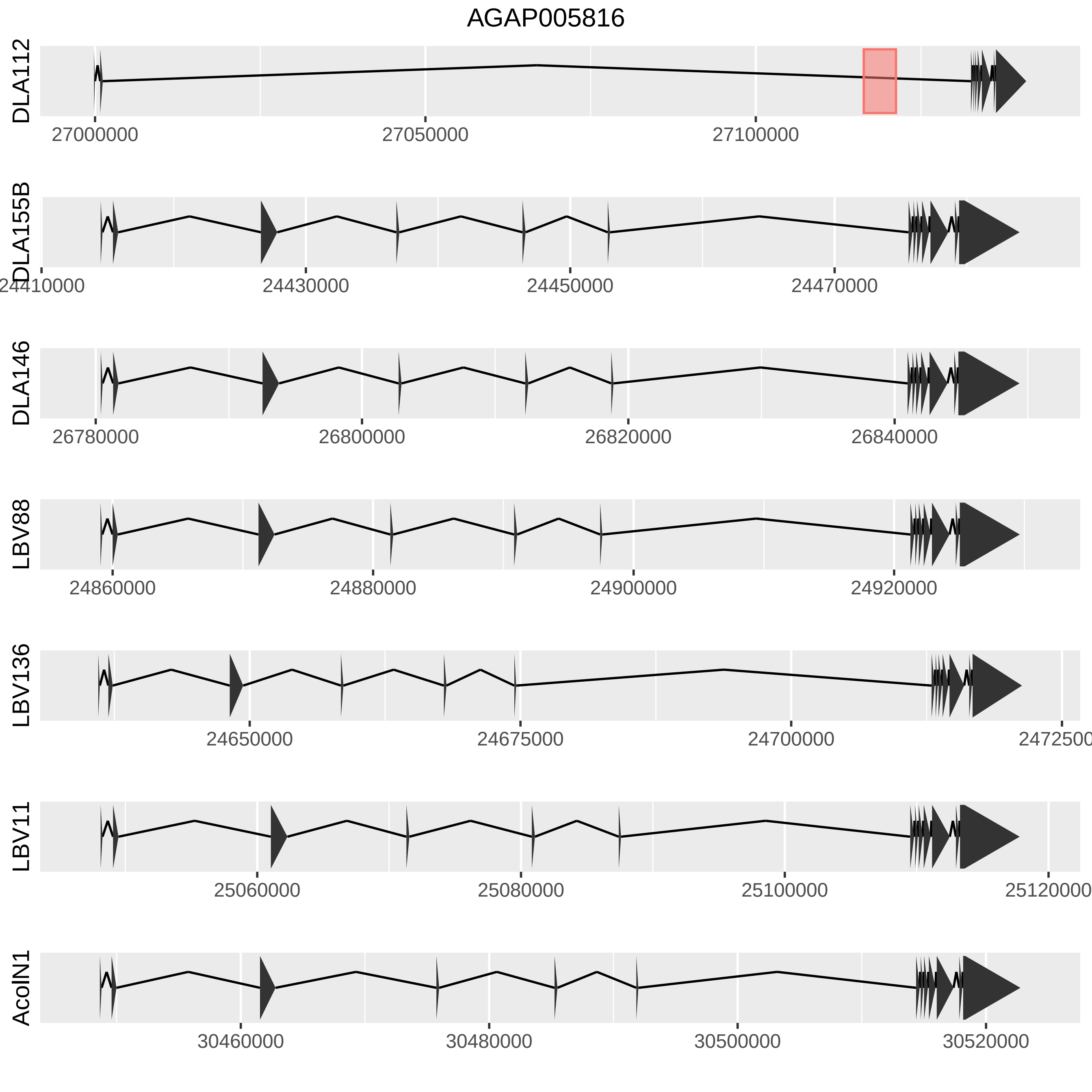
AGAP005816
DLA112
27000000
27050000
27100000
DLA155B
24410000
24430000
24450000
24470000
DLA146
26780000
26800000
26820000
26840000
LBV88
24860000
24880000
24900000
24920000
LBV136
24650000
24675000
24700000
LBV11
25060000
25080000
25100000
AcolN1
30460000
30480000
30500000
30520000

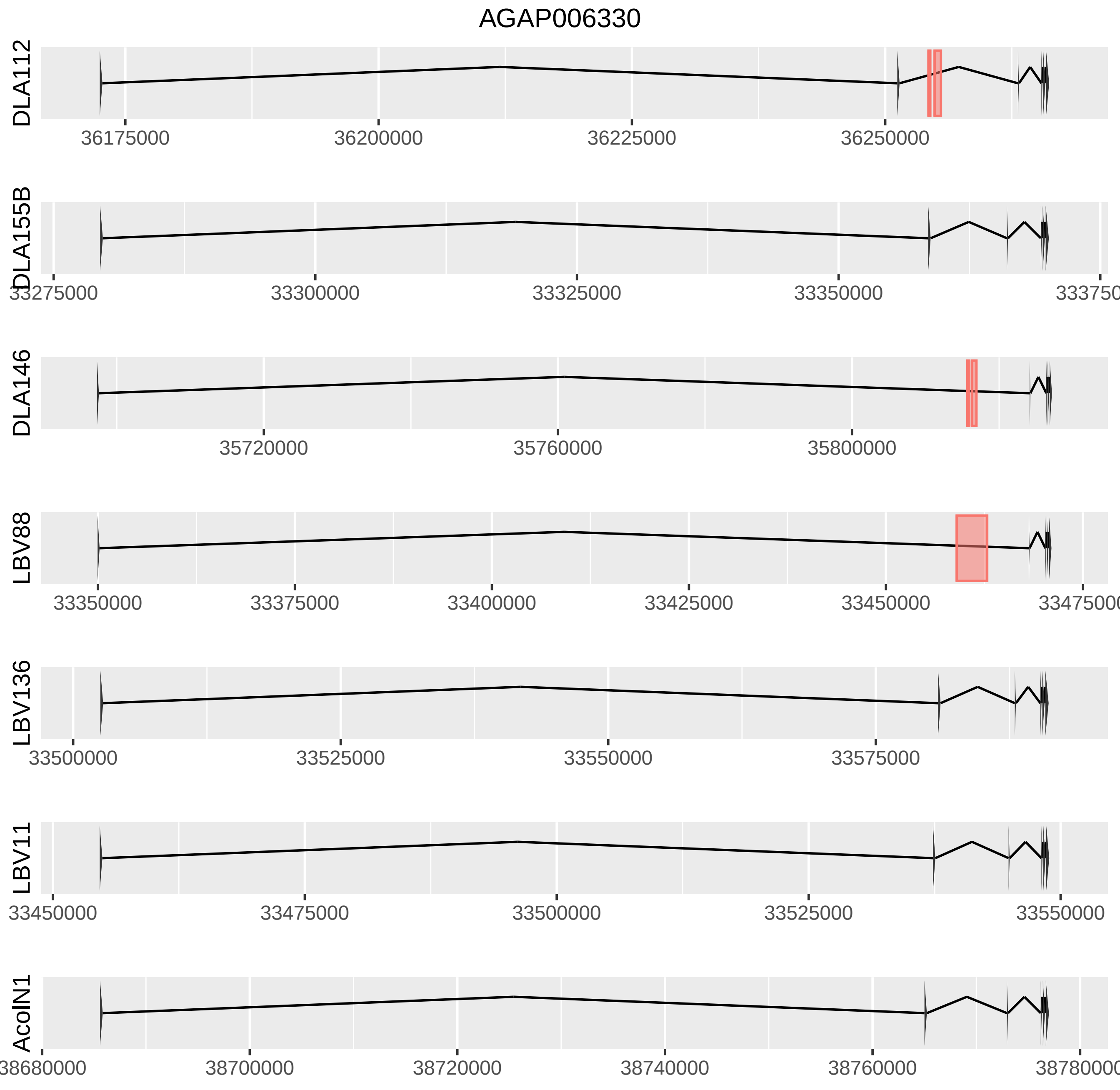
AGAP006330
DLA112
36175000
36200000
36225000
36250000
DLA155B
33275000
33300000
33325000
33350000
DLA146
35720000
35760000
35800000
LBV88
33350000
33375000
33400000
33425000
33450000
LBV136
33500000
33525000
33550000
33575000
LBV11
33450000
33475000
33500000
33525000
33550000
AcolN1
38680000
38700000
38720000
38740000
38760000
38780000

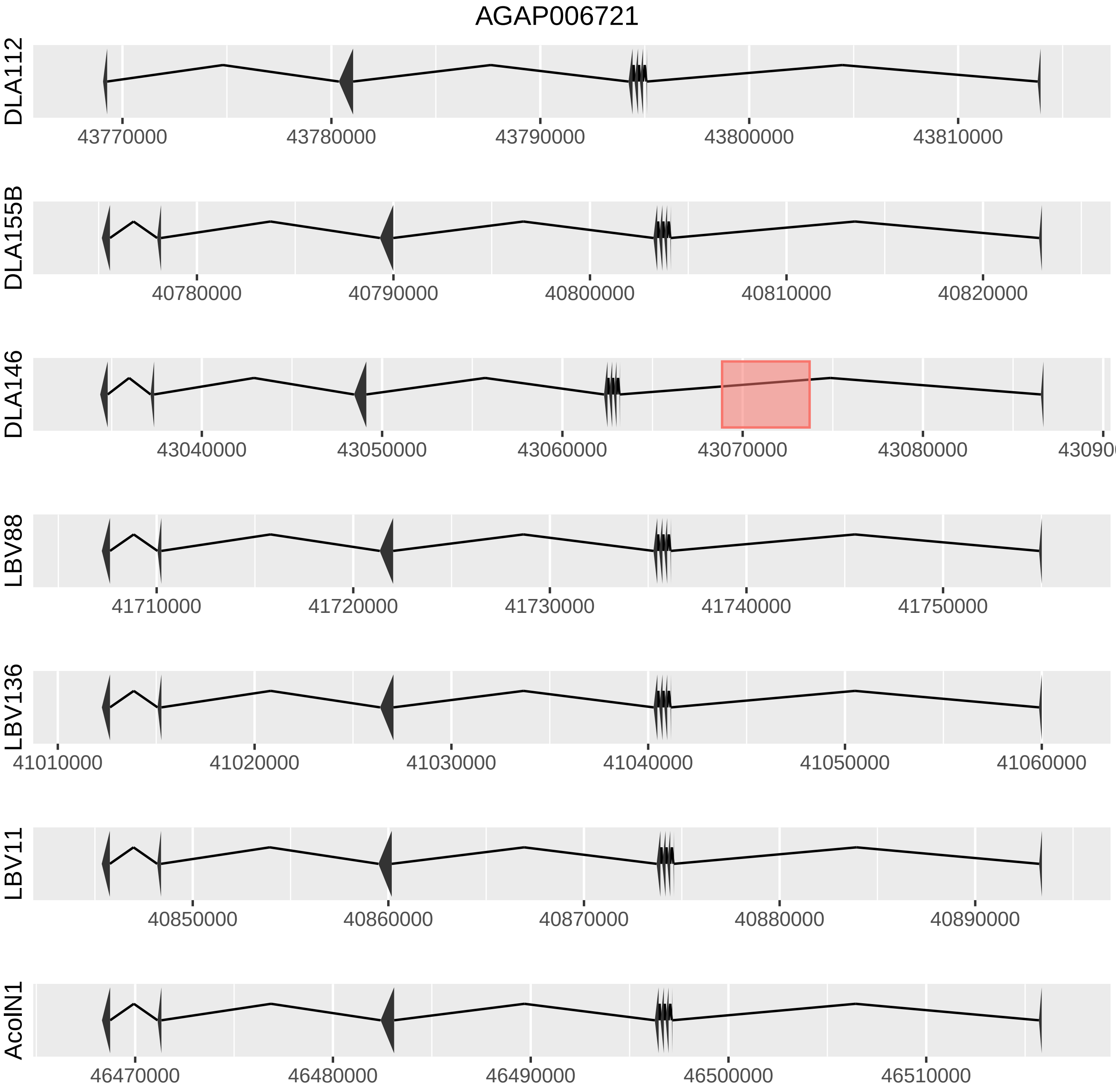
AGAP006721
DLA112
43770000
43780000
43790000
43800000
43810000
DLA155B
40780000
40790000
40800000
40810000
40820000
DLA146
43040000
43050000
43060000
43070000
43080000
LBV88
41710000
41720000
41730000
41740000
41750000
LBV136
41010000
41020000
41030000
41040000
41050000
41060000
LBV11
40850000
40860000
40870000
40880000
40890000
AcolN1
46470000
46480000
46490000
46500000
46510000

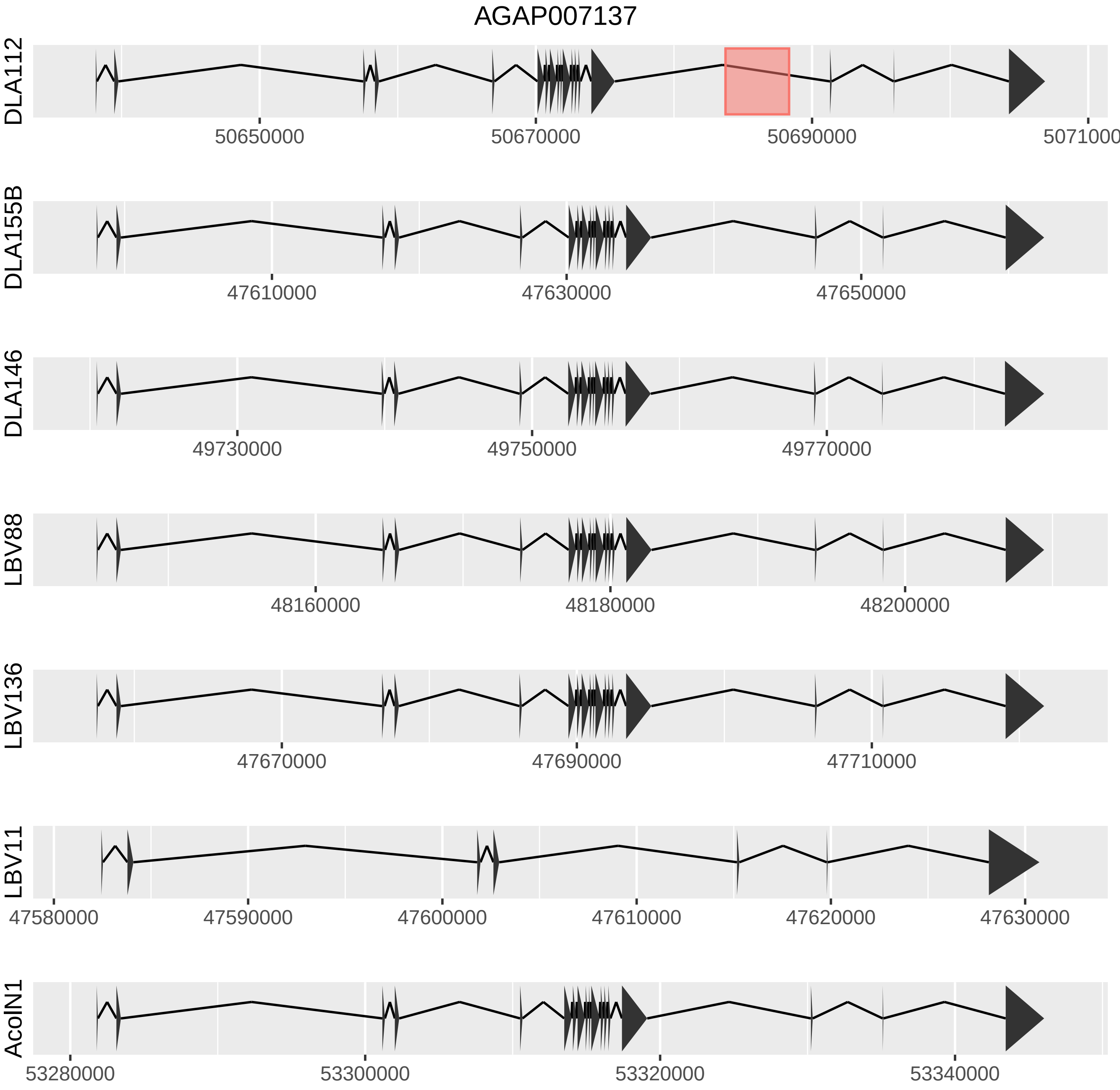
AGAP007137
DLA112
50650000
50670000
50690000
5071000
DLA155B
47610000
47630000
47650000
DLA146
49730000
49750000
49770000
LBV88
48160000
48180000
48200000
LBV136
47670000
47690000
47710000
LBV11
47580000
47590000
47600000
47610000
47620000
47630000
AcolN1
53280000
53300000
53320000
53340000

AGAP007760

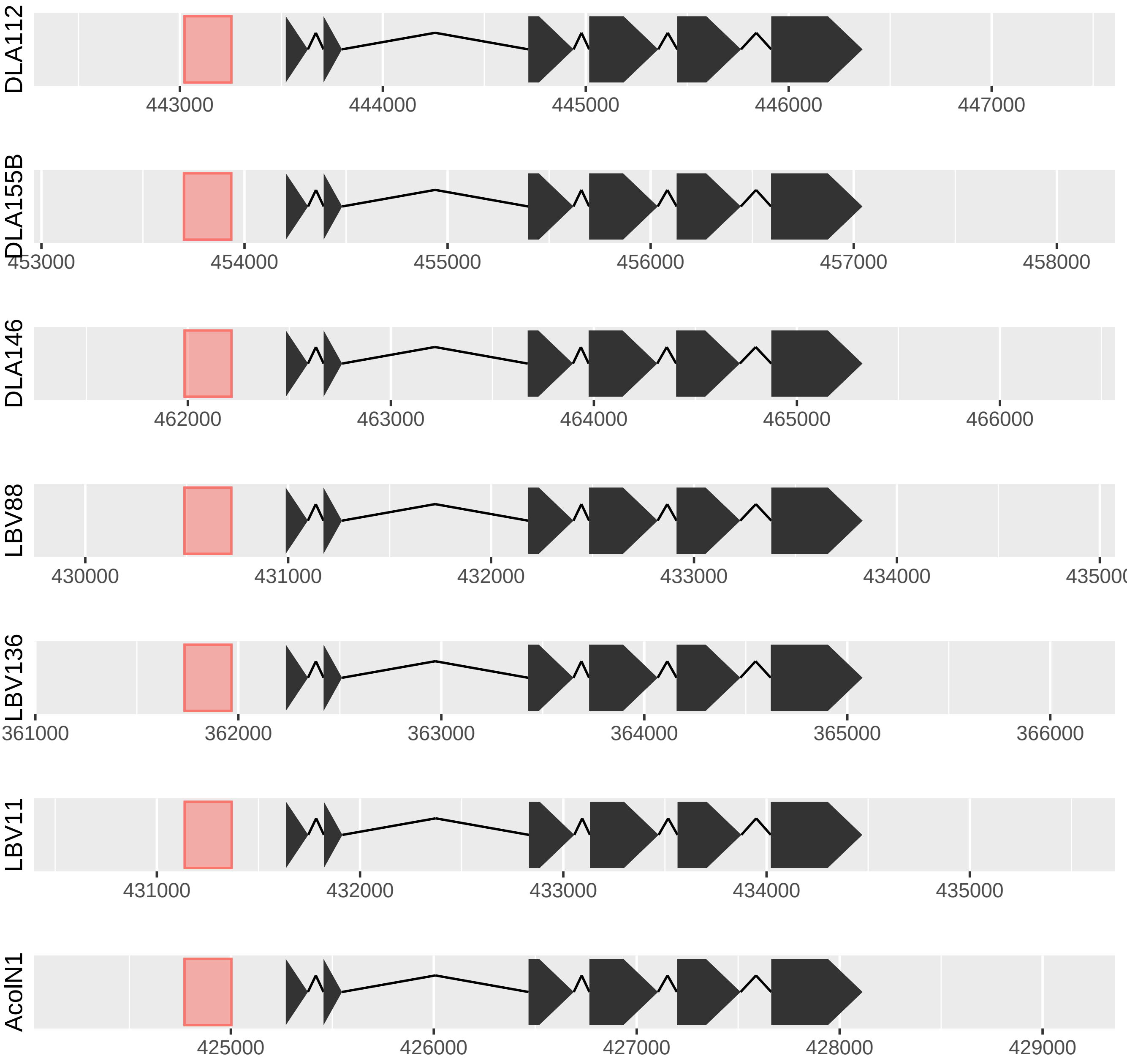

DLA112
443000
444000
445000
446000
447000
DLA155B
453000
454000
455000
456000
457000
458000
DLA146
462000
463000
464000
465000
466000
LBV88
430000
431000
432000
433000
434000
435000
LBV136
361000
362000
363000
364000
365000
366000
LBV11
431000
432000
433000
434000
435000
AcolN1
425000
426000
427000
428000
429000

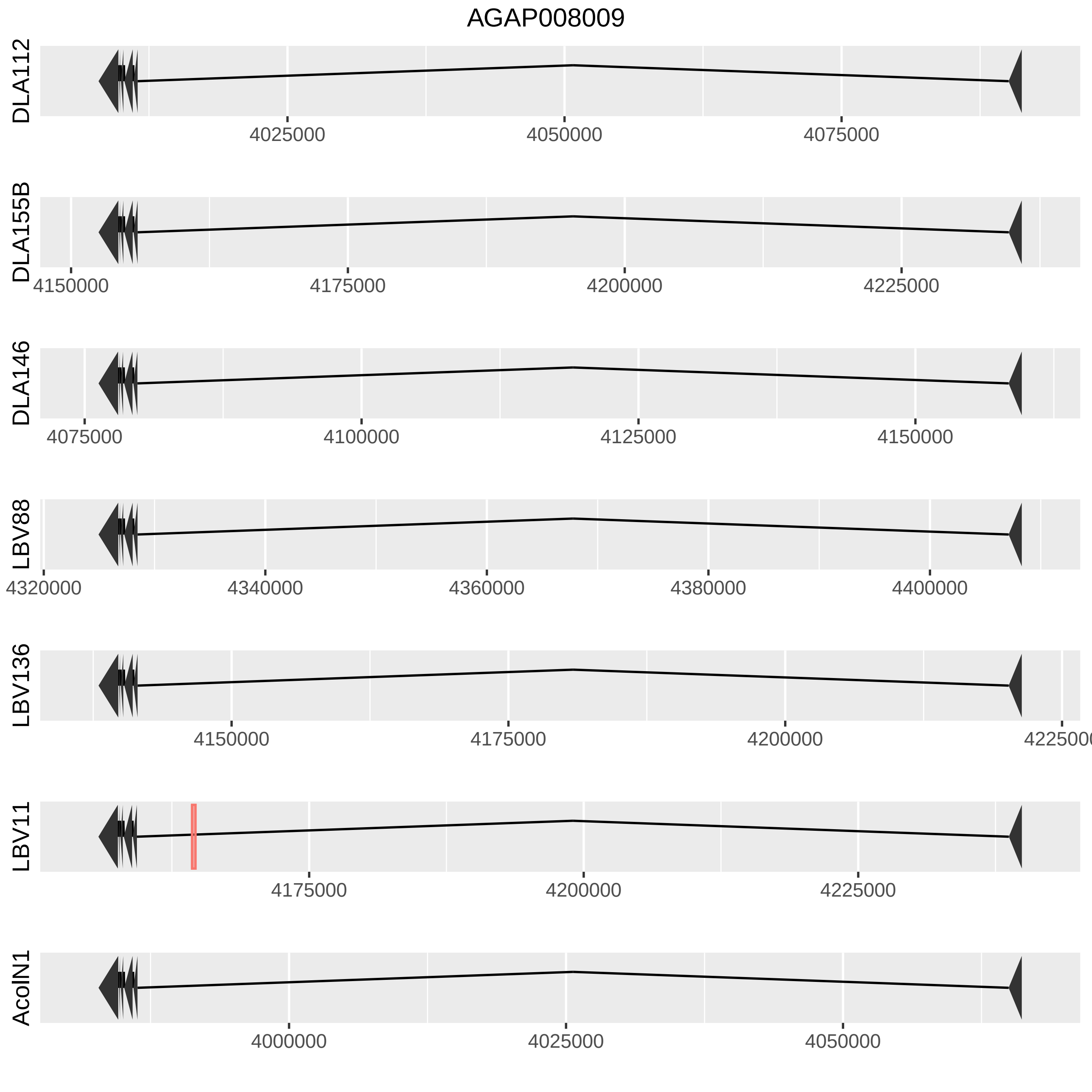
AGAP008009
DLA112
4025000
4050000
4075000
DLA155B
4150000
4175000
4200000
4225000
DLA146
4075000
4100000
4125000
4150000
LBV88
4320000
4340000
4360000
4380000
4400000
LBV136
4150000
4175000
4200000
4225000
LBV11
4175000
4200000
4225000
AcolN1
4000000
4025000
4050000

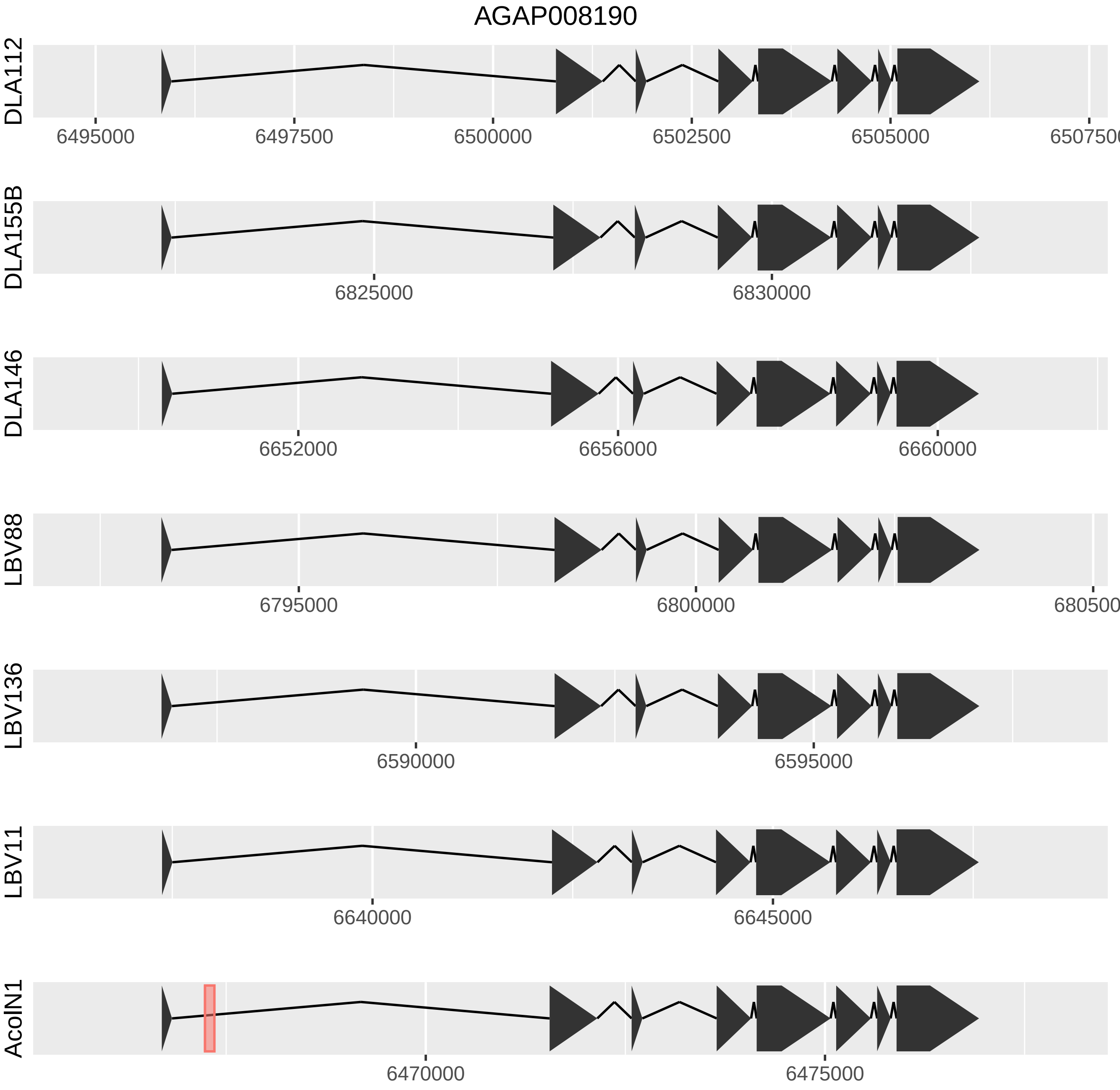

AGAP008190
DLA112
6495000
6497500
6500000
6502500
6505000
6507500
DLA155B
6825000
6830000
DLA146
6652000
6656000
6660000
LBV88
6795000
6800000
680500
LBV136
6590000
6595000
LBV11
6640000
6645000
AcoIN1
6470000
6475000

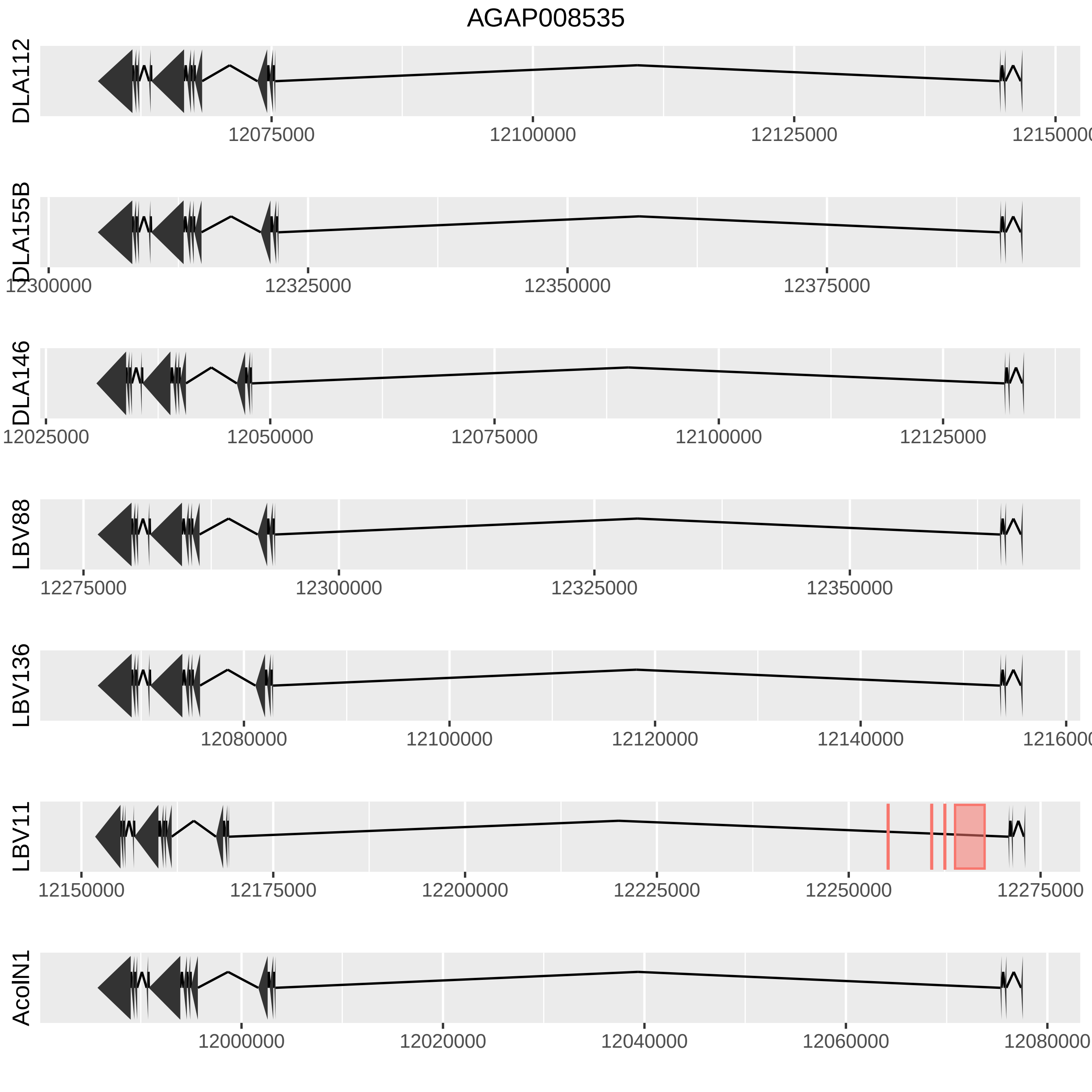
AGAP008535
DLA112
12075000
12100000
12125000
1215000
DLA155B
12300000
12325000
12350000
12375000
DLA146
12025000
12050000
12075000
12100000
12125000
LBV88
12275000
12300000
12325000
12350000
LBV136
12080000
12100000
12120000
12140000
1216000
LBV11
12150000
12175000
12200000
12225000
12250000
12275000
AcolN1
12000000
12020000
12040000
12060000
12080000

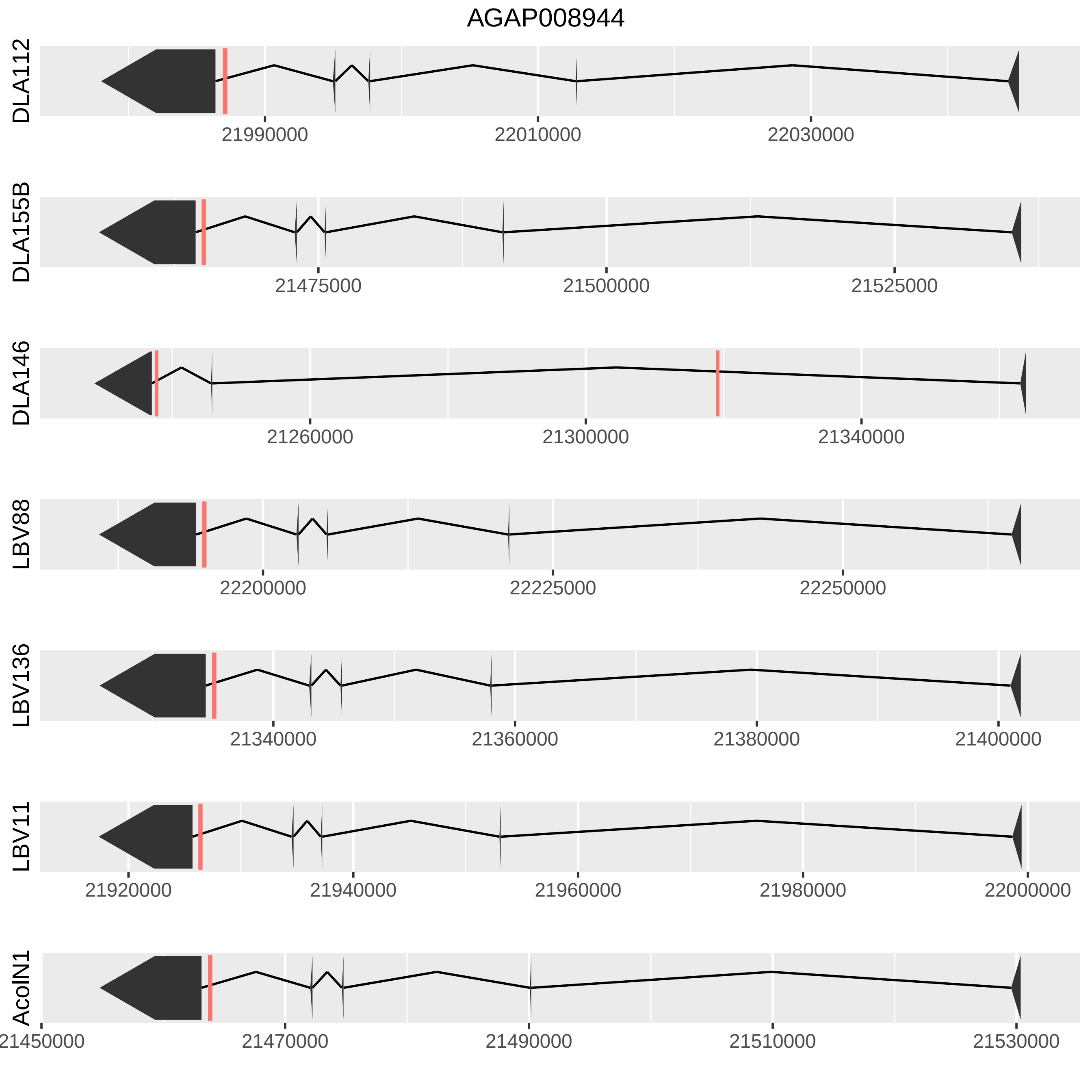
AGAP008944
DLA112
21990000
22010000
22030000
DLA155B
21475000
21500000
21525000
DLA146
21260000
21300000
21340000
LBV88
22200000
22225000
22250000
LBV136
21340000
21360000
21380000
21400000
LBV11
21920000
21940000
21960000
21980000
22000000
AcoIN1
21450000
21470000
21490000
21510000
21530000

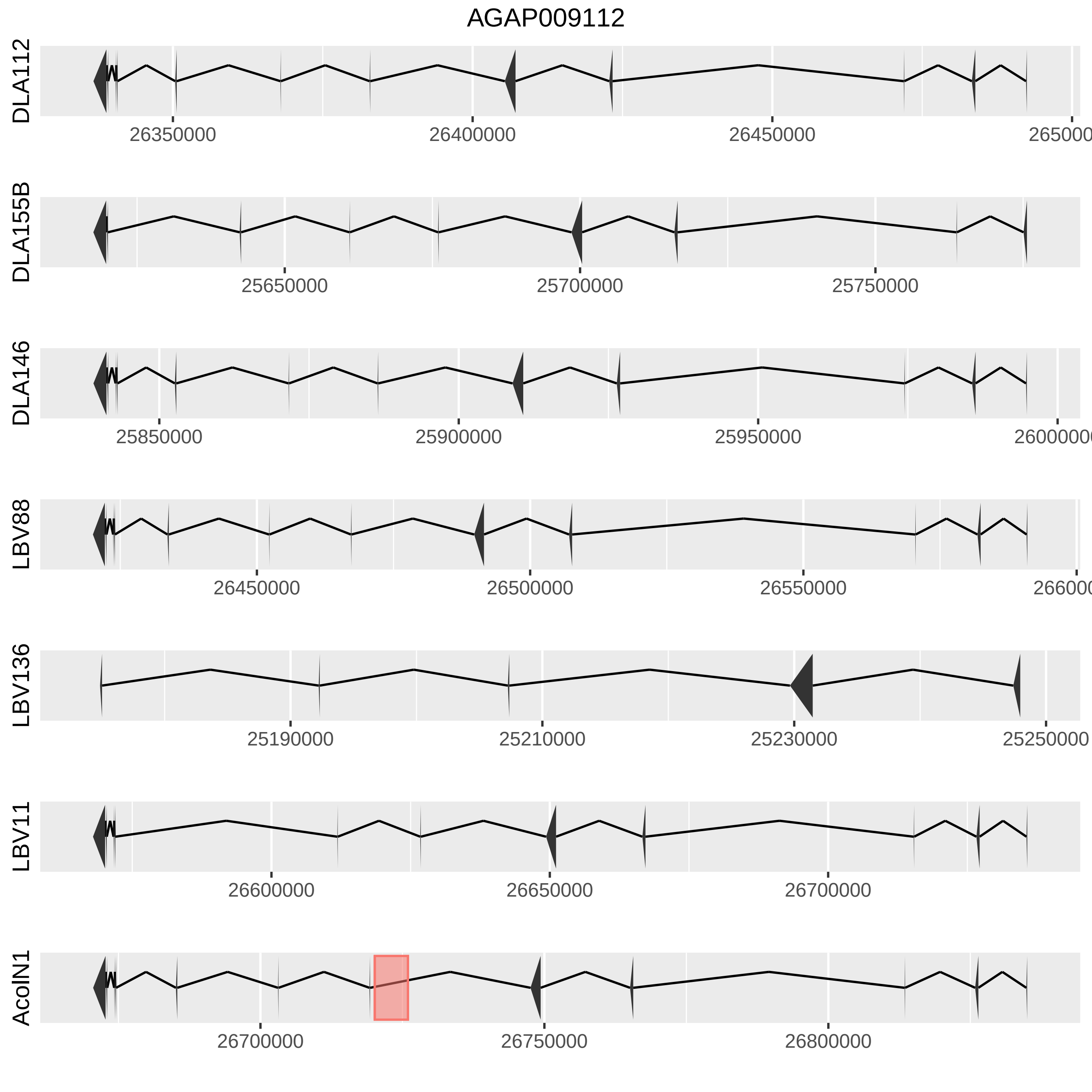
AGAP009112
DLA112
26350000
26400000
26450000
DLA155B
25650000
25700000
25750000
DLA146
25850000
25900000
25950000
LBV88
26450000
26500000
26550000
LBV136
25190000
25210000
25230000
25250000
LBV11
26600000
26650000
26700000
AcolN1
26700000
26750000
26800000

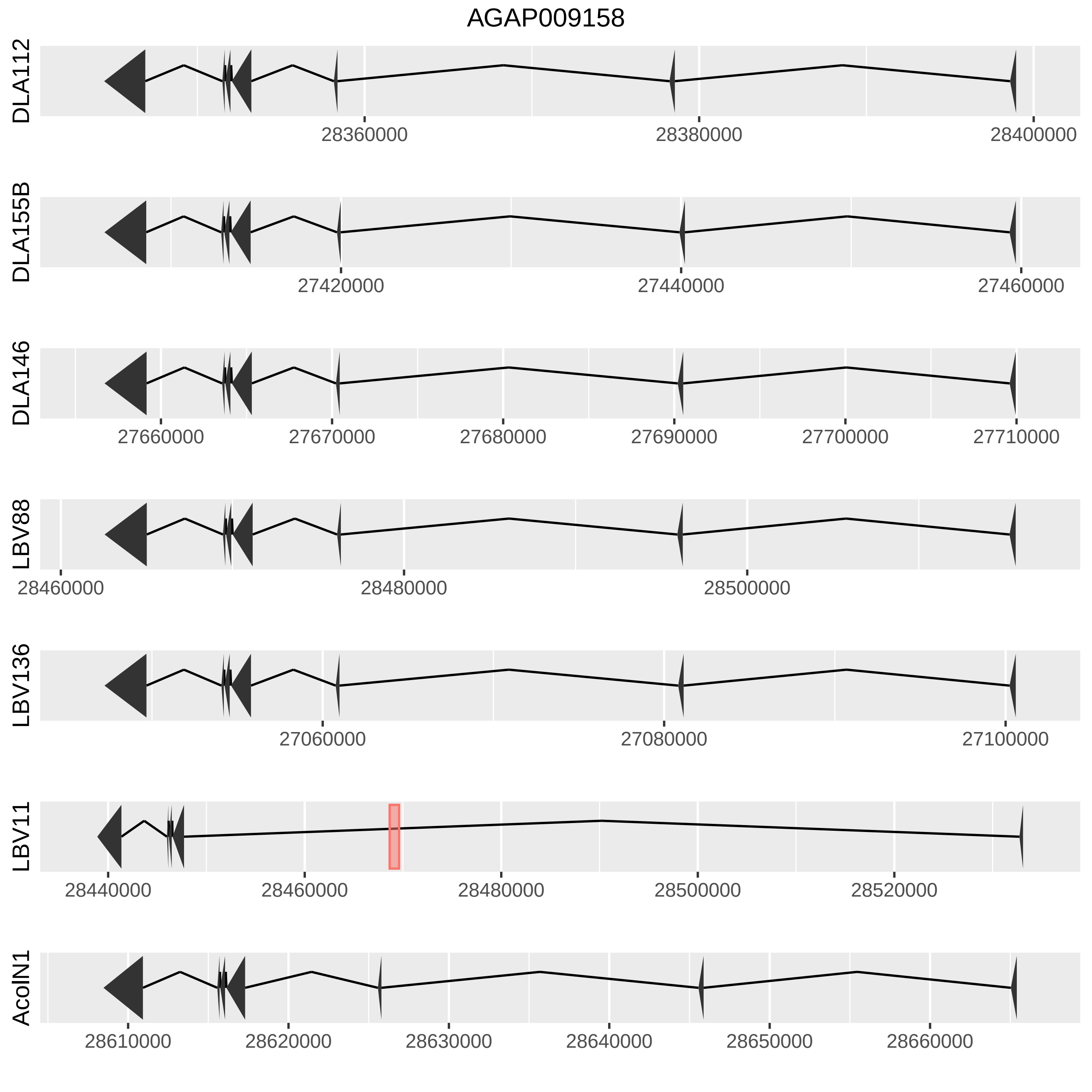

AGAP009158
DLA112
28360000
28380000
28400000
DLA155B
27420000
27440000
27460000
DLA146
27660000
27670000
27680000
27690000
27700000
27710000
LBV88
28460000
28480000
28500000
LBV136
27060000
27080000
27100000
LBV11
28440000
28460000
28480000
28500000
28520000
AcolN1
28610000
28620000
28630000
28640000
28650000
28660000

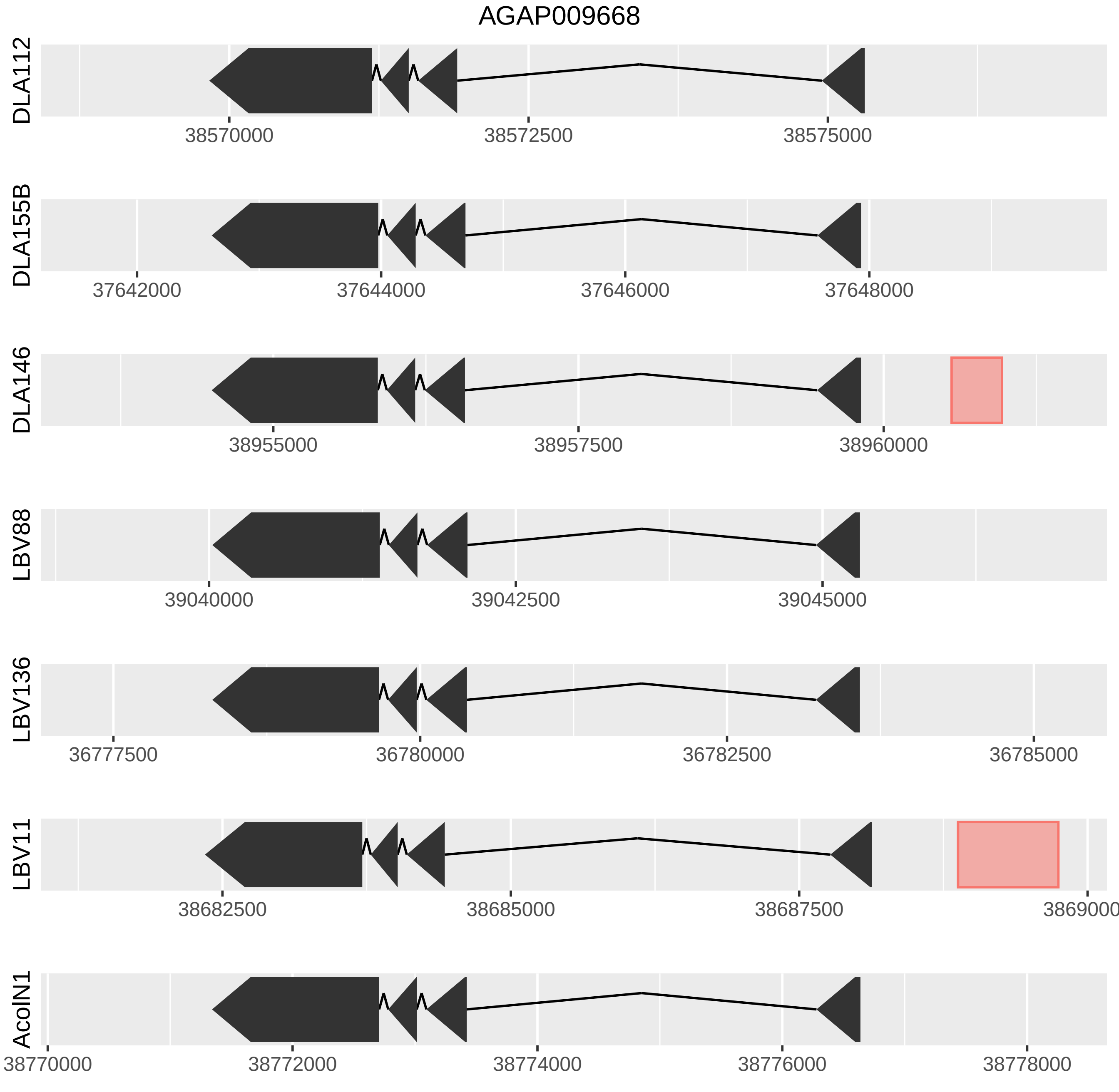

AGAP009668
DLA112
38570000
38572500
38575000
DLA155B
37642000
37644000
37646000
37648000
DLA146
38955000
38957500
38960000
LBV88
39040000
39042500
39045000
LBV136
36777500
36780000
36782500
36785000
LBV11
38682500
38685000
38687500
3869000
AcoIN1
38770000
38772000
38774000
38776000
38778000

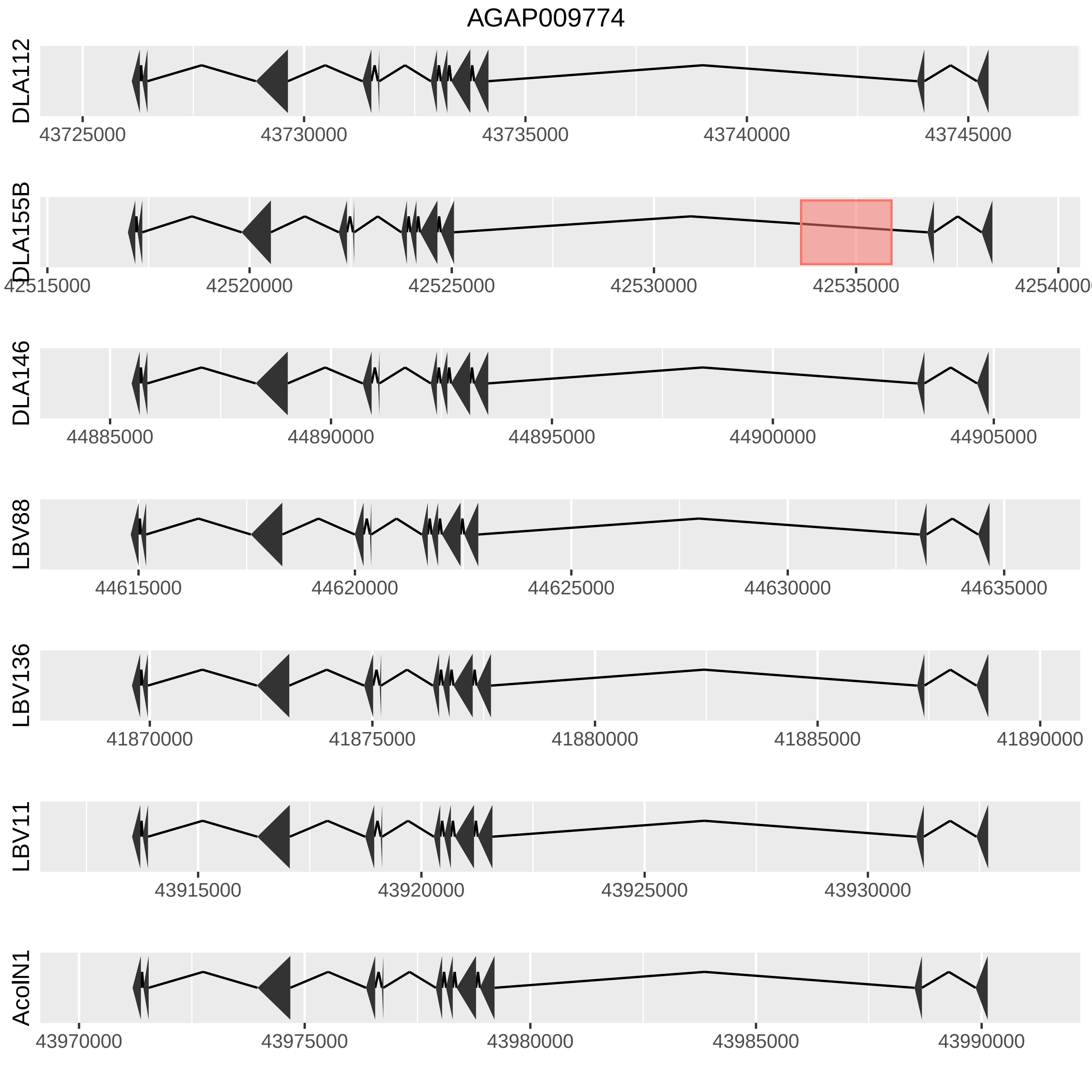
AGAP009774
DLA112
43725000
43730000
43735000
43740000
43745000
DLA155B
42515000
42520000
42525000
42530000
42535000
42540000
DLA146
44885000
44890000
44895000
44900000
44905000
LBV88
44615000
44620000
44625000
44630000
44635000
LBV136
41870000
41875000
41880000
41885000
41890000
LBV11
43915000
43920000
43925000
43930000
AcolN1
43970000
43975000
43980000
43985000
43990000

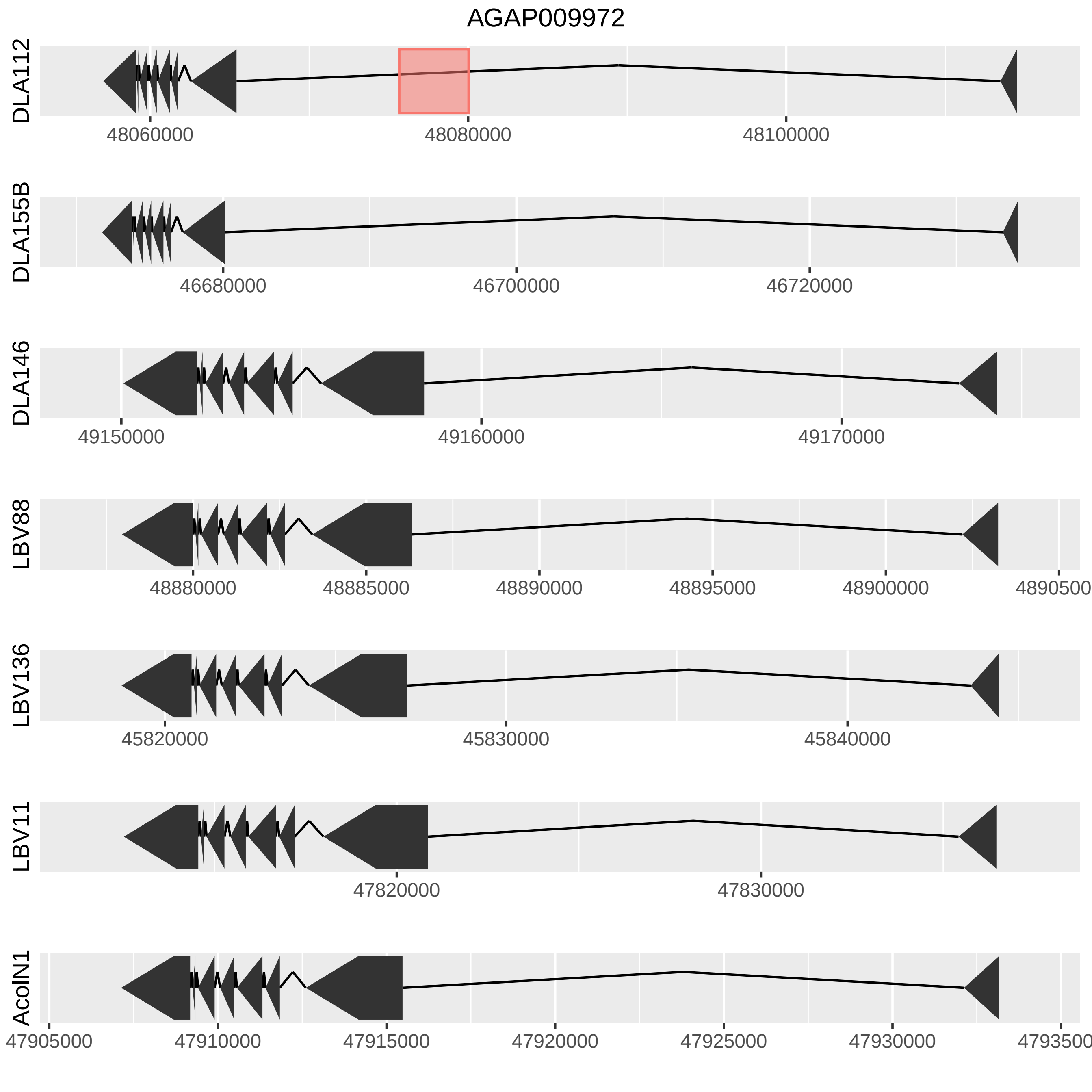

AGAP009972
DLA112
48060000
48080000
48100000
DLA155B
46680000
46700000
46720000
DLA146
49150000
49160000
49170000
LBV88
48880000
48885000
48890000
48895000
48900000
4890500
LBV136
45820000
45830000
45840000
LBV11
47820000
47830000
AcoIN1
47905000
47910000
47915000
47920000
47925000
47930000
4793500

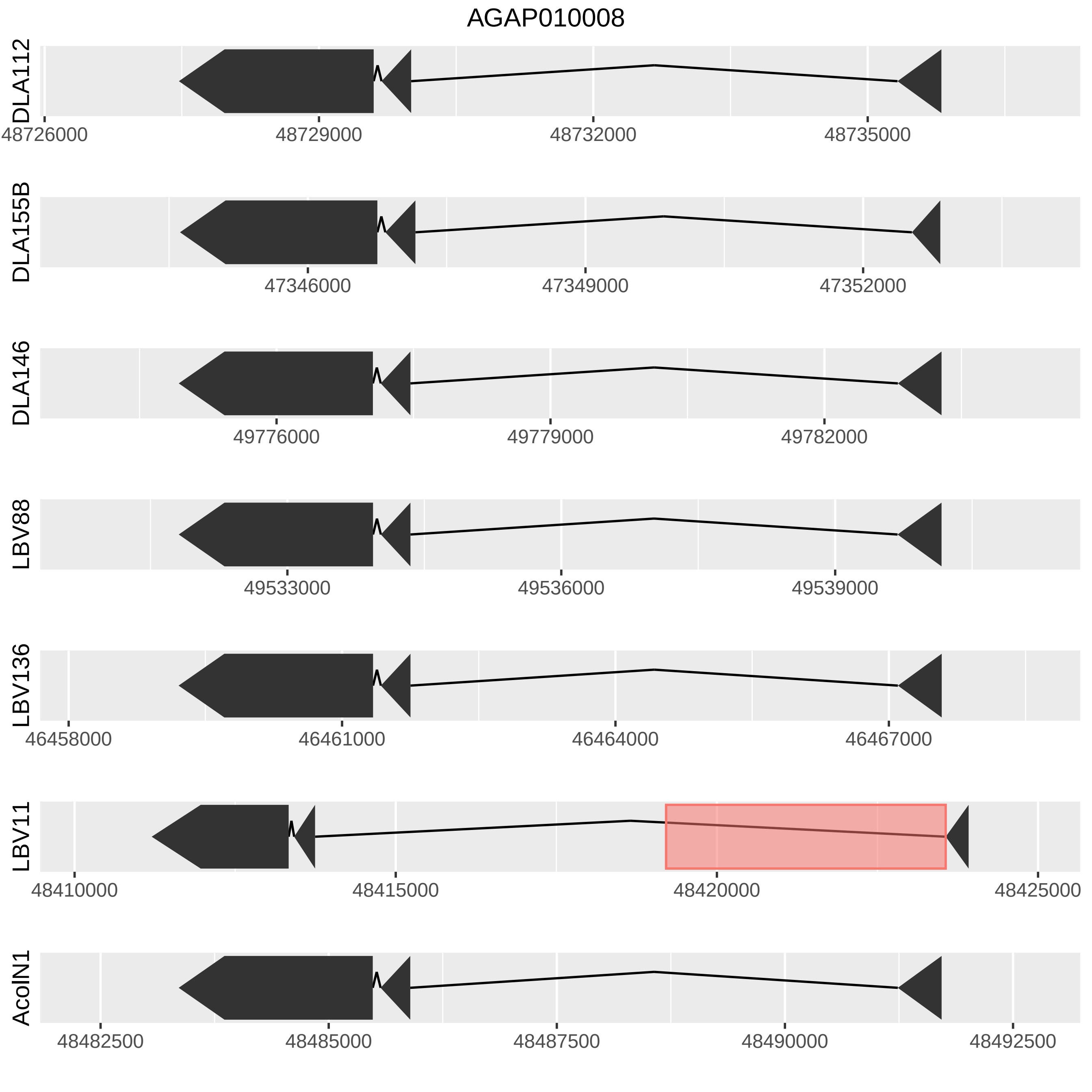

AGAP010008
DLA112
48726000
48729000
48732000
48735000
DLA155B
47346000
47349000
47352000
DLA146
49776000
49779000
49782000
LBV88
49533000
49536000
49539000
LBV136
46458000
46461000
46464000
46467000
LBV11
48410000
48415000
48420000
48425000
AcolN1
48482500
48485000
48487500
48490000
48492500

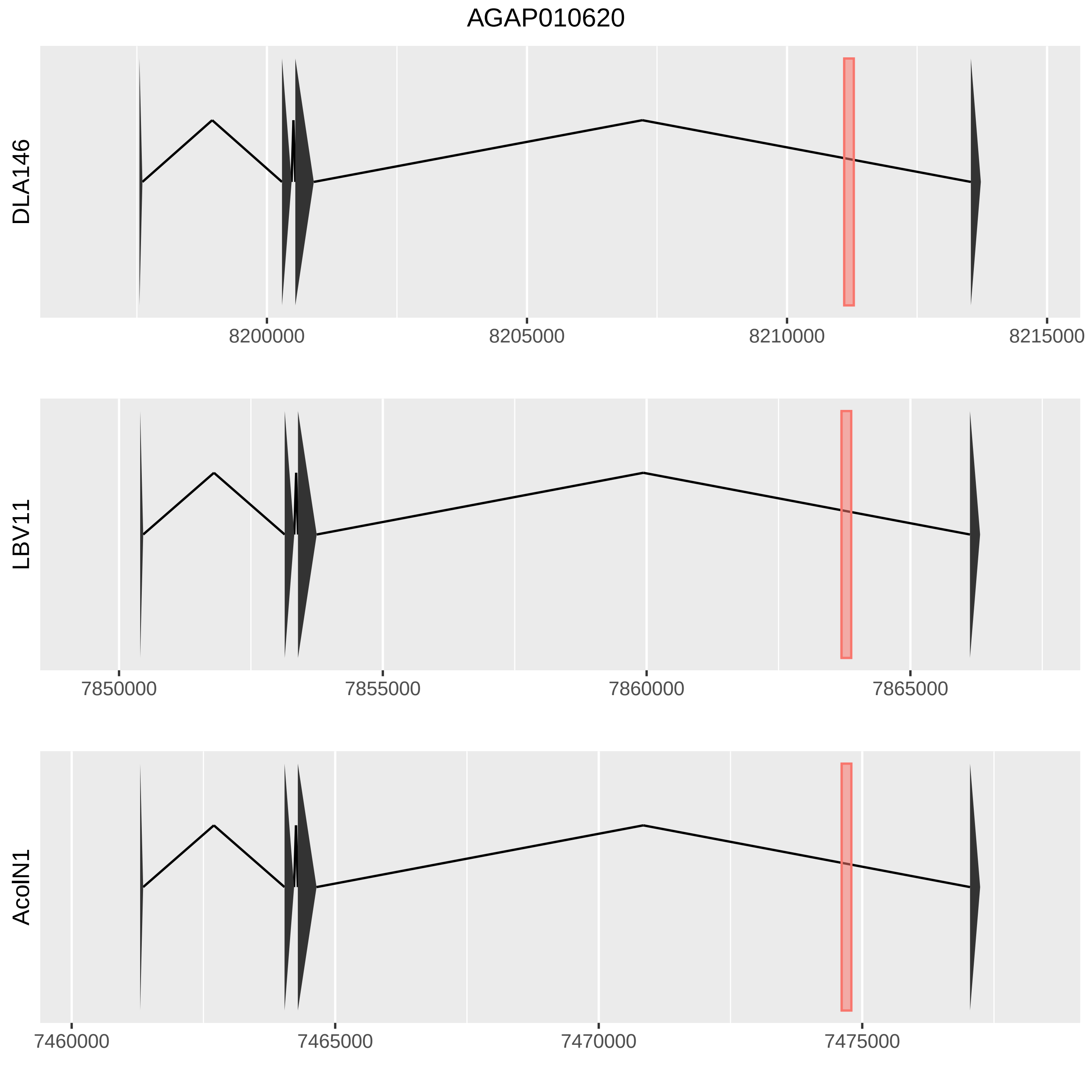
AGAP010620
DLA146
8200000
8205000
8210000
8215000
LBV11
7850000
7855000
7860000
7865000
AcolN1
7460000
7465000
7470000
7475000

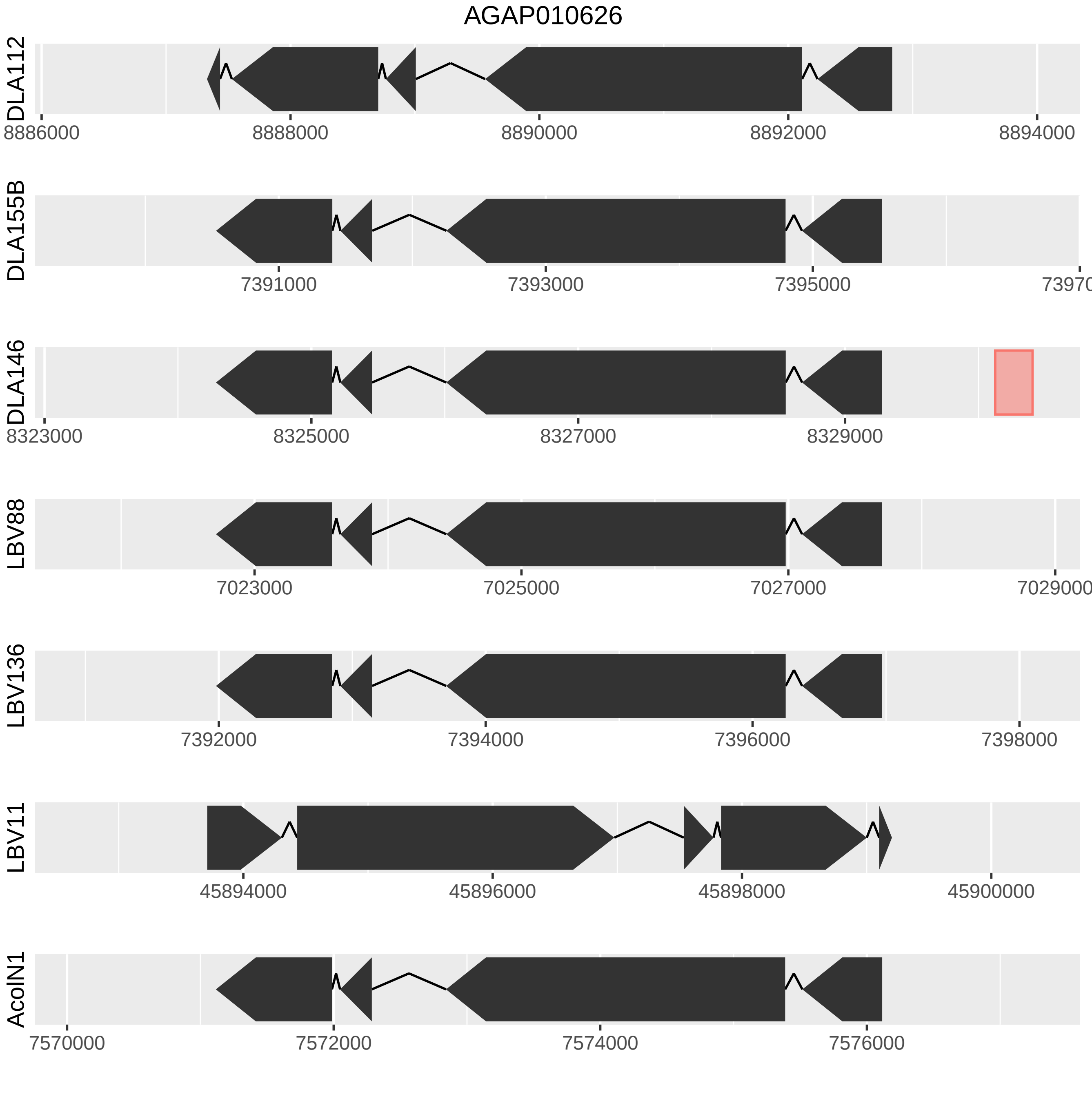
AGAP010626
DLA112
8886000
8888000
8890000
8892000
8894000
DLA155B
7391000
7393000
7395000
7397000
DLA146
8323000
8325000
8327000
8329000
LBV88
7023000
7025000
7027000
7029000
LBV136
7392000
7394000
7396000
7398000
LBV11
45894000
45896000
45898000
45900000
AcolN1
7570000
7572000
7574000
7576000

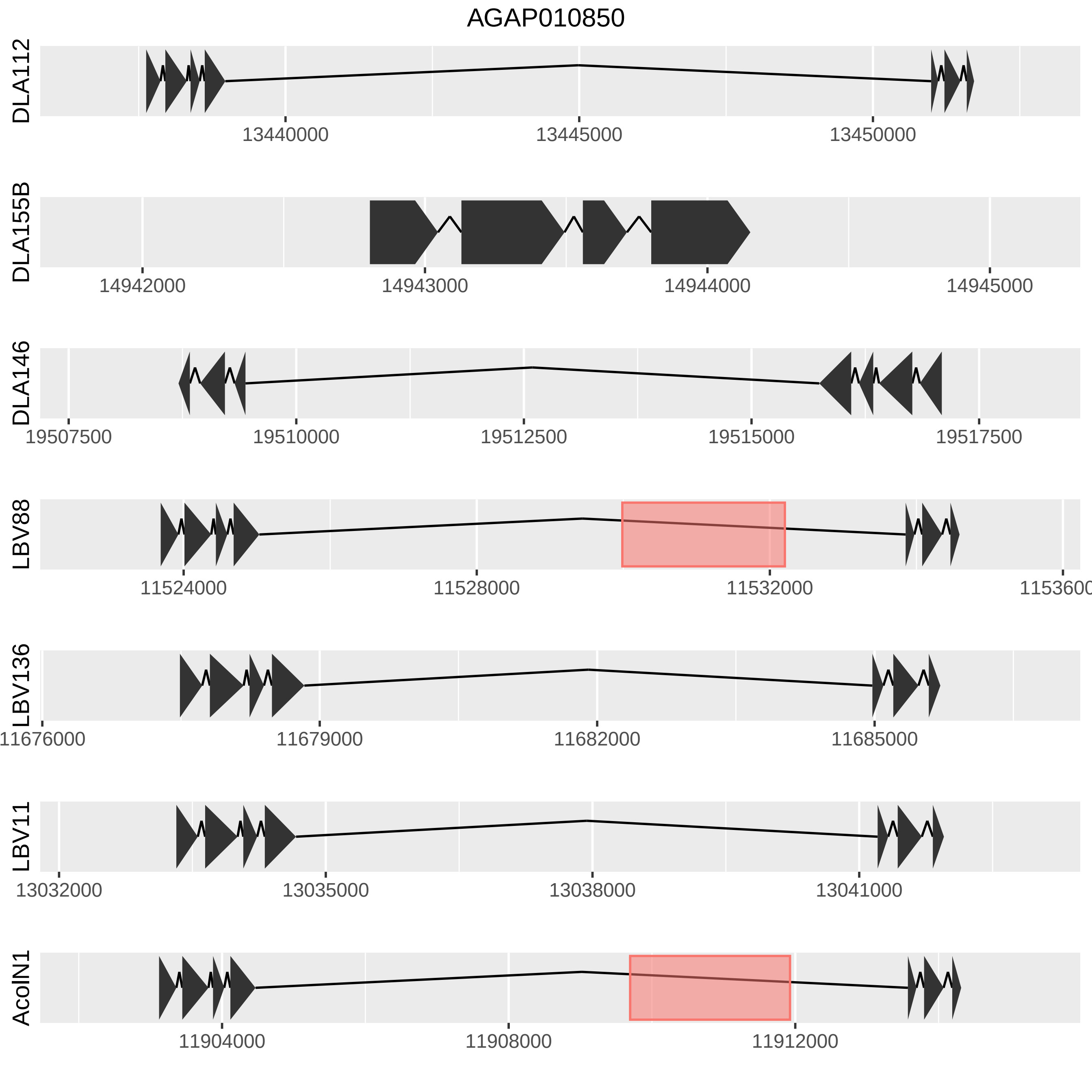
AGAP010850
DLA112
13440000
13445000
13450000
DLA155B
14942000
14943000
14944000
14945000
DLA146
19507500
19510000
19512500
19515000
19517500
LBV88
11524000
11528000
11532000
1153600
LBV136
11676000
11679000
11682000
11685000
LBV11
13032000
13035000
13038000
13041000
AcolN1
11904000
11908000
11912000

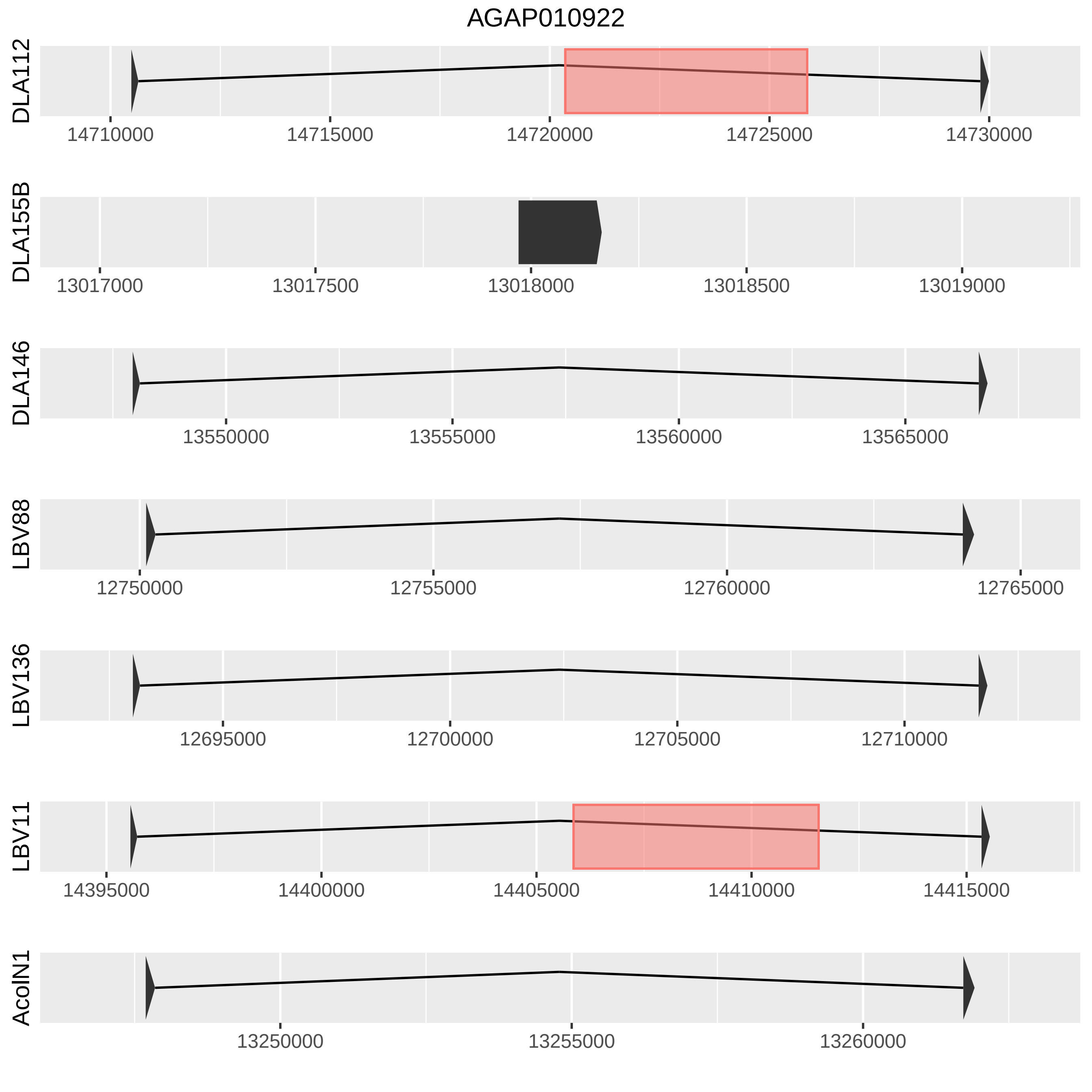
AGAP010922
DLA112
14710000
14715000
14720000
14725000
14730000
DLA155B
13017000
13017500
13018000
13018500
13019000
DLA146
13550000
13555000
13560000
13565000
LBV88
12750000
12755000
12760000
12765000
LBV136
12695000
12700000
12705000
12710000
LBV11
14395000
14400000
14405000
14410000
14415000
AcoIN1
13250000
13255000
13260000

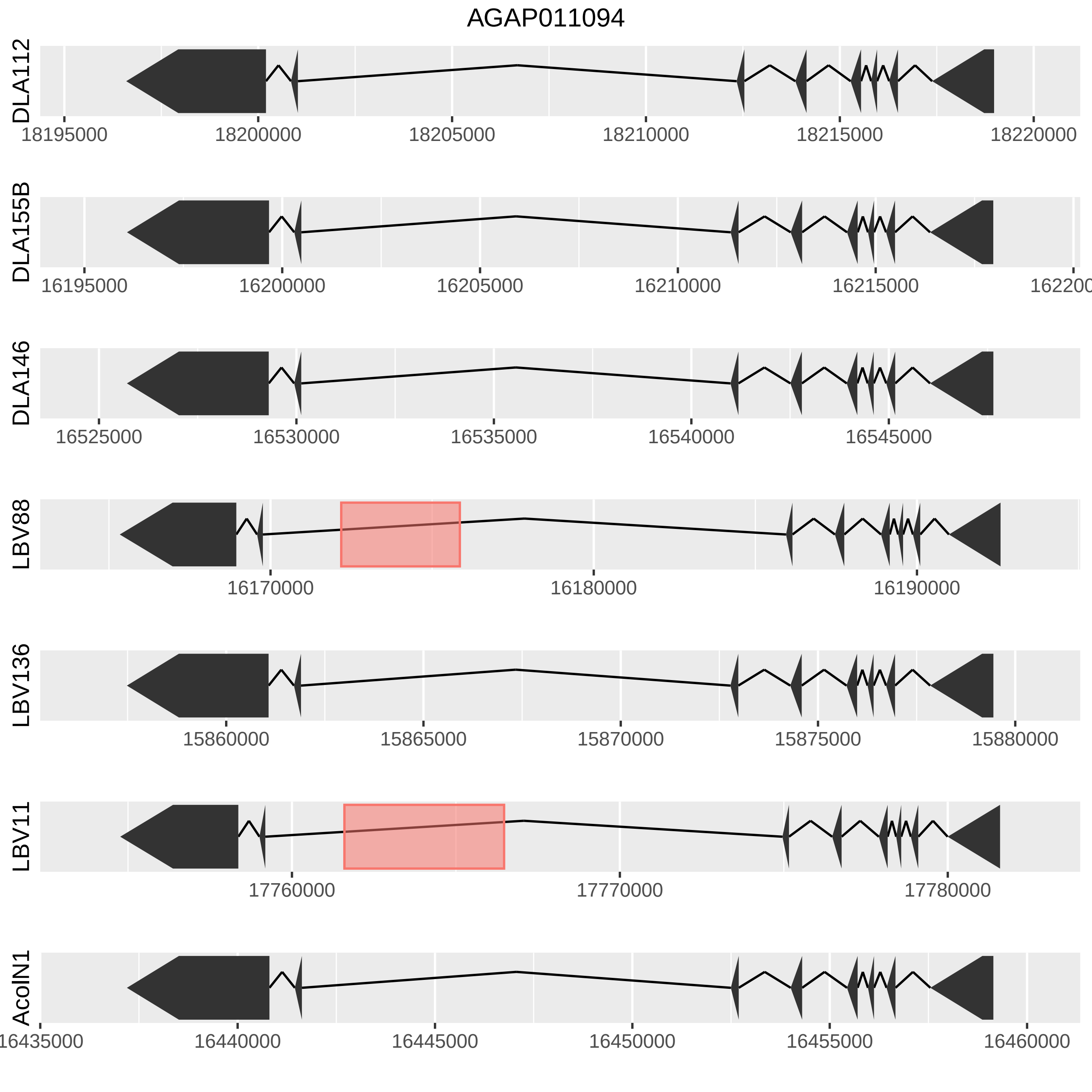

AGAP011094
DLA112
18195000
18200000
18205000
18210000
18215000
18220000
DLA155B
16195000
16200000
16205000
16210000
16215000
DLA146
16525000
16530000
16535000
16540000
16545000
LBV88
16170000
16180000
16190000
LBV136
15860000
15865000
15870000
15875000
15880000
LBV11
17760000
17770000
17780000
AcolN1
16435000
16440000
16445000
16450000
16455000
16460000

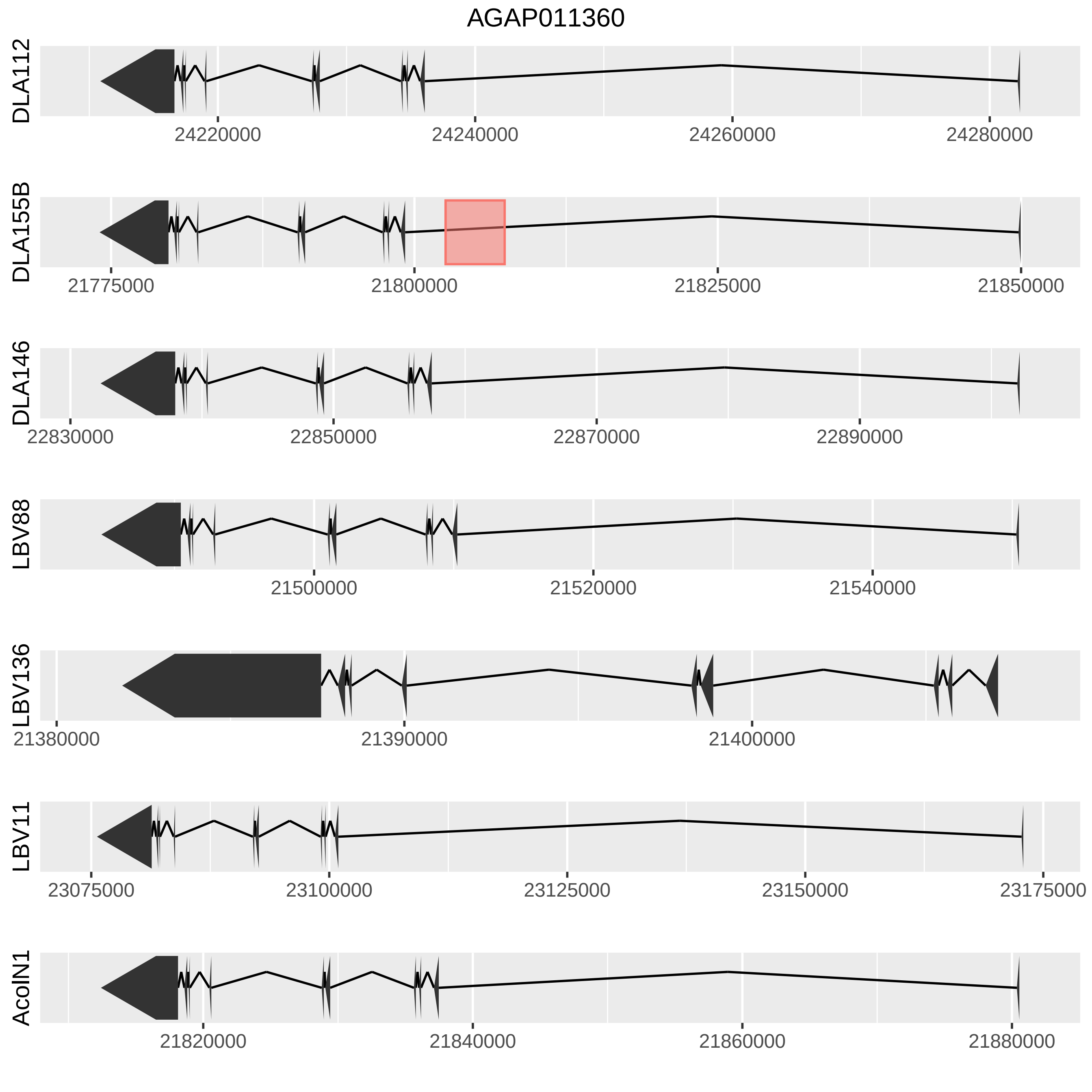

AGAP011360
DLA112
24220000
24240000
24260000
24280000
DLA155B
21775000
21800000
21825000
21850000
DLA146
22830000
22850000
22870000
22890000
LBV88
21500000
21520000
21540000
LBV136
21380000
21390000
21400000
LBV11
23075000
23100000
23125000
23150000
23175000
AcolN1
21820000
21840000
21860000
21880000

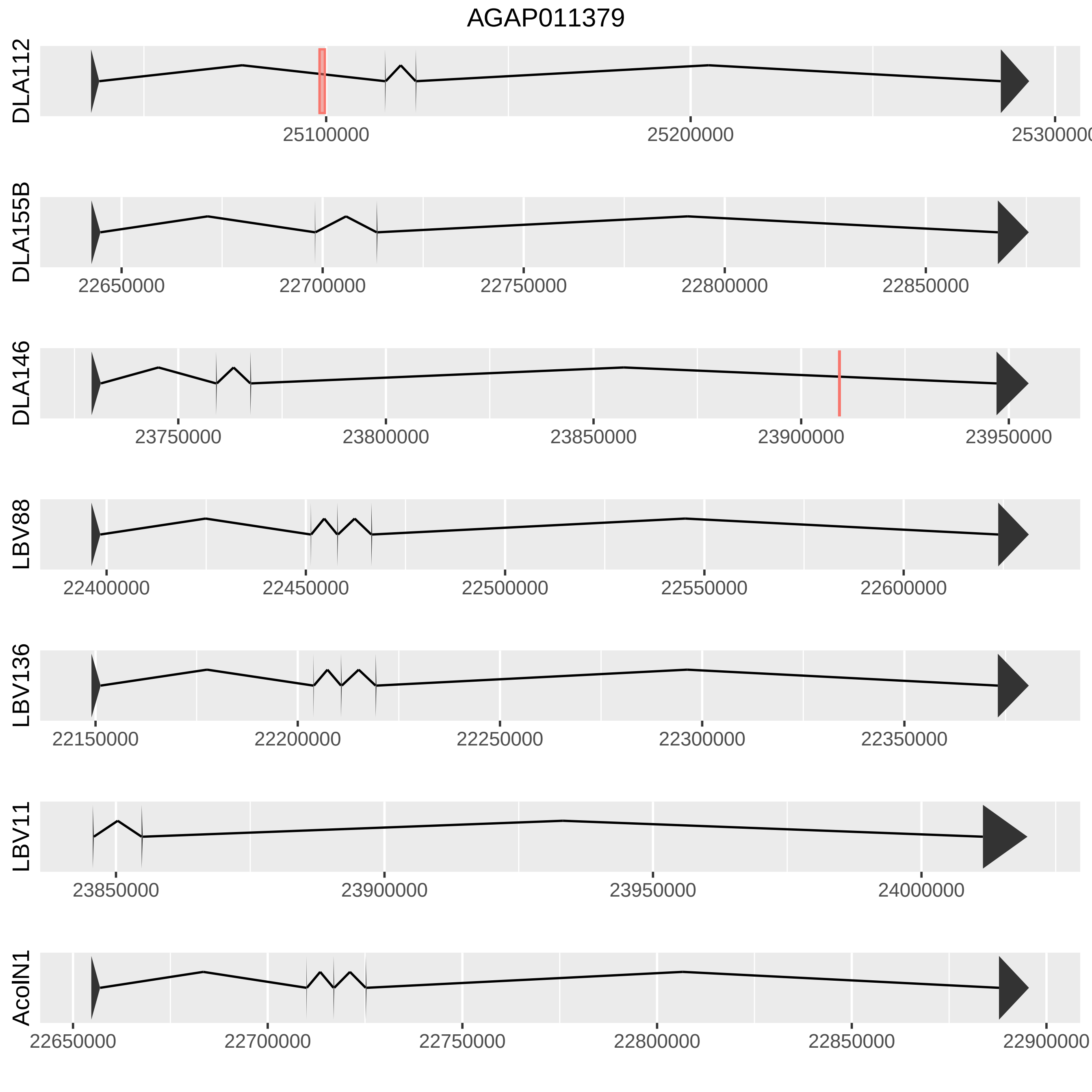
AGAP011379
DLA112
25100000
25200000
25300000
DLA155B
22650000
22700000
22750000
22800000
22850000
DLA146
23750000
23800000
23850000
23900000
23950000
LBV88
22400000
22450000
22500000
22550000
22600000
LBV136
22150000
22200000
22250000
22300000
22350000
LBV11
23850000
23900000
23950000
24000000
AcolN1
22650000
22700000
22750000
22800000
22850000
22900000

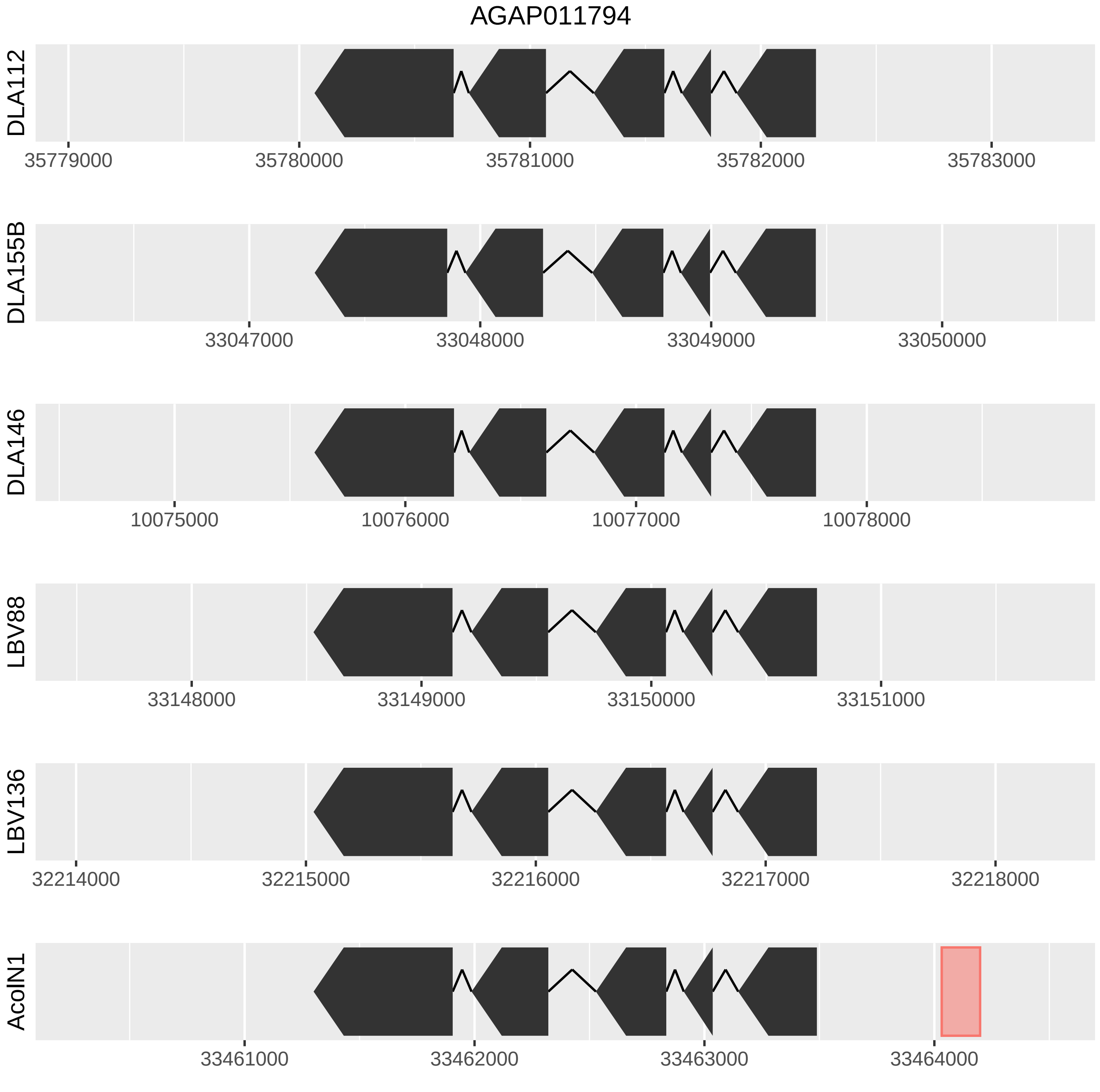

AGAP011794
DLA112
35779000
35780000
35781000
35782000
35783000
DLA155B
33047000
33048000
33049000
33050000
DLA146
10075000
10076000
10077000
10078000
LBV88
33148000
33149000
33150000
33151000
LBV136
32214000
32215000
32216000
32217000
32218000
AcolN1
33461000
33462000
33463000
33464000

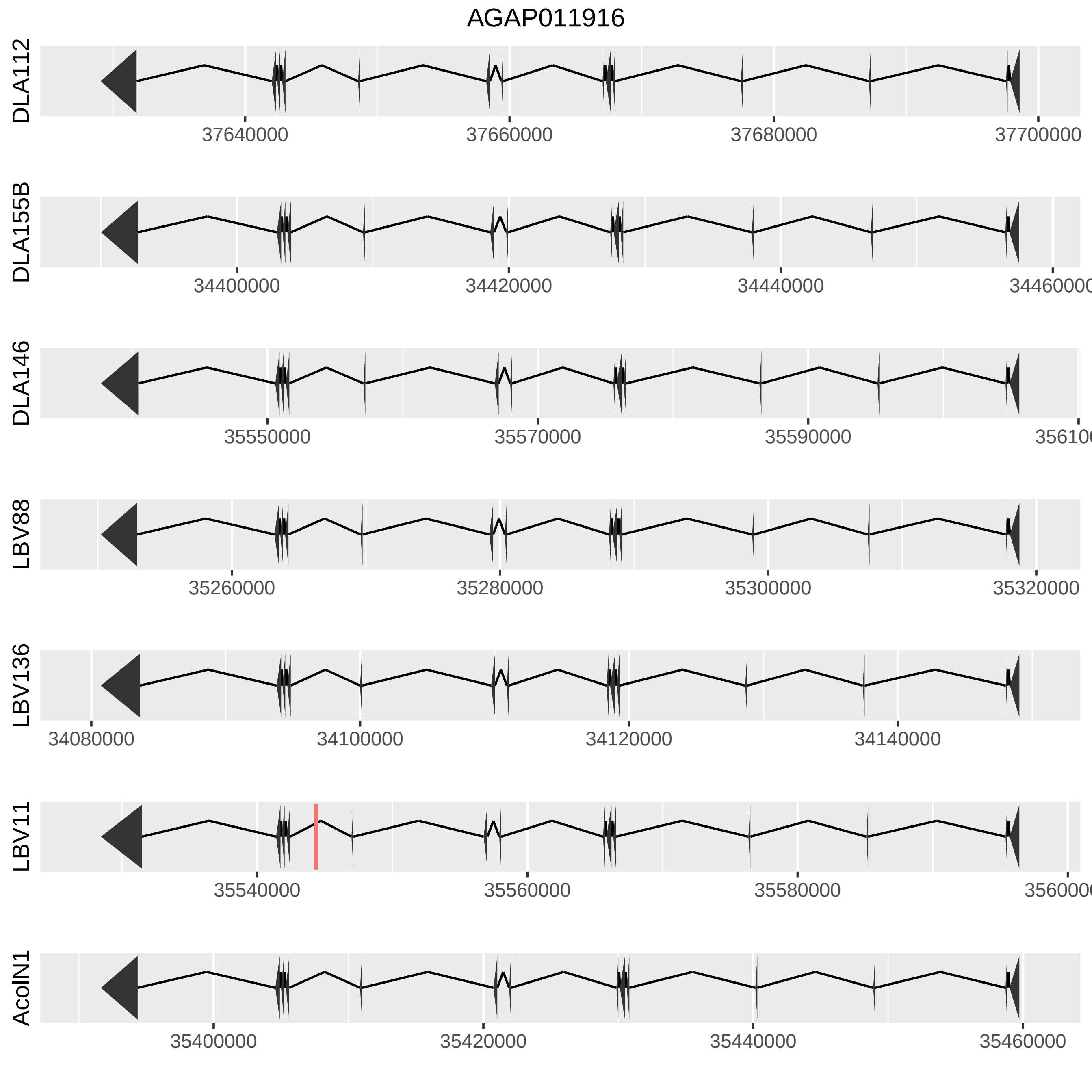
AGAP011916
DLA112
37640000
37660000
37680000
37700000
DLA155B
34400000
34420000
34440000
34460000
DLA146
35550000
35570000
35590000
35610000
LBV88
35260000
35280000
35300000
35320000
LBV136
34080000
34100000
34120000
34140000
LBV11
35540000
35560000
35580000
35600000
AcolN1
35400000
35420000
35440000
35460000

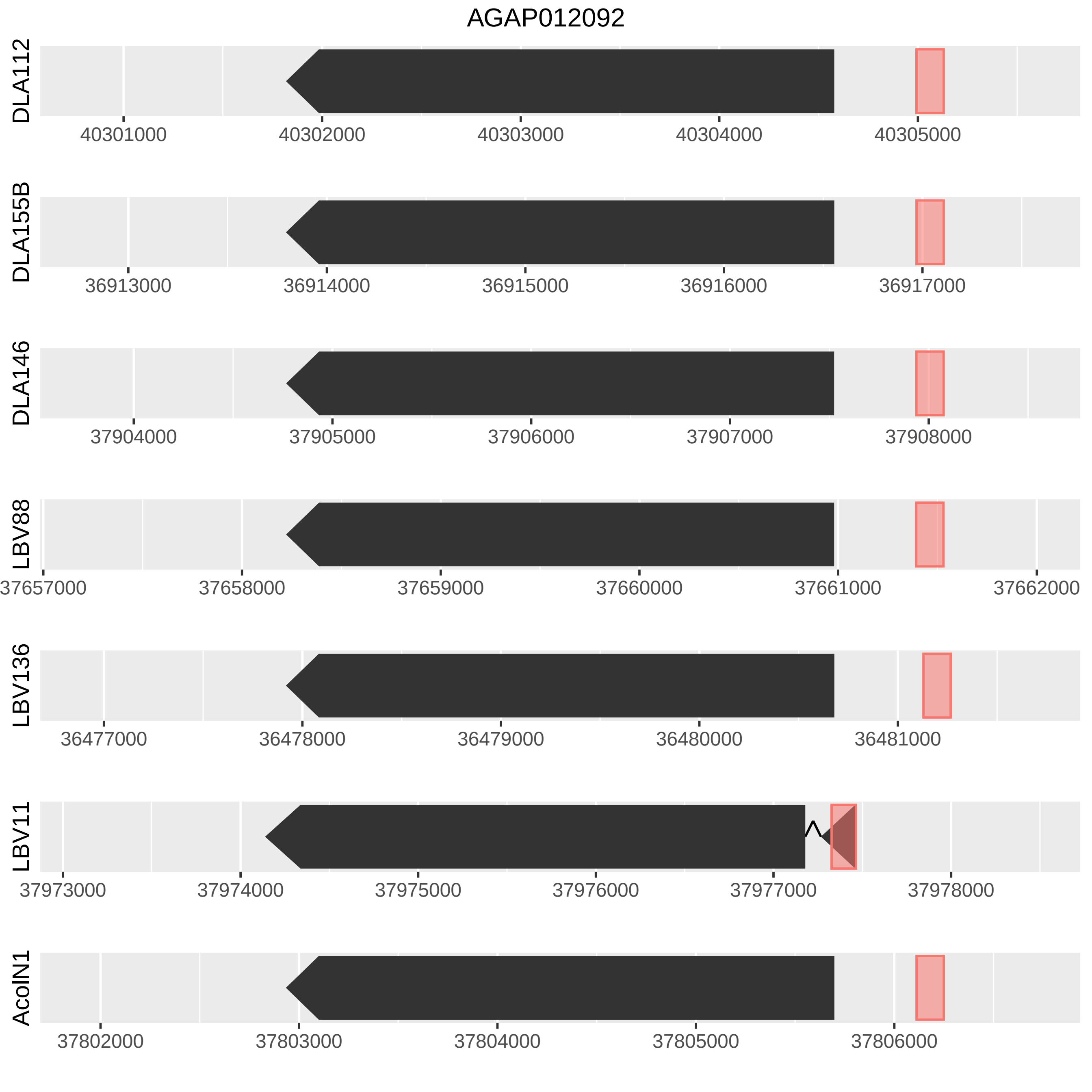
AGAP012092
DLA112
40301000
40302000
40303000
40304000
40305000
DLA155B
36913000
36914000
36915000
36916000
36917000
DLA146
37904000
37905000
37906000
37907000
37908000
LBV88
37657000
37658000
37659000
37660000
37661000
37662000
LBV136
36477000
36478000
36479000
36480000
36481000
LBV11
37973000
37974000
37975000
37976000
37977000
37978000
AcoIN1
37802000
37803000
37804000
37805000
37806000

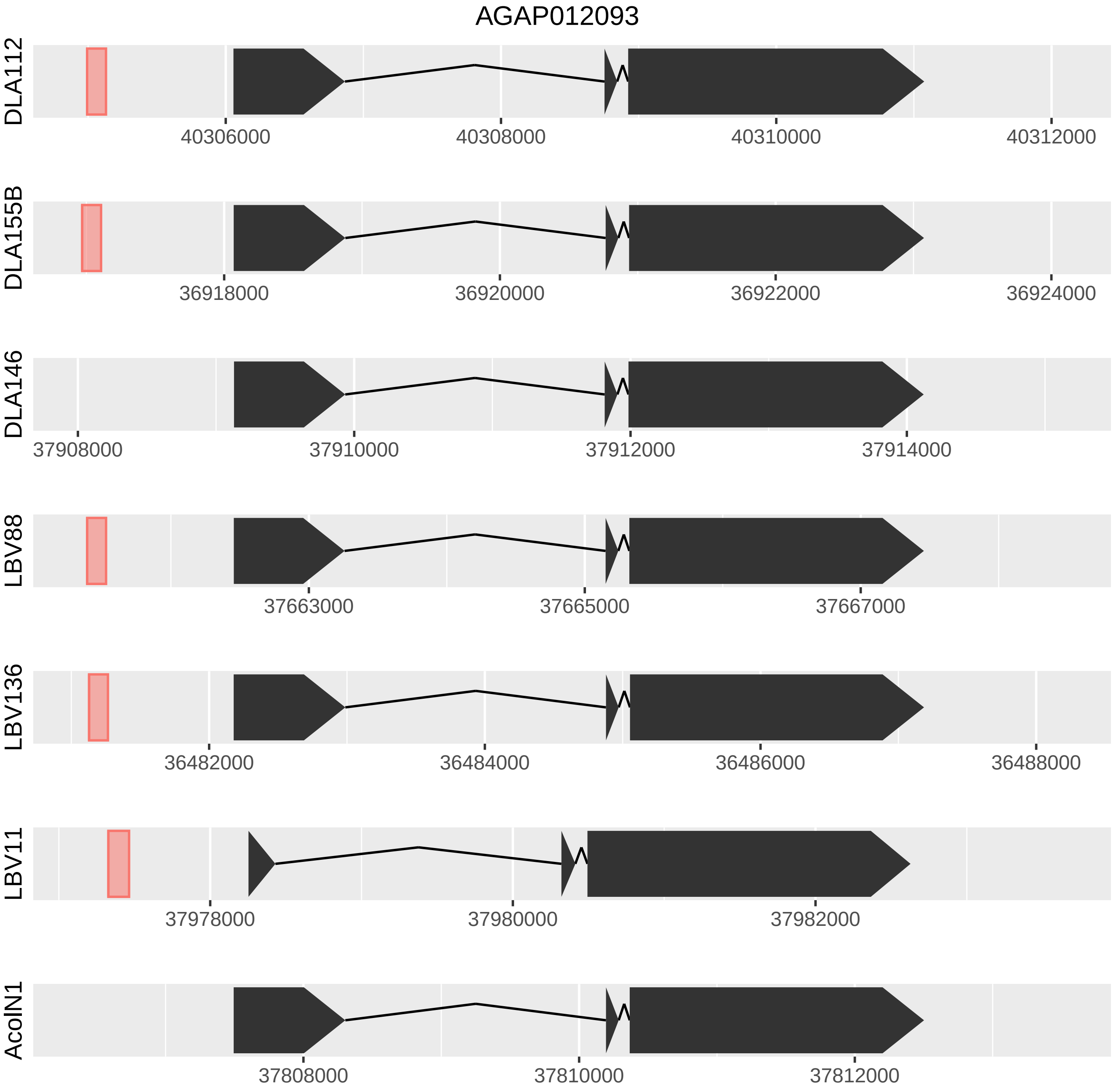
AGAP012093
DLA112
40306000
40308000
40310000
40312000
DLA155B
36918000
36920000
36922000
36924000
DLA146
37908000
37910000
37912000
37914000
LBV88
37663000
37665000
37667000
LBV136
36482000
36484000
36486000
36488000
LBV11
37978000
37980000
37982000
AcolN1
37808000
37810000
37812000

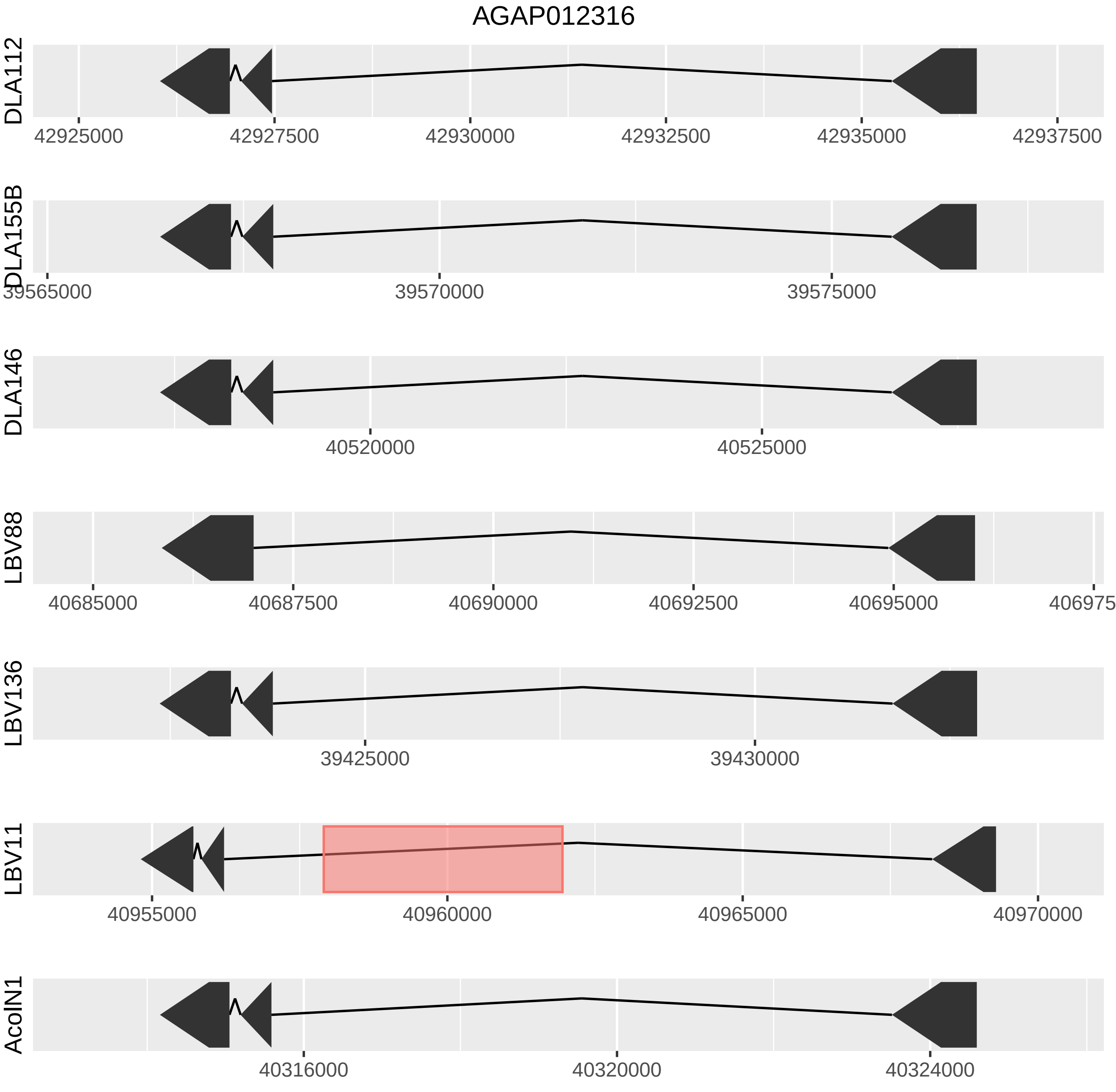
AGAP012316
DLA112
42925000
42927500
42930000
42932500
42935000
42937500
DLA155B
39565000
39570000
39575000
DLA146
40520000
40525000
LBV88
40685000
40687500
40690000
40692500
40695000
LBV136
39425000
39430000
LBV11
40955000
40960000
40965000
40970000
AcoIN1
40316000
40320000
40324000

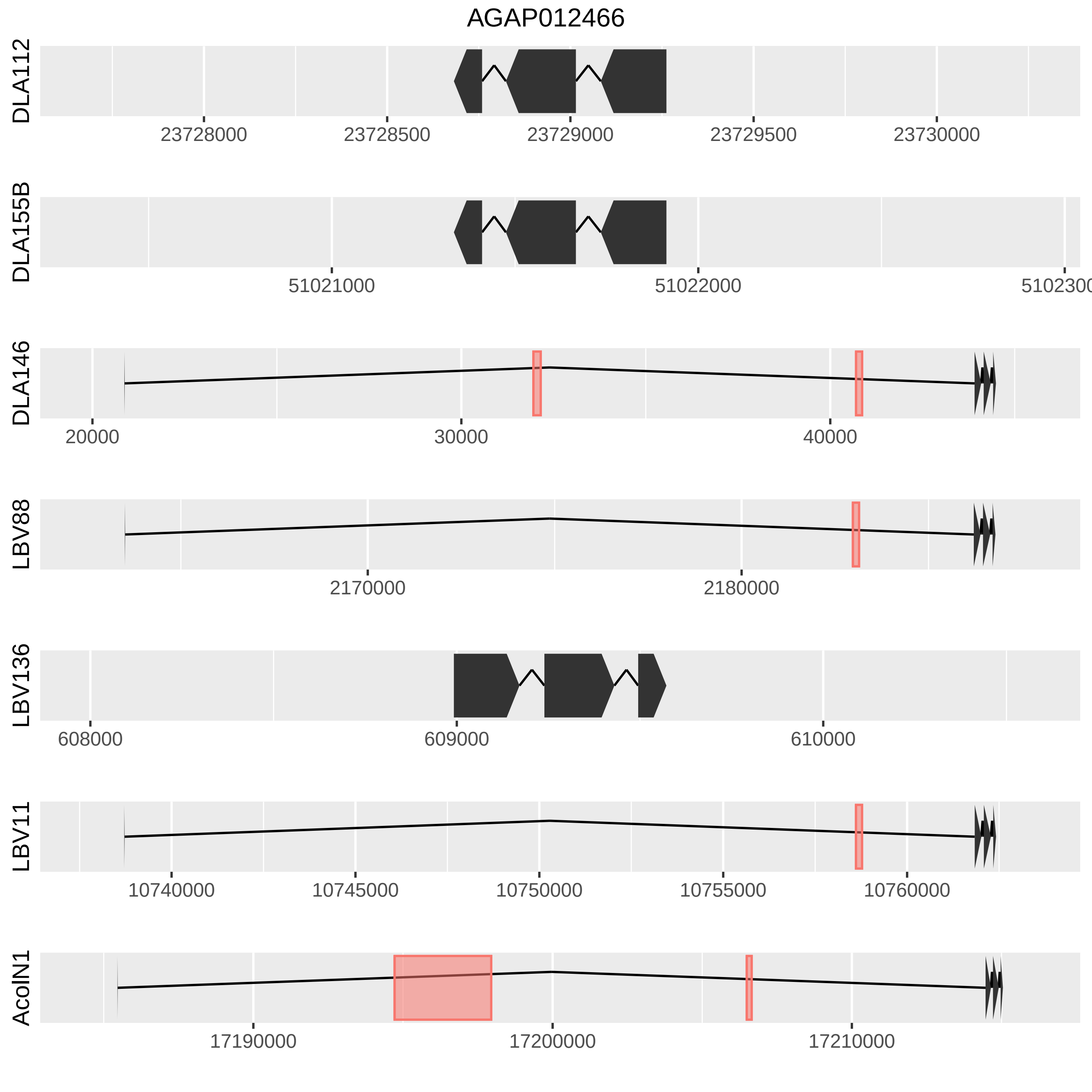

AGAP012466
DLA112
23728000
23728500
23729000
23729500
23730000
DLA155B
51021000
51022000
5102300
DLA146
20000
30000
40000
LBV88
2170000
2180000
LBV136
608000
609000
610000
LBV11
10740000
10745000
10750000
10755000
10760000
AcolN1
17190000
17200000
17210000

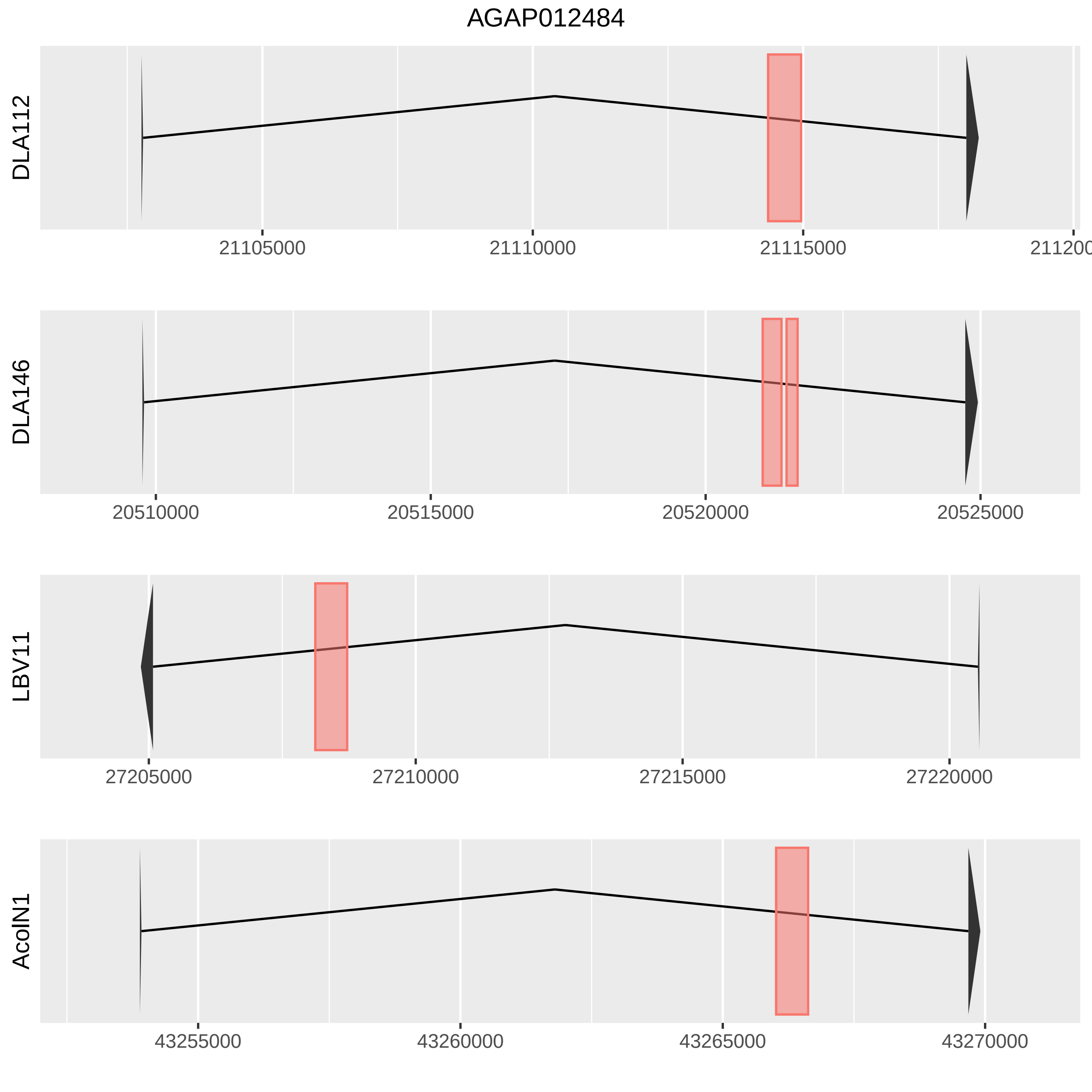
AGAP012484
DLA112
21105000
21110000
21115000
21120000
DLA146
20510000
20515000
20520000
20525000
LBV11
27205000
27210000
27215000
27220000
AcolN1
43255000
43260000
43265000
43270000

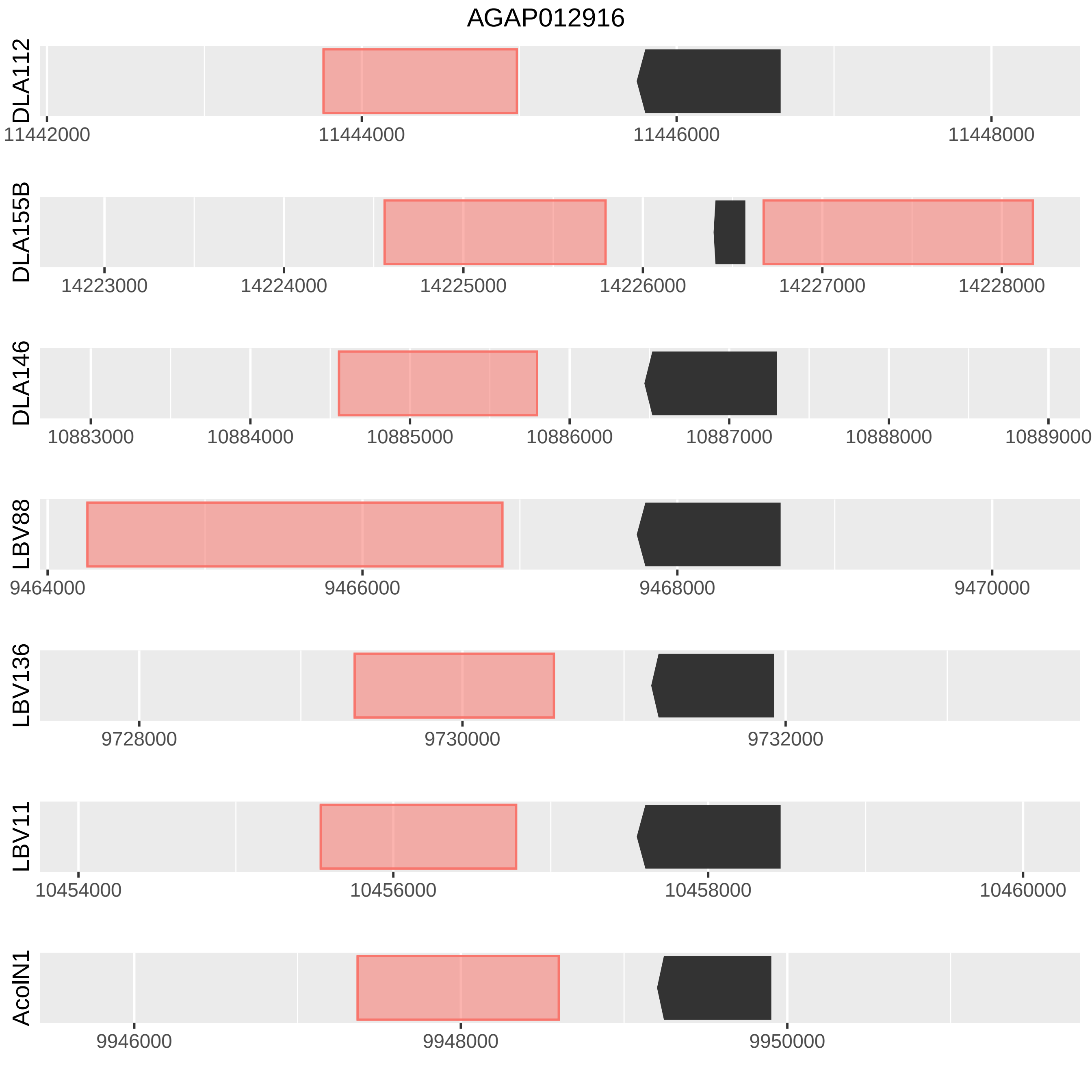
AGAP012916
DLA112
11442000
11444000
11446000
11448000
DLA155B
14223000
14224000
14225000
14226000
14227000
14228000
DLA146
10883000
10884000
10885000
10886000
10887000
10888000
10889000
LBV88
9464000
9466000
9468000
9470000
LBV136
9728000
9730000
9732000
LBV11
10454000
10456000
10458000
10460000
AcoIN1
9946000
9948000
9950000

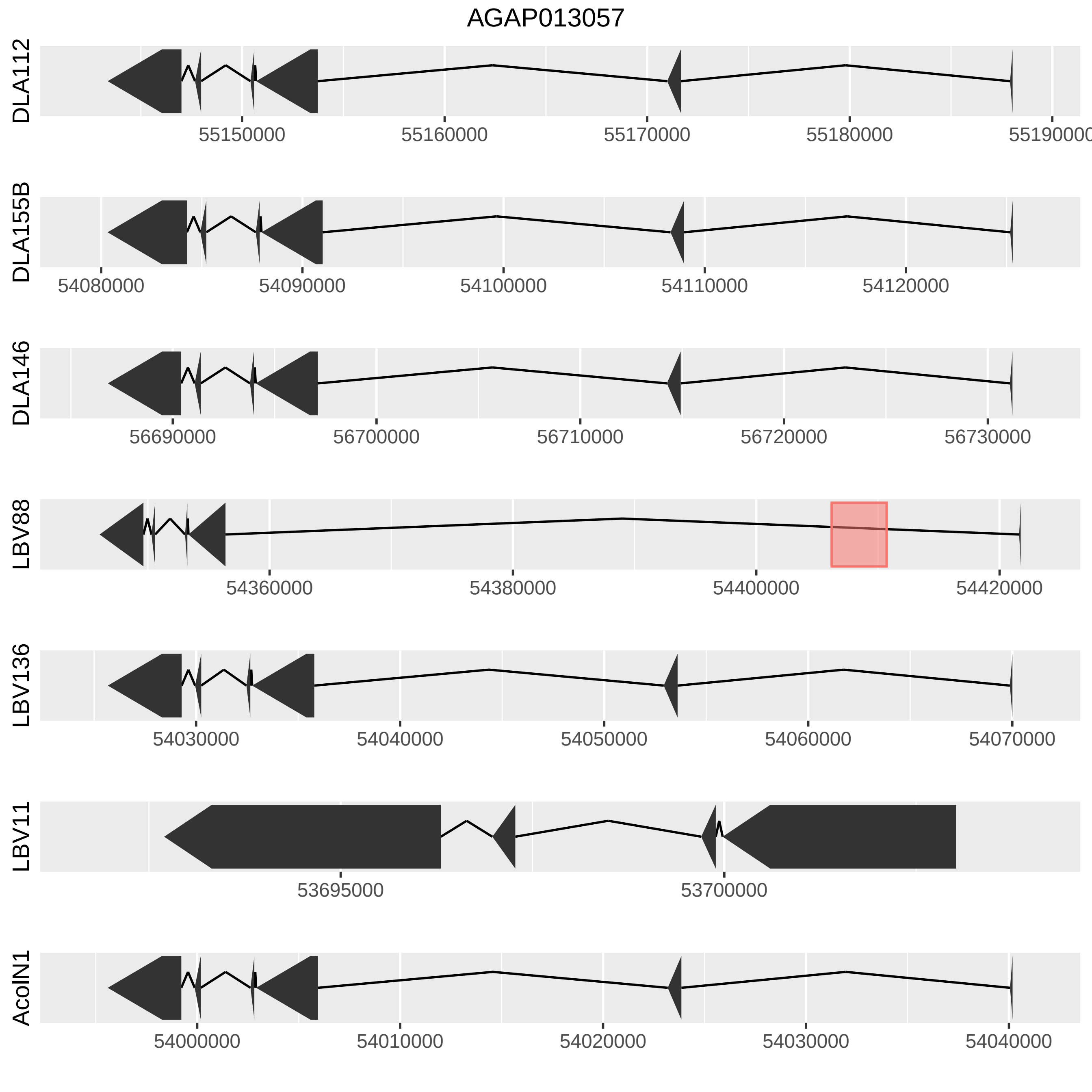

AGAP013057
DLA112
55150000
55160000
55170000
55180000
55190000
DLA155B
54080000
54090000
54100000
54110000
54120000
DLA146
56690000
56700000
56710000
56720000
56730000
LBV88
54360000
54380000
54400000
54420000
LBV136
54030000
54040000
54050000
54060000
54070000
LBV11
53695000
53700000
AcolN1
54000000
54010000
54020000
54030000
54040000

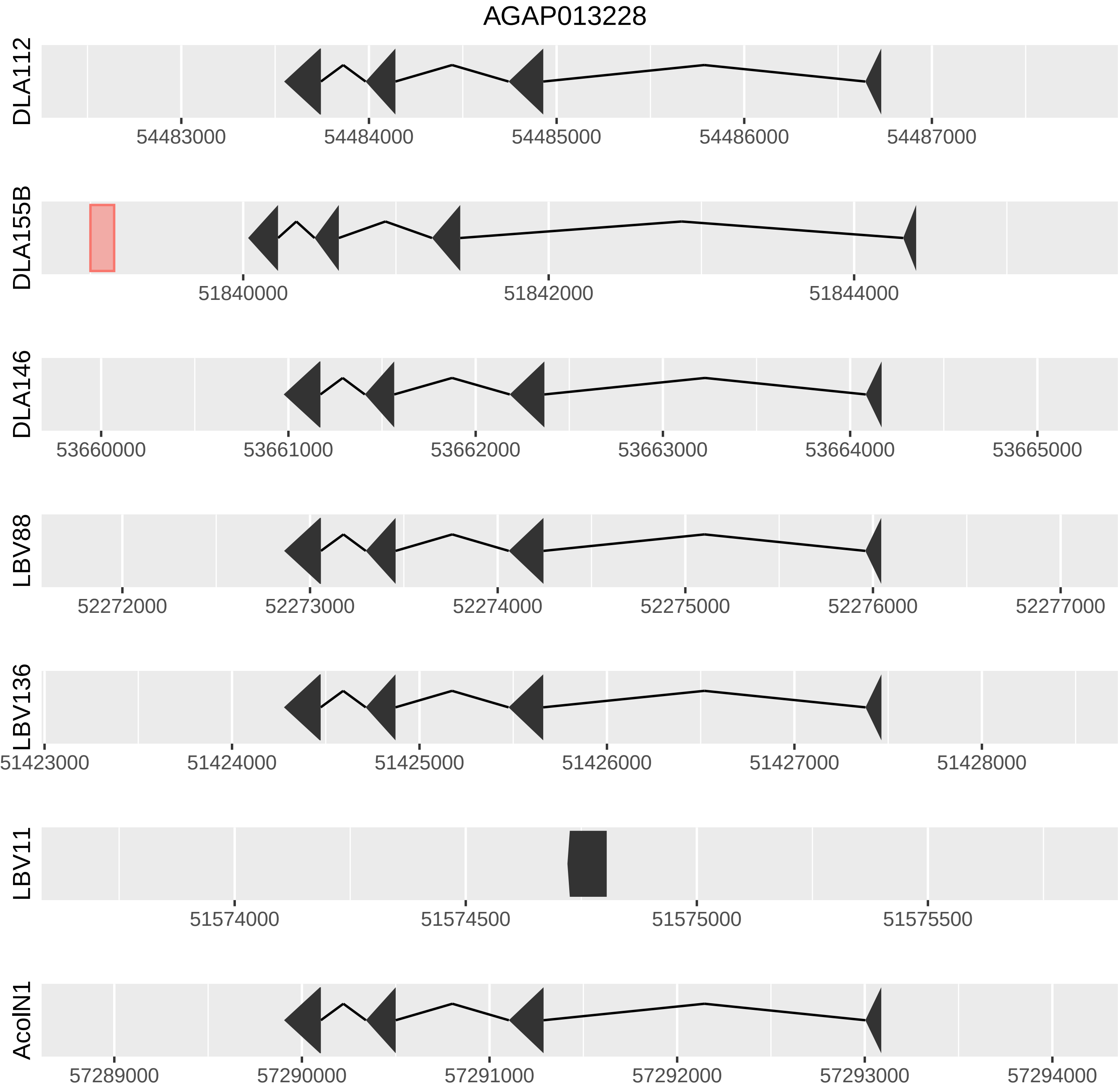
AGAP013228
DLA112
54483000
54484000
54485000
54486000
54487000
DLA155B
51840000
51842000
51844000
DLA146
53660000
53661000
53662000
53663000
53664000
53665000
LBV88
52272000
52273000
52274000
52275000
52276000
52277000
LBV136
51423000
51424000
51425000
51426000
51427000
51428000
LBV11
51574000
51574500
51575000
51575500
AcolN1
57289000
57290000
57291000
57292000
57293000
57294000

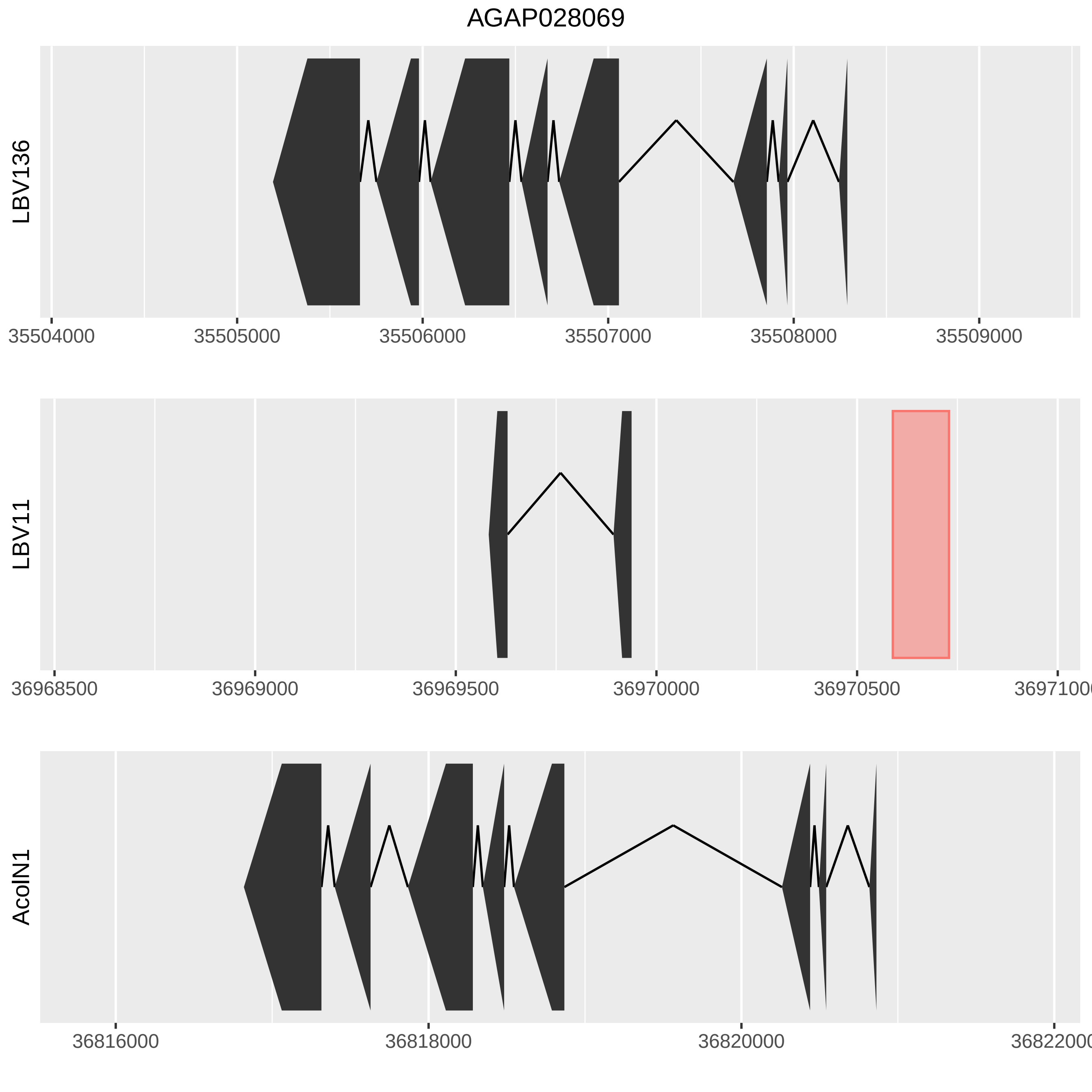
AGAP028069
LBV136
35504000
35505000
35506000
35507000
35508000
35509000
LBV11
36968500
36969000
36969500
36970000
36970500
36971000
AcolN1
36816000
36818000
36820000
36822000

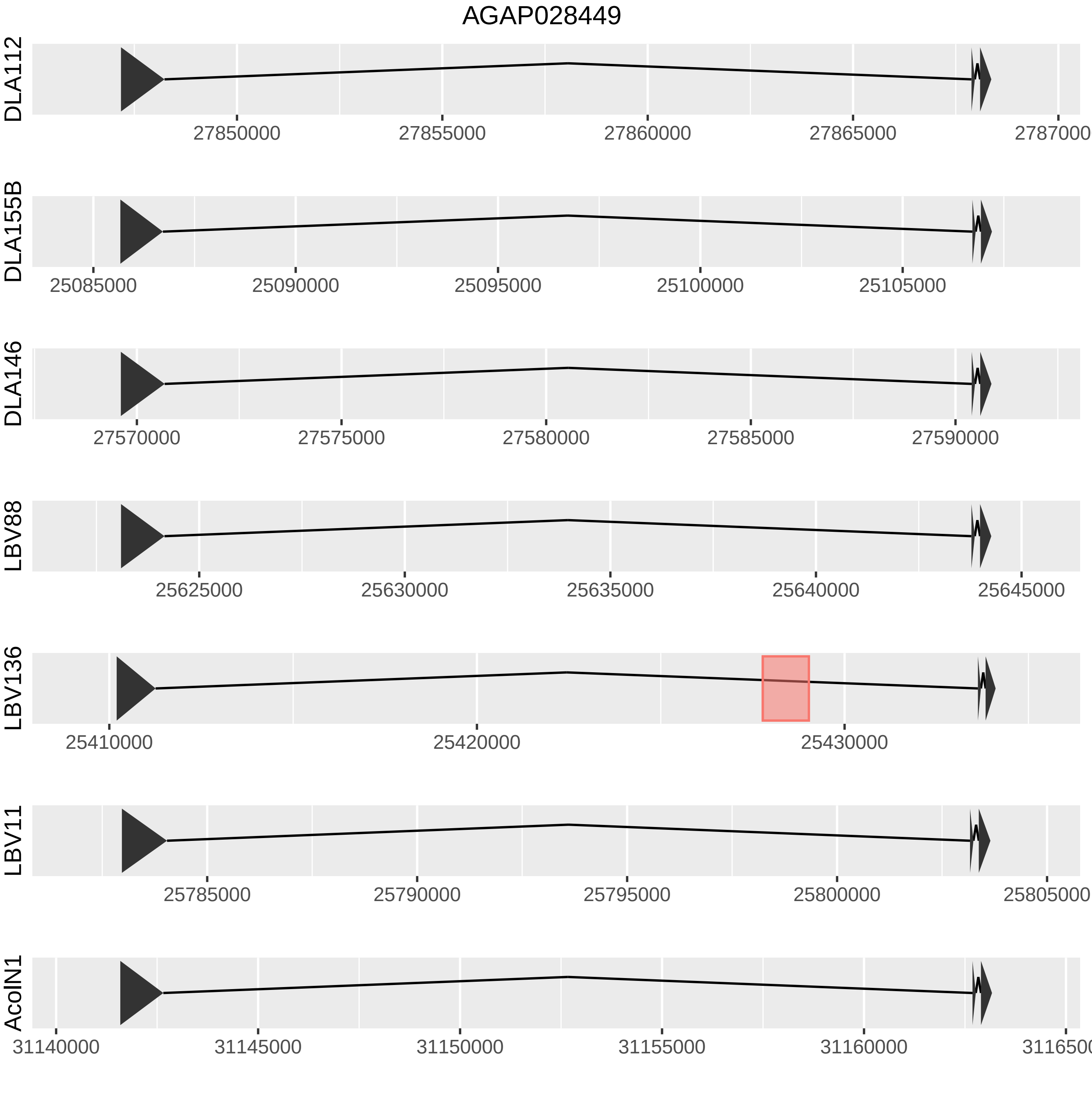

AGAP028449
DLA112
27850000
27855000
27860000
27865000
2787000
DLA155B
25085000
25090000
25095000
25100000
25105000
DLA146
27570000
27575000
27580000
27585000
27590000
LBV88
25625000
25630000
25635000
25640000
25645000
LBV136
25410000
25420000
25430000
LBV11
25785000
25790000
25795000
25800000
25805000
AcolN1
31140000
31145000
31150000
31155000
31160000
311650

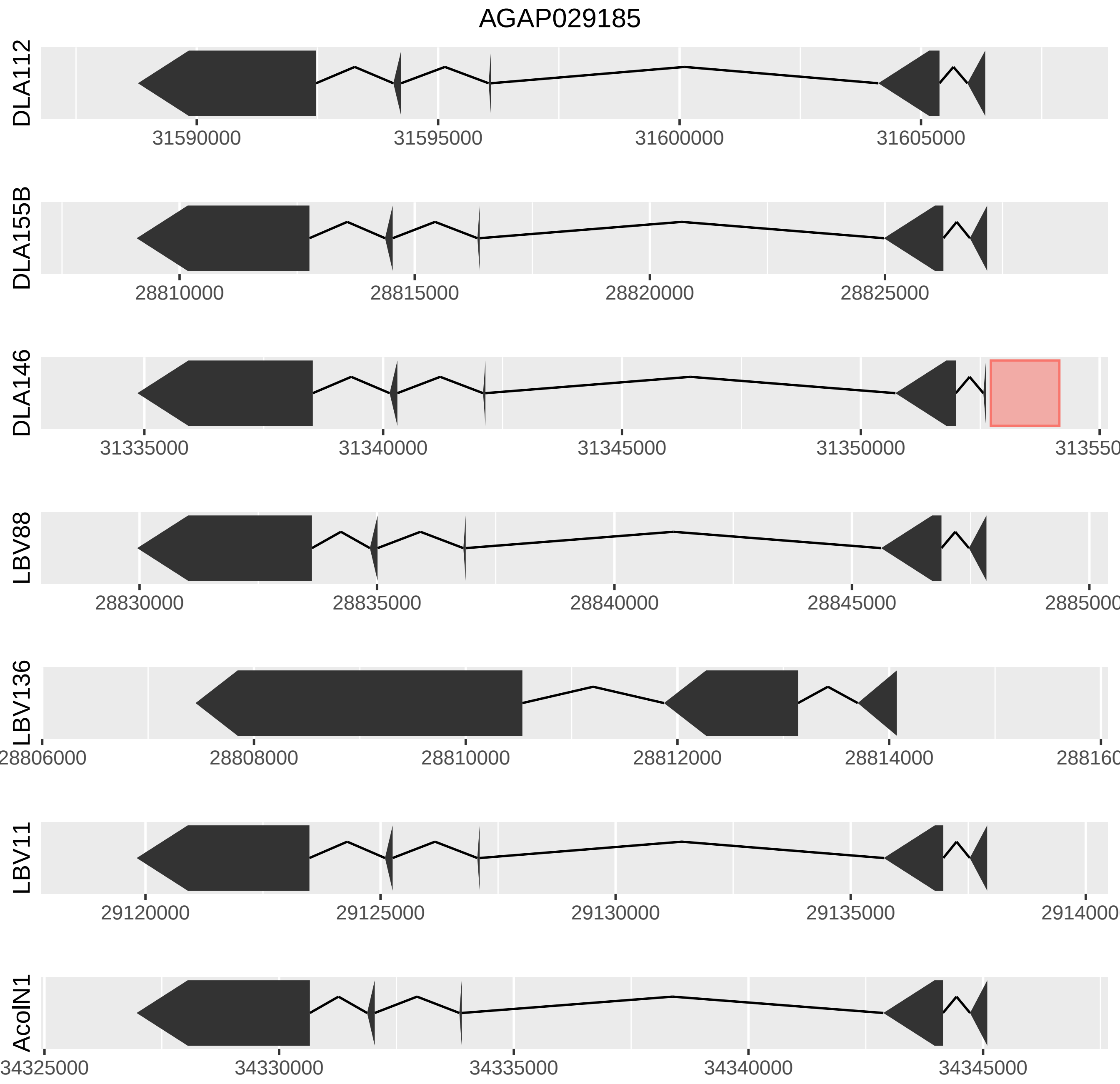

AGAP029185
DLA112
31590000
31595000
31600000
31605000
DLA155B
28810000
28815000
28820000
28825000
DLA146
31335000
31340000
31345000
31350000
313550
LBV88
28830000
28835000
28840000
28845000
2885000
LBV136
28806000
28808000
28810000
28812000
28814000
288160
LBV11
29120000
29125000
29130000
29135000
2914000
AcolN1
34325000
34330000
34335000
34340000
34345000

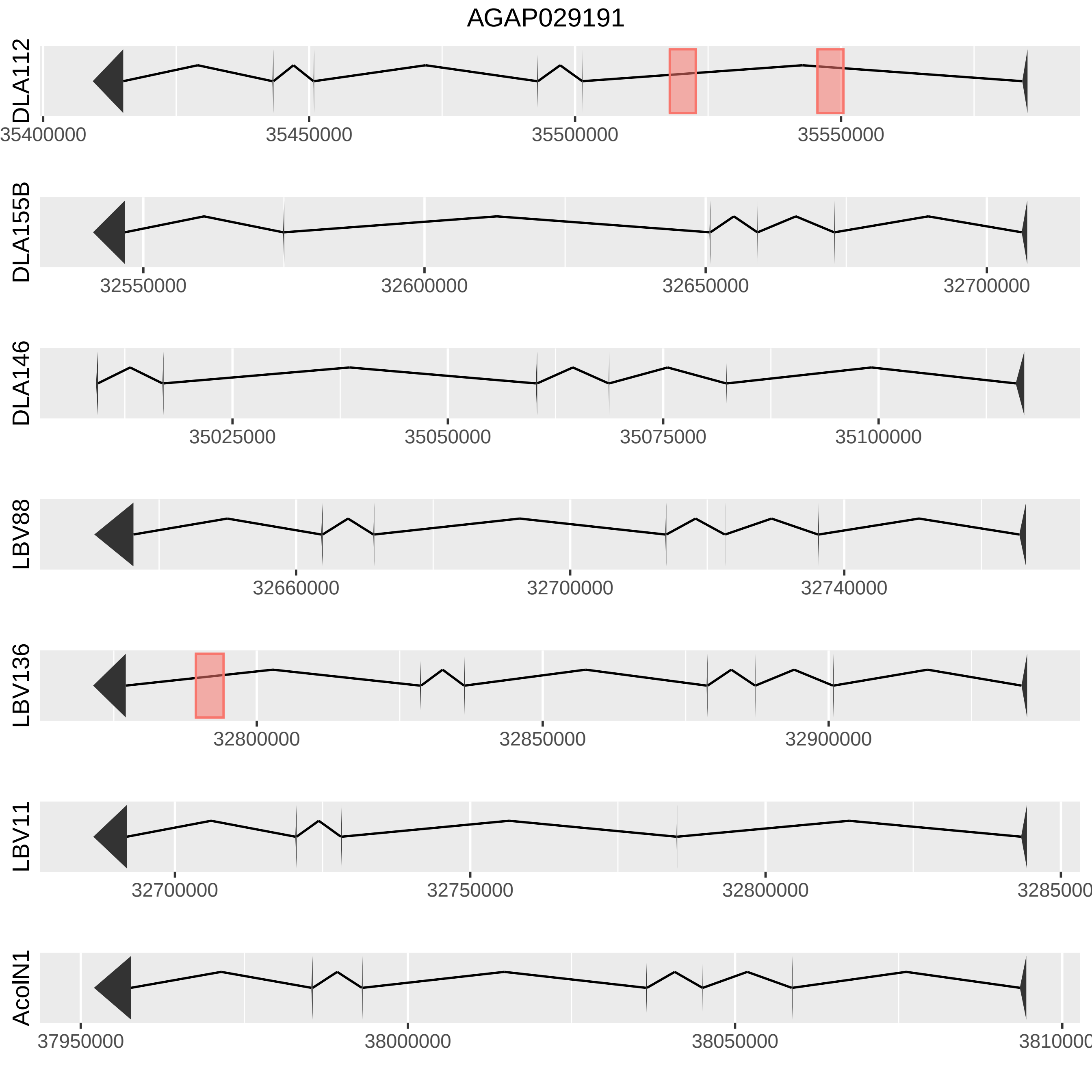
AGAP029191
DLA112
35400000
35450000
35500000
35550000
DLA155B
32550000
32600000
32650000
32700000
DLA146
35025000
35050000
35075000
35100000
LBV88
32660000
32700000
32740000
LBV136
32800000
32850000
32900000
LBV11
32700000
32750000
32800000
AcoIN1
37950000
38000000
38050000

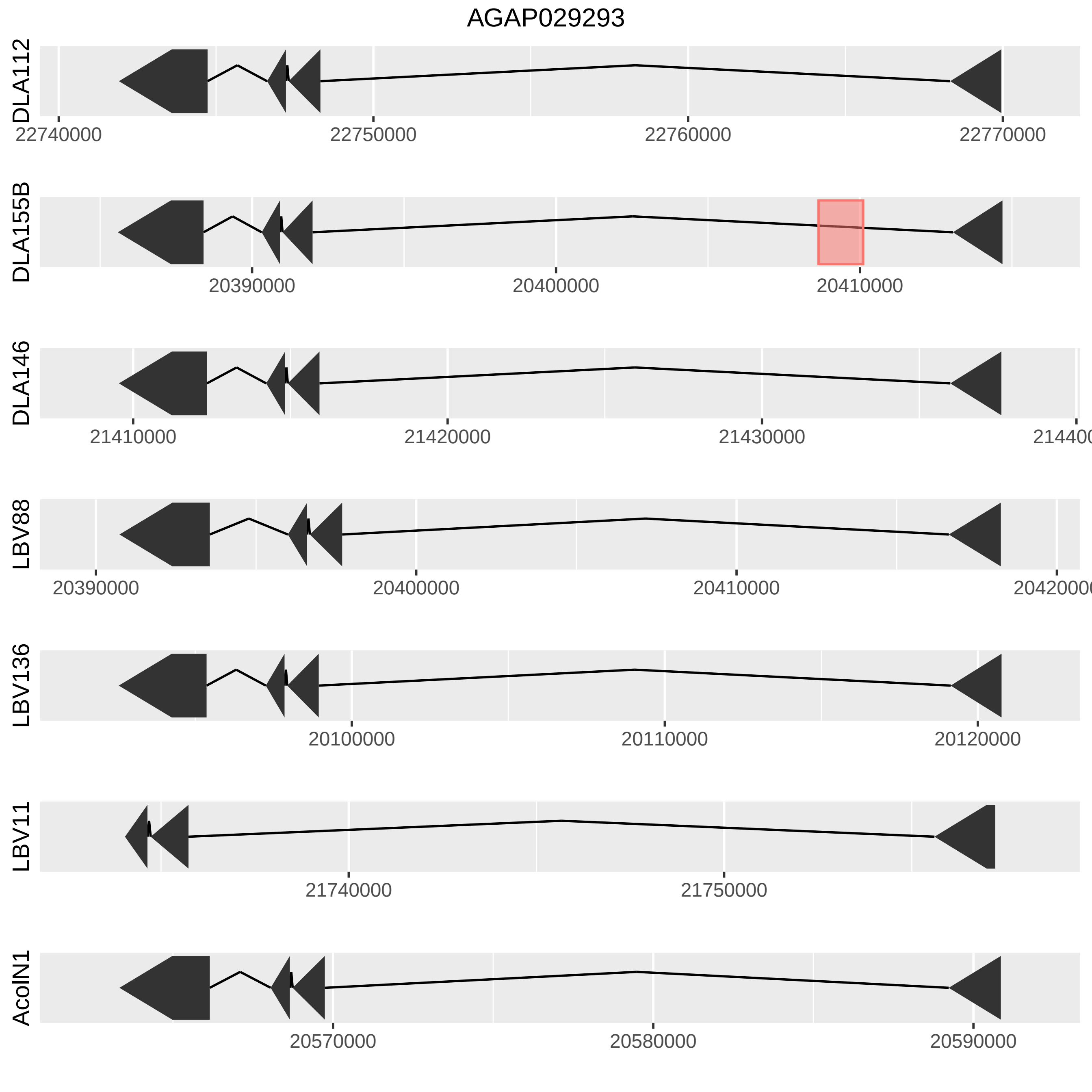

AGAP029293
DLA112
22740000
22750000
22760000
22770000
DLA155B
20390000
20400000
20410000
DLA146
21410000
21420000
21430000
LBV88
20390000
20400000
20410000
LBV136
20100000
20110000
20120000
LBV11
21740000
21750000
AcoIN1
20570000
20580000
20590000

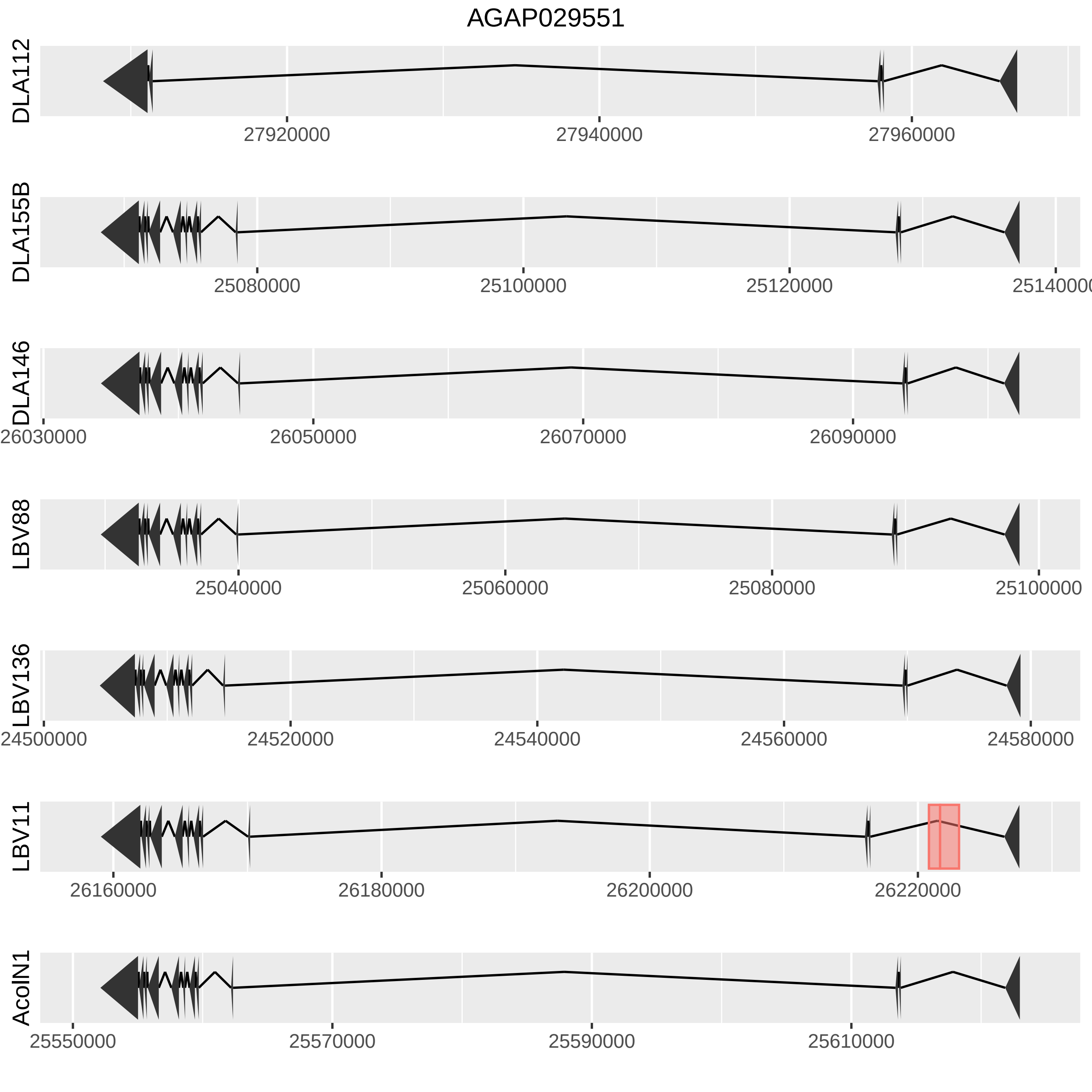

AGAP029551
DLA112
27920000
27940000
27960000
DLA155B
25080000
25100000
25120000
25140000
DLA146
26030000
26050000
26070000
26090000
LBV88
25040000
25060000
25080000
25100000
LBV136
24500000
24520000
24540000
24560000
24580000
LBV11
26160000
26180000
26200000
26220000
AcolN1
25550000
25570000
25590000
25610000

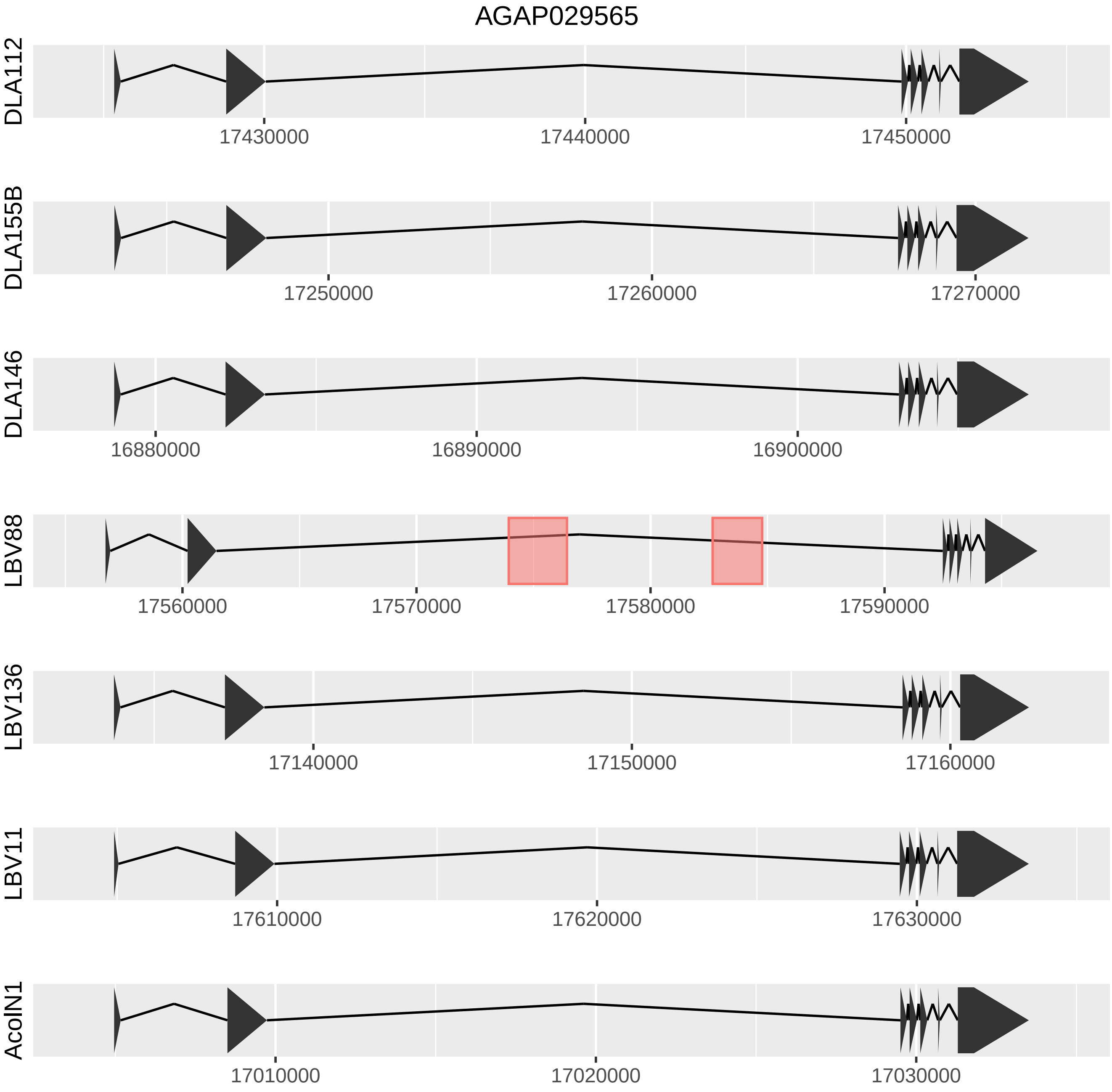

AGAP029565
DLA112
17430000
17440000
17450000
DLA155B
17250000
17260000
17270000
DLA146
16880000
16890000
16900000
LBV88
17560000
17570000
17580000
17590000
LBV136
17140000
17150000
17160000
LBV11
17610000
17620000
17630000
AcolN1
17010000
17020000
17030000

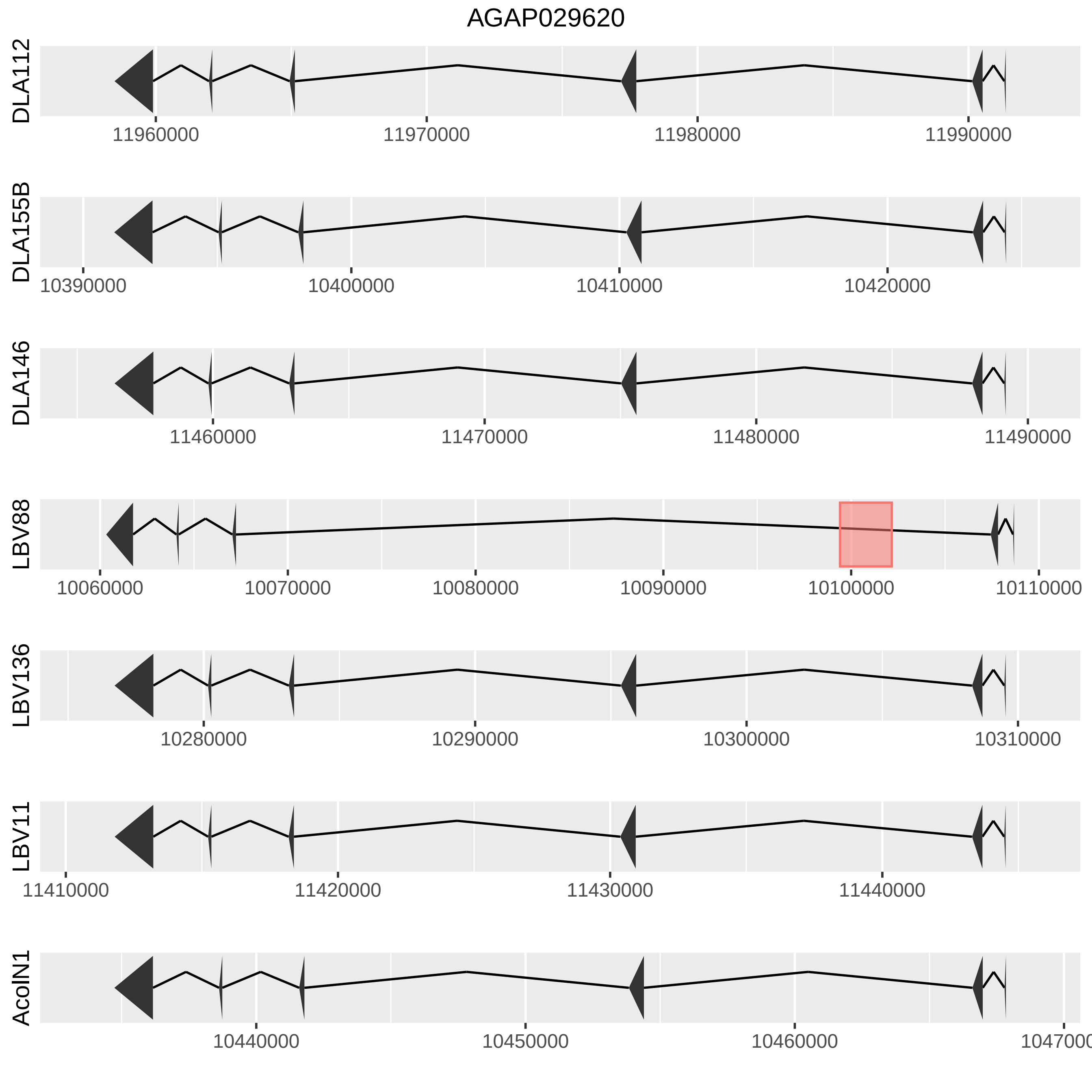
AGAP029620
DLA112
11960000
11970000
11980000
11990000
DLA155B
10390000
10400000
10410000
10420000
DLA146
11460000
11470000
11480000
11490000
LBV88
10060000
10070000
10080000
10090000
10100000
10110000
LBV136
10280000
10290000
10300000
10310000
LBV11
11410000
11420000
11430000
11440000
AcoIN1
10440000
10450000
10460000
