## Additional File 7 for "Uncovering transposable element variants and their potential adaptive impact in urban populations of the malaria vector *Anopheles coluzzii*"

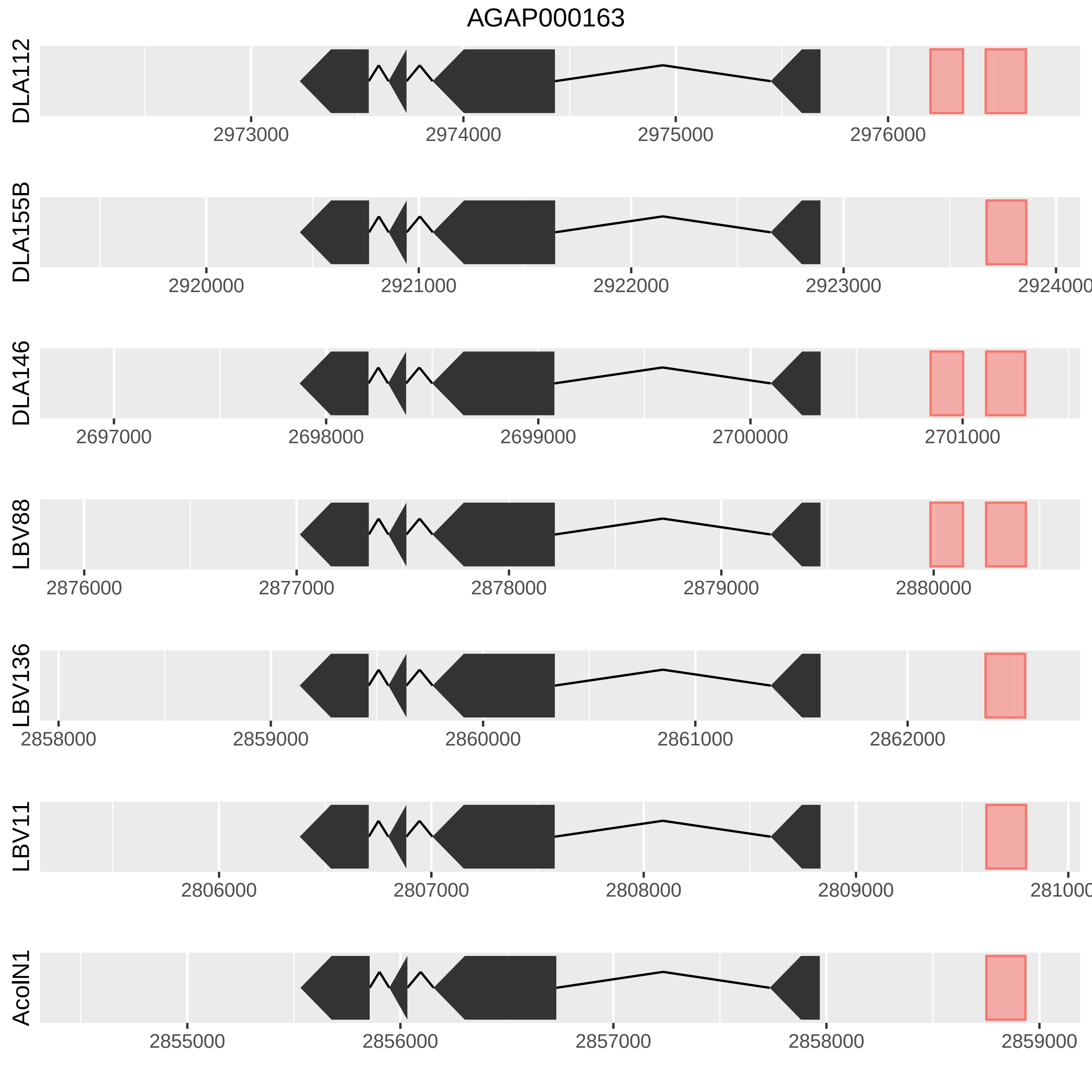
AGAP000163
DLA112
2973000
2974000
2975000
2976000
DLA155B
2920000
2921000
2922000
2923000
2924000
DLA146
2697000
2698000
2699000
2700000
2701000
LBV88
2876000
2877000
2878000
2879000
2880000
LBV136
2858000
2859000
2860000
2861000
2862000
LBV11
2806000
2807000
2808000
2809000
AcoIN1
2855000
2856000
2857000
2858000
2859000

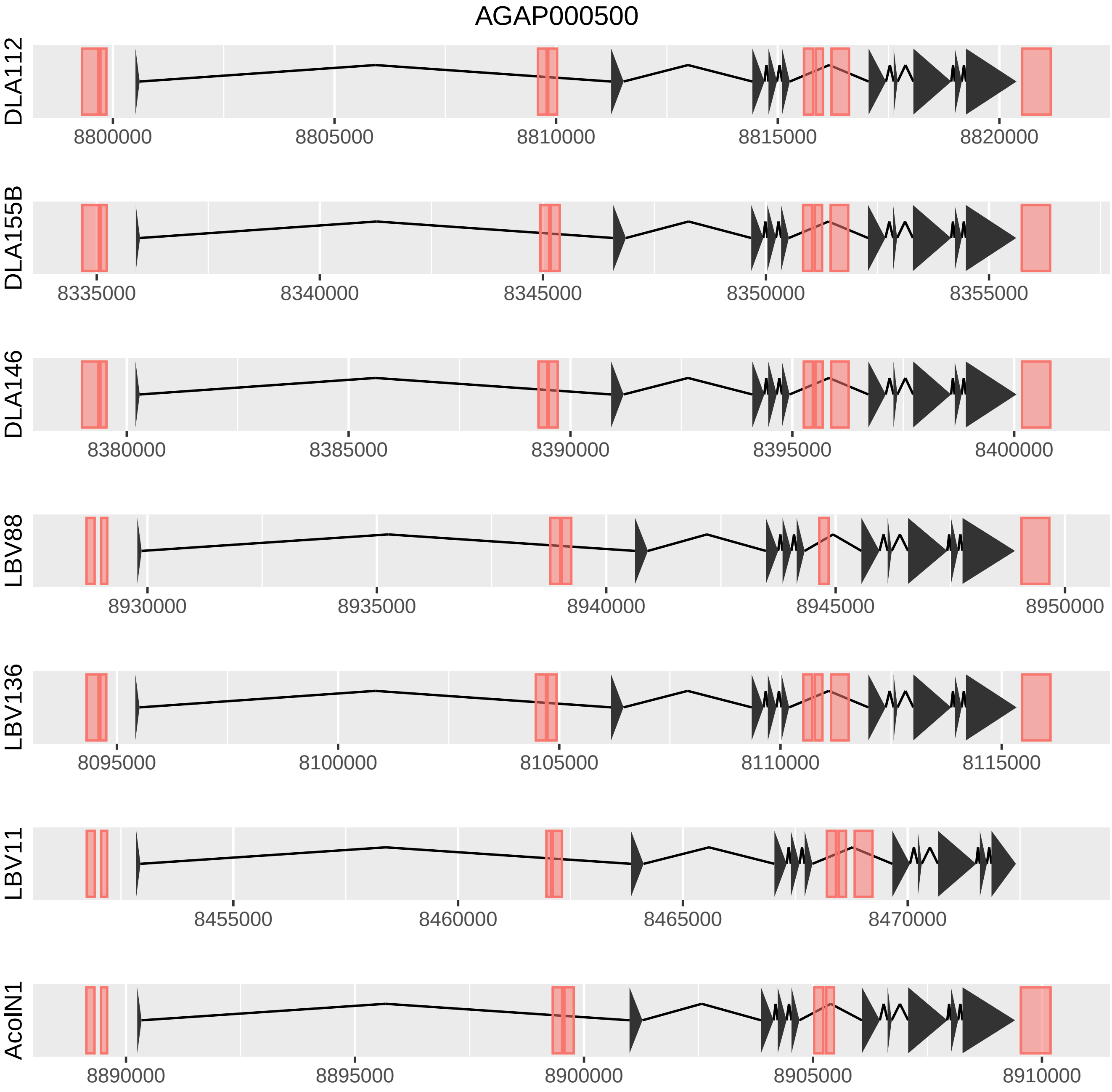
AGAP000500
DLA112
8800000
8805000
8810000
8815000
8820000
DLA155B
8335000
8340000
8345000
8350000
8355000
DLA146
8380000
8385000
8390000
8395000
8400000
LBV88
8930000
8935000
8940000
8945000
8950000
LBV136
8095000
8100000
8105000
8110000
8115000
LBV11
8455000
8460000
8465000
8470000
AcolN1
8890000
8895000
8900000
8905000
8910000

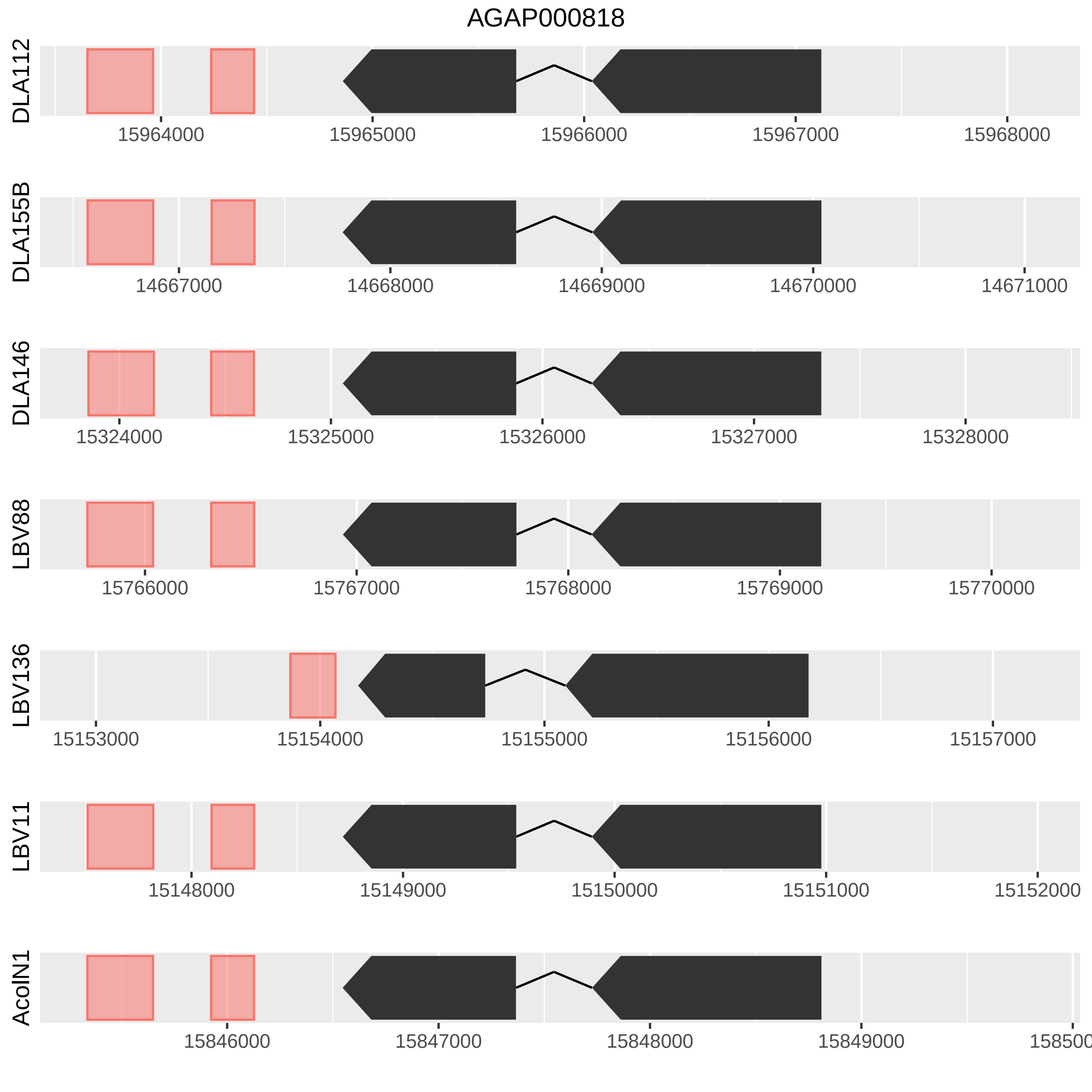
AGAP000818
DLA112
15964000
15965000
15966000
15967000
15968000
DLA155B
14667000
14668000
14669000
14670000
14671000
DLA146
15324000
15325000
15326000
15327000
15328000
LBV88
15766000
15767000
15768000
15769000
15770000
LBV136
15153000
15154000
15155000
15156000
15157000
LBV11
15148000
15149000
15150000
15151000
15152000
AcolN1
15846000
15847000
15848000
15849000

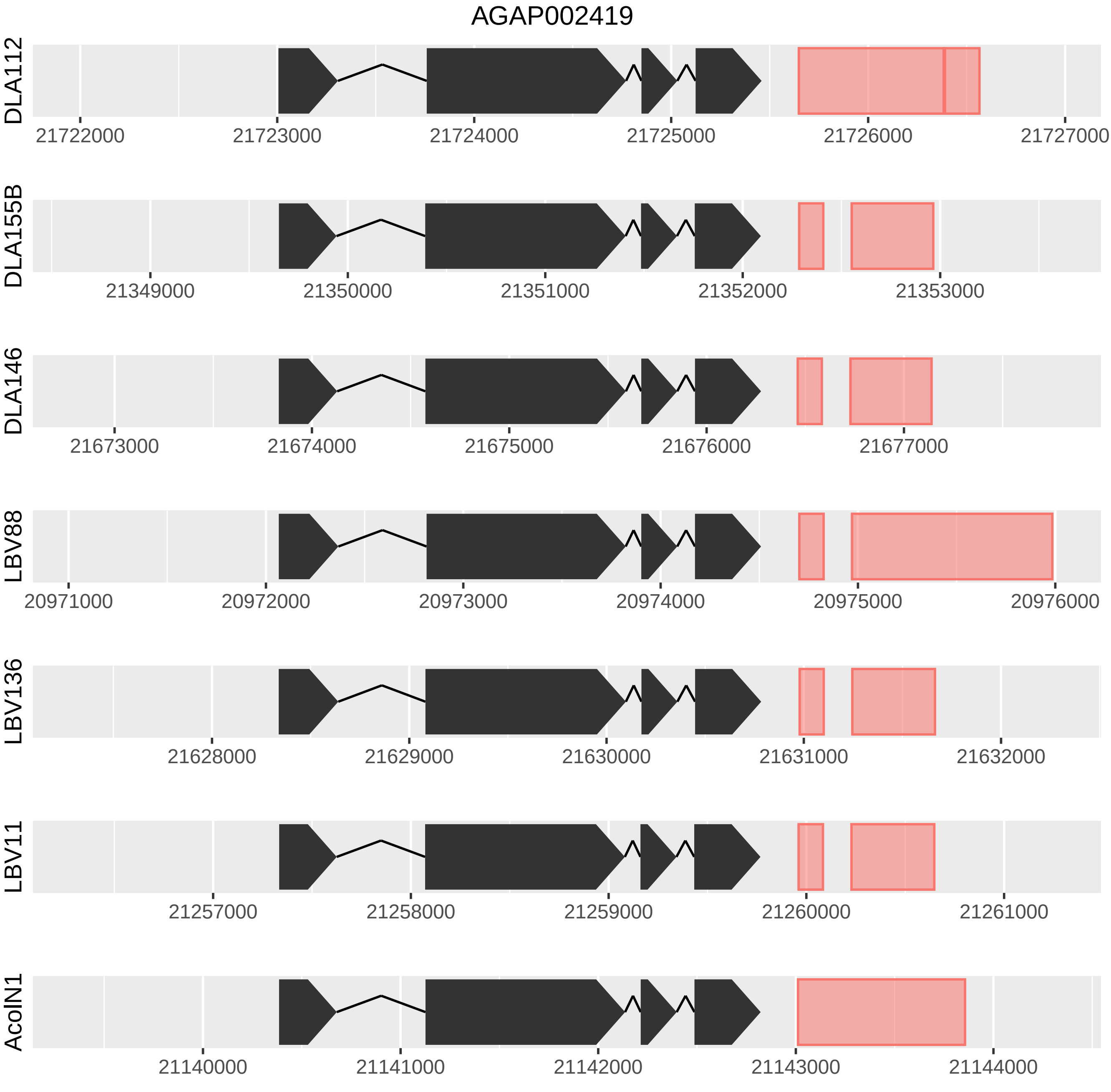
AGAP002419
DLA112
21722000
21723000
21724000
21725000
21726000
21727000
DLA155B
21349000
21350000
21351000
21352000
21353000
DLA146
21673000
21674000
21675000
21676000
21677000
LBV88
20971000
20972000
20973000
20974000
20975000
20976000
LBV136
21628000
21629000
21630000
21631000
21632000
LBV11
21257000
21258000
21259000
21260000
21261000
AcolN1
21140000
21141000
21142000
21143000
21144000

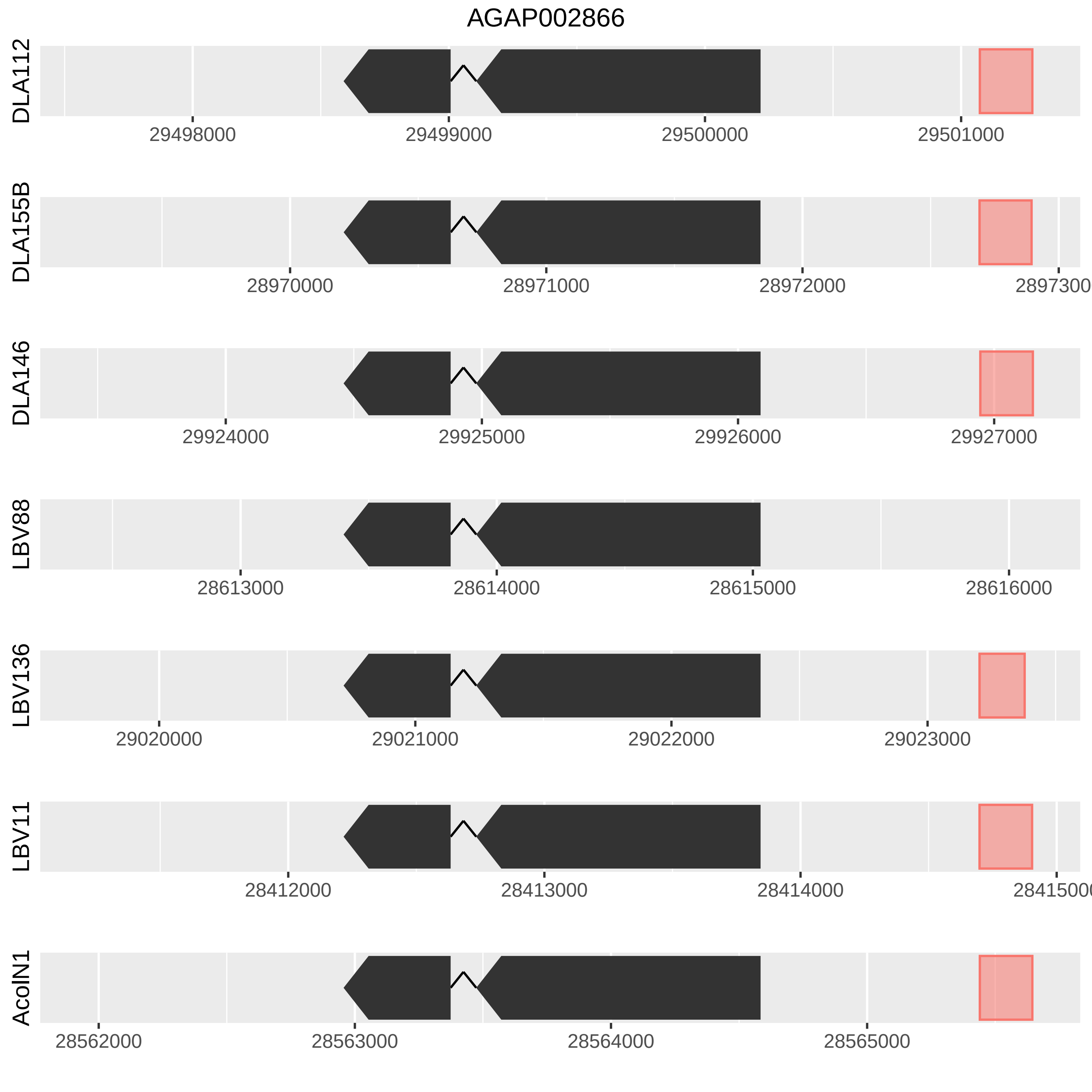

AGAP002866
DLA112
29498000
29499000
29500000
29501000
DLA155B
28970000
28971000
28972000
2897300
DLA146
29924000
29925000
29926000
29927000
LBV88
28613000
28614000
28615000
28616000
LBV136
29020000
29021000
29022000
29023000
LBV11
28412000
28413000
28414000
2841500
AcolN1
28562000
28563000
28564000
28565000

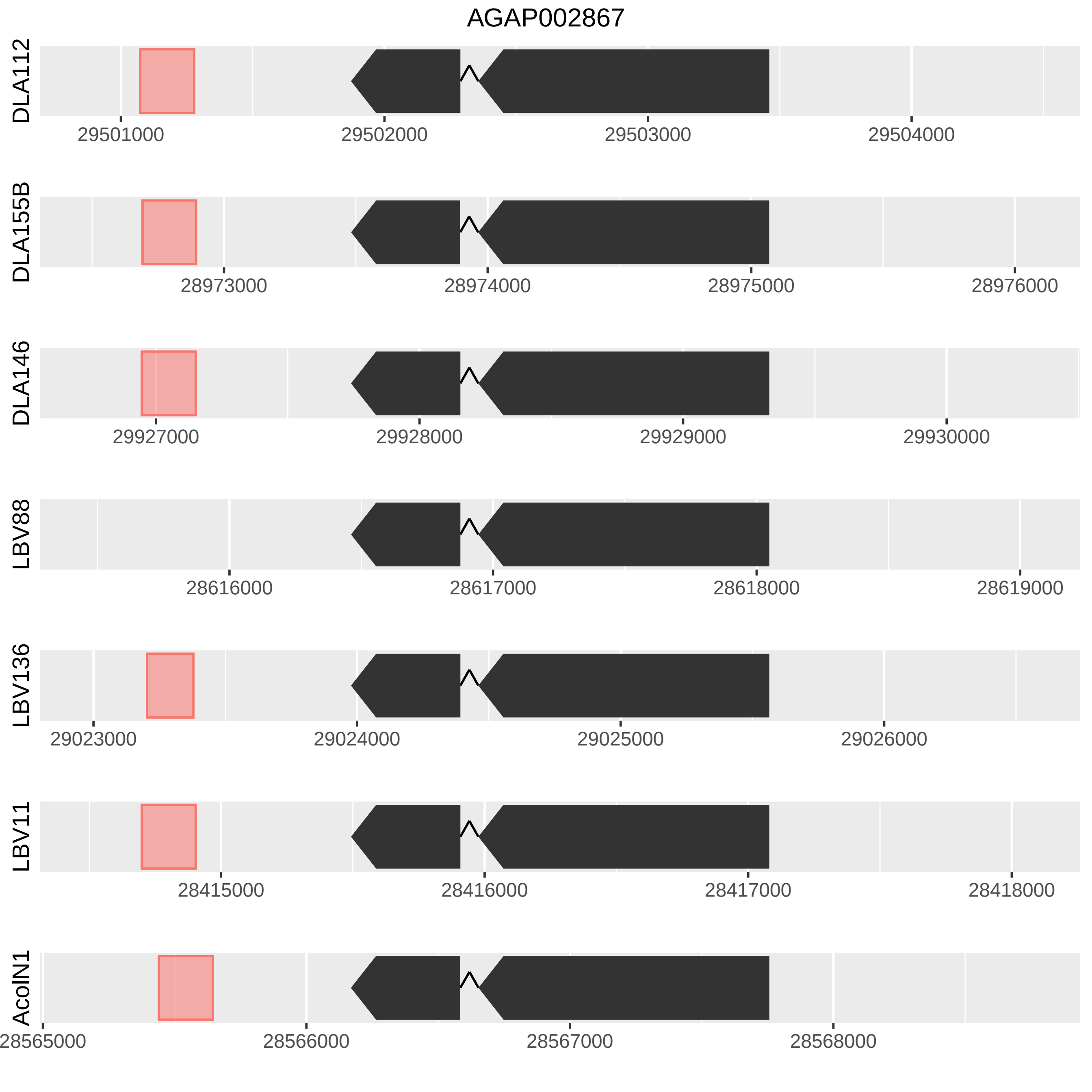
AGAP002867
DLA112
29501000
29502000
29503000
29504000
DLA155B
28973000
28974000
28975000
28976000
DLA146
29927000
29928000
29929000
29930000
LBV88
28616000
28617000
28618000
28619000
LBV136
29023000
29024000
29025000
29026000
LBV11
28415000
28416000
28417000
28418000
AcolN1
28565000
28566000
28567000
28568000

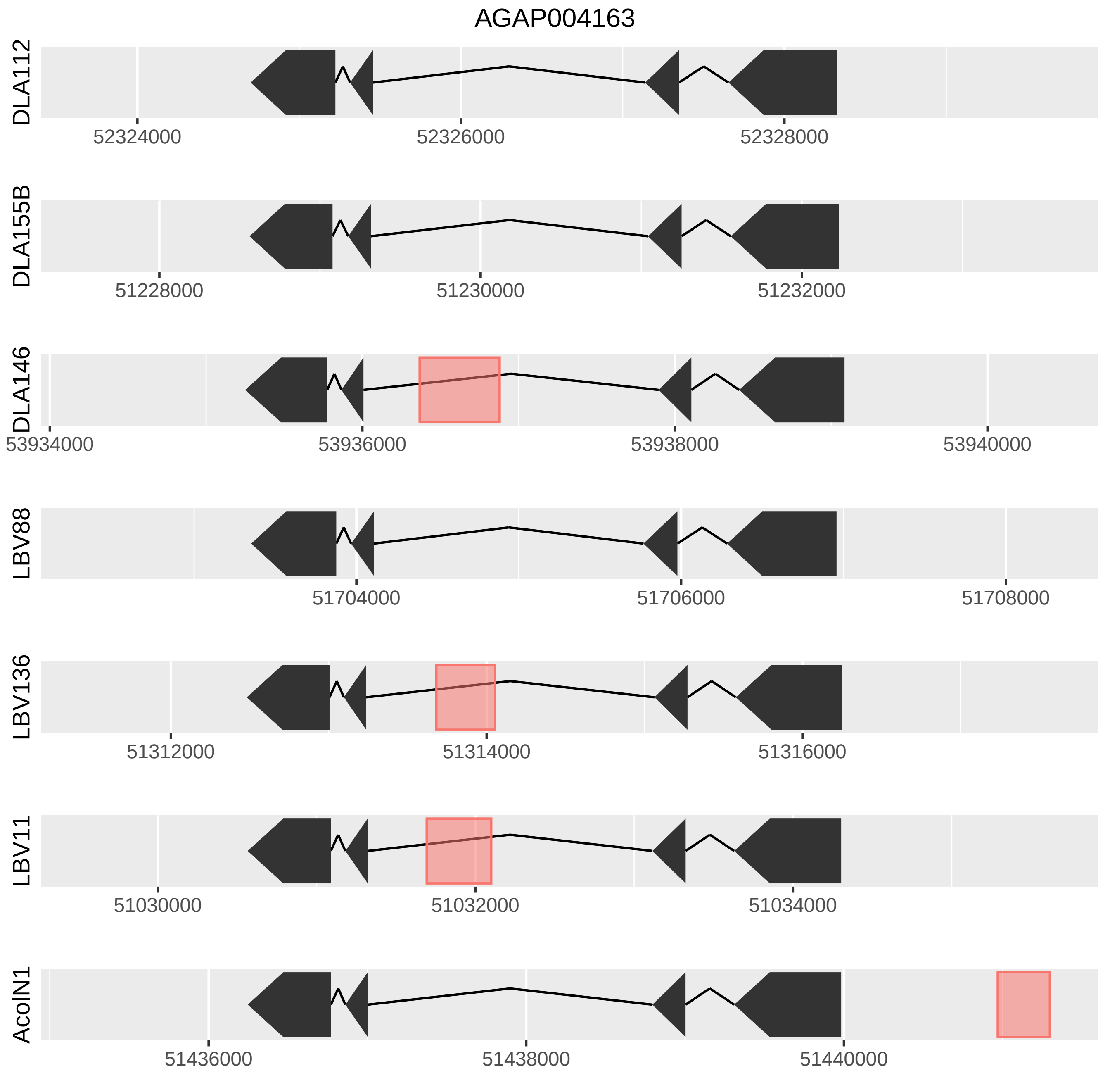
AGAP004163
DLA112
52324000
52326000
52328000
DLA155B
51228000
51230000
51232000
DLA146
53934000
53936000
53938000
53940000
LBV88
51704000
51706000
51708000
LBV136
51312000
51314000
51316000
LBV11
51030000
51032000
51034000
AcolN1
51436000
51438000
51440000

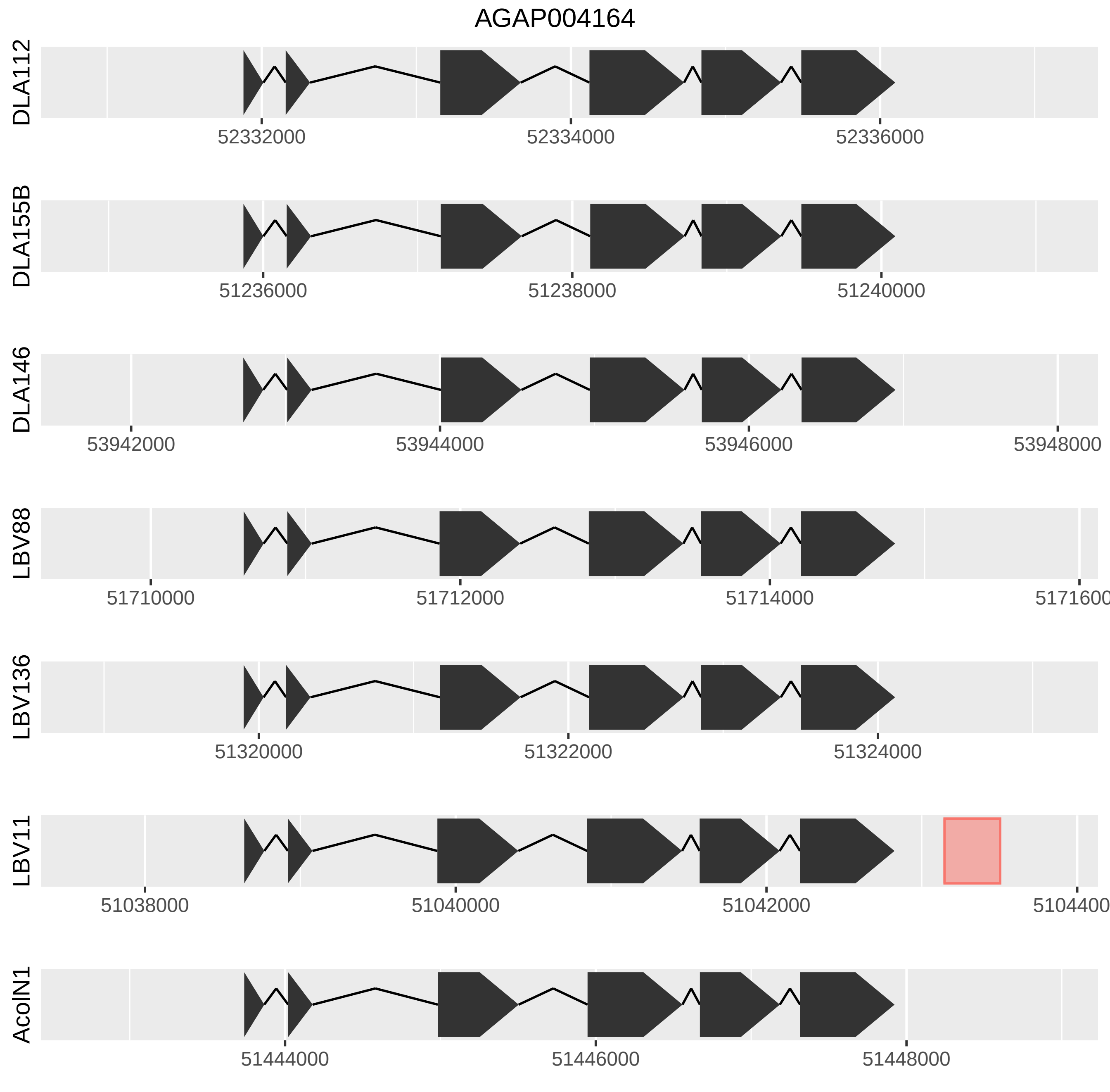
AGAP004164
DLA112
52332000
52334000
52336000
DLA155B
51236000
51238000
51240000
DLA146
53942000
53944000
53946000
53948000
LBV88
51710000
51712000
51714000
5171600
LBV136
51320000
51322000
51324000
LBV11
51038000
51040000
51042000
5104400
AcolN1
51444000
51446000
51448000

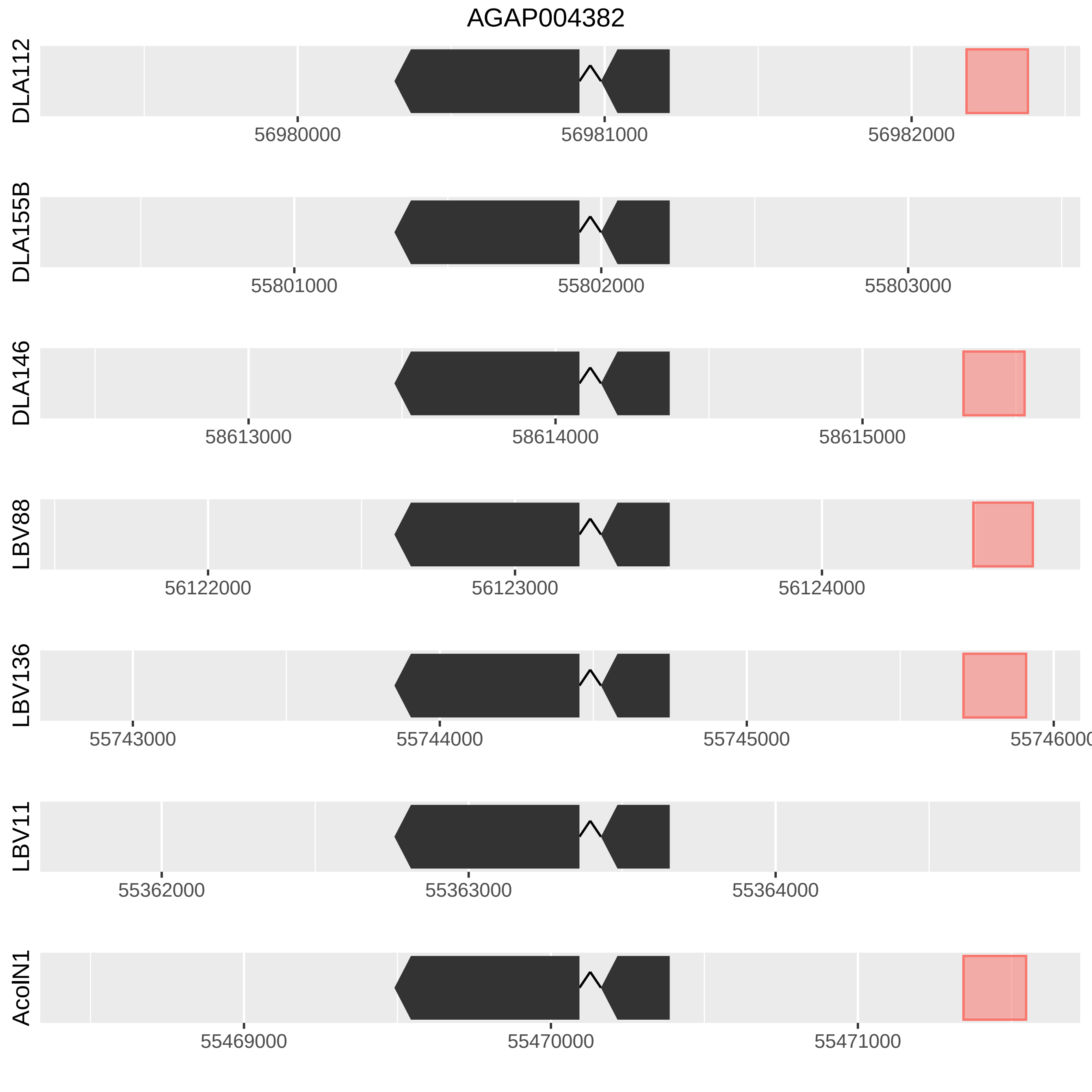
AGAP004382
DLA112
56980000
56981000
56982000
DLA155B
55801000
55802000
55803000
DLA146
58613000
58614000
58615000
LBV88
56122000
56123000
56124000
LBV136
55743000
55744000
55745000
55746000
LBV11
55362000
55363000
55364000
AcoIN1
55469000
55470000
55471000

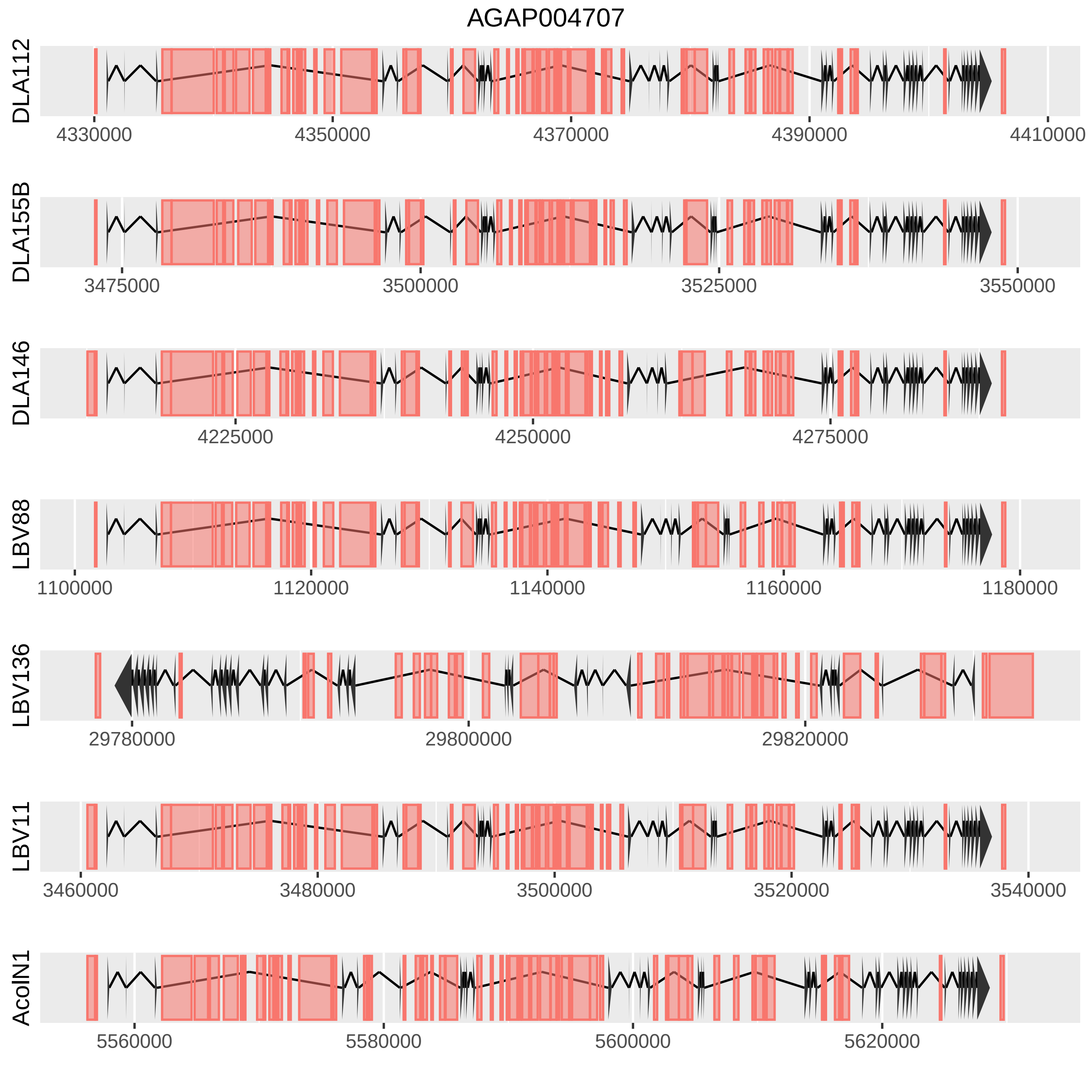
AGAP004707
DLA112
4330000
4350000
4370000
4390000
4410000
DLA155B
3475000
3500000
3525000
3550000
DLA146
4225000
4250000
4275000
LBV88
1100000
1120000
1140000
1160000
1180000
LBV136
29780000
29800000
29820000
LBV11
3460000
3480000
3500000
3520000
3540000
AcolN1
5560000
5580000
5600000
5620000

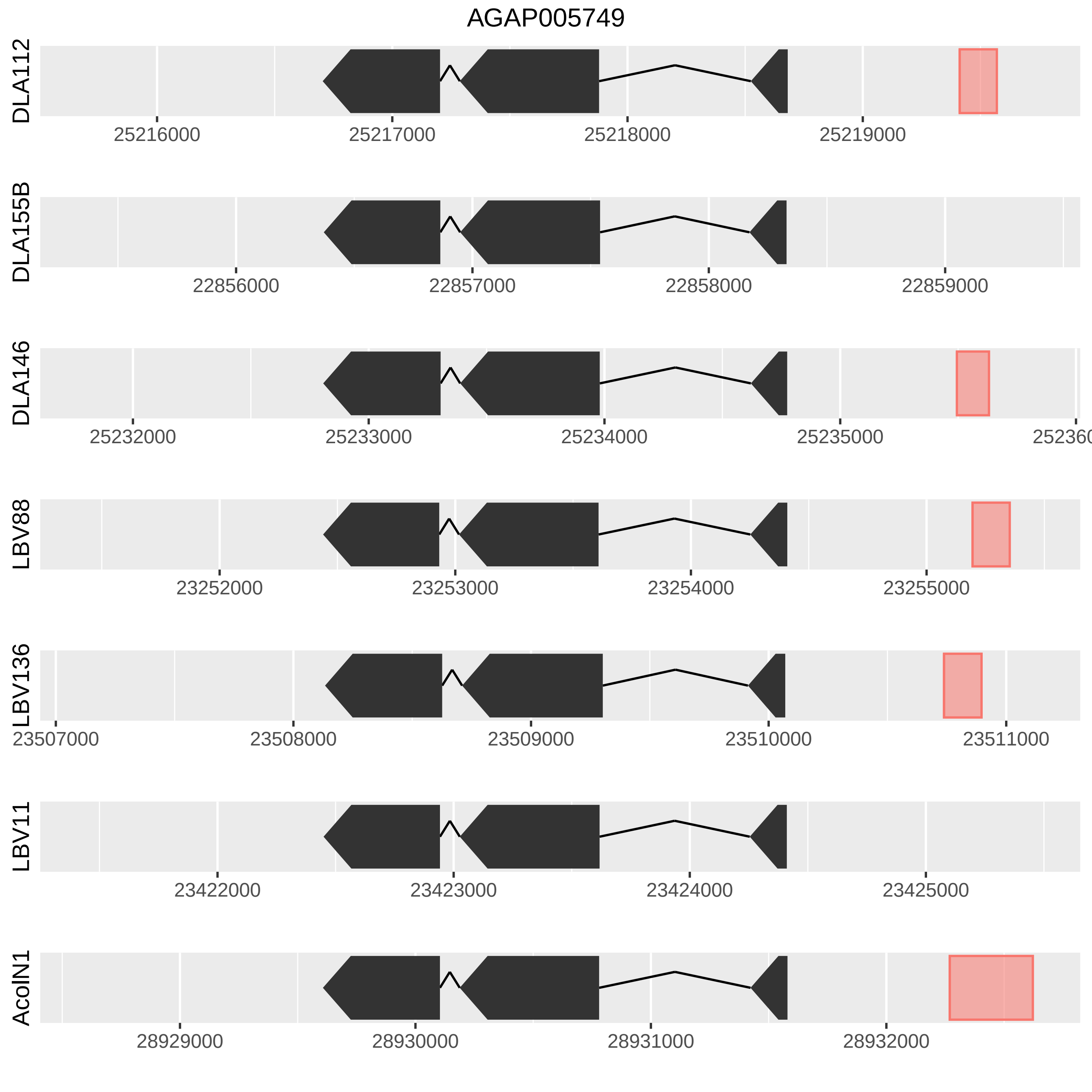
AGAP005749
DLA112
25216000
25217000
25218000
25219000
DLA155B
22856000
22857000
22858000
22859000
DLA146
25232000
25233000
25234000
25235000
DLA138
23252000
23253000
23254000
23255000
LBV136
23507000
23508000
23509000
23510000
23511000
LBV11
23422000
23423000
23424000
23425000
AcoIN1
28929000
28930000
28931000
28932000

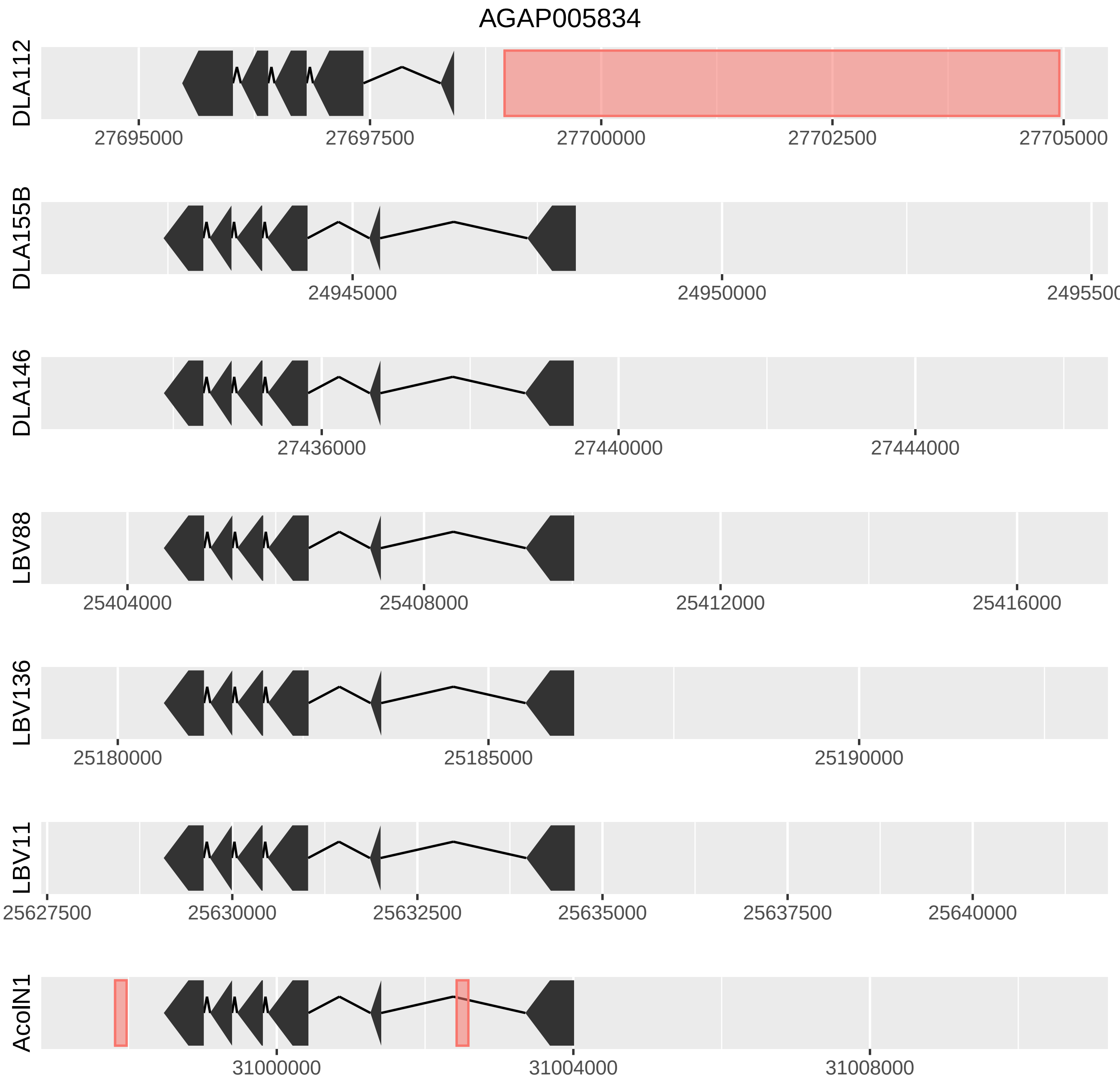

AGAP005834
DLA112
27695000
27697500
27700000
27702500
27705000
DLA155B
24945000
24950000
249550
DLA146
27436000
27440000
27444000
LBV88
25404000
25408000
25412000
25416000
LBV136
25180000
25185000
25190000
LBV11
25627500
25630000
25632500
25635000
25637500
25640000
AcolN1
31000000
31004000
31008000

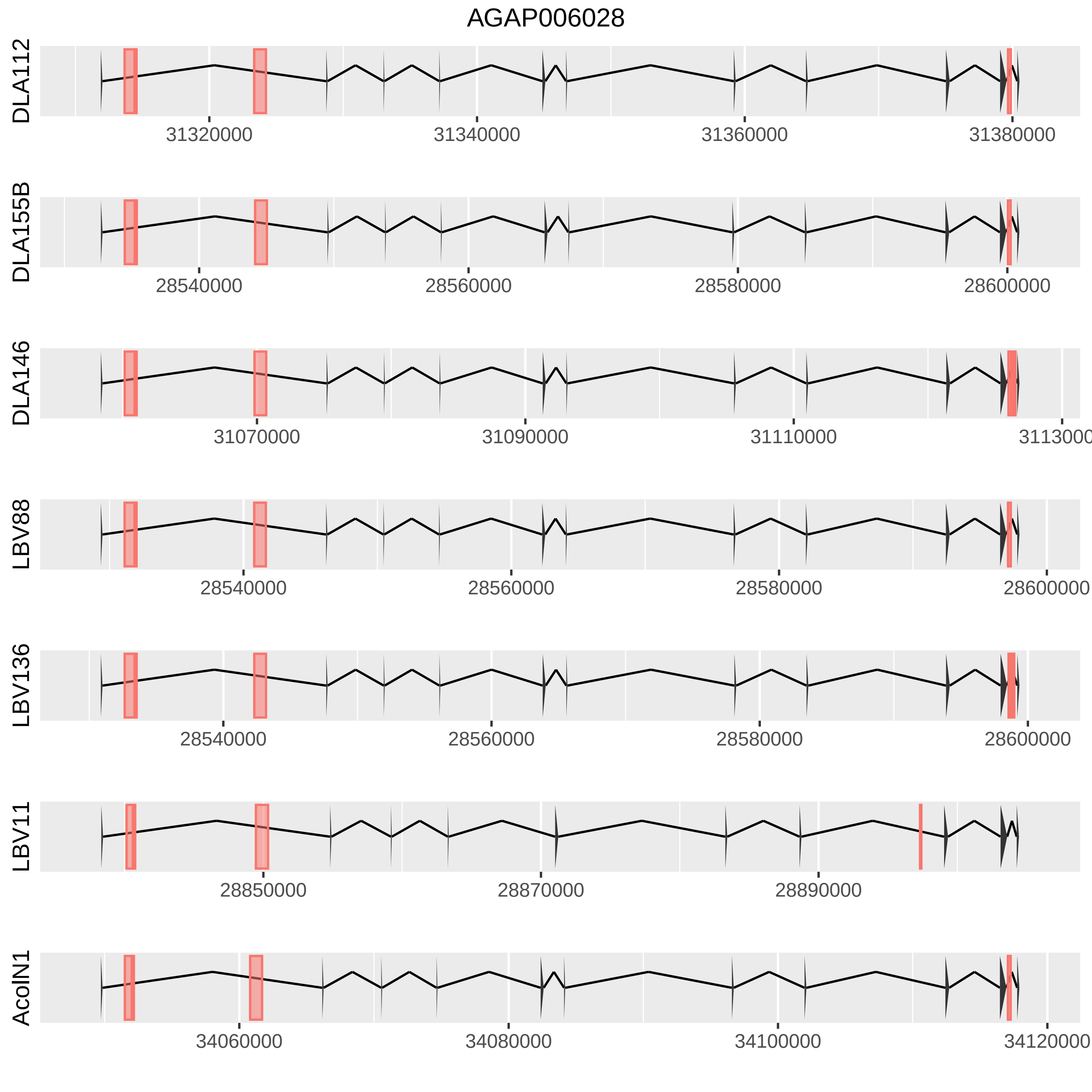

AGAP006028
DLA112
31320000
31340000
31360000
31380000
DLA155B
28540000
28560000
28580000
28600000
DLA146
31070000
31090000
31110000
LBV88
28540000
28560000
28580000
28600000
LBV136
28540000
28560000
28580000
28600000
LBV11
28850000
28870000
28890000
AcolN1
34060000
34080000
34100000
34120000

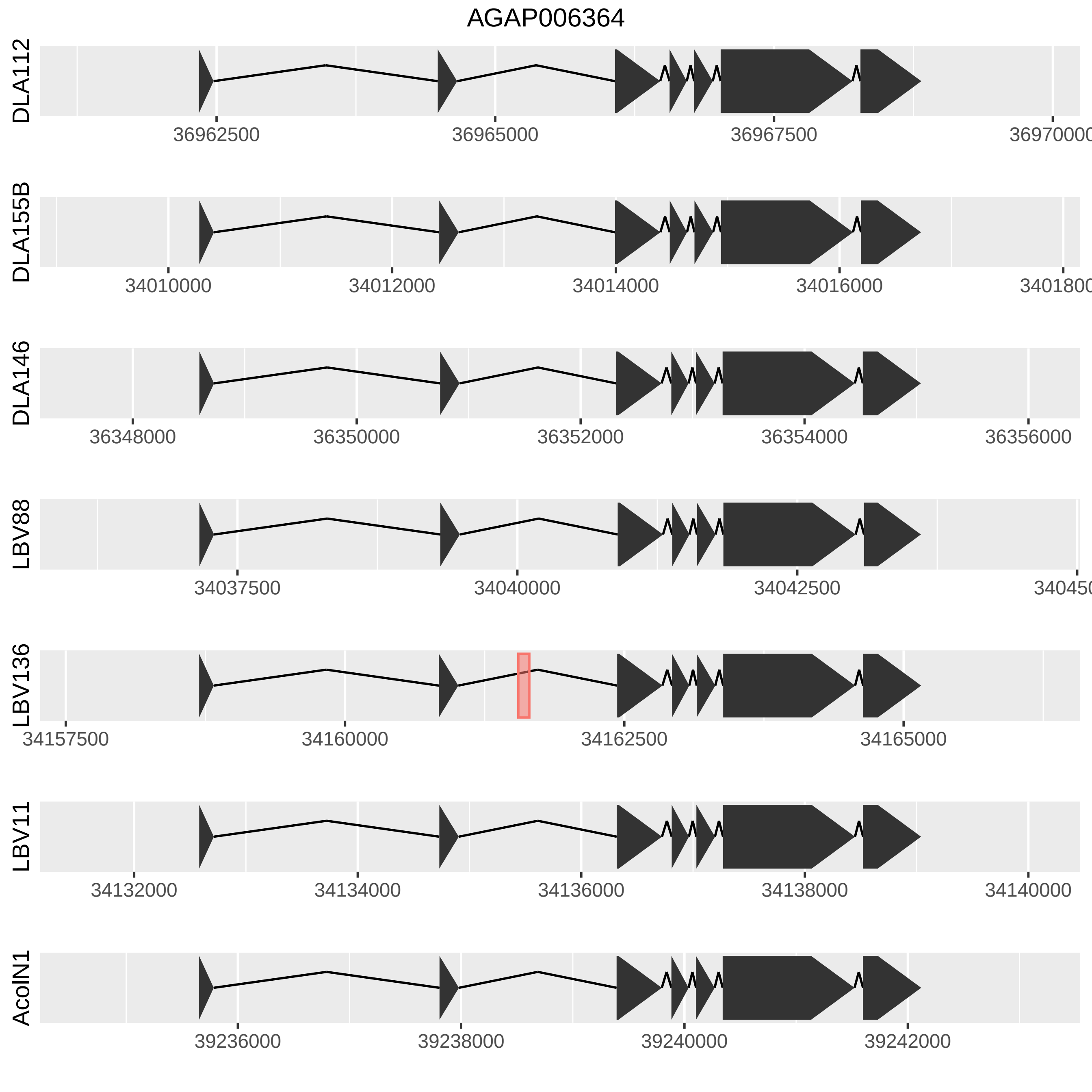
AGAP006364
DLA112
36962500
36965000
36967500
DLA155B
34010000
34012000
34014000
34016000
DLA146
36348000
36350000
36352000
36354000
36356000
LBV88
34037500
34040000
34042500
LBV136
34157500
34160000
34162500
34165000
LBV11
34132000
34134000
34136000
34138000
34140000
AcolN1
39236000
39238000
39240000
39242000

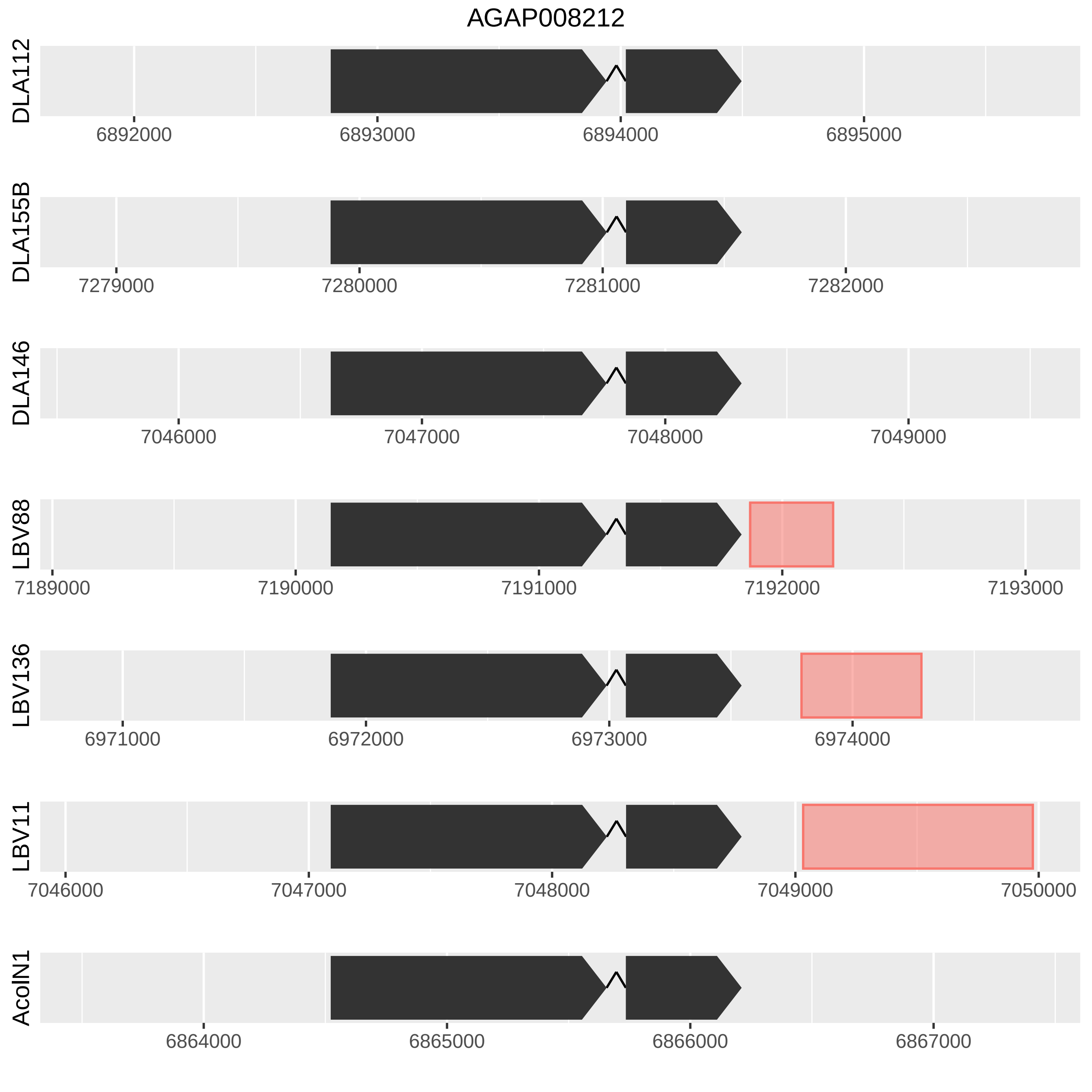
AGAP008212
DLA112
6892000
6893000
6894000
6895000
DLA155B
7279000
7280000
7281000
7282000
DLA146
7046000
7047000
7048000
7049000
LBV88
7189000
7190000
7191000
7192000
7193000
LBV136
6971000
6972000
6973000
6974000
LBV11
7046000
7047000
7048000
7049000
7050000
AcoIN1
6864000
6865000
6866000
6867000

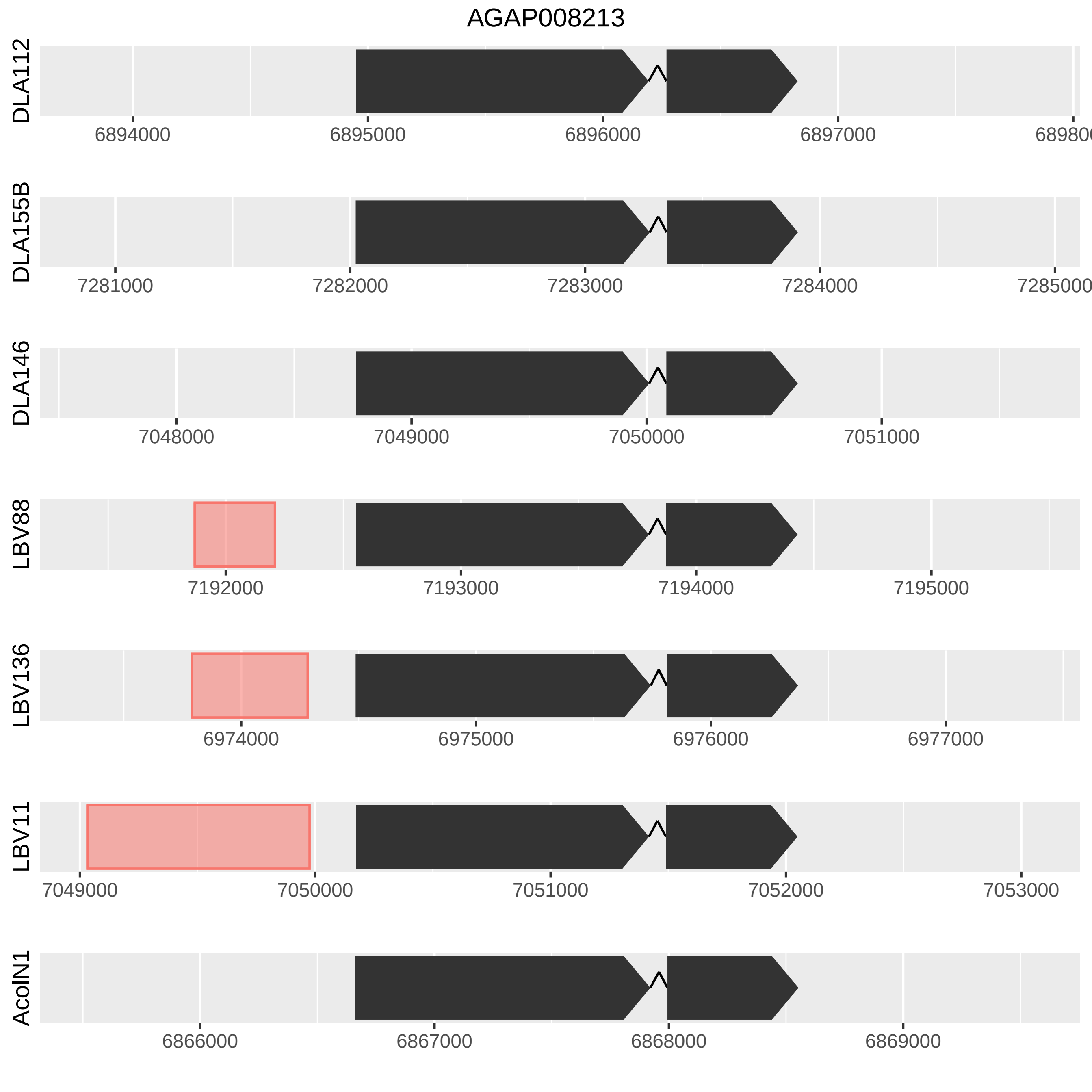

AGAP008213
DLA112
6894000
6895000
6896000
6897000
DLA155B
7281000
7282000
7283000
7284000
DLA146
7048000
7049000
7050000
7051000
LBV88
7192000
7193000
7194000
7195000
LBV136
6974000
6975000
6976000
6977000
LBV11
7049000
7050000
7051000
7052000
7053000
AcoIN1
6866000
6867000
6868000
6869000

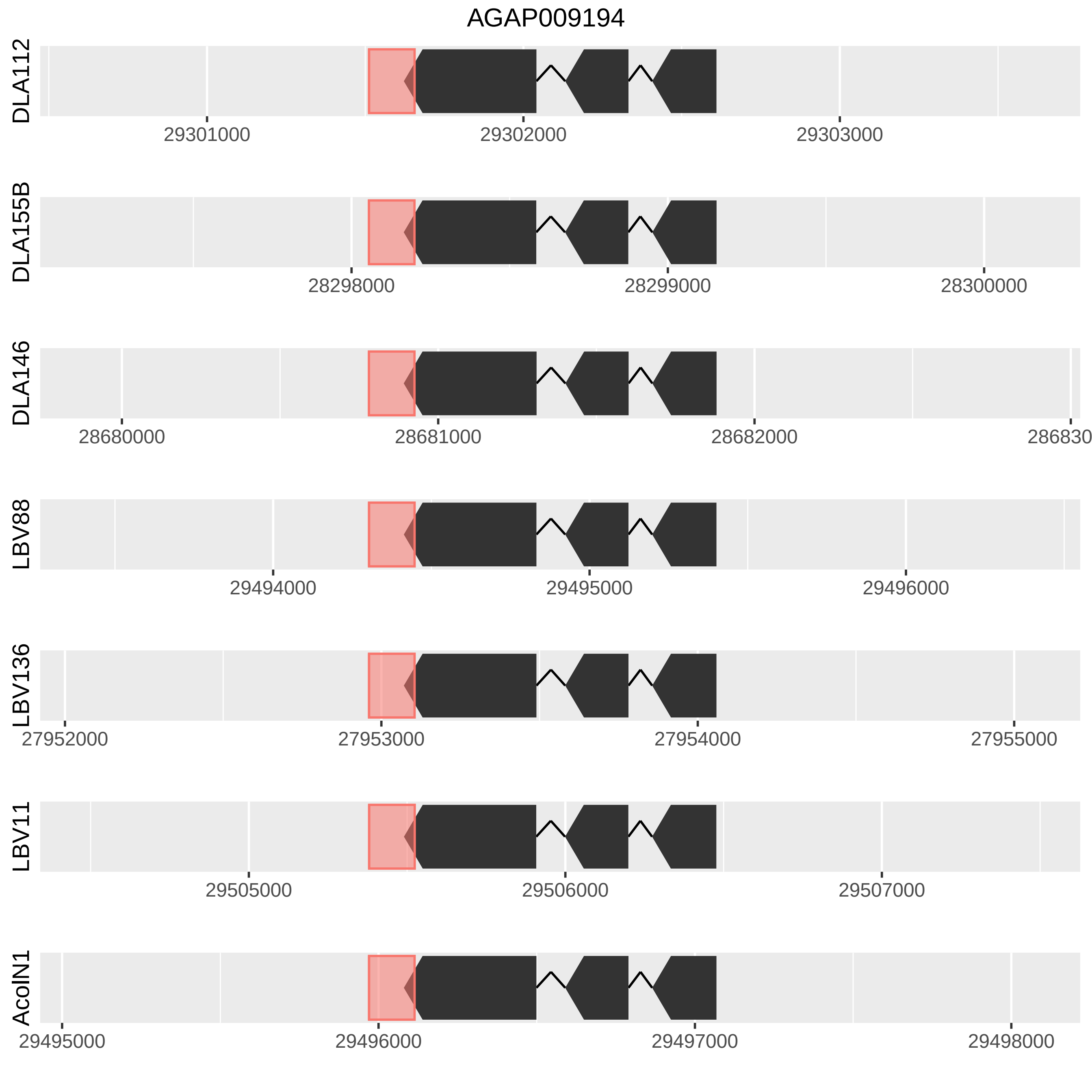

AGAP009194
DLA112
29301000
29302000
29303000
DLA155B
28298000
28299000
28300000
DLA146
28680000
28681000
28682000
LBV88
29494000
29495000
29496000
LBV136
27952000
27953000
27954000
27955000
LBV11
29505000
29506000
29507000
AcolN1
29495000
29496000
29497000
29498000

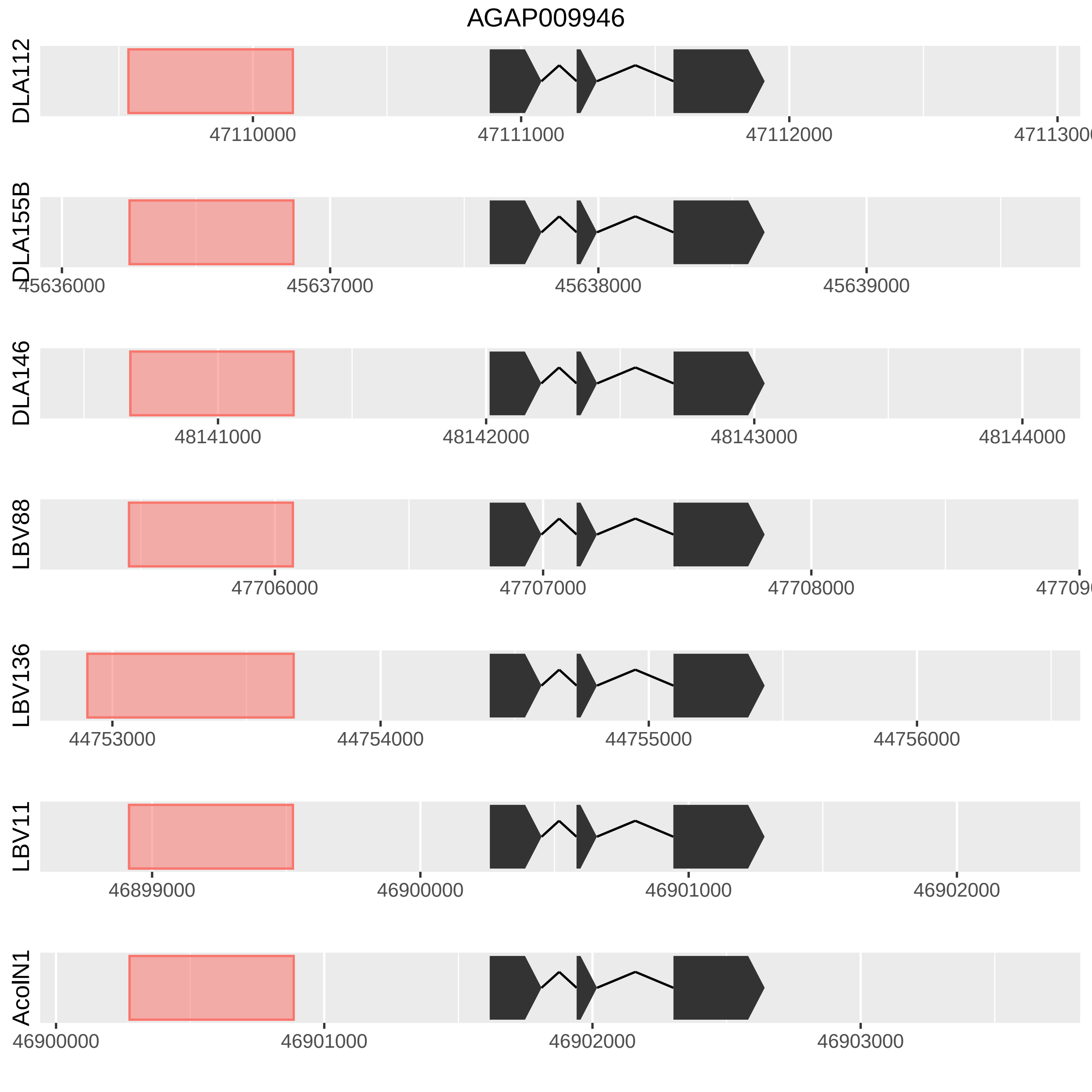
AGAP009946
DLA112
47110000
47111000
47112000
DLA155B
45636000
45637000
45638000
45639000
DLA146
48141000
48142000
48143000
48144000
LBV88
47706000
47707000
47708000
LBV136
44753000
44754000
44755000
44756000
LBV11
46899000
46900000
46901000
46902000
AcolN1
46900000
46901000
46902000
46903000

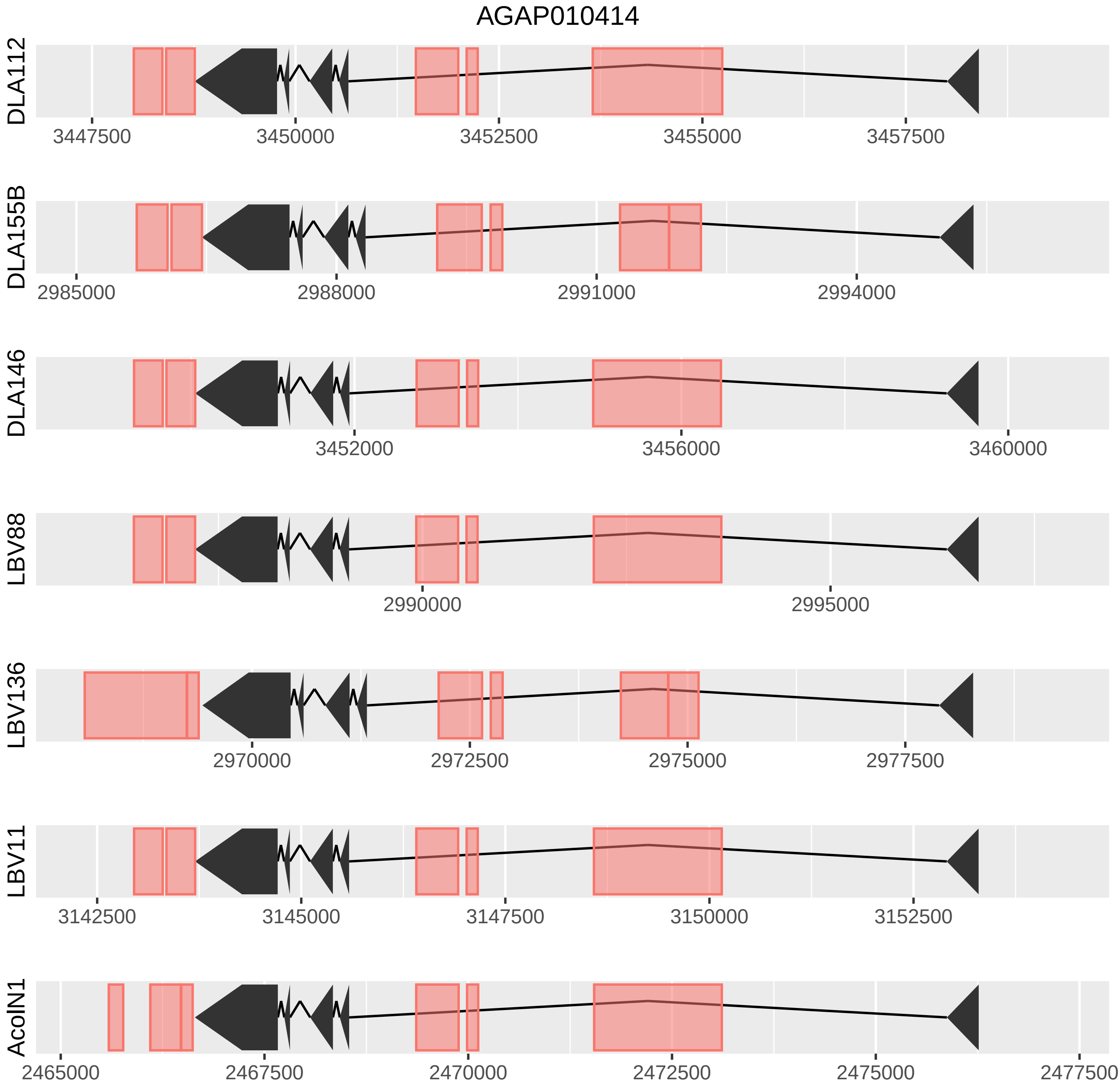
AGAP010414
DLA112
3447500
3450000
3452500
3455000
3457500
DLA155B
2985000
2988000
2991000
2994000
DLA146
3452000
3456000
3460000
LBV88
2990000
2995000
LBV136
2970000
2972500
2975000
2977500
LBV11
3142500
3145000
3147500
3150000
3152500
AcolN1
2465000
2467500
2470000
2472500
2475000
2477500

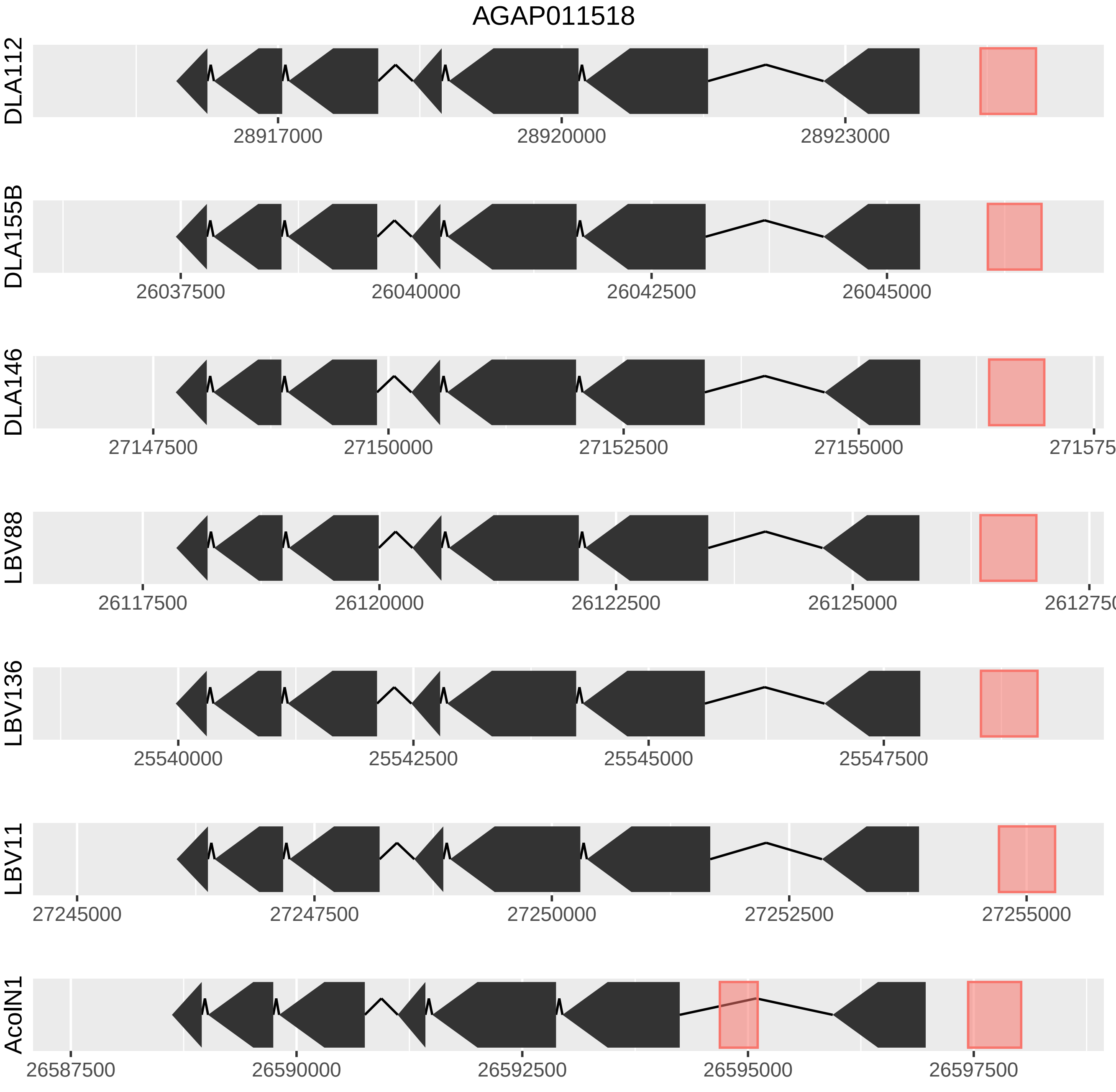

AGAP011518
DLA112
28917000
28920000
28923000
DLA155B
26037500
26040000
26042500
26045000
DLA146
27147500
27150000
27152500
27155000
LBV88
26117500
26120000
26122500
26125000
LBV136
25540000
25542500
25545000
25547500
LBV11
27245000
27247500
27250000
27252500
27255000
AcolN1
26587500
26590000
26592500
26595000
26597500

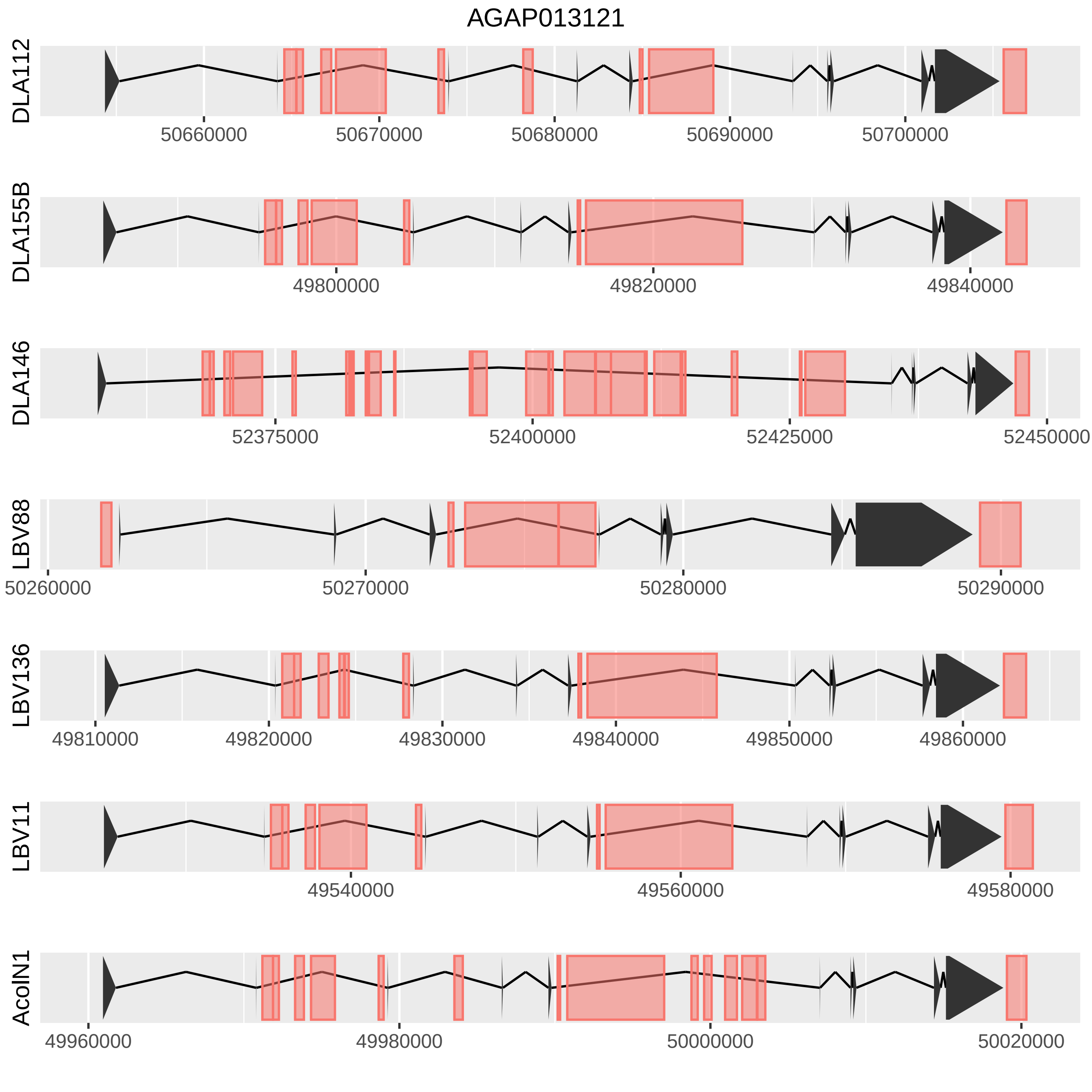

AGAP013121
DLA112
50660000
50670000
50680000
50690000
50700000
DLA155B
49800000
49820000
49840000
DLA146
52375000
52400000
52425000
52450000
LBV88
50260000
50270000
50280000
50290000
LBV136
49810000
49820000
49830000
49840000
49850000
49860000
LBV11
49540000
49560000
49580000
AcolN1
49960000
49980000
50000000
50020000
